# Sex Solves Haldane’s Dilemma

**DOI:** 10.1101/582536

**Authors:** Donal A. Hickey, G. Brian Golding

## Abstract

The cumulative reproductive cost of multi-locus selection has been seen as a potentially limiting factor on the rate of adaptive evolution. In this paper, we show that Haldane’s arguments for the accumulation of reproductive costs over multiple loci are valid only for a clonally reproducing population of asexual genotypes. We show that a sexually reproducing population avoids this accumulation of costs. Thus, sex removes a perceived reproductive constraint on the rate of adaptive evolution. The significance of our results is twofold. First, the results demonstrate that adaptation based on multiple genes – such as selection acting on the standing genetic variation - does not entail a huge reproductive cost as suggested by Haldane, provided of course that the population is reproducing sexually. Secondly, this reduction in the cost of natural selection provides a simple biological explanation for the advantage of sex. Specifically, Haldane’s calculations illustrate the evolutionary disadvantage of asexuality; sexual reproduction frees the population from this disadvantage.

## Introduction

Adaptive evolution depends on reproductive excess because disfavored alleles must be culled in order for favored alleles to increase in frequency within the selected population. Such culling, in the absence of a sufficient reproductive excess, would result in a decrease in population size and, eventually, it could lead to the extinction of the population. Haldane (1957; 1960) referred to the culling of disfavored types as the “cost of natural selection”. In addition to defining this cost, Haldane estimated how great this cost might be during the substitution of one allele for another at a single locus. He then argued that, given independent selection at different genetic loci, this cost would be approximately additive over loci (Haldane 1957; 1960). In other words, if the cost of allelic substitution at one locus equals x, then the cost for L loci would equal Lx. This would mean that evolutionary adaptations that involved allelic substitutions at many genetic loci could prove to be “too costly”, given the available reproductive excess. Thus, simultaneous selection at many loci might lead to extinction of the population or, at best, it would constrain the rate of adaptive change. This problem has been referred to as Haldane’s dilemma (VanValen 1963).

Haldane’s calculations of the cost of natural selection had a major impact on the field of evolutionary biology. For example, Dodson (1962) wondered if the cost were too high to allow the adaptive divergence between humans and chimpanzees, while Kimura used the concept of selective cost to support his theory of neutral evolution (Kimura 1968; 1995; Kimura and Maruyama 1969; Jensen et al 2019). Other studies have pointed out various possible ways in which the cost of allele substitution at many loci might be reduced. But all of these solutions involve changing some of the starting assumptions. For example, Maynard Smith (1968) and Sved (1968) pointed out that the cost could be reduced if we assume some form of truncation selection rather than independent selection, while Grant and Flake (1974) showed that the cost could be reduced in a sub-divided population. These solutions are not very satisfying, however, since they involve modifying Haldane’s original assumptions of independent positive selection in a single undivided population. Moreover, while these modifying assumptions can reduce the accumulation of costs over loci, they do not eliminate it.

In this study, we wished to revisit the problem using Haldane’s original assumptions about population structure and the form of the selection function. We point out, however, that Haldane (1957) did not take sexual reproduction into account. In this paper, we use numerical simulations to demonstrate that Haldane’s argument about the summation of the costs over multiple loci does not apply to a sexually reproducing population – and this is true regardless of the selection scheme or the population structure. In the Discussion below, we explain why Haldane’s concept of cumulative costs applies only to a quasi-infinite asexual population. A randomly outbreeding sexual population, on the other hand, avoids the problem envisaged by Haldane.

Haldane’s reasoning (Haldane 1957) for the cost of natural selection contains two parts. First, there is the calculation of the reproductive cost of substituting one allele for another at a single locus during the process of adaptive evolution. As an example, Haldane cited the rapid spread of industrial melanism among populations of the peppered moth, *Biston betularia*, in the coal-mining areas of northern England (Kettlewell 1956). In order for the melanic types to become common, the non-melanic genotypes must be removed from the population. It is this removal of the existing type that constitutes the cost of natural selection. In general, the time to fixation of the new type is a function of the reproductive capacity and the initial frequency (see Felsenstein 1971).

The second part of Haldane’s argument is about extending the idea of the selective cost of adaptive evolution to many loci. For example, he argues that industrialization might have simultaneously selected for other traits in *B. betularia*, in addition to melanism. Given that the cost is a function of the initial frequency of the favored genotype, and based on the fact that the expected frequency of genotypes carrying multiple favorable alleles (at different loci, given linkage equilibrium between loci) is much lower than the frequency of genotypes carrying a single favorable allele, Haldane reasoned that the cost of natural selection at multiple loci would be larger than at a single locus. In fact, he estimated that there is an approximate summation of these costs over many loci. Thus, it would take approximately twice as long for a genotype carrying two favorable alleles to reach fixation as it would for a single mutant (Grant and Flake 1974). His conclusion was that *B. betularia* might not have sufficient reproductive capacity to allow adaptive evolution at ten loci simultaneously.

As we demonstrate in this study, it is this accumulation of costs over several loci that is avoided in a sexually outbreeding population. Specifically, in a sexual population we do not need to assume an initial population that is large enough to contain at least one individual with the optimum genotype. And it is precisely this assumption of a very large initial population that led Haldane to his conclusion that the costs of selection would accumulate over loci.

## Materials and Methods

In our simulations, we wished to recreate as closely as possible the situation envisaged by Haldane. Therefore, we considered a single undivided population of 100,000 diploid individuals in which rare favorable alleles were already present at several loci. As in Haldane’s model, the initial population was in linkage equilibrium. Also following Haldane’s example, selection at different loci was independent, i.e., fitness effects were multiplicative with no epistasis and no dominance (heterozygotes had a fitness intermediate between the homozygotes). The number of offspring produced by each individual was determined by sampling a Poisson distribution with a mean of 2. Thus, the population conformed to Haldane’s idea of ‘modest reproductive capacity”. All parents were diploid and had the same average fertility; selection was implemented on offspring viability based on the relative genotypic fitness values; i.e., the fittest individuals had the highest probability of survival from zygote to adult.

In addition to replicating Haldane’s model for an asexual population, we did a parallel set of simulations for a diploid, outbreeding sexual population. For the sexual populations, loci are evenly spaced along a linear chromosome. Recombination was implemented as follows: a single recombination event occurred between the two chromosomes in a diploid parent. This is equivalent to one chiasma per meiotic chromosome pair. The position of this recombination event along the chromosome was chosen at random; in the case where we consider 100 loci, this means that the probability of recombination between adjacent loci was 1%. The resulting recombinant represents a sexual gamete. These gametes were then paired at random to produce diploid zygotes for the next generation. All sexual parents produced the same average number of diploid zygotes, i.e., an average of two zygotes per parent. This means that there was no selection at the haploid stage of the life cycle. Viability selection was implemented on the sexual offspring, in the same way as for the asexual population.

We performed numerical simulations of populations containing 100,000 diploid individuals. We did ten replicates of each simulation and the results are shown in the Figures below. The initial frequency of the favored allele is 0.05 at each locus. The fitness advantage of a favored allele is 0.02 and fitness interactions are multiplicative between loci. Individuals produced an average of two offspring each and selection occurred through differential viability that was dependent on the relative genotypic fitness value.

We did a series of simulations in which we varied the number of loci under selection. Specifically, we did simulations for two, four and one hundred selected loci; this illustrates how the system behaves as we increase the number of loci under independent selection. In each case, we compared the outcome of selection in asexual and sexual populations. Following Haldane’s example, we assumed that the favorable alleles were already segregating at a low frequency in the initial population, and that the population was in linkage equilibrium. The source code for the simulations is available on GitHub (https://github.com/gbgolding/evolutionSex).

## Results

The results described below illustrate two main points. First, we show why it was necessary for Haldane (1957) to implicitly assume a progressively larger initial population as the number of loci under selection increased. The reason was that Haldane’s model did not include recombination between the selected loci; therefore, the initial population had to be large enough to contain at least one individual that contained all of the favorable mutations in its genotype. Secondly, we show that there is no need to increase the initial population size for multi-locus selection in a sexually outbreeding population. This is because recombination will automatically produce the genotype with the maximum number of favored alleles later on during the selection process (see Discussion below).

First, we performed simulations of the situation where there is only one locus undergoing selection. This allowed us to check the accuracy of our simulation method against the theoretical predictions of classical population genetics and it also provided a reference point for the subsequent simulations involving more than one locus. The results are presented in Figure 1. The results of ten replicate simulations are shown in Figure 1a, and the expected rise in allele frequency based on classical population genetics theory (Li 1955 page 258) is shown in Figure 1b. From the Figure, we see that the simulation results match the theoretical prediction very well and that it takes approximately 400 generations for the favored allele to rise from its initial frequency of 0.05 to near fixation.

**Figure 1.**
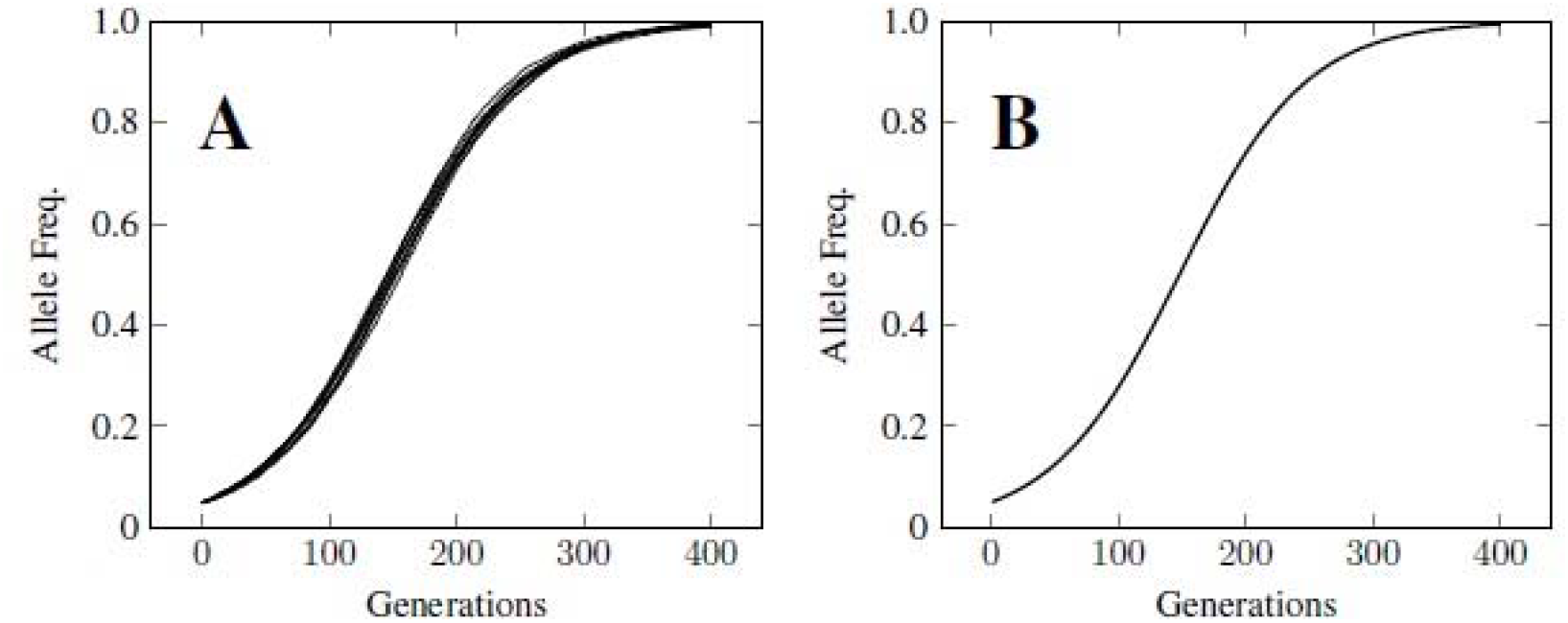
Selection at a single locus, given an initial frequency of 0.05 and a selective advantage of 0.02. The population size was 100,000 diploid individuals. Panel A Shows simulation results for 400 generations (10 replicates, results for individual replicates are shown). Panel B shows the theoretical, predicted rise in frequency of a favored allele at a single locus given an initial frequency of 0.05 and a selective advantage of 0.02.

Having established that the simulation provides the expected results for selection at a single locus, we then performed simulations of selection acting at multiple loci in both sexual and asexual populations. The results for two loci are shown in Figure 2, Panels A and B. We see that the asexual and sexual populations yield different results even when we are dealing with only two loci. This can be explained as follows. Given that the initial frequencies of the favored alleles at the two loci are 0.05, then the expected frequency of chromosomes carrying both favored alleles is one in four 400. But the expected frequency of diploids carrying two such chromosomes is only one in 160,000; therefore, we may not see even a single such double homozygote in the initial population. In most replicates of the simulation, the expected single double homozygote is indeed missing, leaving the fittest genotype with three rather than four favorable alleles (see Figure 2, Panel A). In one “lucky” replicate, the initial population does contain a double homozgote. Of course, if we had assumed a higher initial frequency for the favored alleles – say 0.1 rather than 0.05 – then the initial population would have contained several double homozygotes of the favored combination in all replicates. On the other hand, the problem would have become even more acute if we had used lower initial allele frequencies, say 0.01. In contrast to the asexual population, however, the sexual population can generate diploid genotypes carrying four favorable alleles in all ten replicates as the allele frequencies rise at the individual loci (see Figure 2, Panel B). In other words, recombination compensates for the absence of the optimal genotype in the initial population.

**Figure 2.**
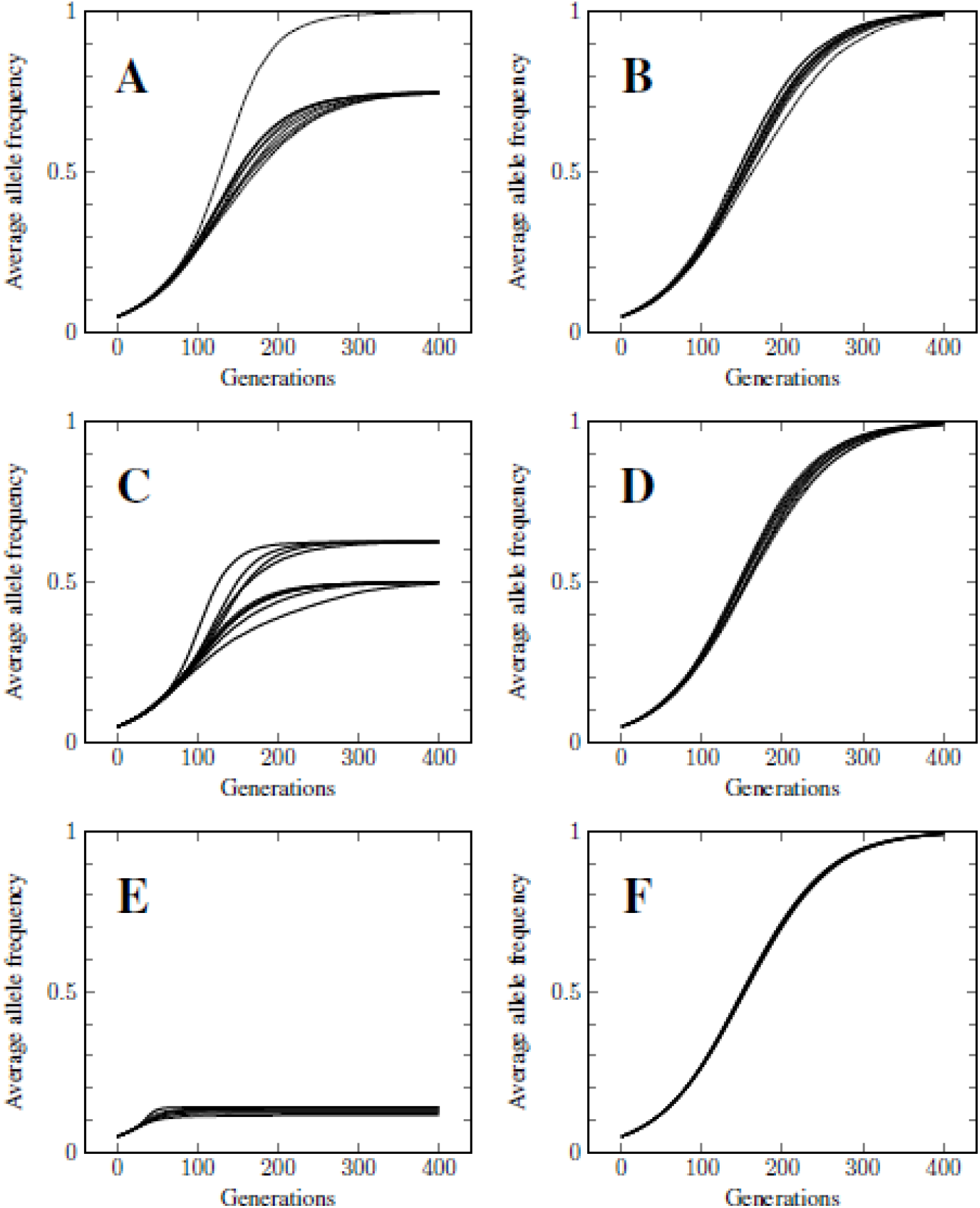
Selection at multiple loci, given an initial frequency of 0.05 for the favorable alleles and a selective advantage of 0.02. The graph shows the rise in the average frequency of favorable alleles during the course of selection. The reason that this frequency does not reach a value of 1.0 in the asexual populations is because the highest fitness genotype in the initial populations did not contain the maximum number of favorable alleles (see text). Panels A and B show the results for asexual and sexual populations, respectively, when selection is acting at two loci. Results for each of ten replicate simulations are shown separately. Panels C and D show comparable results for four loci. Panels E and F show the results for 100 loci.

The results for selection acting simultaneously at four loci are shown in Figure 2, Panels C and D. In this case, we begin to see an even greater difference between the asexual and sexual populations. The reason for this difference is that the initial asexual population lacks many high fitness genotypes. And the reason for this absence is that the expected frequency of such individuals – given the initial allele frequencies of 0.05 and linkage equilibrium – is exceedingly small. For example, the expected frequency of diploid genotypes which are homozygous for the favorable allele at all four loci is less than 10^−10^. Consequently, it would require a population size of billions of diploid individuals in order to see even a single individual of the optimal genotype within the initial population. From Figure 2c we can see that the fittest genotypes in the asexual population contained only four or five out of the eight possible favorable alleles – giving average allele frequencies after selection of 0.5 and 0.625, respectively at the end of the simulation. Of course, the sexual population is also lacking these high fitness genotypes initially, but this is not a problem because random recombination can assemble such genotypes as the individual allele frequencies rise in response to selection. For example, once the allele frequencies have reached a frequency of 0.5 in an outbreeding sexual population, we expect to see some individuals with the optimal genotype begin to appear within the population of 100,000 diploid individuals. This explains why the average allele frequency approaches a value of 1.0 in all replicates of the sexual population (see Figure 2d).

Finally, the results for selection acting on 100 loci are shown in Figure 2, Panels E and F. In this case, there is a striking difference between the asexual and sexual populations. The initial population contains only low fitness genotypes where we expect an average of only five favorable alleles per chromosome – given that there are 100 loci and a frequency of 0.05 for the favorable allele at each locus (see Hickey and Golding 2018). The asexual population is again limited to selecting among such low-fitness genotypes. In contrast to this, the sexual population can use recombination to gradually assemble chromosomes with increasing numbers of favorable alleles per chromosome as the frequencies of the favorable alleles rise in response to selection. At the end of the selection period, the sexual population consists of individuals that are homozygous for the favored allele at all one hundred loci. Table 1 shows a summary of the response to selection in both the asexual and the sexual populations. As can be seen in Table 1, the asexual population is constrained by the maximum fitness in the initial population; in contrast to this recombination continues to produce increasing maximum fitnesses as the selection proceeds. This allows the sexual population to advance beyond the highest fitness value in the initial population. The complete numerical results for the simulations of 100 loci are provided in Supplementary Tables S1 and S2.

**Table 1.** Response to multi-locus selection in asexual and sexual populations. The average fitness and maximum fitness are shown for each tenth generation, up to generation 100; the complete data for all generations are shown in Supplemental Tables T1 and T2. The population size was 100,000 diploid individuals. The initial frequency of the favorable allele at each of 100 the loci was 0.05. The selective advantage of each favorable allele was 0.02. The initial population was in linkage equilibrium. Fitness effects were multiplicative between loci.

| Generation | Asexual |  | Sexual |  |
| --- | --- | --- | --- | --- |
|  | Av. fitness | Max. fitness | Av. fitness | Max. fitness |
| 0 | 1.22 | 1.64 | 1.22 | 1.70 |
| 10 | 1.27 | 1.64 | 1.27 | 1.71 |
| 20 | 1.33 | 1.64 | 1.33 | 1.88 |
| 30 | 1.40 | 1.64 | 1.41 | 2.00 |
| 40 | 1.47 | 1.64 | 1.50 | 2.43 |
| 50 | 1.53 | 1.64 | 1.62 | 2.43 |
| 60 | 1.57 | 1.64 | 1.77 | 2.85 |
| 70 | 1.59 | 1.64 | 1.96 | 3.26 |
| 80 | 1.61 | 1.64 | 2.21 | 3.68 |
| 90 | 1.62 | 1.64 | 2.52 | 4.39 |
| 100 | 1.63 | 1.64 | 2.91 | 4.94 |

These results show that, in a sexually outbreeding population, the frequency of favorable alleles at many loci can respond simultaneously to independent selection at rates that are very similar to the predicted rate at a single locus (compare Figure 2, Panel F and Figure 1, Panel B). This implies that the rate of response to selection at any given locus is not greatly impeded by selection at other loci. In other words, many loci can respond simultaneously to independent selection – but only in a sexually reproducing population. In summary, our results show that an asexual population cannot respond to multi-locus selection as efficiently as a sexual population because it cannot generate higher-fitness genotypic combinations as the allelic frequencies increase in response to selection. Moreover, the relative disadvantage of the asexual population increases as the number of loci under selection increases (compare Panels A, C and E in Figure 2).

## Discussion

As stated in the introduction, Haldane (1957) argued that the costs of natural selection accumulate over loci, i.e., the reproductive cost of allele substitution at two loci would be twice the cost at a single locus. Consequently, the time required for substitution should be twice as long. In general, if it takes t generations for a gene substitution at a single locus, it should take Lt generations for substitutions at L loci. Applying this logic to our model, given that it takes 400 generations for allelic substitution at one locus, it should take 40,000 generations for substitutions at 100 loci. This prediction is in stark contrast to our finding that it takes only 400 generations for allelic substitution at all 100 loci (see Figure 2, Panel F). How can we explain this major difference between Haldane’s prediction and our findings?

Clearly, Haldane’s scenario does not explain our simulation results as shown in Figure 2. The reason for the discrepancy is, we believe, that Haldane’s argument assumed that the optimal genotypic combination had to be already present in the initial population and that genotypes could be considered as fixed entities that reproduce themselves from one generation to the next. But these assumptions do not hold for a sexually outbreeding population. First, the population is usually not large enough to contain even a single individual with the vanishingly rare optimal genotypic combination; this fact has already been pointed out by Ewens (1972) and Maynard Smith (1976). Secondly, specific genotypes are broken down by recombination each generation and replaced by other genotypic combinations (Hickey and Golding 2018). The process of recombination involves the continual disassembly of existing genotypes and reassembly of new genotypes from one generation to the next. Because of this process, however, those genotypes which are expected to be vanishingly rare when the allele frequencies are low will be automatically generated through recombination once the allele frequencies begin to rise in response to selection (Muller 1964; Hickey and Golding 2018). Consequently, there is no need for the massive culling that would be required in a very large initial population that contained an extremely rare optimal genotype. Thus, in an outbreeding sexual population, recombination solves the perceived problem of costs that are cumulative over different genetic loci. In Haldane’s scenario, it is necessary to “grow” the optimal genotypic combination from an initial, vanishingly rare frequency to fixation. In practice, however, this optimal combination is produced by recombination only near the end of the process. This avoids the huge cumulative cost of natural selection.

Haldane’s theory is mathematically very elegant; what happens in nature is mathematically more muddled, but it is much more efficient. In nature, selection favors a wide variety of suboptimal combinations which are then recombined. In this way, recombination can harvest all favorable alleles from any genetic background. It is often not until quite late in the selection process that the optimal genotypic combination first appears. For example, even after the allele frequencies at each of 100 loci have risen to a frequency of 0.9, the probability of randomly generating a chromosome with a favorable allele at all 100 loci is still less than 0.0001. By the time the individual alleles have reached a frequency of 0.99, however, more than one third of all chromosomes in the population are expected to contain the optimal combination - i.e., a favorable allele at all 100 loci. In other words, rather than increasing slowly from an infinitesimally low initial frequency as suggested by Haldane (1957), the optimal combination is generated through recombination near the end of the selection process.

The strategy used by nature is somewhat analogous to the process of parallel computing. Since recombination decouples the allelic changes at a given locus from changes at other loci, it effectively decomposes multi-locus selection into several simultaneous instances of individual selection. Haldane (1957) envisaged that natural selection would have to do an exhaustive search through all possible genotypic combinations; instead, what happens in nature is a massively parallel heuristic search. The process of alternating selection and recombination enables the sexual population to trace a relatively narrow path through the myriad of possible genotypic combinations until, eventually, it arrives at the optimum genotypic combination. A comparison between natural selection and parallel computing has also been made by Wilf and Ewens (2010) although those authors were referring to the sequential fixation of favorable alleles during long-term evolution, rather than to the effects of recombination within a population.

The evolution of sex is a problem that has puzzled biologists for many decades (see Williams 1975; Maynard Smith 1978; Kondrashov 2018 for general reviews). Maynard Smith (1971) asked “what use is sex?” Our answer to that question is that recombination avoids the potentially increasing costs of natural selection as the number of genes under selection increases. This answer was already hinted at in the literature, ever since Fisher (1930) stated that the advantage of sexual reproduction is proportional to the number of genetic loci under selection. Muller (1932; 1964) developed Fisher’s idea and concluded that recombination among many loci could allow a sexual population to evolve “hundreds to millions of times faster” than a comparable asexual population. This same idea of an increasing advantage for recombination as the number of loci under selection increases, has been echoed in later papers (e.g., Crow and Kimura 1965; Otto and Barton 2001; Colegrave 2002; Iles et al 2003; Park and Krug 2013; Edhan et al 2017). Park and Krug (2013) have shown that the advantage of recombination is proportional to L, where L is the number of loci under selection. But the majority of these previous studies deal with the effect of recombination on newly-arising favorable mutations (e.g., Christiansen et al 1998) whereas the focus of this study was on the standing genetic variation that is already present in the population. Also, none of these previous studies offers a biological explanation of why the advantage of sex should increase as we consider more loci under selection. Our explanation for this correlation is that an asexual population would be subject to an increasing reproductive cost as the number of loci increases – as shown by Haldane (1957). But since sexual reproduction eliminates the accumulation of costs over loci (as shown in this study), this leads to an increasing relative advantage of sex as the number of loci under selection increases. Rather than saying that the advantage of sexual reproduction increases as the number of loci under selection increases, it might be more accurate to say that there is an increasing disadvantage to asexual reproduction as the number of loci under selection increases (see Figure 2, Panels A, C and E).

It is interesting to note that Haldane (1957; 1960) did not see the relevance of Muller’s ideas about recombination to the problem of reproductive costs that accumulated over loci. It is perhaps even more interesting that Muller (1964) also did not see the connection between his own ideas and the problem posed by Haldane (1957; 1960). But neither did their colleagues at the time see the connection (e.g. VanValen 1963). Indeed, we believe that we are the first authors to have pointed out the direct link between the ideas of Muller (1932; 1964) and those of Haldane (1957; 1960). And we have to admit that it was only because of the counterintuitive nature of our own simulation results that we were forced to re-examine Haldane’s argument in the light of recombination. One could say that the solution to Haldane’s Dilemma has been “hiding in plain sight” for six decades.

Our results support models of adaptation based on the standing genetic variation (see Barrett and Schluter 2007; Orr 2005; Reznick 2016). As noted by Messer et al (2016) rapid phenotypic evolution suggests that many genomic loci may contribute to strongly selected traits. Molecular studies have confirmed that indeed several loci are generally involved during rapid evolution (Burke et al 2010; Teotónio et al 2009). Moreover, there is direct experimental evidence that recombination facilitates the rapid response to multi-locus selection (McDonald et al 2016; Kosheleva and Desai 2018). The relevance of our work is that it removes any concerns about the high cumulative costs of multi-locus selection as argued by Haldane (1957), provided of course that there is recombination occurring between the selected genes.

In a genetically polymorphic sexual population multi-locus genotypes are highly ephemeral. This is because existing genotypes are merely a small random sample of a much larger set of possible genotypes (Ewens 1972; Edhan 2017). In this study, we assumed – for computational convenience – that there were only 100 biallelic loci. The early, conservative estimates of genetic variation in natural populations concluded that approximately twenty percent of the loci within the eukaryotic genome are polymorphic (Lewontin and Hubby 1966). If we assume that there are 4,000 polymorphic loci – with two alleles at each locus – there are 2^4,000^ possible genotypic combinations, a number which is greater than 10^1,000^. This number is far greater than the total number of atoms in the observable universe – which has been estimated to be approximately 10^80^ (see Hogan 2000). Darwin stated that “endless forms” have evolved as a result of natural selection. We could say that this process has been facilitated through the production of effectively endless genotypic combinations as a result of sexual reproduction and recombination.

In conclusion, our results show that outbreeding and recombination allow many genes to respond rapidly to independent selection at the same time. In a sexual population, selection on one gene does not slow the rate of adaptive response at another genetic locus. This is because recombination allows allelic change at one locus to become independent of the changes at other loci. Consequently, even those organisms with limited population sizes and modest reproductive capacity can evolve efficiently. Haldane (1957) predicted that the cost of multi-locus selection would pose a particularly difficult challenge for such species. Perhaps this explains why sex tends to be most common among species with limited population sizes and modest reproductive capacity.

## Acknowledgements

This work was supported by a Discovery Grant from NSERC Canada to GBG (grant no. 140221-10). We are grateful to Richard Lewontin for stimulating discussions during the early development of this work. We would like to thank Chi Kuen Cheung for his support and encouragement.

