## Supplementary Table 1 for "Sex Solves Haldane’s Dilemma"

```
Supplemental Table S1.  Simulation results for ten replicate runs with asexual reproduction.

Each simulation was run for 400 generations (see main manuscript text).
Note that in some replicates, the fittest genotype was represented by only a single individual and was lost by random drift in the early generations of the simulation.


gen  popSize    avgFit     maxFit maxAlleles avgAlleles
  0  100000     1.22131    1.64061     25    10.0069
  1  100000     1.22131    1.64061     25    10.0069
  2  100000     1.22599    1.64061     25    10.1983
  3  100000     1.23078    1.63998     25    10.3937
  4  100000     1.23573    1.60844     24    10.5953
  5  100000     1.24031    1.60844     24    10.7809
  6  100000     1.24517    1.60844     24    10.9769
  7  100000     1.25019    1.60844     24    11.1785
  8  100000     1.25537    1.60844     24    11.3855
  9  100000     1.26060    1.60844     24    11.5943
 10  100000     1.26598    1.60844     24    11.8078
 11  100000     1.27114    1.60844     24    12.0115
 12  100000     1.27665    1.60844     24    12.2283
 13  100000     1.28238    1.60844     24    12.4529
 14  100000     1.28826    1.60844     24    12.6829
 15  100000     1.29405    1.60844     24    12.9091
 16  100000     1.30012    1.60844     24    13.1449
 17  100000     1.30635    1.60844     24    13.3851
 18  100000     1.31250    1.60844     24    13.6209
 19  100000     1.31894    1.60844     24    13.8671
 20  100000     1.32520    1.60844     24    14.1054
 21  100000     1.33173    1.60844     24    14.3528
 22  100000     1.33820    1.60844     24    14.5972
 23  100000     1.34476    1.60844     24    14.8439
 24  100000     1.35150    1.60844     24    15.0967
 25  100000     1.35823    1.60844     24    15.3471
 26  100000     1.36490    1.60844     24    15.5947
 27  100000     1.37156    1.60844     24    15.8411
 28  100000     1.37834    1.60844     24    16.0913
 29  100000     1.38484    1.60844     24    16.3304
 30  100000     1.39187    1.60844     24    16.5882
 31  100000     1.39874    1.60844     24    16.8392
 32  100000     1.40559    1.60844     24    17.0881
 33  100000     1.41225    1.60844     24    17.3292
 34  100000     1.41879    1.60844     24    17.5645
 35  100000     1.42510    1.60844     24    17.7907
 36  100000     1.43148    1.60844     24    18.0201
 37  100000     1.43762    1.60844     24    18.2390
 38  100000     1.44330    1.60844     24    18.4407
 39  100000     1.44908    1.60844     24    18.6462
 40  100000     1.45511    1.60844     24    18.8594
 41  100000     1.46031    1.60844     24    19.0429
 42  100000     1.46549    1.60844     24    19.2251
 43  100000     1.47056    1.60844     24    19.4028
 44  100000     1.47558    1.60844     24    19.5784
 45  100000     1.48044    1.60844     24    19.7478
 46  100000     1.48504    1.60844     24    19.9078
 47  100000     1.48956    1.60844     24    20.0651
 48  100000     1.49390    1.60844     24    20.2155
 49  100000     1.49762    1.60844     24    20.3443
 50  100000     1.50173    1.60844     24    20.4857
 51  100000     1.50519    1.60844     24    20.6047
 52  100000     1.50874    1.60844     24    20.7264
 53  100000     1.51231    1.60844     24    20.8489
 54  100000     1.51533    1.60844     24    20.9523
 55  100000     1.51827    1.60844     24    21.0525
 56  100000     1.52145    1.60844     24    21.1605
 57  100000     1.52425    1.60844     24    21.2558
 58  100000     1.52707    1.60844     24    21.3514
 59  100000     1.52961    1.60844     24    21.4377
 60  100000     1.53202    1.60844     24    21.5188
 61  100000     1.53454    1.60844     24    21.6040
 62  100000     1.53666    1.60844     24    21.6753
 63  100000     1.53865    1.60844     24    21.7422
 64  100000     1.54064    1.60844     24    21.8091
 65  100000     1.54250    1.60844     24    21.8715
 66  100000     1.54428    1.60844     24    21.9309
 67  100000     1.54580    1.60844     24    21.9818
 68  100000     1.54752    1.60844     24    22.0392
 69  100000     1.54925    1.60844     24    22.0965
 70  100000     1.55087    1.60844     24    22.1503
 71  100000     1.55244    1.60844     24    22.2025
 72  100000     1.55381    1.60844     24    22.2479
 73  100000     1.55539    1.60844     24    22.3004
 74  100000     1.55669    1.60844     24    22.3436
 75  100000     1.55802    1.60844     24    22.3873
 76  100000     1.55927    1.60844     24    22.4283
 77  100000     1.56044    1.60844     24    22.4670
 78  100000     1.56180    1.60844     24    22.5117
 79  100000     1.56300    1.60844     24    22.5510
 80  100000     1.56411    1.60844     24    22.5877
 81  100000     1.56497    1.60844     24    22.6157
 82  100000     1.56606    1.60844     24    22.6511
 83  100000     1.56691    1.60844     24    22.6789
 84  100000     1.56793    1.60844     24    22.7120
 85  100000     1.56896    1.60844     24    22.7458
 86  100000     1.56969    1.60844     24    22.7693
 87  100000     1.57062    1.60844     24    22.7995
 88  100000     1.57129    1.60844     24    22.8212
 89  100000     1.57201    1.60844     24    22.8446
 90  100000     1.57299    1.60844     24    22.8763
 91  100000     1.57369    1.60844     24    22.8991
 92  100000     1.57444    1.60844     24    22.9234
 93  100000     1.57519    1.60844     24    22.9476
 94  100000     1.57598    1.60844     24    22.9734
 95  100000     1.57659    1.60844     24    22.9930
 96  100000     1.57728    1.60844     24    23.0151
 97  100000     1.57783    1.60844     24    23.0329
 98  100000     1.57856    1.60844     24    23.0564
 99  100000     1.57923    1.60844     24    23.0780
100  100000     1.57984    1.60844     24    23.0978
101  100000     1.58042    1.60844     24    23.1163
102  100000     1.58094    1.60844     24    23.1330
103  100000     1.58144    1.60844     24    23.1488
104  100000     1.58200    1.60844     24    23.1669
105  100000     1.58244    1.60844     24    23.1807
106  100000     1.58291    1.60844     24    23.1960
107  100000     1.58356    1.60844     24    23.2169
108  100000     1.58408    1.60844     24    23.2336
109  100000     1.58462    1.60844     24    23.2508
110  100000     1.58507    1.60844     24    23.2652
111  100000     1.58555    1.60844     24    23.2804
112  100000     1.58598    1.60844     24    23.2942
113  100000     1.58636    1.60844     24    23.3062
114  100000     1.58681    1.60844     24    23.3206
115  100000     1.58717    1.60844     24    23.3319
116  100000     1.58754    1.60844     24    23.3439
117  100000     1.58796    1.60844     24    23.3572
118  100000     1.58832    1.60844     24    23.3688
119  100000     1.58877    1.60844     24    23.3829
120  100000     1.58909    1.60844     24    23.3934
121  100000     1.58945    1.60844     24    23.4048
122  100000     1.58990    1.60844     24    23.4189
123  100000     1.59026    1.60844     24    23.4303
124  100000     1.59057    1.60844     24    23.4403
125  100000     1.59099    1.60844     24    23.4535
126  100000     1.59130    1.60844     24    23.4633
127  100000     1.59173    1.60844     24    23.4769
128  100000     1.59206    1.60844     24    23.4873
129  100000     1.59231    1.60844     24    23.4952
130  100000     1.59257    1.60844     24    23.5035
131  100000     1.59283    1.60844     24    23.5115
132  100000     1.59309    1.60844     24    23.5199
133  100000     1.59341    1.60844     24    23.5298
134  100000     1.59378    1.60844     24    23.5414
135  100000     1.59403    1.60844     24    23.5495
136  100000     1.59432    1.60844     24    23.5586
137  100000     1.59451    1.60844     24    23.5644
138  100000     1.59478    1.60844     24    23.5729
139  100000     1.59500    1.60844     24    23.5798
140  100000     1.59526    1.60844     24    23.5883
141  100000     1.59556    1.60844     24    23.5976
142  100000     1.59578    1.60844     24    23.6045
143  100000     1.59610    1.60844     24    23.6145
144  100000     1.59637    1.60844     24    23.6231
145  100000     1.59669    1.60844     24    23.6331
146  100000     1.59691    1.60844     24    23.6401
147  100000     1.59711    1.60844     24    23.6464
148  100000     1.59734    1.60844     24    23.6536
149  100000     1.59757    1.60844     24    23.6608
150  100000     1.59778    1.60844     24    23.6674
151  100000     1.59792    1.60844     24    23.6720
152  100000     1.59815    1.60844     24    23.6791
153  100000     1.59836    1.60844     24    23.6856
154  100000     1.59855    1.60844     24    23.6916
155  100000     1.59877    1.60844     24    23.6984
156  100000     1.59893    1.60844     24    23.7034
157  100000     1.59914    1.60844     24    23.7103
158  100000     1.59927    1.60844     24    23.7142
159  100000     1.59946    1.60844     24    23.7201
160  100000     1.59963    1.60844     24    23.7255
161  100000     1.59983    1.60844     24    23.7317
162  100000     1.59999    1.60844     24    23.7367
163  100000     1.60010    1.60844     24    23.7401
164  100000     1.60026    1.60844     24    23.7453
165  100000     1.60042    1.60844     24    23.7503
166  100000     1.60059    1.60844     24    23.7554
167  100000     1.60075    1.60844     24    23.7604
168  100000     1.60085    1.60844     24    23.7637
169  100000     1.60101    1.60844     24    23.7687
170  100000     1.60116    1.60844     24    23.7733
171  100000     1.60131    1.60844     24    23.7780
172  100000     1.60143    1.60844     24    23.7818
173  100000     1.60151    1.60844     24    23.7841
174  100000     1.60161    1.60844     24    23.7875
175  100000     1.60169    1.60844     24    23.7900
176  100000     1.60186    1.60844     24    23.7951
177  100000     1.60198    1.60844     24    23.7988
178  100000     1.60209    1.60844     24    23.8023
179  100000     1.60224    1.60844     24    23.8072
180  100000     1.60238    1.60844     24    23.8114
181  100000     1.60247    1.60844     24    23.8141
182  100000     1.60257    1.60844     24    23.8173
183  100000     1.60269    1.60844     24    23.8210
184  100000     1.60276    1.60844     24    23.8234
185  100000     1.60285    1.60844     24    23.8261
186  100000     1.60290    1.60844     24    23.8275
187  100000     1.60304    1.60844     24    23.8319
188  100000     1.60312    1.60844     24    23.8345
189  100000     1.60323    1.60844     24    23.8380
190  100000     1.60332    1.60844     24    23.8406
191  100000     1.60340    1.60844     24    23.8432
192  100000     1.60349    1.60844     24    23.8459
193  100000     1.60360    1.60844     24    23.8496
194  100000     1.60372    1.60844     24    23.8532
195  100000     1.60378    1.60844     24    23.8552
196  100000     1.60387    1.60844     24    23.8580
197  100000     1.60397    1.60844     24    23.8611
198  100000     1.60405    1.60844     24    23.8635
199  100000     1.60413    1.60844     24    23.8659
200  100000     1.60420    1.60844     24    23.8681
201  100000     1.60430    1.60844     24    23.8714
202  100000     1.60439    1.60844     24    23.8740
203  100000     1.60447    1.60844     24    23.8767
204  100000     1.60453    1.60844     24    23.8786
205  100000     1.60463    1.60844     24    23.8815
206  100000     1.60471    1.60844     24    23.8842
207  100000     1.60480    1.60844     24    23.8867
208  100000     1.60488    1.60844     24    23.8894
209  100000     1.60494    1.60844     24    23.8911
210  100000     1.60494    1.60844     24    23.8912
211  100000     1.60501    1.60844     24    23.8932
212  100000     1.60505    1.60844     24    23.8947
213  100000     1.60513    1.60844     24    23.8971
214  100000     1.60516    1.60844     24    23.8980
215  100000     1.60528    1.60844     24    23.9017
216  100000     1.60534    1.60844     24    23.9037
217  100000     1.60540    1.60844     24    23.9056
218  100000     1.60543    1.60844     24    23.9063
219  100000     1.60547    1.60844     24    23.9076
220  100000     1.60554    1.60844     24    23.9099
221  100000     1.60558    1.60844     24    23.9111
222  100000     1.60562    1.60844     24    23.9125
223  100000     1.60567    1.60844     24    23.9139
224  100000     1.60575    1.60844     24    23.9163
225  100000     1.60582    1.60844     24    23.9186
226  100000     1.60588    1.60844     24    23.9205
227  100000     1.60594    1.60844     24    23.9225
228  100000     1.60597    1.60844     24    23.9233
229  100000     1.60601    1.60844     24    23.9246
230  100000     1.60608    1.60844     24    23.9268
231  100000     1.60612    1.60844     24    23.9279
232  100000     1.60615    1.60844     24    23.9289
233  100000     1.60622    1.60844     24    23.9310
234  100000     1.60627    1.60844     24    23.9328
235  100000     1.60631    1.60844     24    23.9340
236  100000     1.60637    1.60844     24    23.9358
237  100000     1.60637    1.60844     24    23.9357
238  100000     1.60641    1.60844     24    23.9370
239  100000     1.60641    1.60844     24    23.9369
240  100000     1.60643    1.60844     24    23.9376
241  100000     1.60646    1.60844     24    23.9386
242  100000     1.60648    1.60844     24    23.9393
243  100000     1.60650    1.60844     24    23.9400
244  100000     1.60654    1.60844     24    23.9410
245  100000     1.60658    1.60844     24    23.9422
246  100000     1.60661    1.60844     24    23.9433
247  100000     1.60666    1.60844     24    23.9449
248  100000     1.60669    1.60844     24    23.9456
249  100000     1.60670    1.60844     24    23.9461
250  100000     1.60673    1.60844     24    23.9471
251  100000     1.60676    1.60844     24    23.9478
252  100000     1.60678    1.60844     24    23.9486
253  100000     1.60681    1.60844     24    23.9494
254  100000     1.60684    1.60844     24    23.9505
255  100000     1.60687    1.60844     24    23.9515
256  100000     1.60690    1.60844     24    23.9523
257  100000     1.60694    1.60844     24    23.9536
258  100000     1.60696    1.60844     24    23.9541
259  100000     1.60699    1.60844     24    23.9550
260  100000     1.60701    1.60844     24    23.9557
261  100000     1.60703    1.60844     24    23.9562
262  100000     1.60707    1.60844     24    23.9574
263  100000     1.60712    1.60844     24    23.9592
264  100000     1.60715    1.60844     24    23.9599
265  100000     1.60718    1.60844     24    23.9610
266  100000     1.60721    1.60844     24    23.9619
267  100000     1.60724    1.60844     24    23.9627
268  100000     1.60724    1.60844     24    23.9627
269  100000     1.60725    1.60844     24    23.9632
270  100000     1.60727    1.60844     24    23.9638
271  100000     1.60730    1.60844     24    23.9646
272  100000     1.60733    1.60844     24    23.9656
273  100000     1.60734    1.60844     24    23.9660
274  100000     1.60737    1.60844     24    23.9669
275  100000     1.60738    1.60844     24    23.9672
276  100000     1.60738    1.60844     24    23.9672
277  100000     1.60738    1.60844     24    23.9673
278  100000     1.60742    1.60844     24    23.9685
279  100000     1.60743    1.60844     24    23.9689
280  100000     1.60745    1.60844     24    23.9693
281  100000     1.60747    1.60844     24    23.9699
282  100000     1.60747    1.60844     24    23.9701
283  100000     1.60747    1.60844     24    23.9699
284  100000     1.60749    1.60844     24    23.9707
285  100000     1.60751    1.60844     24    23.9713
286  100000     1.60755    1.60844     24    23.9724
287  100000     1.60756    1.60844     24    23.9727
288  100000     1.60757    1.60844     24    23.9731
289  100000     1.60760    1.60844     24    23.9741
290  100000     1.60762    1.60844     24    23.9746
291  100000     1.60762    1.60844     24    23.9748
292  100000     1.60763    1.60844     24    23.9750
293  100000     1.60764    1.60844     24    23.9753
294  100000     1.60764    1.60844     24    23.9752
295  100000     1.60765    1.60844     24    23.9755
296  100000     1.60769    1.60844     24    23.9768
297  100000     1.60770    1.60844     24    23.9773
298  100000     1.60772    1.60844     24    23.9776
299  100000     1.60774    1.60844     24    23.9785
300  100000     1.60774    1.60844     24    23.9784
301  100000     1.60774    1.60844     24    23.9785
302  100000     1.60777    1.60844     24    23.9794
303  100000     1.60780    1.60844     24    23.9803
304  100000     1.60781    1.60844     24    23.9806
305  100000     1.60781    1.60844     24    23.9806
306  100000     1.60782    1.60844     24    23.9808
307  100000     1.60782    1.60844     24    23.9809
308  100000     1.60784    1.60844     24    23.9815
309  100000     1.60785    1.60844     24    23.9819
310  100000     1.60785    1.60844     24    23.9820
311  100000     1.60787    1.60844     24    23.9825
312  100000     1.60787    1.60844     24    23.9825
313  100000     1.60788    1.60844     24    23.9828
314  100000     1.60790    1.60844     24    23.9834
315  100000     1.60791    1.60844     24    23.9837
316  100000     1.60791    1.60844     24    23.9838
317  100000     1.60793    1.60844     24    23.9842
318  100000     1.60793    1.60844     24    23.9844
319  100000     1.60795    1.60844     24    23.9848
320  100000     1.60795    1.60844     24    23.9850
321  100000     1.60796    1.60844     24    23.9852
322  100000     1.60796    1.60844     24    23.9854
323  100000     1.60797    1.60844     24    23.9856
324  100000     1.60797    1.60844     24    23.9856
325  100000     1.60798    1.60844     24    23.9858
326  100000     1.60800    1.60844     24    23.9864
327  100000     1.60801    1.60844     24    23.9868
328  100000     1.60802    1.60844     24    23.9869
329  100000     1.60802    1.60844     24    23.9871
330  100000     1.60802    1.60844     24    23.9870
331  100000     1.60802    1.60844     24    23.9871
332  100000     1.60804    1.60844     24    23.9875
333  100000     1.60804    1.60844     24    23.9877
334  100000     1.60805    1.60844     24    23.9881
335  100000     1.60807    1.60844     24    23.9885
336  100000     1.60808    1.60844     24    23.9889
337  100000     1.60808    1.60844     24    23.9888
338  100000     1.60808    1.60844     24    23.9890
339  100000     1.60810    1.60844     24    23.9894
340  100000     1.60809    1.60844     24    23.9893
341  100000     1.60811    1.60844     24    23.9898
342  100000     1.60812    1.60844     24    23.9900
343  100000     1.60811    1.60844     24    23.9899
344  100000     1.60811    1.60844     24    23.9897
345  100000     1.60811    1.60844     24    23.9900
346  100000     1.60812    1.60844     24    23.9900
347  100000     1.60811    1.60844     24    23.9899
348  100000     1.60811    1.60844     24    23.9898
349  100000     1.60811    1.60844     24    23.9900
350  100000     1.60811    1.60844     24    23.9899
351  100000     1.60813    1.60844     24    23.9905
352  100000     1.60814    1.60844     24    23.9907
353  100000     1.60815    1.60844     24    23.9911
354  100000     1.60816    1.60844     24    23.9914
355  100000     1.60817    1.60844     24    23.9917
356  100000     1.60817    1.60844     24    23.9918
357  100000     1.60817    1.60844     24    23.9918
358  100000     1.60818    1.60844     24    23.9918
359  100000     1.60819    1.60844     24    23.9922
360  100000     1.60820    1.60844     24    23.9926
361  100000     1.60821    1.60844     24    23.9929
362  100000     1.60822    1.60844     24    23.9932
363  100000     1.60822    1.60844     24    23.9933
364  100000     1.60823    1.60844     24    23.9935
365  100000     1.60824    1.60844     24    23.9938
366  100000     1.60825    1.60844     24    23.9942
367  100000     1.60825    1.60844     24    23.9941
368  100000     1.60825    1.60844     24    23.9942
369  100000     1.60824    1.60844     24    23.9940
370  100000     1.60825    1.60844     24    23.9943
371  100000     1.60826    1.60844     24    23.9945
372  100000     1.60826    1.60844     24    23.9946
373  100000     1.60827    1.60844     24    23.9949
374  100000     1.60827    1.60844     24    23.9949
375  100000     1.60827    1.60844     24    23.9949
376  100000     1.60828    1.60844     24    23.9951
377  100000     1.60827    1.60844     24    23.9949
378  100000     1.60828    1.60844     24    23.9951
379  100000     1.60829    1.60844     24    23.9954
380  100000     1.60830    1.60844     24    23.9956
381  100000     1.60830    1.60844     24    23.9956
382  100000     1.60830    1.60844     24    23.9958
383  100000     1.60830    1.60844     24    23.9958
384  100000     1.60831    1.60844     24    23.9959
385  100000     1.60831    1.60844     24    23.9962
386  100000     1.60832    1.60844     24    23.9963
387  100000     1.60832    1.60844     24    23.9963
388  100000     1.60832    1.60844     24    23.9964
389  100000     1.60832    1.60844     24    23.9965
390  100000     1.60833    1.60844     24    23.9966
391  100000     1.60834    1.60844     24    23.9971
392  100000     1.60835    1.60844     24    23.9973
393  100000     1.60836    1.60844     24    23.9975
394  100000     1.60836    1.60844     24    23.9976
395  100000     1.60836    1.60844     24    23.9976
396  100000     1.60836    1.60844     24    23.9975
397  100000     1.60835    1.60844     24    23.9974
398  100000     1.60836    1.60844     24    23.9977
399  100000     1.60836    1.60844     24    23.9976
400  100000     1.60836    1.60844     24    23.9976

Asexual Model with selection (random number based on
relative fitness to determine who survives).  Mutation rate is      1e-08,
Recomb rate is ONE per asexual individual per generation
Starting values at generation   0 are; population size 100000,
with avg fitness of    1.22142.

gen  popSize    avgFit     maxFit maxAlleles avgAlleles
  0  100000     1.22142    1.60720     24    10.0105
  1  100000     1.22142    1.60720     24    10.0105
  2  100000     1.22595    1.57690     23    10.1954
  3  100000     1.23079    1.57690     23    10.3926
  4  100000     1.23560    1.57690     23    10.5885
  5  100000     1.24037    1.57629     23    10.7815
  6  100000     1.24530    1.57629     23    10.9807
  7  100000     1.25042    1.57629     23    11.1866
  8  100000     1.25568    1.57629     23    11.3967
  9  100000     1.26085    1.57629     23    11.6025
 10  100000     1.26636    1.57629     23    11.8209
 11  100000     1.27205    1.57629     23    12.0454
 12  100000     1.27808    1.57629     23    12.2825
 13  100000     1.28377    1.57629     23    12.5053
 14  100000     1.28962    1.57629     23    12.7336
 15  100000     1.29546    1.57629     23    12.9608
 16  100000     1.30136    1.57629     23    13.1892
 17  100000     1.30771    1.57629     23    13.4337
 18  100000     1.31425    1.57629     23    13.6850
 19  100000     1.32068    1.57629     23    13.9314
 20  100000     1.32672    1.57629     23    14.1617
 21  100000     1.33317    1.57629     23    14.4064
 22  100000     1.33929    1.57629     23    14.6375
 23  100000     1.34567    1.57629     23    14.8775
 24  100000     1.35224    1.57629     23    15.1235
 25  100000     1.35883    1.57629     23    15.3703
 26  100000     1.36537    1.57629     23    15.6142
 27  100000     1.37191    1.57629     23    15.8576
 28  100000     1.37820    1.57629     23    16.0908
 29  100000     1.38484    1.57629     23    16.3350
 30  100000     1.39158    1.57629     23    16.5832
 31  100000     1.39814    1.57629     23    16.8228
 32  100000     1.40460    1.57629     23    17.0591
 33  100000     1.41076    1.57629     23    17.2836
 34  100000     1.41703    1.57629     23    17.5111
 35  100000     1.42273    1.57629     23    17.7167
 36  100000     1.42868    1.57629     23    17.9315
 37  100000     1.43444    1.57629     23    18.1386
 38  100000     1.43974    1.57629     23    18.3288
 39  100000     1.44508    1.57629     23    18.5195
 40  100000     1.44985    1.57629     23    18.6889
 41  100000     1.45451    1.57629     23    18.8547
 42  100000     1.45933    1.57629     23    19.0256
 43  100000     1.46413    1.57629     23    19.1953
 44  100000     1.46863    1.57629     23    19.3540
 45  100000     1.47257    1.57629     23    19.4927
 46  100000     1.47660    1.57629     23    19.6342
 47  100000     1.48039    1.57629     23    19.7673
 48  100000     1.48375    1.57629     23    19.8848
 49  100000     1.48709    1.57629     23    20.0012
 50  100000     1.49017    1.57629     23    20.1085
 51  100000     1.49302    1.57629     23    20.2078
 52  100000     1.49603    1.57629     23    20.3126
 53  100000     1.49894    1.57629     23    20.4133
 54  100000     1.50158    1.57629     23    20.5047
 55  100000     1.50446    1.57629     23    20.6040
 56  100000     1.50709    1.57629     23    20.6945
 57  100000     1.50944    1.57629     23    20.7752
 58  100000     1.51179    1.57629     23    20.8562
 59  100000     1.51367    1.57629     23    20.9204
 60  100000     1.51577    1.57629     23    20.9926
 61  100000     1.51767    1.57629     23    21.0575
 62  100000     1.51958    1.57629     23    21.1227
 63  100000     1.52131    1.57629     23    21.1818
 64  100000     1.52298    1.57629     23    21.2389
 65  100000     1.52476    1.57629     23    21.2994
 66  100000     1.52625    1.57629     23    21.3498
 67  100000     1.52751    1.57629     23    21.3926
 68  100000     1.52895    1.57629     23    21.4416
 69  100000     1.53017    1.57629     23    21.4828
 70  100000     1.53138    1.57629     23    21.5238
 71  100000     1.53267    1.57629     23    21.5670
 72  100000     1.53375    1.57629     23    21.6033
 73  100000     1.53490    1.57629     23    21.6422
 74  100000     1.53581    1.57629     23    21.6729
 75  100000     1.53691    1.57629     23    21.7098
 76  100000     1.53784    1.57629     23    21.7411
 77  100000     1.53880    1.57629     23    21.7733
 78  100000     1.53958    1.57629     23    21.7994
 79  100000     1.54055    1.57629     23    21.8320
 80  100000     1.54142    1.57629     23    21.8610
 81  100000     1.54230    1.57629     23    21.8905
 82  100000     1.54314    1.57629     23    21.9188
 83  100000     1.54380    1.57629     23    21.9405
 84  100000     1.54466    1.57629     23    21.9692
 85  100000     1.54534    1.57629     23    21.9920
 86  100000     1.54610    1.57629     23    22.0172
 87  100000     1.54689    1.57629     23    22.0435
 88  100000     1.54754    1.57629     23    22.0651
 89  100000     1.54821    1.57629     23    22.0873
 90  100000     1.54885    1.57629     23    22.1085
 91  100000     1.54941    1.57629     23    22.1272
 92  100000     1.55009    1.57629     23    22.1495
 93  100000     1.55059    1.57629     23    22.1661
 94  100000     1.55099    1.57629     23    22.1793
 95  100000     1.55156    1.57629     23    22.1983
 96  100000     1.55210    1.57629     23    22.2160
 97  100000     1.55261    1.57629     23    22.2329
 98  100000     1.55310    1.57629     23    22.2493
 99  100000     1.55345    1.57629     23    22.2605
100  100000     1.55394    1.57629     23    22.2769
101  100000     1.55437    1.57629     23    22.2911
102  100000     1.55483    1.57629     23    22.3062
103  100000     1.55519    1.57629     23    22.3180
104  100000     1.55545    1.57629     23    22.3267
105  100000     1.55584    1.57629     23    22.3392
106  100000     1.55625    1.57629     23    22.3529
107  100000     1.55660    1.57629     23    22.3644
108  100000     1.55691    1.57629     23    22.3746
109  100000     1.55720    1.57629     23    22.3842
110  100000     1.55759    1.57629     23    22.3969
111  100000     1.55795    1.57629     23    22.4086
112  100000     1.55830    1.57629     23    22.4201
113  100000     1.55859    1.57629     23    22.4296
114  100000     1.55885    1.57629     23    22.4381
115  100000     1.55918    1.57629     23    22.4489
116  100000     1.55950    1.57629     23    22.4595
117  100000     1.55985    1.57629     23    22.4707
118  100000     1.56010    1.57629     23    22.4788
119  100000     1.56042    1.57629     23    22.4893
120  100000     1.56072    1.57629     23    22.4992
121  100000     1.56101    1.57629     23    22.5086
122  100000     1.56124    1.57629     23    22.5163
123  100000     1.56152    1.57629     23    22.5254
124  100000     1.56178    1.57629     23    22.5338
125  100000     1.56205    1.57629     23    22.5428
126  100000     1.56232    1.57629     23    22.5514
127  100000     1.56257    1.57629     23    22.5596
128  100000     1.56285    1.57629     23    22.5687
129  100000     1.56306    1.57629     23    22.5756
130  100000     1.56327    1.57629     23    22.5823
131  100000     1.56350    1.57629     23    22.5896
132  100000     1.56371    1.57629     23    22.5968
133  100000     1.56401    1.57629     23    22.6065
134  100000     1.56426    1.57629     23    22.6148
135  100000     1.56451    1.57629     23    22.6227
136  100000     1.56475    1.57629     23    22.6308
137  100000     1.56494    1.57629     23    22.6368
138  100000     1.56514    1.57629     23    22.6433
139  100000     1.56532    1.57629     23    22.6491
140  100000     1.56552    1.57629     23    22.6556
141  100000     1.56571    1.57629     23    22.6618
142  100000     1.56594    1.57629     23    22.6691
143  100000     1.56614    1.57629     23    22.6757
144  100000     1.56634    1.57629     23    22.6821
145  100000     1.56646    1.57629     23    22.6860
146  100000     1.56660    1.57629     23    22.6907
147  100000     1.56681    1.57629     23    22.6973
148  100000     1.56700    1.57629     23    22.7035
149  100000     1.56715    1.57629     23    22.7084
150  100000     1.56726    1.57629     23    22.7121
151  100000     1.56745    1.57629     23    22.7180
152  100000     1.56762    1.57629     23    22.7235
153  100000     1.56780    1.57629     23    22.7294
154  100000     1.56797    1.57629     23    22.7349
155  100000     1.56812    1.57629     23    22.7398
156  100000     1.56824    1.57629     23    22.7437
157  100000     1.56838    1.57629     23    22.7482
158  100000     1.56852    1.57629     23    22.7527
159  100000     1.56864    1.57629     23    22.7567
160  100000     1.56872    1.57629     23    22.7592
161  100000     1.56881    1.57629     23    22.7622
162  100000     1.56896    1.57629     23    22.7670
163  100000     1.56910    1.57629     23    22.7717
164  100000     1.56921    1.57629     23    22.7751
165  100000     1.56930    1.57629     23    22.7781
166  100000     1.56937    1.57629     23    22.7803
167  100000     1.56950    1.57629     23    22.7843
168  100000     1.56958    1.57629     23    22.7870
169  100000     1.56970    1.57629     23    22.7908
170  100000     1.56975    1.57629     23    22.7924
171  100000     1.56987    1.57629     23    22.7963
172  100000     1.56996    1.57629     23    22.7993
173  100000     1.57010    1.57629     23    22.8038
174  100000     1.57019    1.57629     23    22.8068
175  100000     1.57028    1.57629     23    22.8097
176  100000     1.57040    1.57629     23    22.8134
177  100000     1.57053    1.57629     23    22.8177
178  100000     1.57067    1.57629     23    22.8222
179  100000     1.57075    1.57629     23    22.8248
180  100000     1.57086    1.57629     23    22.8286
181  100000     1.57098    1.57629     23    22.8322
182  100000     1.57102    1.57629     23    22.8336
183  100000     1.57113    1.57629     23    22.8372
184  100000     1.57119    1.57629     23    22.8389
185  100000     1.57132    1.57629     23    22.8432
186  100000     1.57136    1.57629     23    22.8447
187  100000     1.57144    1.57629     23    22.8472
188  100000     1.57155    1.57629     23    22.8506
189  100000     1.57162    1.57629     23    22.8529
190  100000     1.57171    1.57629     23    22.8559
191  100000     1.57177    1.57629     23    22.8579
192  100000     1.57186    1.57629     23    22.8607
193  100000     1.57193    1.57629     23    22.8628
194  100000     1.57200    1.57629     23    22.8652
195  100000     1.57208    1.57629     23    22.8677
196  100000     1.57217    1.57629     23    22.8705
197  100000     1.57225    1.57629     23    22.8734
198  100000     1.57231    1.57629     23    22.8751
199  100000     1.57243    1.57629     23    22.8790
200  100000     1.57249    1.57629     23    22.8810
201  100000     1.57254    1.57629     23    22.8827
202  100000     1.57263    1.57629     23    22.8855
203  100000     1.57269    1.57629     23    22.8875
204  100000     1.57276    1.57629     23    22.8898
205  100000     1.57282    1.57629     23    22.8917
206  100000     1.57286    1.57629     23    22.8932
207  100000     1.57288    1.57629     23    22.8939
208  100000     1.57294    1.57629     23    22.8958
209  100000     1.57300    1.57629     23    22.8976
210  100000     1.57306    1.57629     23    22.8995
211  100000     1.57313    1.57629     23    22.9020
212  100000     1.57318    1.57629     23    22.9034
213  100000     1.57323    1.57629     23    22.9053
214  100000     1.57326    1.57629     23    22.9063
215  100000     1.57331    1.57629     23    22.9078
216  100000     1.57336    1.57629     23    22.9095
217  100000     1.57342    1.57629     23    22.9114
218  100000     1.57349    1.57629     23    22.9138
219  100000     1.57356    1.57629     23    22.9158
220  100000     1.57360    1.57629     23    22.9173
221  100000     1.57364    1.57629     23    22.9186
222  100000     1.57369    1.57629     23    22.9200
223  100000     1.57376    1.57629     23    22.9223
224  100000     1.57377    1.57629     23    22.9228
225  100000     1.57380    1.57629     23    22.9236
226  100000     1.57384    1.57629     23    22.9251
227  100000     1.57390    1.57629     23    22.9271
228  100000     1.57394    1.57629     23    22.9281
229  100000     1.57400    1.57629     23    22.9302
230  100000     1.57403    1.57629     23    22.9309
231  100000     1.57409    1.57629     23    22.9330
232  100000     1.57414    1.57629     23    22.9345
233  100000     1.57418    1.57629     23    22.9360
234  100000     1.57420    1.57629     23    22.9366
235  100000     1.57425    1.57629     23    22.9382
236  100000     1.57431    1.57629     23    22.9400
237  100000     1.57435    1.57629     23    22.9413
238  100000     1.57440    1.57629     23    22.9431
239  100000     1.57443    1.57629     23    22.9439
240  100000     1.57445    1.57629     23    22.9446
241  100000     1.57446    1.57629     23    22.9450
242  100000     1.57451    1.57629     23    22.9466
243  100000     1.57452    1.57629     23    22.9467
244  100000     1.57454    1.57629     23    22.9473
245  100000     1.57458    1.57629     23    22.9487
246  100000     1.57461    1.57629     23    22.9498
247  100000     1.57465    1.57629     23    22.9510
248  100000     1.57470    1.57629     23    22.9525
249  100000     1.57474    1.57629     23    22.9538
250  100000     1.57477    1.57629     23    22.9547
251  100000     1.57477    1.57629     23    22.9548
252  100000     1.57479    1.57629     23    22.9555
253  100000     1.57482    1.57629     23    22.9565
254  100000     1.57484    1.57629     23    22.9571
255  100000     1.57488    1.57629     23    22.9583
256  100000     1.57490    1.57629     23    22.9591
257  100000     1.57492    1.57629     23    22.9597
258  100000     1.57496    1.57629     23    22.9611
259  100000     1.57499    1.57629     23    22.9620
260  100000     1.57503    1.57629     23    22.9631
261  100000     1.57504    1.57629     23    22.9636
262  100000     1.57507    1.57629     23    22.9646
263  100000     1.57511    1.57629     23    22.9659
264  100000     1.57511    1.57629     23    22.9659
265  100000     1.57514    1.57629     23    22.9666
266  100000     1.57515    1.57629     23    22.9672
267  100000     1.57517    1.57629     23    22.9677
268  100000     1.57520    1.57629     23    22.9688
269  100000     1.57523    1.57629     23    22.9696
270  100000     1.57523    1.57629     23    22.9698
271  100000     1.57522    1.57629     23    22.9693
272  100000     1.57522    1.57629     23    22.9692
273  100000     1.57523    1.57629     23    22.9694
274  100000     1.57523    1.57629     23    22.9697
275  100000     1.57524    1.57629     23    22.9700
276  100000     1.57526    1.57629     23    22.9703
277  100000     1.57526    1.57629     23    22.9703
278  100000     1.57526    1.57629     23    22.9703
279  100000     1.57528    1.57629     23    22.9710
280  100000     1.57529    1.57629     23    22.9713
281  100000     1.57531    1.57629     23    22.9719
282  100000     1.57533    1.57629     23    22.9727
283  100000     1.57536    1.57629     23    22.9735
284  100000     1.57536    1.57629     23    22.9737
285  100000     1.57539    1.57629     23    22.9745
286  100000     1.57540    1.57629     23    22.9748
287  100000     1.57541    1.57629     23    22.9753
288  100000     1.57544    1.57629     23    22.9761
289  100000     1.57547    1.57629     23    22.9769
290  100000     1.57548    1.57629     23    22.9773
291  100000     1.57549    1.57629     23    22.9777
292  100000     1.57553    1.57629     23    22.9788
293  100000     1.57554    1.57629     23    22.9792
294  100000     1.57556    1.57629     23    22.9801
295  100000     1.57559    1.57629     23    22.9808
296  100000     1.57560    1.57629     23    22.9813
297  100000     1.57562    1.57629     23    22.9820
298  100000     1.57563    1.57629     23    22.9822
299  100000     1.57564    1.57629     23    22.9826
300  100000     1.57566    1.57629     23    22.9831
301  100000     1.57567    1.57629     23    22.9835
302  100000     1.57569    1.57629     23    22.9842
303  100000     1.57570    1.57629     23    22.9846
304  100000     1.57572    1.57629     23    22.9850
305  100000     1.57573    1.57629     23    22.9854
306  100000     1.57575    1.57629     23    22.9859
307  100000     1.57576    1.57629     23    22.9863
308  100000     1.57576    1.57629     23    22.9864
309  100000     1.57576    1.57629     23    22.9863
310  100000     1.57577    1.57629     23    22.9867
311  100000     1.57578    1.57629     23    22.9870
312  100000     1.57579    1.57629     23    22.9873
313  100000     1.57578    1.57629     23    22.9870
314  100000     1.57579    1.57629     23    22.9874
315  100000     1.57581    1.57629     23    22.9878
316  100000     1.57582    1.57629     23    22.9881
317  100000     1.57582    1.57629     23    22.9883
318  100000     1.57583    1.57629     23    22.9884
319  100000     1.57584    1.57629     23    22.9887
320  100000     1.57584    1.57629     23    22.9888
321  100000     1.57585    1.57629     23    22.9889
322  100000     1.57585    1.57629     23    22.9891
323  100000     1.57586    1.57629     23    22.9894
324  100000     1.57587    1.57629     23    22.9897
325  100000     1.57589    1.57629     23    22.9903
326  100000     1.57589    1.57629     23    22.9903
327  100000     1.57590    1.57629     23    22.9905
328  100000     1.57589    1.57629     23    22.9904
329  100000     1.57590    1.57629     23    22.9905
330  100000     1.57591    1.57629     23    22.9908
331  100000     1.57590    1.57629     23    22.9907
332  100000     1.57591    1.57629     23    22.9910
333  100000     1.57593    1.57629     23    22.9914
334  100000     1.57594    1.57629     23    22.9919
335  100000     1.57595    1.57629     23    22.9922
336  100000     1.57595    1.57629     23    22.9920
337  100000     1.57594    1.57629     23    22.9920
338  100000     1.57594    1.57629     23    22.9920
339  100000     1.57594    1.57629     23    22.9920
340  100000     1.57595    1.57629     23    22.9922
341  100000     1.57595    1.57629     23    22.9923
342  100000     1.57596    1.57629     23    22.9925
343  100000     1.57597    1.57629     23    22.9930
344  100000     1.57597    1.57629     23    22.9931
345  100000     1.57598    1.57629     23    22.9931
346  100000     1.57597    1.57629     23    22.9930
347  100000     1.57598    1.57629     23    22.9933
348  100000     1.57598    1.57629     23    22.9932
349  100000     1.57599    1.57629     23    22.9933
350  100000     1.57598    1.57629     23    22.9931
351  100000     1.57598    1.57629     23    22.9931
352  100000     1.57599    1.57629     23    22.9935
353  100000     1.57600    1.57629     23    22.9938
354  100000     1.57600    1.57629     23    22.9938
355  100000     1.57600    1.57629     23    22.9937
356  100000     1.57601    1.57629     23    22.9942
357  100000     1.57601    1.57629     23    22.9942
358  100000     1.57602    1.57629     23    22.9944
359  100000     1.57602    1.57629     23    22.9944
360  100000     1.57603    1.57629     23    22.9947
361  100000     1.57603    1.57629     23    22.9949
362  100000     1.57604    1.57629     23    22.9950
363  100000     1.57604    1.57629     23    22.9951
364  100000     1.57604    1.57629     23    22.9950
365  100000     1.57604    1.57629     23    22.9951
366  100000     1.57605    1.57629     23    22.9952
367  100000     1.57604    1.57629     23    22.9951
368  100000     1.57604    1.57629     23    22.9950
369  100000     1.57605    1.57629     23    22.9952
370  100000     1.57605    1.57629     23    22.9952
371  100000     1.57605    1.57629     23    22.9953
372  100000     1.57606    1.57629     23    22.9955
373  100000     1.57607    1.57629     23    22.9958
374  100000     1.57607    1.57629     23    22.9960
375  100000     1.57607    1.57629     23    22.9959
376  100000     1.57607    1.57629     23    22.9959
377  100000     1.57608    1.57629     23    22.9962
378  100000     1.57609    1.57629     23    22.9965
379  100000     1.57609    1.57629     23    22.9964
380  100000     1.57609    1.57629     23    22.9965
381  100000     1.57610    1.57629     23    22.9968
382  100000     1.57610    1.57629     23    22.9967
383  100000     1.57610    1.57629     23    22.9968
384  100000     1.57611    1.57629     23    22.9970
385  100000     1.57611    1.57629     23    22.9971
386  100000     1.57611    1.57629     23    22.9970
387  100000     1.57610    1.57629     23    22.9969
388  100000     1.57611    1.57629     23    22.9970
389  100000     1.57611    1.57629     23    22.9971
390  100000     1.57611    1.57629     23    22.9970
391  100000     1.57610    1.57629     23    22.9967
392  100000     1.57610    1.57629     23    22.9967
393  100000     1.57611    1.57629     23    22.9969
394  100000     1.57611    1.57629     23    22.9970
395  100000     1.57610    1.57629     23    22.9969
396  100000     1.57611    1.57629     23    22.9969
397  100000     1.57611    1.57629     23    22.9970
398  100000     1.57611    1.57629     23    22.9971
399  100000     1.57612    1.57629     23    22.9973
400  100000     1.57612    1.57629     23    22.9975

Asexual Model with selection (random number based on
relative fitness to determine who survives).  Mutation rate is      1e-08,
Recomb rate is ONE per asexual individual per generation
Starting values at generation   0 are; population size 100000,
with avg fitness of    1.22144.

gen  popSize    avgFit     maxFit maxAlleles avgAlleles
  0  100000     1.22144    1.67213     26    10.0116
  1  100000     1.22144    1.67213     26    10.0116
  2  100000     1.22609    1.67213     26    10.2024
  3  100000     1.23082    1.67213     26    10.3953
  4  100000     1.23575    1.67213     26    10.5953
  5  100000     1.24105    1.67213     26    10.8101
  6  100000     1.24614    1.67213     26    11.0153
  7  100000     1.25119    1.67213     26    11.2183
  8  100000     1.25595    1.67213     26    11.4085
  9  100000     1.26170    1.67213     26    11.6372
 10  100000     1.26699    1.67213     26    11.8469
 11  100000     1.27257    1.67213     26    12.0669
 12  100000     1.27833    1.67213     26    12.2941
 13  100000     1.28427    1.67213     26    12.5265
 14  100000     1.29014    1.67213     26    12.7550
 15  100000     1.29627    1.67213     26    12.9925
 16  100000     1.30252    1.67213     26    13.2338
 17  100000     1.30902    1.67213     26    13.4835
 18  100000     1.31541    1.67213     26    13.7273
 19  100000     1.32240    1.67213     26    13.9940
 20  100000     1.32886    1.67213     26    14.2383
 21  100000     1.33559    1.67213     26    14.4926
 22  100000     1.34242    1.67213     26    14.7487
 23  100000     1.34920    1.67213     26    15.0007
 24  100000     1.35646    1.67213     26    15.2694
 25  100000     1.36414    1.67213     26    15.5529
 26  100000     1.37178    1.67213     26    15.8328
 27  100000     1.38043    1.67213     26    16.1483
 28  100000     1.38846    1.67213     26    16.4392
 29  100000     1.39635    1.67213     26    16.7230
 30  100000     1.40542    1.67213     26    17.0489
 31  100000     1.41411    1.67213     26    17.3593
 32  100000     1.42301    1.67213     26    17.6749
 33  100000     1.43192    1.67213     26    17.9905
 34  100000     1.44073    1.67213     26    18.3003
 35  100000     1.44979    1.67213     26    18.6169
 36  100000     1.45925    1.67213     26    18.9471
 37  100000     1.46832    1.67213     26    19.2624
 38  100000     1.47740    1.67213     26    19.5759
 39  100000     1.48618    1.67213     26    19.8781
 40  100000     1.49513    1.67213     26    20.1840
 41  100000     1.50398    1.67213     26    20.4861
 42  100000     1.51258    1.67213     26    20.7787
 43  100000     1.52091    1.67213     26    21.0607
 44  100000     1.52946    1.67213     26    21.3498
 45  100000     1.53744    1.67213     26    21.6185
 46  100000     1.54506    1.67213     26    21.8741
 47  100000     1.55233    1.67213     26    22.1171
 48  100000     1.55907    1.67213     26    22.3418
 49  100000     1.56623    1.67213     26    22.5799
 50  100000     1.57247    1.67213     26    22.7868
 51  100000     1.57890    1.67213     26    23.0002
 52  100000     1.58451    1.67213     26    23.1855
 53  100000     1.58956    1.67213     26    23.3512
 54  100000     1.59472    1.67213     26    23.5208
 55  100000     1.59964    1.67213     26    23.6814
 56  100000     1.60446    1.67213     26    23.8391
 57  100000     1.60890    1.67213     26    23.9838
 58  100000     1.61270    1.67213     26    24.1072
 59  100000     1.61617    1.67213     26    24.2198
 60  100000     1.61950    1.67213     26    24.3276
 61  100000     1.62281    1.67213     26    24.4345
 62  100000     1.62589    1.67213     26    24.5339
 63  100000     1.62870    1.67213     26    24.6248
 64  100000     1.63148    1.67213     26    24.7140
 65  100000     1.63426    1.67213     26    24.8033
 66  100000     1.63655    1.67213     26    24.8766
 67  100000     1.63878    1.67213     26    24.9479
 68  100000     1.64067    1.67213     26    25.0084
 69  100000     1.64286    1.67213     26    25.0781
 70  100000     1.64448    1.67213     26    25.1298
 71  100000     1.64627    1.67213     26    25.1866
 72  100000     1.64771    1.67213     26    25.2328
 73  100000     1.64916    1.67213     26    25.2787
 74  100000     1.65058    1.67213     26    25.3237
 75  100000     1.65189    1.67213     26    25.3652
 76  100000     1.65294    1.67213     26    25.3981
 77  100000     1.65394    1.67213     26    25.4299
 78  100000     1.65489    1.67213     26    25.4599
 79  100000     1.65591    1.67213     26    25.4920
 80  100000     1.65679    1.67213     26    25.5199
 81  100000     1.65746    1.67213     26    25.5410
 82  100000     1.65831    1.67213     26    25.5679
 83  100000     1.65907    1.67213     26    25.5918
 84  100000     1.65980    1.67213     26    25.6148
 85  100000     1.66051    1.67213     26    25.6370
 86  100000     1.66121    1.67213     26    25.6591
 87  100000     1.66175    1.67213     26    25.6760
 88  100000     1.66236    1.67213     26    25.6951
 89  100000     1.66278    1.67213     26    25.7084
 90  100000     1.66311    1.67213     26    25.7187
 91  100000     1.66371    1.67213     26    25.7375
 92  100000     1.66411    1.67213     26    25.7502
 93  100000     1.66454    1.67213     26    25.7633
 94  100000     1.66489    1.67213     26    25.7745
 95  100000     1.66528    1.67213     26    25.7867
 96  100000     1.66567    1.67213     26    25.7987
 97  100000     1.66606    1.67213     26    25.8109
 98  100000     1.66651    1.67213     26    25.8252
 99  100000     1.66684    1.67213     26    25.8353
100  100000     1.66711    1.67213     26    25.8437
101  100000     1.66741    1.67213     26    25.8530
102  100000     1.66768    1.67213     26    25.8616
103  100000     1.66793    1.67213     26    25.8693
104  100000     1.66814    1.67213     26    25.8758
105  100000     1.66831    1.67213     26    25.8811
106  100000     1.66847    1.67213     26    25.8862
107  100000     1.66871    1.67213     26    25.8936
108  100000     1.66887    1.67213     26    25.8986
109  100000     1.66913    1.67213     26    25.9068
110  100000     1.66924    1.67213     26    25.9102
111  100000     1.66940    1.67213     26    25.9152
112  100000     1.66951    1.67213     26    25.9185
113  100000     1.66971    1.67213     26    25.9246
114  100000     1.66984    1.67213     26    25.9289
115  100000     1.66992    1.67213     26    25.9314
116  100000     1.67004    1.67213     26    25.9352
117  100000     1.67019    1.67213     26    25.9399
118  100000     1.67024    1.67213     26    25.9412
119  100000     1.67036    1.67213     26    25.9450
120  100000     1.67042    1.67213     26    25.9469
121  100000     1.67051    1.67213     26    25.9497
122  100000     1.67062    1.67213     26    25.9530
123  100000     1.67071    1.67213     26    25.9559
124  100000     1.67079    1.67213     26    25.9583
125  100000     1.67089    1.67213     26    25.9614
126  100000     1.67092    1.67213     26    25.9625
127  100000     1.67099    1.67213     26    25.9647
128  100000     1.67106    1.67213     26    25.9668
129  100000     1.67110    1.67213     26    25.9682
130  100000     1.67122    1.67213     26    25.9718
131  100000     1.67127    1.67213     26    25.9733
132  100000     1.67132    1.67213     26    25.9749
133  100000     1.67137    1.67213     26    25.9763
134  100000     1.67138    1.67213     26    25.9767
135  100000     1.67141    1.67213     26    25.9776
136  100000     1.67144    1.67213     26    25.9786
137  100000     1.67149    1.67213     26    25.9802
138  100000     1.67153    1.67213     26    25.9815
139  100000     1.67157    1.67213     26    25.9828
140  100000     1.67160    1.67213     26    25.9836
141  100000     1.67165    1.67213     26    25.9852
142  100000     1.67166    1.67213     26    25.9854
143  100000     1.67169    1.67213     26    25.9864
144  100000     1.67172    1.67213     26    25.9872
145  100000     1.67174    1.67213     26    25.9878
146  100000     1.67175    1.67213     26    25.9884
147  100000     1.67176    1.67213     26    25.9886
148  100000     1.67179    1.67213     26    25.9894
149  100000     1.67179    1.67213     26    25.9894
150  100000     1.67179    1.67213     26    25.9894
151  100000     1.67181    1.67213     26    25.9900
152  100000     1.67183    1.67213     26    25.9908
153  100000     1.67185    1.67213     26    25.9914
154  100000     1.67187    1.67213     26    25.9919
155  100000     1.67189    1.67213     26    25.9925
156  100000     1.67192    1.67213     26    25.9935
157  100000     1.67193    1.67213     26    25.9938
158  100000     1.67194    1.67213     26    25.9941
159  100000     1.67195    1.67213     26    25.9944
160  100000     1.67195    1.67213     26    25.9945
161  100000     1.67197    1.67213     26    25.9949
162  100000     1.67197    1.67213     26    25.9951
163  100000     1.67198    1.67213     26    25.9953
164  100000     1.67200    1.67213     26    25.9959
165  100000     1.67199    1.67213     26    25.9957
166  100000     1.67200    1.67213     26    25.9961
167  100000     1.67201    1.67213     26    25.9964
168  100000     1.67202    1.67213     26    25.9965
169  100000     1.67203    1.67213     26    25.9970
170  100000     1.67204    1.67213     26    25.9972
171  100000     1.67205    1.67213     26    25.9974
172  100000     1.67204    1.67213     26    25.9973
173  100000     1.67205    1.67213     26    25.9976
174  100000     1.67205    1.67213     26    25.9974
175  100000     1.67205    1.67213     26    25.9974
176  100000     1.67205    1.67213     26    25.9975
177  100000     1.67205    1.67213     26    25.9974
178  100000     1.67205    1.67213     26    25.9976
179  100000     1.67206    1.67213     26    25.9979
180  100000     1.67206    1.67213     26    25.9979
181  100000     1.67207    1.67213     26    25.9981
182  100000     1.67207    1.67213     26    25.9980
183  100000     1.67208    1.67213     26    25.9983
184  100000     1.67207    1.67213     26    25.9982
185  100000     1.67207    1.67213     26    25.9981
186  100000     1.67208    1.67213     26    25.9984
187  100000     1.67209    1.67213     26    25.9987
188  100000     1.67209    1.67213     26    25.9988
189  100000     1.67209    1.67213     26    25.9987
190  100000     1.67209    1.67213     26    25.9986
191  100000     1.67209    1.67213     26    25.9987
192  100000     1.67209    1.67213     26    25.9986
193  100000     1.67209    1.67213     26    25.9988
194  100000     1.67209    1.67213     26    25.9987
195  100000     1.67209    1.67213     26    25.9987
196  100000     1.67209    1.67213     26    25.9988
197  100000     1.67209    1.67213     26    25.9988
198  100000     1.67209    1.67213     26    25.9988
199  100000     1.67210    1.67213     26    25.9990
200  100000     1.67210    1.67213     26    25.9991
201  100000     1.67210    1.67213     26    25.9990
202  100000     1.67210    1.67213     26    25.9990
203  100000     1.67210    1.67213     26    25.9990
204  100000     1.67211    1.67213     26    25.9992
205  100000     1.67211    1.67213     26    25.9993
206  100000     1.67211    1.67213     26    25.9993
207  100000     1.67211    1.67213     26    25.9993
208  100000     1.67211    1.67213     26    25.9994
209  100000     1.67211    1.67213     26    25.9994
210  100000     1.67211    1.67213     26    25.9994
211  100000     1.67211    1.67213     26    25.9994
212  100000     1.67211    1.67213     26    25.9995
213  100000     1.67212    1.67213     26    25.9996
214  100000     1.67212    1.67213     26    25.9997
215  100000     1.67212    1.67213     26    25.9997
216  100000     1.67212    1.67213     26    25.9997
217  100000     1.67212    1.67213     26    25.9997
218  100000     1.67212    1.67213     26    25.9998
219  100000     1.67213    1.67213     26    25.9998
220  100000     1.67213    1.67213     26    25.9998
221  100000     1.67213    1.67213     26    25.9999
222  100000     1.67213    1.67213     26    25.9999
223  100000     1.67213    1.67213     26    25.9999
224  100000     1.67213    1.67213     26    25.9998
225  100000     1.67213    1.67213     26    25.9999
226  100000     1.67213    1.67213     26    25.9999
227  100000     1.67213    1.67213     26    25.9999
228  100000     1.67213    1.67213     26    25.9999
229  100000     1.67213    1.67213     26    25.9999
230  100000     1.67213    1.67213     26    25.9999
231  100000     1.67213    1.67213     26    25.9999
232  100000     1.67213    1.67213     26    25.9999
233  100000     1.67213    1.67213     26    25.9999
234  100000     1.67213    1.67213     26    25.9998
235  100000     1.67213    1.67213     26    25.9998
236  100000     1.67213    1.67213     26    25.9999
237  100000     1.67213    1.67213     26    25.9998
238  100000     1.67213    1.67213     26    25.9999
239  100000     1.67213    1.67213     26    25.9999
240  100000     1.67213    1.67213     26    25.9999
241  100000     1.67213    1.67213     26    25.9999
242  100000     1.67213    1.67213     26    25.9999
243  100000     1.67213    1.67213     26    26.0000
244  100000     1.67213    1.67213     26    26.0000
245  100000     1.67213    1.67213     26    26.0000
246  100000     1.67213    1.67213     26    26.0000
247  100000     1.67213    1.67213     26    26.0000
248  100000     1.67213    1.67213     26    26.0000
249  100000     1.67213    1.67213     26    26.0000
250  100000     1.67213    1.67213     26    26.0000
251  100000     1.67213    1.67213     26    26.0000
252  100000     1.67213    1.67213     26    26.0000
253  100000     1.67213    1.67213     26    26.0000
254  100000     1.67213    1.67213     26    26.0000
255  100000     1.67213    1.67213     26    26.0000
256  100000     1.67213    1.67213     26    26.0000
257  100000     1.67213    1.67213     26    26.0000
258  100000     1.67213    1.67213     26    26.0000
259  100000     1.67213    1.67213     26    26.0000
260  100000     1.67213    1.67213     26    26.0000
261  100000     1.67213    1.67213     26    26.0000
262  100000     1.67213    1.67213     26    26.0000
263  100000     1.67213    1.67213     26    26.0000
264  100000     1.67213    1.67213     26    26.0000
265  100000     1.67213    1.67213     26    26.0000
266  100000     1.67213    1.67213     26    26.0000
267  100000     1.67213    1.67213     26    26.0000
268  100000     1.67213    1.67213     26    26.0000
269  100000     1.67213    1.67213     26    26.0000
270  100000     1.67213    1.67213     26    26.0000
271  100000     1.67213    1.67213     26    26.0000
272  100000     1.67213    1.67213     26    26.0000
273  100000     1.67213    1.67213     26    26.0000
274  100000     1.67213    1.67213     26    26.0000
275  100000     1.67213    1.67213     26    26.0000
276  100000     1.67213    1.67213     26    26.0000
277  100000     1.67213    1.67213     26    26.0000
278  100000     1.67213    1.67213     26    26.0000
279  100000     1.67213    1.67213     26    26.0000
280  100000     1.67213    1.67213     26    26.0000
281  100000     1.67213    1.67213     26    26.0000
282  100000     1.67213    1.67213     26    26.0000
283  100000     1.67213    1.67213     26    26.0000
284  100000     1.67213    1.67213     26    26.0000
285  100000     1.67213    1.67213     26    26.0000
286  100000     1.67213    1.67213     26    26.0000
287  100000     1.67213    1.67213     26    26.0000
288  100000     1.67213    1.67213     26    26.0000
289  100000     1.67213    1.67213     26    26.0000
290  100000     1.67213    1.67213     26    26.0000
291  100000     1.67213    1.67213     26    26.0000
292  100000     1.67213    1.67213     26    26.0000
293  100000     1.67213    1.67213     26    26.0000
294  100000     1.67213    1.67213     26    26.0000
295  100000     1.67213    1.67213     26    26.0000
296  100000     1.67213    1.67213     26    26.0000
297  100000     1.67213    1.67213     26    26.0000
298  100000     1.67213    1.67213     26    26.0000
299  100000     1.67213    1.67213     26    26.0000
300  100000     1.67213    1.67213     26    26.0000
301  100000     1.67213    1.67213     26    26.0000
302  100000     1.67213    1.67213     26    26.0000
303  100000     1.67213    1.67213     26    26.0000
304  100000     1.67213    1.67213     26    26.0000
305  100000     1.67213    1.67213     26    26.0000
306  100000     1.67213    1.67213     26    26.0000
307  100000     1.67213    1.67213     26    26.0000
308  100000     1.67213    1.67213     26    26.0000
309  100000     1.67213    1.67213     26    26.0000
310  100000     1.67213    1.67213     26    26.0000
311  100000     1.67213    1.67213     26    26.0000
312  100000     1.67213    1.67213     26    26.0000
313  100000     1.67213    1.67213     26    26.0000
314  100000     1.67213    1.67213     26    26.0000
315  100000     1.67213    1.67213     26    26.0000
316  100000     1.67213    1.67213     26    26.0000
317  100000     1.67213    1.67213     26    26.0000
318  100000     1.67213    1.67213     26    26.0000
319  100000     1.67213    1.67213     26    26.0000
320  100000     1.67213    1.67213     26    26.0000
321  100000     1.67213    1.67213     26    26.0000
322  100000     1.67213    1.67213     26    26.0000
323  100000     1.67213    1.67213     26    26.0000
324  100000     1.67213    1.67213     26    26.0000
325  100000     1.67213    1.67213     26    26.0000
326  100000     1.67213    1.67213     26    26.0000
327  100000     1.67213    1.67213     26    26.0000
328  100000     1.67213    1.67213     26    26.0000
329  100000     1.67213    1.67213     26    26.0000
330  100000     1.67213    1.67213     26    26.0000
331  100000     1.67213    1.67213     26    26.0000
332  100000     1.67213    1.67213     26    26.0000
333  100000     1.67213    1.67213     26    26.0000
334  100000     1.67213    1.67213     26    26.0000
335  100000     1.67213    1.67213     26    26.0000
336  100000     1.67213    1.67213     26    26.0000
337  100000     1.67213    1.67213     26    26.0000
338  100000     1.67213    1.67213     26    26.0000
339  100000     1.67213    1.67213     26    26.0000
340  100000     1.67213    1.67213     26    26.0000
341  100000     1.67213    1.67213     26    26.0000
342  100000     1.67213    1.67213     26    26.0000
343  100000     1.67213    1.67213     26    26.0000
344  100000     1.67213    1.67213     26    26.0000
345  100000     1.67213    1.67213     26    26.0000
346  100000     1.67213    1.67213     26    26.0000
347  100000     1.67213    1.67213     26    26.0000
348  100000     1.67213    1.67213     26    26.0000
349  100000     1.67213    1.67213     26    26.0000
350  100000     1.67213    1.67213     26    26.0000
351  100000     1.67213    1.67213     26    26.0000
352  100000     1.67213    1.67213     26    26.0000
353  100000     1.67213    1.67213     26    26.0000
354  100000     1.67213    1.67213     26    26.0000
355  100000     1.67213    1.67213     26    26.0000
356  100000     1.67213    1.67213     26    26.0000
357  100000     1.67213    1.67213     26    26.0000
358  100000     1.67213    1.67213     26    26.0000
359  100000     1.67213    1.67213     26    26.0000
360  100000     1.67213    1.67213     26    26.0000
361  100000     1.67213    1.67213     26    26.0000
362  100000     1.67213    1.67213     26    26.0000
363  100000     1.67213    1.67213     26    26.0000
364  100000     1.67213    1.67213     26    26.0000
365  100000     1.67213    1.67213     26    26.0000
366  100000     1.67213    1.67213     26    26.0000
367  100000     1.67213    1.67213     26    26.0000
368  100000     1.67213    1.67213     26    26.0000
369  100000     1.67213    1.67213     26    26.0000
370  100000     1.67213    1.67213     26    26.0000
371  100000     1.67213    1.67213     26    26.0000
372  100000     1.67213    1.67213     26    26.0000
373  100000     1.67213    1.67213     26    26.0000
374  100000     1.67213    1.67213     26    26.0000
375  100000     1.67213    1.67213     26    26.0000
376  100000     1.67213    1.67213     26    26.0000
377  100000     1.67213    1.67213     26    26.0000
378  100000     1.67213    1.67213     26    26.0000
379  100000     1.67213    1.67213     26    26.0000
380  100000     1.67213    1.67213     26    26.0000
381  100000     1.67213    1.67213     26    26.0000
382  100000     1.67213    1.67213     26    26.0000
383  100000     1.67213    1.67213     26    26.0000
384  100000     1.67213    1.67213     26    26.0000
385  100000     1.67213    1.67213     26    26.0000
386  100000     1.67213    1.67213     26    26.0000
387  100000     1.67213    1.67213     26    26.0000
388  100000     1.67213    1.67213     26    26.0000
389  100000     1.67213    1.67213     26    26.0000
390  100000     1.67213    1.67213     26    26.0000
391  100000     1.67213    1.67213     26    26.0000
392  100000     1.67213    1.67213     26    26.0000
393  100000     1.67213    1.67213     26    26.0000
394  100000     1.67213    1.67213     26    26.0000
395  100000     1.67213    1.67213     26    26.0000
396  100000     1.67213    1.67213     26    26.0000
397  100000     1.67213    1.67213     26    26.0000
398  100000     1.67213    1.67213     26    26.0000
399  100000     1.67213    1.67213     26    26.0000
400  100000     1.67213    1.67213     26    26.0000

Asexual Model with selection (random number based on
relative fitness to determine who survives).  Mutation rate is      1e-08,
Recomb rate is ONE per asexual individual per generation
Starting values at generation   0 are; population size 100000,
with avg fitness of    1.22112.

gen  popSize    avgFit     maxFit maxAlleles avgAlleles
  0  100000     1.22112    1.73969     28     9.9979
  1  100000     1.22112    1.73969     28     9.9979
  2  100000     1.22563    1.73969     28    10.1827
  3  100000     1.23042    1.73969     28    10.3788
  4  100000     1.23496    1.73969     28    10.5633
  5  100000     1.23997    1.73969     28    10.7662
  6  100000     1.24503    1.73969     28    10.9704
  7  100000     1.25026    1.73969     28    11.1804
  8  100000     1.25570    1.73969     28    11.3975
  9  100000     1.26112    1.73969     28    11.6133
 10  100000     1.26648    1.73969     28    11.8259
 11  100000     1.27187    1.73969     28    12.0381
 12  100000     1.27785    1.73969     28    12.2727
 13  100000     1.28413    1.73969     28    12.5183
 14  100000     1.29039    1.73969     28    12.7611
 15  100000     1.29673    1.73969     28    13.0060
 16  100000     1.30313    1.73969     28    13.2513
 17  100000     1.30986    1.73969     28    13.5083
 18  100000     1.31651    1.73969     28    13.7604
 19  100000     1.32371    1.73969     28    14.0314
 20  100000     1.33119    1.73969     28    14.3110
 21  100000     1.33901    1.73969     28    14.6011
 22  100000     1.34742    1.73969     28    14.9102
 23  100000     1.35582    1.73969     28    15.2169
 24  100000     1.36548    1.73969     28    15.5664
 25  100000     1.37536    1.73969     28    15.9232
 26  100000     1.38607    1.73969     28    16.3043
 27  100000     1.39802    1.73969     28    16.7274
 28  100000     1.40977    1.73969     28    17.1398
 29  100000     1.42277    1.73969     28    17.5925
 30  100000     1.43557    1.73969     28    18.0365
 31  100000     1.44950    1.73969     28    18.5155
 32  100000     1.46413    1.73969     28    19.0145
 33  100000     1.47994    1.73969     28    19.5501
 34  100000     1.49554    1.73969     28    20.0747
 35  100000     1.51110    1.73969     28    20.5957
 36  100000     1.52784    1.73969     28    21.1539
 37  100000     1.54462    1.73969     28    21.7108
 38  100000     1.56081    1.73969     28    22.2461
 39  100000     1.57702    1.73969     28    22.7799
 40  100000     1.59233    1.73969     28    23.2816
 41  100000     1.60700    1.73969     28    23.7612
 42  100000     1.61985    1.73969     28    24.1789
 43  100000     1.63190    1.73969     28    24.5698
 44  100000     1.64332    1.73969     28    24.9398
 45  100000     1.65408    1.73969     28    25.2872
 46  100000     1.66389    1.73969     28    25.6031
 47  100000     1.67231    1.73969     28    25.8736
 48  100000     1.68025    1.73969     28    26.1281
 49  100000     1.68733    1.73969     28    26.3543
 50  100000     1.69327    1.73969     28    26.5441
 51  100000     1.69815    1.73969     28    26.6994
 52  100000     1.70298    1.73969     28    26.8534
 53  100000     1.70706    1.73969     28    26.9825
 54  100000     1.71066    1.73969     28    27.0961
 55  100000     1.71399    1.73969     28    27.2013
 56  100000     1.71693    1.73969     28    27.2939
 57  100000     1.71939    1.73969     28    27.3714
 58  100000     1.72134    1.73969     28    27.4323
 59  100000     1.72351    1.73969     28    27.5006
 60  100000     1.72528    1.73969     28    27.5558
 61  100000     1.72666    1.73969     28    27.5989
 62  100000     1.72795    1.73969     28    27.6395
 63  100000     1.72902    1.73969     28    27.6726
 64  100000     1.73000    1.73969     28    27.7030
 65  100000     1.73079    1.73969     28    27.7278
 66  100000     1.73171    1.73969     28    27.7561
 67  100000     1.73253    1.73969     28    27.7815
 68  100000     1.73317    1.73969     28    27.8014
 69  100000     1.73362    1.73969     28    27.8152
 70  100000     1.73402    1.73969     28    27.8276
 71  100000     1.73446    1.73969     28    27.8412
 72  100000     1.73478    1.73969     28    27.8512
 73  100000     1.73519    1.73969     28    27.8637
 74  100000     1.73547    1.73969     28    27.8723
 75  100000     1.73574    1.73969     28    27.8805
 76  100000     1.73611    1.73969     28    27.8920
 77  100000     1.73635    1.73969     28    27.8992
 78  100000     1.73660    1.73969     28    27.9071
 79  100000     1.73681    1.73969     28    27.9134
 80  100000     1.73702    1.73969     28    27.9197
 81  100000     1.73719    1.73969     28    27.9248
 82  100000     1.73739    1.73969     28    27.9309
 83  100000     1.73756    1.73969     28    27.9361
 84  100000     1.73765    1.73969     28    27.9389
 85  100000     1.73778    1.73969     28    27.9429
 86  100000     1.73790    1.73969     28    27.9464
 87  100000     1.73803    1.73969     28    27.9505
 88  100000     1.73814    1.73969     28    27.9538
 89  100000     1.73822    1.73969     28    27.9562
 90  100000     1.73831    1.73969     28    27.9589
 91  100000     1.73837    1.73969     28    27.9607
 92  100000     1.73846    1.73969     28    27.9632
 93  100000     1.73854    1.73969     28    27.9659
 94  100000     1.73858    1.73969     28    27.9670
 95  100000     1.73862    1.73969     28    27.9683
 96  100000     1.73868    1.73969     28    27.9699
 97  100000     1.73873    1.73969     28    27.9714
 98  100000     1.73876    1.73969     28    27.9723
 99  100000     1.73883    1.73969     28    27.9744
100  100000     1.73885    1.73969     28    27.9751
101  100000     1.73887    1.73969     28    27.9757
102  100000     1.73890    1.73969     28    27.9765
103  100000     1.73898    1.73969     28    27.9790
104  100000     1.73900    1.73969     28    27.9796
105  100000     1.73905    1.73969     28    27.9811
106  100000     1.73908    1.73969     28    27.9819
107  100000     1.73912    1.73969     28    27.9833
108  100000     1.73914    1.73969     28    27.9837
109  100000     1.73917    1.73969     28    27.9846
110  100000     1.73918    1.73969     28    27.9849
111  100000     1.73919    1.73969     28    27.9853
112  100000     1.73923    1.73969     28    27.9865
113  100000     1.73925    1.73969     28    27.9870
114  100000     1.73927    1.73969     28    27.9875
115  100000     1.73930    1.73969     28    27.9885
116  100000     1.73935    1.73969     28    27.9899
117  100000     1.73936    1.73969     28    27.9904
118  100000     1.73938    1.73969     28    27.9910
119  100000     1.73938    1.73969     28    27.9910
120  100000     1.73939    1.73969     28    27.9913
121  100000     1.73939    1.73969     28    27.9913
122  100000     1.73941    1.73969     28    27.9919
123  100000     1.73942    1.73969     28    27.9921
124  100000     1.73944    1.73969     28    27.9927
125  100000     1.73945    1.73969     28    27.9931
126  100000     1.73945    1.73969     28    27.9929
127  100000     1.73946    1.73969     28    27.9934
128  100000     1.73948    1.73969     28    27.9938
129  100000     1.73948    1.73969     28    27.9939
130  100000     1.73950    1.73969     28    27.9944
131  100000     1.73952    1.73969     28    27.9950
132  100000     1.73952    1.73969     28    27.9952
133  100000     1.73951    1.73969     28    27.9948
134  100000     1.73954    1.73969     28    27.9956
135  100000     1.73954    1.73969     28    27.9957
136  100000     1.73955    1.73969     28    27.9961
137  100000     1.73955    1.73969     28    27.9960
138  100000     1.73957    1.73969     28    27.9965
139  100000     1.73958    1.73969     28    27.9969
140  100000     1.73958    1.73969     28    27.9969
141  100000     1.73959    1.73969     28    27.9971
142  100000     1.73959    1.73969     28    27.9973
143  100000     1.73960    1.73969     28    27.9974
144  100000     1.73960    1.73969     28    27.9975
145  100000     1.73961    1.73969     28    27.9976
146  100000     1.73961    1.73969     28    27.9978
147  100000     1.73962    1.73969     28    27.9980
148  100000     1.73962    1.73969     28    27.9980
149  100000     1.73961    1.73969     28    27.9978
150  100000     1.73962    1.73969     28    27.9980
151  100000     1.73962    1.73969     28    27.9981
152  100000     1.73963    1.73969     28    27.9984
153  100000     1.73963    1.73969     28    27.9984
154  100000     1.73963    1.73969     28    27.9984
155  100000     1.73964    1.73969     28    27.9986
156  100000     1.73964    1.73969     28    27.9988
157  100000     1.73965    1.73969     28    27.9989
158  100000     1.73965    1.73969     28    27.9988
159  100000     1.73965    1.73969     28    27.9988
160  100000     1.73965    1.73969     28    27.9988
161  100000     1.73964    1.73969     28    27.9987
162  100000     1.73964    1.73969     28    27.9987
163  100000     1.73964    1.73969     28    27.9988
164  100000     1.73964    1.73969     28    27.9988
165  100000     1.73964    1.73969     28    27.9986
166  100000     1.73964    1.73969     28    27.9986
167  100000     1.73963    1.73969     28    27.9985
168  100000     1.73964    1.73969     28    27.9986
169  100000     1.73964    1.73969     28    27.9986
170  100000     1.73964    1.73969     28    27.9986
171  100000     1.73964    1.73969     28    27.9987
172  100000     1.73964    1.73969     28    27.9988
173  100000     1.73965    1.73969     28    27.9990
174  100000     1.73966    1.73969     28    27.9991
175  100000     1.73966    1.73969     28    27.9992
176  100000     1.73965    1.73969     28    27.9991
177  100000     1.73965    1.73969     28    27.9991
178  100000     1.73965    1.73969     28    27.9990
179  100000     1.73966    1.73969     28    27.9991
180  100000     1.73966    1.73969     28    27.9992
181  100000     1.73966    1.73969     28    27.9993
182  100000     1.73966    1.73969     28    27.9993
183  100000     1.73966    1.73969     28    27.9993
184  100000     1.73967    1.73969     28    27.9995
185  100000     1.73967    1.73969     28    27.9994
186  100000     1.73967    1.73969     28    27.9995
187  100000     1.73967    1.73969     28    27.9996
188  100000     1.73968    1.73969     28    27.9997
189  100000     1.73968    1.73969     28    27.9997
190  100000     1.73968    1.73969     28    27.9997
191  100000     1.73968    1.73969     28    27.9998
192  100000     1.73968    1.73969     28    27.9998
193  100000     1.73968    1.73969     28    27.9998
194  100000     1.73968    1.73969     28    27.9998
195  100000     1.73968    1.73969     28    27.9998
196  100000     1.73968    1.73969     28    27.9999
197  100000     1.73968    1.73969     28    27.9999
198  100000     1.73968    1.73969     28    27.9999
199  100000     1.73968    1.73969     28    27.9999
200  100000     1.73968    1.73969     28    28.0000
201  100000     1.73969    1.73969     28    28.0000
202  100000     1.73969    1.73969     28    28.0000
203  100000     1.73969    1.73969     28    28.0000
204  100000     1.73969    1.73969     28    28.0000
205  100000     1.73969    1.73969     28    28.0000
206  100000     1.73969    1.73969     28    28.0000
207  100000     1.73969    1.73969     28    28.0000
208  100000     1.73969    1.73969     28    28.0000
209  100000     1.73969    1.73969     28    28.0000
210  100000     1.73969    1.73969     28    28.0000
211  100000     1.73969    1.73969     28    28.0000
212  100000     1.73969    1.73969     28    28.0000
213  100000     1.73969    1.73969     28    28.0000
214  100000     1.73969    1.73969     28    28.0000
215  100000     1.73969    1.73969     28    28.0000
216  100000     1.73969    1.73969     28    28.0000
217  100000     1.73969    1.73969     28    28.0000
218  100000     1.73968    1.73969     28    28.0000
219  100000     1.73969    1.73969     28    28.0000
220  100000     1.73969    1.73969     28    28.0000
221  100000     1.73969    1.73969     28    28.0000
222  100000     1.73969    1.73969     28    28.0000
223  100000     1.73969    1.73969     28    28.0000
224  100000     1.73969    1.73969     28    28.0000
225  100000     1.73969    1.73969     28    28.0000
226  100000     1.73969    1.73969     28    28.0000
227  100000     1.73969    1.73969     28    28.0000
228  100000     1.73969    1.73969     28    28.0000
229  100000     1.73969    1.73969     28    28.0000
230  100000     1.73969    1.73969     28    28.0000
231  100000     1.73969    1.73969     28    28.0000
232  100000     1.73969    1.73969     28    28.0000
233  100000     1.73969    1.73969     28    28.0000
234  100000     1.73969    1.73969     28    28.0000
235  100000     1.73969    1.73969     28    28.0000
236  100000     1.73969    1.73969     28    28.0000
237  100000     1.73969    1.73969     28    28.0000
238  100000     1.73969    1.73969     28    28.0000
239  100000     1.73969    1.73969     28    28.0000
240  100000     1.73969    1.73969     28    28.0000
241  100000     1.73969    1.73969     28    28.0000
242  100000     1.73969    1.73969     28    28.0000
243  100000     1.73969    1.73969     28    28.0000
244  100000     1.73969    1.73969     28    28.0000
245  100000     1.73969    1.73969     28    28.0000
246  100000     1.73969    1.73969     28    28.0000
247  100000     1.73969    1.73969     28    28.0000
248  100000     1.73969    1.73969     28    28.0000
249  100000     1.73969    1.73969     28    28.0000
250  100000     1.73969    1.73969     28    28.0000
251  100000     1.73969    1.73969     28    28.0000
252  100000     1.73969    1.73969     28    28.0000
253  100000     1.73969    1.73969     28    28.0000
254  100000     1.73969    1.73969     28    28.0000
255  100000     1.73969    1.73969     28    28.0000
256  100000     1.73969    1.73969     28    28.0000
257  100000     1.73969    1.73969     28    28.0000
258  100000     1.73969    1.73969     28    28.0000
259  100000     1.73969    1.73969     28    28.0000
260  100000     1.73969    1.73969     28    28.0000
261  100000     1.73969    1.73969     28    28.0000
262  100000     1.73969    1.73969     28    28.0000
263  100000     1.73969    1.73969     28    28.0000
264  100000     1.73969    1.73969     28    28.0000
265  100000     1.73969    1.73969     28    28.0000
266  100000     1.73969    1.73969     28    28.0000
267  100000     1.73969    1.73969     28    28.0000
268  100000     1.73969    1.73969     28    28.0000
269  100000     1.73969    1.73969     28    28.0000
270  100000     1.73969    1.73969     28    28.0000
271  100000     1.73969    1.73969     28    28.0000
272  100000     1.73969    1.73969     28    28.0000
273  100000     1.73969    1.73969     28    28.0000
274  100000     1.73969    1.73969     28    28.0000
275  100000     1.73969    1.73969     28    28.0000
276  100000     1.73969    1.73969     28    28.0000
277  100000     1.73969    1.73969     28    28.0000
278  100000     1.73969    1.73969     28    28.0000
279  100000     1.73969    1.73969     28    28.0000
280  100000     1.73969    1.73969     28    28.0000
281  100000     1.73969    1.73969     28    28.0000
282  100000     1.73969    1.73969     28    28.0000
283  100000     1.73969    1.73969     28    28.0000
284  100000     1.73969    1.73969     28    28.0000
285  100000     1.73969    1.73969     28    28.0000
286  100000     1.73969    1.73969     28    28.0000
287  100000     1.73969    1.73969     28    28.0000
288  100000     1.73969    1.73969     28    28.0000
289  100000     1.73969    1.73969     28    28.0000
290  100000     1.73969    1.73969     28    28.0000
291  100000     1.73969    1.73969     28    28.0000
292  100000     1.73969    1.73969     28    28.0000
293  100000     1.73969    1.73969     28    28.0000
294  100000     1.73969    1.73969     28    28.0000
295  100000     1.73969    1.73969     28    28.0000
296  100000     1.73969    1.73969     28    28.0000
297  100000     1.73969    1.73969     28    28.0000
298  100000     1.73969    1.73969     28    28.0000
299  100000     1.73969    1.73969     28    28.0000
300  100000     1.73969    1.73969     28    28.0000
301  100000     1.73969    1.73969     28    28.0000
302  100000     1.73969    1.73969     28    28.0000
303  100000     1.73969    1.73969     28    28.0000
304  100000     1.73969    1.73969     28    28.0000
305  100000     1.73969    1.73969     28    28.0000
306  100000     1.73969    1.73969     28    28.0000
307  100000     1.73969    1.73969     28    28.0000
308  100000     1.73969    1.73969     28    28.0000
309  100000     1.73969    1.73969     28    28.0000
310  100000     1.73969    1.73969     28    28.0000
311  100000     1.73969    1.73969     28    28.0000
312  100000     1.73969    1.73969     28    28.0000
313  100000     1.73969    1.73969     28    28.0000
314  100000     1.73969    1.73969     28    28.0000
315  100000     1.73969    1.73969     28    28.0000
316  100000     1.73969    1.73969     28    28.0000
317  100000     1.73969    1.73969     28    28.0000
318  100000     1.73969    1.73969     28    28.0000
319  100000     1.73969    1.73969     28    28.0000
320  100000     1.73969    1.73969     28    28.0000
321  100000     1.73969    1.73969     28    28.0000
322  100000     1.73969    1.73969     28    28.0000
323  100000     1.73969    1.73969     28    28.0000
324  100000     1.73969    1.73969     28    28.0000
325  100000     1.73969    1.73969     28    28.0000
326  100000     1.73969    1.73969     28    28.0000
327  100000     1.73969    1.73969     28    28.0000
328  100000     1.73969    1.73969     28    28.0000
329  100000     1.73969    1.73969     28    28.0000
330  100000     1.73969    1.73969     28    28.0000
331  100000     1.73969    1.73969     28    28.0000
332  100000     1.73969    1.73969     28    28.0000
333  100000     1.73969    1.73969     28    28.0000
334  100000     1.73969    1.73969     28    28.0000
335  100000     1.73969    1.73969     28    28.0000
336  100000     1.73969    1.73969     28    28.0000
337  100000     1.73969    1.73969     28    28.0000
338  100000     1.73969    1.73969     28    28.0000
339  100000     1.73969    1.73969     28    28.0000
340  100000     1.73969    1.73969     28    28.0000
341  100000     1.73969    1.73969     28    28.0000
342  100000     1.73969    1.73969     28    28.0000
343  100000     1.73969    1.73969     28    28.0000
344  100000     1.73969    1.73969     28    28.0000
345  100000     1.73969    1.73969     28    28.0000
346  100000     1.73969    1.73969     28    28.0000
347  100000     1.73969    1.73969     28    28.0000
348  100000     1.73969    1.73969     28    28.0000
349  100000     1.73969    1.73969     28    28.0000
350  100000     1.73969    1.73969     28    28.0000
351  100000     1.73969    1.73969     28    28.0000
352  100000     1.73969    1.73969     28    28.0000
353  100000     1.73969    1.73969     28    28.0000
354  100000     1.73969    1.73969     28    28.0000
355  100000     1.73969    1.73969     28    28.0000
356  100000     1.73969    1.73969     28    28.0000
357  100000     1.73969    1.73969     28    28.0000
358  100000     1.73969    1.73969     28    28.0000
359  100000     1.73969    1.73969     28    28.0000
360  100000     1.73969    1.73969     28    28.0000
361  100000     1.73969    1.73969     28    28.0000
362  100000     1.73969    1.73969     28    28.0000
363  100000     1.73969    1.73969     28    28.0000
364  100000     1.73969    1.73969     28    28.0000
365  100000     1.73969    1.73969     28    28.0000
366  100000     1.73969    1.73969     28    28.0000
367  100000     1.73969    1.73969     28    28.0000
368  100000     1.73969    1.73969     28    28.0000
369  100000     1.73969    1.73969     28    28.0000
370  100000     1.73969    1.73969     28    28.0000
371  100000     1.73969    1.73969     28    28.0000
372  100000     1.73969    1.73969     28    28.0000
373  100000     1.73969    1.73969     28    28.0000
374  100000     1.73969    1.73969     28    28.0000
375  100000     1.73969    1.73969     28    28.0000
376  100000     1.73969    1.73969     28    28.0000
377  100000     1.73969    1.73969     28    28.0000
378  100000     1.73969    1.73969     28    28.0000
379  100000     1.73969    1.73969     28    28.0000
380  100000     1.73969    1.73969     28    28.0000
381  100000     1.73969    1.73969     28    28.0000
382  100000     1.73969    1.73969     28    28.0000
383  100000     1.73969    1.73969     28    28.0000
384  100000     1.73969    1.73969     28    28.0000
385  100000     1.73969    1.73969     28    28.0000
386  100000     1.73969    1.73969     28    28.0000
387  100000     1.73969    1.73969     28    28.0000
388  100000     1.73969    1.73969     28    28.0000
389  100000     1.73969    1.73969     28    28.0000
390  100000     1.73969    1.73969     28    28.0000
391  100000     1.73969    1.73969     28    28.0000
392  100000     1.73969    1.73969     28    28.0000
393  100000     1.73969    1.73969     28    28.0000
394  100000     1.73969    1.73969     28    28.0000
395  100000     1.73969    1.73969     28    28.0000
396  100000     1.73969    1.73969     28    28.0000
397  100000     1.73968    1.73969     28    28.0000
398  100000     1.73969    1.73969     28    28.0000
399  100000     1.73969    1.73969     28    28.0000
400  100000     1.73969    1.73969     28    28.0000

Asexual Model with selection (random number based on
relative fitness to determine who survives).  Mutation rate is      1e-08,
Recomb rate is ONE per asexual individual per generation
Starting values at generation   0 are; population size 100000,
with avg fitness of    1.22091.

gen  popSize    avgFit     maxFit maxAlleles avgAlleles
  0  100000     1.22091    1.64061     25     9.9902
  1  100000     1.22091    1.64061     25     9.9902
  2  100000     1.22604    1.64061     25    10.2003
  3  100000     1.23100    1.64061     25    10.4020
  4  100000     1.23567    1.64061     25    10.5919
  5  100000     1.24038    1.64061     25    10.7829
  6  100000     1.24552    1.64061     25    10.9898
  7  100000     1.25057    1.64061     25    11.1925
  8  100000     1.25583    1.64061     25    11.4029
  9  100000     1.26122    1.64061     25    11.6176
 10  100000     1.26698    1.64061     25    11.8460
 11  100000     1.27247    1.64061     25    12.0621
 12  100000     1.27835    1.64061     25    12.2936
 13  100000     1.28390    1.64061     25    12.5118
 14  100000     1.28941    1.64061     25    12.7261
 15  100000     1.29560    1.64061     25    12.9665
 16  100000     1.30217    1.64061     25    13.2210
 17  100000     1.30861    1.64061     25    13.4682
 18  100000     1.31532    1.64061     25    13.7242
 19  100000     1.32202    1.64061     25    13.9800
 20  100000     1.32886    1.64061     25    14.2400
 21  100000     1.33571    1.64061     25    14.4986
 22  100000     1.34270    1.64061     25    14.7619
 23  100000     1.34962    1.64061     25    15.0206
 24  100000     1.35674    1.64061     25    15.2866
 25  100000     1.36457    1.64061     25    15.5763
 26  100000     1.37187    1.64061     25    15.8462
 27  100000     1.37902    1.64061     25    16.1087
 28  100000     1.38617    1.64061     25    16.3706
 29  100000     1.39355    1.64061     25    16.6389
 30  100000     1.40085    1.64061     25    16.9040
 31  100000     1.40787    1.64061     25    17.1576
 32  100000     1.41488    1.64061     25    17.4101
 33  100000     1.42209    1.64061     25    17.6683
 34  100000     1.42932    1.64061     25    17.9264
 35  100000     1.43635    1.64061     25    18.1761
 36  100000     1.44336    1.64061     25    18.4245
 37  100000     1.44984    1.64061     25    18.6533
 38  100000     1.45659    1.64061     25    18.8904
 39  100000     1.46313    1.64061     25    19.1193
 40  100000     1.46988    1.64061     25    19.3551
 41  100000     1.47663    1.64061     25    19.5896
 42  100000     1.48330    1.64061     25    19.8209
 43  100000     1.48940    1.64061     25    20.0310
 44  100000     1.49544    1.64061     25    20.2386
 45  100000     1.50099    1.64061     25    20.4284
 46  100000     1.50682    1.64061     25    20.6277
 47  100000     1.51254    1.64061     25    20.8221
 48  100000     1.51819    1.64061     25    21.0138
 49  100000     1.52341    1.64061     25    21.1905
 50  100000     1.52834    1.64061     25    21.3574
 51  100000     1.53310    1.64061     25    21.5174
 52  100000     1.53789    1.64061     25    21.6779
 53  100000     1.54231    1.64061     25    21.8260
 54  100000     1.54638    1.64061     25    21.9623
 55  100000     1.55049    1.64061     25    22.0988
 56  100000     1.55482    1.64061     25    22.2433
 57  100000     1.55865    1.64061     25    22.3700
 58  100000     1.56247    1.64061     25    22.4968
 59  100000     1.56564    1.64061     25    22.6009
 60  100000     1.56896    1.64061     25    22.7105
 61  100000     1.57202    1.64061     25    22.8109
 62  100000     1.57489    1.64061     25    22.9047
 63  100000     1.57743    1.64061     25    22.9880
 64  100000     1.58003    1.64061     25    23.0729
 65  100000     1.58240    1.64061     25    23.1505
 66  100000     1.58477    1.64061     25    23.2275
 67  100000     1.58695    1.64061     25    23.2986
 68  100000     1.58920    1.64061     25    23.3716
 69  100000     1.59144    1.64061     25    23.4441
 70  100000     1.59359    1.64061     25    23.5138
 71  100000     1.59558    1.64061     25    23.5780
 72  100000     1.59734    1.64061     25    23.6349
 73  100000     1.59895    1.64061     25    23.6864
 74  100000     1.60052    1.64061     25    23.7373
 75  100000     1.60202    1.64061     25    23.7853
 76  100000     1.60368    1.64061     25    23.8384
 77  100000     1.60515    1.64061     25    23.8853
 78  100000     1.60636    1.64061     25    23.9240
 79  100000     1.60762    1.64061     25    23.9643
 80  100000     1.60890    1.64061     25    24.0050
 81  100000     1.61011    1.64061     25    24.0436
 82  100000     1.61141    1.64061     25    24.0852
 83  100000     1.61271    1.64061     25    24.1269
 84  100000     1.61377    1.64061     25    24.1604
 85  100000     1.61476    1.64061     25    24.1918
 86  100000     1.61576    1.64061     25    24.2233
 87  100000     1.61672    1.64061     25    24.2539
 88  100000     1.61756    1.64061     25    24.2805
 89  100000     1.61856    1.64061     25    24.3122
 90  100000     1.61949    1.64061     25    24.3416
 91  100000     1.62021    1.64061     25    24.3642
 92  100000     1.62102    1.64061     25    24.3897
 93  100000     1.62171    1.64061     25    24.4115
 94  100000     1.62245    1.64061     25    24.4348
 95  100000     1.62311    1.64061     25    24.4555
 96  100000     1.62383    1.64061     25    24.4782
 97  100000     1.62433    1.64061     25    24.4939
 98  100000     1.62485    1.64061     25    24.5104
 99  100000     1.62551    1.64061     25    24.5309
100  100000     1.62598    1.64061     25    24.5456
101  100000     1.62650    1.64061     25    24.5620
102  100000     1.62701    1.64061     25    24.5777
103  100000     1.62737    1.64061     25    24.5892
104  100000     1.62788    1.64061     25    24.6049
105  100000     1.62834    1.64061     25    24.6194
106  100000     1.62888    1.64061     25    24.6365
107  100000     1.62923    1.64061     25    24.6472
108  100000     1.62961    1.64061     25    24.6591
109  100000     1.62994    1.64061     25    24.6695
110  100000     1.63030    1.64061     25    24.6806
111  100000     1.63067    1.64061     25    24.6920
112  100000     1.63100    1.64061     25    24.7023
113  100000     1.63141    1.64061     25    24.7151
114  100000     1.63166    1.64061     25    24.7229
115  100000     1.63199    1.64061     25    24.7333
116  100000     1.63233    1.64061     25    24.7438
117  100000     1.63255    1.64061     25    24.7508
118  100000     1.63290    1.64061     25    24.7614
119  100000     1.63307    1.64061     25    24.7667
120  100000     1.63337    1.64061     25    24.7762
121  100000     1.63356    1.64061     25    24.7822
122  100000     1.63375    1.64061     25    24.7880
123  100000     1.63399    1.64061     25    24.7956
124  100000     1.63414    1.64061     25    24.8002
125  100000     1.63433    1.64061     25    24.8060
126  100000     1.63456    1.64061     25    24.8132
127  100000     1.63475    1.64061     25    24.8190
128  100000     1.63483    1.64061     25    24.8216
129  100000     1.63495    1.64061     25    24.8252
130  100000     1.63520    1.64061     25    24.8330
131  100000     1.63532    1.64061     25    24.8369
132  100000     1.63557    1.64061     25    24.8445
133  100000     1.63571    1.64061     25    24.8489
134  100000     1.63586    1.64061     25    24.8534
135  100000     1.63604    1.64061     25    24.8591
136  100000     1.63615    1.64061     25    24.8626
137  100000     1.63628    1.64061     25    24.8666
138  100000     1.63647    1.64061     25    24.8726
139  100000     1.63660    1.64061     25    24.8763
140  100000     1.63676    1.64061     25    24.8814
141  100000     1.63692    1.64061     25    24.8864
142  100000     1.63703    1.64061     25    24.8898
143  100000     1.63713    1.64061     25    24.8930
144  100000     1.63725    1.64061     25    24.8966
145  100000     1.63735    1.64061     25    24.8998
146  100000     1.63743    1.64061     25    24.9022
147  100000     1.63751    1.64061     25    24.9045
148  100000     1.63758    1.64061     25    24.9068
149  100000     1.63769    1.64061     25    24.9103
150  100000     1.63781    1.64061     25    24.9137
151  100000     1.63791    1.64061     25    24.9170
152  100000     1.63801    1.64061     25    24.9199
153  100000     1.63807    1.64061     25    24.9220
154  100000     1.63817    1.64061     25    24.9251
155  100000     1.63826    1.64061     25    24.9276
156  100000     1.63830    1.64061     25    24.9289
157  100000     1.63837    1.64061     25    24.9311
158  100000     1.63842    1.64061     25    24.9329
159  100000     1.63845    1.64061     25    24.9336
160  100000     1.63848    1.64061     25    24.9346
161  100000     1.63853    1.64061     25    24.9361
162  100000     1.63857    1.64061     25    24.9373
163  100000     1.63863    1.64061     25    24.9393
164  100000     1.63869    1.64061     25    24.9410
165  100000     1.63873    1.64061     25    24.9422
166  100000     1.63877    1.64061     25    24.9437
167  100000     1.63885    1.64061     25    24.9459
168  100000     1.63891    1.64061     25    24.9480
169  100000     1.63895    1.64061     25    24.9492
170  100000     1.63900    1.64061     25    24.9507
171  100000     1.63905    1.64061     25    24.9520
172  100000     1.63909    1.64061     25    24.9534
173  100000     1.63914    1.64061     25    24.9550
174  100000     1.63918    1.64061     25    24.9561
175  100000     1.63921    1.64061     25    24.9570
176  100000     1.63919    1.64061     25    24.9566
177  100000     1.63924    1.64061     25    24.9581
178  100000     1.63927    1.64061     25    24.9590
179  100000     1.63929    1.64061     25    24.9597
180  100000     1.63935    1.64061     25    24.9613
181  100000     1.63941    1.64061     25    24.9632
182  100000     1.63945    1.64061     25    24.9646
183  100000     1.63945    1.64061     25    24.9644
184  100000     1.63947    1.64061     25    24.9649
185  100000     1.63949    1.64061     25    24.9658
186  100000     1.63953    1.64061     25    24.9670
187  100000     1.63956    1.64061     25    24.9680
188  100000     1.63957    1.64061     25    24.9682
189  100000     1.63960    1.64061     25    24.9691
190  100000     1.63964    1.64061     25    24.9704
191  100000     1.63966    1.64061     25    24.9709
192  100000     1.63967    1.64061     25    24.9713
193  100000     1.63969    1.64061     25    24.9719
194  100000     1.63974    1.64061     25    24.9735
195  100000     1.63978    1.64061     25    24.9747
196  100000     1.63982    1.64061     25    24.9758
197  100000     1.63982    1.64061     25    24.9759
198  100000     1.63985    1.64061     25    24.9767
199  100000     1.63987    1.64061     25    24.9774
200  100000     1.63990    1.64061     25    24.9783
201  100000     1.63989    1.64061     25    24.9780
202  100000     1.63990    1.64061     25    24.9783
203  100000     1.63991    1.64061     25    24.9787
204  100000     1.63995    1.64061     25    24.9799
205  100000     1.63998    1.64061     25    24.9808
206  100000     1.64001    1.64061     25    24.9816
207  100000     1.64003    1.64061     25    24.9822
208  100000     1.64003    1.64061     25    24.9825
209  100000     1.64005    1.64061     25    24.9828
210  100000     1.64005    1.64061     25    24.9830
211  100000     1.64006    1.64061     25    24.9833
212  100000     1.64007    1.64061     25    24.9836
213  100000     1.64008    1.64061     25    24.9839
214  100000     1.64010    1.64061     25    24.9846
215  100000     1.64012    1.64061     25    24.9852
216  100000     1.64012    1.64061     25    24.9851
217  100000     1.64015    1.64061     25    24.9859
218  100000     1.64016    1.64061     25    24.9863
219  100000     1.64017    1.64061     25    24.9866
220  100000     1.64018    1.64061     25    24.9871
221  100000     1.64019    1.64061     25    24.9873
222  100000     1.64021    1.64061     25    24.9879
223  100000     1.64023    1.64061     25    24.9884
224  100000     1.64024    1.64061     25    24.9886
225  100000     1.64024    1.64061     25    24.9888
226  100000     1.64025    1.64061     25    24.9891
227  100000     1.64026    1.64061     25    24.9893
228  100000     1.64026    1.64061     25    24.9896
229  100000     1.64026    1.64061     25    24.9894
230  100000     1.64027    1.64061     25    24.9898
231  100000     1.64028    1.64061     25    24.9901
232  100000     1.64029    1.64061     25    24.9903
233  100000     1.64029    1.64061     25    24.9902
234  100000     1.64030    1.64061     25    24.9906
235  100000     1.64030    1.64061     25    24.9908
236  100000     1.64032    1.64061     25    24.9913
237  100000     1.64032    1.64061     25    24.9913
238  100000     1.64032    1.64061     25    24.9912
239  100000     1.64032    1.64061     25    24.9913
240  100000     1.64033    1.64061     25    24.9915
241  100000     1.64033    1.64061     25    24.9916
242  100000     1.64034    1.64061     25    24.9920
243  100000     1.64035    1.64061     25    24.9922
244  100000     1.64035    1.64061     25    24.9921
245  100000     1.64034    1.64061     25    24.9920
246  100000     1.64035    1.64061     25    24.9921
247  100000     1.64035    1.64061     25    24.9922
248  100000     1.64036    1.64061     25    24.9925
249  100000     1.64037    1.64061     25    24.9928
250  100000     1.64037    1.64061     25    24.9927
251  100000     1.64036    1.64061     25    24.9926
252  100000     1.64038    1.64061     25    24.9932
253  100000     1.64040    1.64061     25    24.9937
254  100000     1.64041    1.64061     25    24.9940
255  100000     1.64041    1.64061     25    24.9941
256  100000     1.64042    1.64061     25    24.9944
257  100000     1.64043    1.64061     25    24.9945
258  100000     1.64043    1.64061     25    24.9945
259  100000     1.64043    1.64061     25    24.9946
260  100000     1.64043    1.64061     25    24.9945
261  100000     1.64042    1.64061     25    24.9942
262  100000     1.64044    1.64061     25    24.9949
263  100000     1.64044    1.64061     25    24.9949
264  100000     1.64044    1.64061     25    24.9949
265  100000     1.64045    1.64061     25    24.9953
266  100000     1.64045    1.64061     25    24.9954
267  100000     1.64046    1.64061     25    24.9955
268  100000     1.64046    1.64061     25    24.9955
269  100000     1.64045    1.64061     25    24.9954
270  100000     1.64046    1.64061     25    24.9956
271  100000     1.64047    1.64061     25    24.9958
272  100000     1.64047    1.64061     25    24.9958
273  100000     1.64047    1.64061     25    24.9958
274  100000     1.64047    1.64061     25    24.9959
275  100000     1.64047    1.64061     25    24.9959
276  100000     1.64048    1.64061     25    24.9961
277  100000     1.64048    1.64061     25    24.9961
278  100000     1.64048    1.64061     25    24.9963
279  100000     1.64048    1.64061     25    24.9963
280  100000     1.64049    1.64061     25    24.9965
281  100000     1.64049    1.64061     25    24.9963
282  100000     1.64049    1.64061     25    24.9966
283  100000     1.64050    1.64061     25    24.9969
284  100000     1.64050    1.64061     25    24.9968
285  100000     1.64051    1.64061     25    24.9970
286  100000     1.64051    1.64061     25    24.9970
287  100000     1.64050    1.64061     25    24.9967
288  100000     1.64051    1.64061     25    24.9969
289  100000     1.64051    1.64061     25    24.9971
290  100000     1.64051    1.64061     25    24.9970
291  100000     1.64051    1.64061     25    24.9972
292  100000     1.64052    1.64061     25    24.9974
293  100000     1.64053    1.64061     25    24.9975
294  100000     1.64053    1.64061     25    24.9976
295  100000     1.64053    1.64061     25    24.9977
296  100000     1.64053    1.64061     25    24.9978
297  100000     1.64054    1.64061     25    24.9979
298  100000     1.64054    1.64061     25    24.9979
299  100000     1.64054    1.64061     25    24.9980
300  100000     1.64055    1.64061     25    24.9982
301  100000     1.64055    1.64061     25    24.9982
302  100000     1.64055    1.64061     25    24.9984
303  100000     1.64055    1.64061     25    24.9984
304  100000     1.64056    1.64061     25    24.9985
305  100000     1.64056    1.64061     25    24.9985
306  100000     1.64056    1.64061     25    24.9985
307  100000     1.64056    1.64061     25    24.9985
308  100000     1.64055    1.64061     25    24.9984
309  100000     1.64056    1.64061     25    24.9985
310  100000     1.64056    1.64061     25    24.9986
311  100000     1.64056    1.64061     25    24.9986
312  100000     1.64056    1.64061     25    24.9987
313  100000     1.64056    1.64061     25    24.9986
314  100000     1.64056    1.64061     25    24.9987
315  100000     1.64056    1.64061     25    24.9986
316  100000     1.64057    1.64061     25    24.9988
317  100000     1.64057    1.64061     25    24.9988
318  100000     1.64057    1.64061     25    24.9990
319  100000     1.64057    1.64061     25    24.9990
320  100000     1.64057    1.64061     25    24.9990
321  100000     1.64057    1.64061     25    24.9990
322  100000     1.64058    1.64061     25    24.9991
323  100000     1.64057    1.64061     25    24.9989
324  100000     1.64057    1.64061     25    24.9989
325  100000     1.64057    1.64061     25    24.9988
326  100000     1.64057    1.64061     25    24.9989
327  100000     1.64057    1.64061     25    24.9990
328  100000     1.64057    1.64061     25    24.9990
329  100000     1.64058    1.64061     25    24.9991
330  100000     1.64058    1.64061     25    24.9991
331  100000     1.64058    1.64061     25    24.9991
332  100000     1.64058    1.64061     25    24.9991
333  100000     1.64058    1.64061     25    24.9991
334  100000     1.64058    1.64061     25    24.9992
335  100000     1.64058    1.64061     25    24.9991
336  100000     1.64058    1.64061     25    24.9992
337  100000     1.64058    1.64061     25    24.9992
338  100000     1.64059    1.64061     25    24.9994
339  100000     1.64059    1.64061     25    24.9994
340  100000     1.64059    1.64061     25    24.9995
341  100000     1.64059    1.64061     25    24.9995
342  100000     1.64059    1.64061     25    24.9995
343  100000     1.64059    1.64061     25    24.9995
344  100000     1.64059    1.64061     25    24.9994
345  100000     1.64059    1.64061     25    24.9994
346  100000     1.64059    1.64061     25    24.9994
347  100000     1.64059    1.64061     25    24.9994
348  100000     1.64059    1.64061     25    24.9995
349  100000     1.64059    1.64061     25    24.9995
350  100000     1.64059    1.64061     25    24.9996
351  100000     1.64059    1.64061     25    24.9996
352  100000     1.64060    1.64061     25    24.9997
353  100000     1.64060    1.64061     25    24.9997
354  100000     1.64060    1.64061     25    24.9997
355  100000     1.64060    1.64061     25    24.9997
356  100000     1.64060    1.64061     25    24.9998
357  100000     1.64060    1.64061     25    24.9998
358  100000     1.64060    1.64061     25    24.9998
359  100000     1.64060    1.64061     25    24.9998
360  100000     1.64060    1.64061     25    24.9998
361  100000     1.64060    1.64061     25    24.9998
362  100000     1.64060    1.64061     25    24.9999
363  100000     1.64060    1.64061     25    24.9999
364  100000     1.64060    1.64061     25    24.9999
365  100000     1.64060    1.64061     25    24.9999
366  100000     1.64060    1.64061     25    24.9999
367  100000     1.64060    1.64061     25    24.9999
368  100000     1.64060    1.64061     25    24.9999
369  100000     1.64060    1.64061     25    24.9999
370  100000     1.64060    1.64061     25    24.9998
371  100000     1.64060    1.64061     25    24.9998
372  100000     1.64060    1.64061     25    24.9998
373  100000     1.64060    1.64061     25    24.9999
374  100000     1.64060    1.64061     25    24.9998
375  100000     1.64060    1.64061     25    24.9998
376  100000     1.64060    1.64061     25    24.9998
377  100000     1.64060    1.64061     25    24.9998
378  100000     1.64060    1.64061     25    24.9999
379  100000     1.64060    1.64061     25    24.9999
380  100000     1.64060    1.64061     25    24.9999
381  100000     1.64060    1.64061     25    25.0000
382  100000     1.64061    1.64061     25    25.0000
383  100000     1.64061    1.64061     25    25.0000
384  100000     1.64061    1.64061     25    25.0000
385  100000     1.64061    1.64061     25    25.0000
386  100000     1.64060    1.64061     25    25.0000
387  100000     1.64061    1.64061     25    25.0000
388  100000     1.64061    1.64061     25    25.0000
389  100000     1.64061    1.64061     25    25.0000
390  100000     1.64061    1.64061     25    25.0000
391  100000     1.64061    1.64061     25    25.0000
392  100000     1.64061    1.64061     25    25.0000
393  100000     1.64061    1.64061     25    25.0000
394  100000     1.64061    1.64061     25    25.0000
395  100000     1.64061    1.64061     25    25.0000
396  100000     1.64061    1.64061     25    25.0000
397  100000     1.64061    1.64061     25    25.0000
398  100000     1.64061    1.64061     25    25.0000
399  100000     1.64061    1.64061     25    25.0000
400  100000     1.64060    1.64061     25    24.9999

Asexual Model with selection (random number based on
relative fitness to determine who survives).  Mutation rate is      1e-08,
Recomb rate is ONE per asexual individual per generation
Starting values at generation   0 are; population size 100000,
with avg fitness of    1.22117.

gen  popSize    avgFit     maxFit maxAlleles avgAlleles
  0  100000     1.22117    1.70623     27    10.0003
  1  100000     1.22117    1.70623     27    10.0003
  2  100000     1.22592    1.70623     27    10.1954
  3  100000     1.23027    1.70623     27    10.3731
  4  100000     1.23486    1.70623     27    10.5595
  5  100000     1.23971    1.70623     27    10.7560
  6  100000     1.24459    1.70623     27    10.9530
  7  100000     1.24970    1.70623     27    11.1580
  8  100000     1.25505    1.70623     27    11.3715
  9  100000     1.26049    1.70623     27    11.5879
 10  100000     1.26576    1.70623     27    11.7963
 11  100000     1.27135    1.70623     27    12.0162
 12  100000     1.27691    1.70623     27    12.2336
 13  100000     1.28339    1.70623     27    12.4863
 14  100000     1.29011    1.70623     27    12.7478
 15  100000     1.29640    1.70623     27    12.9903
 16  100000     1.30355    1.70623     27    13.2660
 17  100000     1.31027    1.70623     27    13.5228
 18  100000     1.31743    1.70623     27    13.7961
 19  100000     1.32494    1.70623     27    14.0796
 20  100000     1.33207    1.70623     27    14.3472
 21  100000     1.33955    1.70623     27    14.6259
 22  100000     1.34742    1.70623     27    14.9182
 23  100000     1.35616    1.70623     27    15.2417
 24  100000     1.36460    1.70623     27    15.5506
 25  100000     1.37348    1.70623     27    15.8744
 26  100000     1.38234    1.70623     27    16.1953
 27  100000     1.39126    1.70623     27    16.5157
 28  100000     1.40061    1.70623     27    16.8508
 29  100000     1.41028    1.70623     27    17.1963
 30  100000     1.42010    1.70623     27    17.5439
 31  100000     1.43072    1.70623     27    17.9187
 32  100000     1.44092    1.70623     27    18.2752
 33  100000     1.45168    1.70623     27    18.6502
 34  100000     1.46257    1.70623     27    19.0292
 35  100000     1.47338    1.70623     27    19.4018
 36  100000     1.48384    1.70623     27    19.7606
 37  100000     1.49492    1.70623     27    20.1384
 38  100000     1.50642    1.70623     27    20.5298
 39  100000     1.51651    1.70623     27    20.8706
 40  100000     1.52680    1.70623     27    21.2181
 41  100000     1.53679    1.70623     27    21.5535
 42  100000     1.54638    1.70623     27    21.8748
 43  100000     1.55540    1.70623     27    22.1761
 44  100000     1.56457    1.70623     27    22.4798
 45  100000     1.57302    1.70623     27    22.7589
 46  100000     1.58074    1.70623     27    23.0135
 47  100000     1.58808    1.70623     27    23.2553
 48  100000     1.59516    1.70623     27    23.4875
 49  100000     1.60226    1.70623     27    23.7193
 50  100000     1.60866    1.70623     27    23.9284
 51  100000     1.61465    1.70623     27    24.1232
 52  100000     1.62021    1.70623     27    24.3029
 53  100000     1.62519    1.70623     27    24.4644
 54  100000     1.63025    1.70623     27    24.6279
 55  100000     1.63435    1.70623     27    24.7598
 56  100000     1.63859    1.70623     27    24.8960
 57  100000     1.64264    1.70623     27    25.0258
 58  100000     1.64599    1.70623     27    25.1326
 59  100000     1.64896    1.70623     27    25.2271
 60  100000     1.65195    1.70623     27    25.3225
 61  100000     1.65444    1.70623     27    25.4013
 62  100000     1.65692    1.70623     27    25.4800
 63  100000     1.65939    1.70623     27    25.5586
 64  100000     1.66170    1.70623     27    25.6317
 65  100000     1.66365    1.70623     27    25.6931
 66  100000     1.66563    1.70623     27    25.7554
 67  100000     1.66736    1.70623     27    25.8098
 68  100000     1.66885    1.70623     27    25.8564
 69  100000     1.67042    1.70623     27    25.9053
 70  100000     1.67169    1.70623     27    25.9453
 71  100000     1.67316    1.70623     27    25.9912
 72  100000     1.67428    1.70623     27    26.0259
 73  100000     1.67542    1.70623     27    26.0617
 74  100000     1.67641    1.70623     27    26.0924
 75  100000     1.67760    1.70623     27    26.1294
 76  100000     1.67860    1.70623     27    26.1603
 77  100000     1.67951    1.70623     27    26.1887
 78  100000     1.68045    1.70623     27    26.2175
 79  100000     1.68117    1.70623     27    26.2397
 80  100000     1.68204    1.70623     27    26.2666
 81  100000     1.68276    1.70623     27    26.2889
 82  100000     1.68365    1.70623     27    26.3164
 83  100000     1.68430    1.70623     27    26.3363
 84  100000     1.68500    1.70623     27    26.3579
 85  100000     1.68554    1.70623     27    26.3744
 86  100000     1.68604    1.70623     27    26.3898
 87  100000     1.68653    1.70623     27    26.4049
 88  100000     1.68719    1.70623     27    26.4250
 89  100000     1.68770    1.70623     27    26.4404
 90  100000     1.68814    1.70623     27    26.4541
 91  100000     1.68863    1.70623     27    26.4690
 92  100000     1.68905    1.70623     27    26.4817
 93  100000     1.68957    1.70623     27    26.4977
 94  100000     1.68999    1.70623     27    26.5103
 95  100000     1.69039    1.70623     27    26.5225
 96  100000     1.69077    1.70623     27    26.5341
 97  100000     1.69108    1.70623     27    26.5435
 98  100000     1.69134    1.70623     27    26.5515
 99  100000     1.69167    1.70623     27    26.5616
100  100000     1.69205    1.70623     27    26.5730
101  100000     1.69238    1.70623     27    26.5830
102  100000     1.69267    1.70623     27    26.5920
103  100000     1.69300    1.70623     27    26.6018
104  100000     1.69326    1.70623     27    26.6097
105  100000     1.69352    1.70623     27    26.6179
106  100000     1.69379    1.70623     27    26.6258
107  100000     1.69409    1.70623     27    26.6348
108  100000     1.69433    1.70623     27    26.6424
109  100000     1.69465    1.70623     27    26.6520
110  100000     1.69495    1.70623     27    26.6611
111  100000     1.69526    1.70623     27    26.6705
112  100000     1.69553    1.70623     27    26.6787
113  100000     1.69571    1.70623     27    26.6841
114  100000     1.69603    1.70623     27    26.6938
115  100000     1.69626    1.70623     27    26.7005
116  100000     1.69646    1.70623     27    26.7066
117  100000     1.69663    1.70623     27    26.7118
118  100000     1.69682    1.70623     27    26.7176
119  100000     1.69697    1.70623     27    26.7219
120  100000     1.69714    1.70623     27    26.7270
121  100000     1.69734    1.70623     27    26.7333
122  100000     1.69745    1.70623     27    26.7366
123  100000     1.69773    1.70623     27    26.7449
124  100000     1.69790    1.70623     27    26.7500
125  100000     1.69798    1.70623     27    26.7524
126  100000     1.69816    1.70623     27    26.7579
127  100000     1.69832    1.70623     27    26.7628
128  100000     1.69856    1.70623     27    26.7699
129  100000     1.69867    1.70623     27    26.7731
130  100000     1.69878    1.70623     27    26.7766
131  100000     1.69895    1.70623     27    26.7818
132  100000     1.69913    1.70623     27    26.7871
133  100000     1.69924    1.70623     27    26.7903
134  100000     1.69944    1.70623     27    26.7963
135  100000     1.69961    1.70623     27    26.8014
136  100000     1.69974    1.70623     27    26.8054
137  100000     1.69987    1.70623     27    26.8095
138  100000     1.70004    1.70623     27    26.8143
139  100000     1.70017    1.70623     27    26.8183
140  100000     1.70026    1.70623     27    26.8210
141  100000     1.70031    1.70623     27    26.8227
142  100000     1.70042    1.70623     27    26.8258
143  100000     1.70051    1.70623     27    26.8287
144  100000     1.70059    1.70623     27    26.8309
145  100000     1.70069    1.70623     27    26.8339
146  100000     1.70080    1.70623     27    26.8373
147  100000     1.70093    1.70623     27    26.8412
148  100000     1.70108    1.70623     27    26.8456
149  100000     1.70114    1.70623     27    26.8474
150  100000     1.70128    1.70623     27    26.8518
151  100000     1.70137    1.70623     27    26.8545
152  100000     1.70139    1.70623     27    26.8551
153  100000     1.70146    1.70623     27    26.8572
154  100000     1.70151    1.70623     27    26.8586
155  100000     1.70160    1.70623     27    26.8614
156  100000     1.70169    1.70623     27    26.8639
157  100000     1.70179    1.70623     27    26.8670
158  100000     1.70189    1.70623     27    26.8701
159  100000     1.70196    1.70623     27    26.8720
160  100000     1.70203    1.70623     27    26.8743
161  100000     1.70209    1.70623     27    26.8760
162  100000     1.70214    1.70623     27    26.8776
163  100000     1.70224    1.70623     27    26.8805
164  100000     1.70232    1.70623     27    26.8828
165  100000     1.70238    1.70623     27    26.8847
166  100000     1.70247    1.70623     27    26.8873
167  100000     1.70253    1.70623     27    26.8892
168  100000     1.70262    1.70623     27    26.8919
169  100000     1.70271    1.70623     27    26.8946
170  100000     1.70275    1.70623     27    26.8960
171  100000     1.70279    1.70623     27    26.8972
172  100000     1.70288    1.70623     27    26.8997
173  100000     1.70294    1.70623     27    26.9014
174  100000     1.70303    1.70623     27    26.9041
175  100000     1.70309    1.70623     27    26.9061
176  100000     1.70315    1.70623     27    26.9079
177  100000     1.70318    1.70623     27    26.9088
178  100000     1.70327    1.70623     27    26.9113
179  100000     1.70333    1.70623     27    26.9134
180  100000     1.70340    1.70623     27    26.9154
181  100000     1.70341    1.70623     27    26.9157
182  100000     1.70346    1.70623     27    26.9170
183  100000     1.70354    1.70623     27    26.9196
184  100000     1.70357    1.70623     27    26.9205
185  100000     1.70367    1.70623     27    26.9234
186  100000     1.70372    1.70623     27    26.9250
187  100000     1.70379    1.70623     27    26.9269
188  100000     1.70384    1.70623     27    26.9284
189  100000     1.70391    1.70623     27    26.9306
190  100000     1.70396    1.70623     27    26.9321
191  100000     1.70398    1.70623     27    26.9328
192  100000     1.70400    1.70623     27    26.9334
193  100000     1.70405    1.70623     27    26.9347
194  100000     1.70406    1.70623     27    26.9349
195  100000     1.70411    1.70623     27    26.9365
196  100000     1.70417    1.70623     27    26.9384
197  100000     1.70421    1.70623     27    26.9395
198  100000     1.70423    1.70623     27    26.9400
199  100000     1.70428    1.70623     27    26.9418
200  100000     1.70434    1.70623     27    26.9434
201  100000     1.70436    1.70623     27    26.9442
202  100000     1.70440    1.70623     27    26.9454
203  100000     1.70442    1.70623     27    26.9458
204  100000     1.70444    1.70623     27    26.9464
205  100000     1.70447    1.70623     27    26.9475
206  100000     1.70452    1.70623     27    26.9488
207  100000     1.70454    1.70623     27    26.9494
208  100000     1.70461    1.70623     27    26.9517
209  100000     1.70463    1.70623     27    26.9521
210  100000     1.70465    1.70623     27    26.9529
211  100000     1.70468    1.70623     27    26.9536
212  100000     1.70471    1.70623     27    26.9546
213  100000     1.70474    1.70623     27    26.9554
214  100000     1.70476    1.70623     27    26.9560
215  100000     1.70479    1.70623     27    26.9569
216  100000     1.70480    1.70623     27    26.9572
217  100000     1.70483    1.70623     27    26.9581
218  100000     1.70484    1.70623     27    26.9585
219  100000     1.70489    1.70623     27    26.9598
220  100000     1.70490    1.70623     27    26.9602
221  100000     1.70493    1.70623     27    26.9610
222  100000     1.70494    1.70623     27    26.9613
223  100000     1.70496    1.70623     27    26.9619
224  100000     1.70499    1.70623     27    26.9628
225  100000     1.70502    1.70623     27    26.9637
226  100000     1.70506    1.70623     27    26.9650
227  100000     1.70508    1.70623     27    26.9658
228  100000     1.70511    1.70623     27    26.9665
229  100000     1.70513    1.70623     27    26.9672
230  100000     1.70514    1.70623     27    26.9673
231  100000     1.70516    1.70623     27    26.9679
232  100000     1.70519    1.70623     27    26.9688
233  100000     1.70520    1.70623     27    26.9692
234  100000     1.70523    1.70623     27    26.9702
235  100000     1.70525    1.70623     27    26.9708
236  100000     1.70527    1.70623     27    26.9713
237  100000     1.70530    1.70623     27    26.9722
238  100000     1.70532    1.70623     27    26.9728
239  100000     1.70534    1.70623     27    26.9733
240  100000     1.70536    1.70623     27    26.9739
241  100000     1.70538    1.70623     27    26.9745
242  100000     1.70538    1.70623     27    26.9747
243  100000     1.70539    1.70623     27    26.9749
244  100000     1.70541    1.70623     27    26.9756
245  100000     1.70541    1.70623     27    26.9756
246  100000     1.70543    1.70623     27    26.9759
247  100000     1.70543    1.70623     27    26.9762
248  100000     1.70546    1.70623     27    26.9769
249  100000     1.70549    1.70623     27    26.9778
250  100000     1.70550    1.70623     27    26.9782
251  100000     1.70551    1.70623     27    26.9786
252  100000     1.70554    1.70623     27    26.9794
253  100000     1.70555    1.70623     27    26.9796
254  100000     1.70557    1.70623     27    26.9802
255  100000     1.70559    1.70623     27    26.9807
256  100000     1.70561    1.70623     27    26.9815
257  100000     1.70562    1.70623     27    26.9819
258  100000     1.70564    1.70623     27    26.9823
259  100000     1.70565    1.70623     27    26.9825
260  100000     1.70566    1.70623     27    26.9831
261  100000     1.70568    1.70623     27    26.9837
262  100000     1.70571    1.70623     27    26.9845
263  100000     1.70571    1.70623     27    26.9844
264  100000     1.70573    1.70623     27    26.9850
265  100000     1.70574    1.70623     27    26.9853
266  100000     1.70576    1.70623     27    26.9858
267  100000     1.70576    1.70623     27    26.9861
268  100000     1.70577    1.70623     27    26.9864
269  100000     1.70579    1.70623     27    26.9868
270  100000     1.70581    1.70623     27    26.9874
271  100000     1.70582    1.70623     27    26.9877
272  100000     1.70582    1.70623     27    26.9879
273  100000     1.70584    1.70623     27    26.9882
274  100000     1.70584    1.70623     27    26.9882
275  100000     1.70584    1.70623     27    26.9885
276  100000     1.70586    1.70623     27    26.9889
277  100000     1.70586    1.70623     27    26.9890
278  100000     1.70587    1.70623     27    26.9891
279  100000     1.70587    1.70623     27    26.9891
280  100000     1.70588    1.70623     27    26.9896
281  100000     1.70589    1.70623     27    26.9898
282  100000     1.70589    1.70623     27    26.9900
283  100000     1.70589    1.70623     27    26.9897
284  100000     1.70589    1.70623     27    26.9898
285  100000     1.70590    1.70623     27    26.9902
286  100000     1.70591    1.70623     27    26.9905
287  100000     1.70592    1.70623     27    26.9907
288  100000     1.70594    1.70623     27    26.9913
289  100000     1.70596    1.70623     27    26.9918
290  100000     1.70597    1.70623     27    26.9922
291  100000     1.70596    1.70623     27    26.9918
292  100000     1.70596    1.70623     27    26.9919
293  100000     1.70595    1.70623     27    26.9917
294  100000     1.70596    1.70623     27    26.9918
295  100000     1.70595    1.70623     27    26.9917
296  100000     1.70596    1.70623     27    26.9918
297  100000     1.70596    1.70623     27    26.9920
298  100000     1.70597    1.70623     27    26.9923
299  100000     1.70597    1.70623     27    26.9922
300  100000     1.70598    1.70623     27    26.9924
301  100000     1.70598    1.70623     27    26.9926
302  100000     1.70599    1.70623     27    26.9929
303  100000     1.70600    1.70623     27    26.9931
304  100000     1.70600    1.70623     27    26.9932
305  100000     1.70601    1.70623     27    26.9934
306  100000     1.70602    1.70623     27    26.9938
307  100000     1.70603    1.70623     27    26.9941
308  100000     1.70604    1.70623     27    26.9945
309  100000     1.70605    1.70623     27    26.9946
310  100000     1.70605    1.70623     27    26.9946
311  100000     1.70605    1.70623     27    26.9947
312  100000     1.70606    1.70623     27    26.9950
313  100000     1.70606    1.70623     27    26.9950
314  100000     1.70607    1.70623     27    26.9952
315  100000     1.70607    1.70623     27    26.9952
316  100000     1.70607    1.70623     27    26.9951
317  100000     1.70608    1.70623     27    26.9954
318  100000     1.70609    1.70623     27    26.9957
319  100000     1.70609    1.70623     27    26.9958
320  100000     1.70609    1.70623     27    26.9958
321  100000     1.70609    1.70623     27    26.9958
322  100000     1.70610    1.70623     27    26.9960
323  100000     1.70610    1.70623     27    26.9961
324  100000     1.70610    1.70623     27    26.9961
325  100000     1.70611    1.70623     27    26.9964
326  100000     1.70611    1.70623     27    26.9963
327  100000     1.70611    1.70623     27    26.9965
328  100000     1.70611    1.70623     27    26.9964
329  100000     1.70611    1.70623     27    26.9965
330  100000     1.70612    1.70623     27    26.9966
331  100000     1.70612    1.70623     27    26.9968
332  100000     1.70612    1.70623     27    26.9968
333  100000     1.70613    1.70623     27    26.9971
334  100000     1.70613    1.70623     27    26.9971
335  100000     1.70614    1.70623     27    26.9974
336  100000     1.70615    1.70623     27    26.9975
337  100000     1.70615    1.70623     27    26.9977
338  100000     1.70616    1.70623     27    26.9979
339  100000     1.70616    1.70623     27    26.9978
340  100000     1.70616    1.70623     27    26.9980
341  100000     1.70616    1.70623     27    26.9980
342  100000     1.70617    1.70623     27    26.9981
343  100000     1.70616    1.70623     27    26.9980
344  100000     1.70616    1.70623     27    26.9980
345  100000     1.70616    1.70623     27    26.9980
346  100000     1.70616    1.70623     27    26.9980
347  100000     1.70617    1.70623     27    26.9982
348  100000     1.70617    1.70623     27    26.9981
349  100000     1.70617    1.70623     27    26.9983
350  100000     1.70617    1.70623     27    26.9983
351  100000     1.70617    1.70623     27    26.9982
352  100000     1.70618    1.70623     27    26.9984
353  100000     1.70618    1.70623     27    26.9984
354  100000     1.70618    1.70623     27    26.9984
355  100000     1.70618    1.70623     27    26.9984
356  100000     1.70618    1.70623     27    26.9986
357  100000     1.70618    1.70623     27    26.9986
358  100000     1.70618    1.70623     27    26.9986
359  100000     1.70618    1.70623     27    26.9984
360  100000     1.70618    1.70623     27    26.9984
361  100000     1.70618    1.70623     27    26.9984
362  100000     1.70618    1.70623     27    26.9984
363  100000     1.70618    1.70623     27    26.9985
364  100000     1.70618    1.70623     27    26.9986
365  100000     1.70619    1.70623     27    26.9987
366  100000     1.70619    1.70623     27    26.9988
367  100000     1.70619    1.70623     27    26.9988
368  100000     1.70619    1.70623     27    26.9989
369  100000     1.70619    1.70623     27    26.9989
370  100000     1.70620    1.70623     27    26.9990
371  100000     1.70620    1.70623     27    26.9990
372  100000     1.70619    1.70623     27    26.9989
373  100000     1.70620    1.70623     27    26.9991
374  100000     1.70620    1.70623     27    26.9992
375  100000     1.70620    1.70623     27    26.9992
376  100000     1.70620    1.70623     27    26.9992
377  100000     1.70620    1.70623     27    26.9992
378  100000     1.70620    1.70623     27    26.9991
379  100000     1.70620    1.70623     27    26.9992
380  100000     1.70621    1.70623     27    26.9993
381  100000     1.70620    1.70623     27    26.9992
382  100000     1.70621    1.70623     27    26.9993
383  100000     1.70621    1.70623     27    26.9993
384  100000     1.70621    1.70623     27    26.9994
385  100000     1.70621    1.70623     27    26.9995
386  100000     1.70622    1.70623     27    26.9996
387  100000     1.70622    1.70623     27    26.9997
388  100000     1.70622    1.70623     27    26.9996
389  100000     1.70622    1.70623     27    26.9997
390  100000     1.70622    1.70623     27    26.9997
391  100000     1.70622    1.70623     27    26.9997
392  100000     1.70622    1.70623     27    26.9997
393  100000     1.70622    1.70623     27    26.9998
394  100000     1.70622    1.70623     27    26.9997
395  100000     1.70622    1.70623     27    26.9997
396  100000     1.70622    1.70623     27    26.9998
397  100000     1.70622    1.70623     27    26.9998
398  100000     1.70622    1.70623     27    26.9998
399  100000     1.70622    1.70623     27    26.9998
400  100000     1.70623    1.70623     27    26.9998

Asexual Model with selection (random number based on
relative fitness to determine who survives).  Mutation rate is      1e-08,
Recomb rate is ONE per asexual individual per generation
Starting values at generation   0 are; population size 100000,
with avg fitness of    1.22083.

gen  popSize    avgFit     maxFit maxAlleles avgAlleles
  0  100000     1.22083    1.67277     26     9.9863
  1  100000     1.22083    1.67277     26     9.9863
  2  100000     1.22544    1.67277     26    10.1750
  3  100000     1.23029    1.67277     26    10.3726
  4  100000     1.23514    1.67277     26    10.5694
  5  100000     1.24023    1.67277     26    10.7759
  6  100000     1.24507    1.67277     26    10.9706
  7  100000     1.24991    1.67277     26    11.1655
  8  100000     1.25504    1.67277     26    11.3702
  9  100000     1.26022    1.67277     26    11.5767
 10  100000     1.26570    1.67277     26    11.7947
 11  100000     1.27095    1.67277     26    12.0023
 12  100000     1.27720    1.67277     26    12.2480
 13  100000     1.28293    1.67277     26    12.4718
 14  100000     1.28895    1.67277     26    12.7064
 15  100000     1.29509    1.67277     26    12.9454
 16  100000     1.30118    1.67277     26    13.1804
 17  100000     1.30743    1.67277     26    13.4211
 18  100000     1.31435    1.67277     26    13.6868
 19  100000     1.32065    1.67277     26    13.9265
 20  100000     1.32754    1.67277     26    14.1886
 21  100000     1.33401    1.67277     26    14.4342
 22  100000     1.34091    1.67277     26    14.6944
 23  100000     1.34756    1.67277     26    14.9437
 24  100000     1.35471    1.67277     26    15.2098
 25  100000     1.36156    1.67277     26    15.4649
 26  100000     1.36889    1.67277     26    15.7349
 27  100000     1.37629    1.67277     26    16.0071
 28  100000     1.38370    1.67277     26    16.2782
 29  100000     1.39176    1.67277     26    16.5720
 30  100000     1.39923    1.67277     26    16.8430
 31  100000     1.40687    1.67277     26    17.1177
 32  100000     1.41466    1.67277     26    17.3969
 33  100000     1.42243    1.67277     26    17.6746
 34  100000     1.43027    1.67277     26    17.9530
 35  100000     1.43816    1.67277     26    18.2318
 36  100000     1.44626    1.67277     26    18.5160
 37  100000     1.45438    1.67277     26    18.7992
 38  100000     1.46208    1.67277     26    19.0664
 39  100000     1.46965    1.67277     26    19.3283
 40  100000     1.47742    1.67277     26    19.5954
 41  100000     1.48522    1.67277     26    19.8630
 42  100000     1.49301    1.67277     26    20.1293
 43  100000     1.50077    1.67277     26    20.3931
 44  100000     1.50879    1.67277     26    20.6652
 45  100000     1.51656    1.67277     26    20.9278
 46  100000     1.52420    1.67277     26    21.1849
 47  100000     1.53178    1.67277     26    21.4395
 48  100000     1.53903    1.67277     26    21.6819
 49  100000     1.54619    1.67277     26    21.9207
 50  100000     1.55332    1.67277     26    22.1581
 51  100000     1.56035    1.67277     26    22.3915
 52  100000     1.56710    1.67277     26    22.6152
 53  100000     1.57320    1.67277     26    22.8167
 54  100000     1.57922    1.67277     26    23.0152
 55  100000     1.58506    1.67277     26    23.2075
 56  100000     1.59037    1.67277     26    23.3815
 57  100000     1.59562    1.67277     26    23.5537
 58  100000     1.60056    1.67277     26    23.7150
 59  100000     1.60499    1.67277     26    23.8600
 60  100000     1.60944    1.67277     26    24.0049
 61  100000     1.61328    1.67277     26    24.1297
 62  100000     1.61699    1.67277     26    24.2499
 63  100000     1.62058    1.67277     26    24.3664
 64  100000     1.62387    1.67277     26    24.4731
 65  100000     1.62678    1.67277     26    24.5670
 66  100000     1.62982    1.67277     26    24.6653
 67  100000     1.63214    1.67277     26    24.7400
 68  100000     1.63470    1.67277     26    24.8226
 69  100000     1.63705    1.67277     26    24.8983
 70  100000     1.63920    1.67277     26    24.9673
 71  100000     1.64126    1.67277     26    25.0335
 72  100000     1.64331    1.67277     26    25.0995
 73  100000     1.64500    1.67277     26    25.1539
 74  100000     1.64677    1.67277     26    25.2107
 75  100000     1.64844    1.67277     26    25.2640
 76  100000     1.64987    1.67277     26    25.3098
 77  100000     1.65127    1.67277     26    25.3542
 78  100000     1.65250    1.67277     26    25.3937
 79  100000     1.65370    1.67277     26    25.4317
 80  100000     1.65481    1.67277     26    25.4671
 81  100000     1.65580    1.67277     26    25.4985
 82  100000     1.65674    1.67277     26    25.5283
 83  100000     1.65750    1.67277     26    25.5523
 84  100000     1.65808    1.67277     26    25.5709
 85  100000     1.65883    1.67277     26    25.5949
 86  100000     1.65959    1.67277     26    25.6187
 87  100000     1.66036    1.67277     26    25.6430
 88  100000     1.66097    1.67277     26    25.6624
 89  100000     1.66167    1.67277     26    25.6843
 90  100000     1.66227    1.67277     26    25.7035
 91  100000     1.66269    1.67277     26    25.7167
 92  100000     1.66315    1.67277     26    25.7314
 93  100000     1.66370    1.67277     26    25.7486
 94  100000     1.66419    1.67277     26    25.7642
 95  100000     1.66459    1.67277     26    25.7767
 96  100000     1.66492    1.67277     26    25.7870
 97  100000     1.66526    1.67277     26    25.7977
 98  100000     1.66553    1.67277     26    25.8062
 99  100000     1.66595    1.67277     26    25.8193
100  100000     1.66622    1.67277     26    25.8277
101  100000     1.66647    1.67277     26    25.8357
102  100000     1.66669    1.67277     26    25.8426
103  100000     1.66696    1.67277     26    25.8509
104  100000     1.66722    1.67277     26    25.8592
105  100000     1.66748    1.67277     26    25.8673
106  100000     1.66770    1.67277     26    25.8741
107  100000     1.66791    1.67277     26    25.8807
108  100000     1.66815    1.67277     26    25.8882
109  100000     1.66820    1.67277     26    25.8897
110  100000     1.66830    1.67277     26    25.8929
111  100000     1.66842    1.67277     26    25.8966
112  100000     1.66858    1.67277     26    25.9018
113  100000     1.66867    1.67277     26    25.9044
114  100000     1.66877    1.67277     26    25.9077
115  100000     1.66888    1.67277     26    25.9109
116  100000     1.66902    1.67277     26    25.9154
117  100000     1.66918    1.67277     26    25.9205
118  100000     1.66922    1.67277     26    25.9219
119  100000     1.66933    1.67277     26    25.9252
120  100000     1.66943    1.67277     26    25.9284
121  100000     1.66957    1.67277     26    25.9328
122  100000     1.66969    1.67277     26    25.9363
123  100000     1.66977    1.67277     26    25.9389
124  100000     1.66992    1.67277     26    25.9435
125  100000     1.67001    1.67277     26    25.9466
126  100000     1.67010    1.67277     26    25.9493
127  100000     1.67019    1.67277     26    25.9521
128  100000     1.67024    1.67277     26    25.9535
129  100000     1.67031    1.67277     26    25.9556
130  100000     1.67038    1.67277     26    25.9579
131  100000     1.67046    1.67277     26    25.9602
132  100000     1.67053    1.67277     26    25.9625
133  100000     1.67059    1.67277     26    25.9644
134  100000     1.67068    1.67277     26    25.9672
135  100000     1.67073    1.67277     26    25.9686
136  100000     1.67077    1.67277     26    25.9699
137  100000     1.67082    1.67277     26    25.9715
138  100000     1.67085    1.67277     26    25.9723
139  100000     1.67087    1.67277     26    25.9730
140  100000     1.67092    1.67277     26    25.9745
141  100000     1.67091    1.67277     26    25.9743
142  100000     1.67094    1.67277     26    25.9755
143  100000     1.67096    1.67277     26    25.9759
144  100000     1.67099    1.67277     26    25.9769
145  100000     1.67104    1.67277     26    25.9784
146  100000     1.67107    1.67277     26    25.9796
147  100000     1.67110    1.67277     26    25.9805
148  100000     1.67114    1.67277     26    25.9816
149  100000     1.67117    1.67277     26    25.9828
150  100000     1.67120    1.67277     26    25.9836
151  100000     1.67122    1.67277     26    25.9843
152  100000     1.67123    1.67277     26    25.9846
153  100000     1.67123    1.67277     26    25.9845
154  100000     1.67125    1.67277     26    25.9852
155  100000     1.67125    1.67277     26    25.9853
156  100000     1.67129    1.67277     26    25.9866
157  100000     1.67132    1.67277     26    25.9873
158  100000     1.67134    1.67277     26    25.9881
159  100000     1.67136    1.67277     26    25.9885
160  100000     1.67135    1.67277     26    25.9883
161  100000     1.67137    1.67277     26    25.9889
162  100000     1.67139    1.67277     26    25.9895
163  100000     1.67140    1.67277     26    25.9898
164  100000     1.67140    1.67277     26    25.9899
165  100000     1.67142    1.67277     26    25.9903
166  100000     1.67142    1.67277     26    25.9904
167  100000     1.67142    1.67277     26    25.9903
168  100000     1.67143    1.67277     26    25.9907
169  100000     1.67144    1.67277     26    25.9909
170  100000     1.67147    1.67277     26    25.9920
171  100000     1.67149    1.67277     26    25.9927
172  100000     1.67150    1.67277     26    25.9930
173  100000     1.67151    1.67277     26    25.9932
174  100000     1.67152    1.67277     26    25.9935
175  100000     1.67152    1.67277     26    25.9936
176  100000     1.67153    1.67277     26    25.9940
177  100000     1.67154    1.67277     26    25.9943
178  100000     1.67153    1.67277     26    25.9940
179  100000     1.67154    1.67277     26    25.9943
180  100000     1.67154    1.67277     26    25.9944
181  100000     1.67154    1.67277     26    25.9944
182  100000     1.67156    1.67277     26    25.9949
183  100000     1.67156    1.67277     26    25.9949
184  100000     1.67156    1.67277     26    25.9949
185  100000     1.67157    1.67277     26    25.9950
186  100000     1.67158    1.67277     26    25.9952
187  100000     1.67158    1.67277     26    25.9955
188  100000     1.67159    1.67277     26    25.9959
189  100000     1.67160    1.67277     26    25.9960
190  100000     1.67159    1.67277     26    25.9958
191  100000     1.67159    1.67277     26    25.9958
192  100000     1.67161    1.67277     26    25.9964
193  100000     1.67162    1.67277     26    25.9967
194  100000     1.67163    1.67277     26    25.9970
195  100000     1.67162    1.67277     26    25.9969
196  100000     1.67164    1.67277     26    25.9974
197  100000     1.67165    1.67277     26    25.9975
198  100000     1.67166    1.67277     26    25.9978
199  100000     1.67166    1.67277     26    25.9979
200  100000     1.67167    1.67277     26    25.9982
201  100000     1.67167    1.67277     26    25.9981
202  100000     1.67167    1.67277     26    25.9984
203  100000     1.67168    1.67277     26    25.9985
204  100000     1.67168    1.67277     26    25.9986
205  100000     1.67167    1.67277     26    25.9985
206  100000     1.67168    1.67277     26    25.9986
207  100000     1.67168    1.67277     26    25.9987
208  100000     1.67168    1.67277     26    25.9987
209  100000     1.67169    1.67277     26    25.9988
210  100000     1.67169    1.67277     26    25.9989
211  100000     1.67169    1.67277     26    25.9990
212  100000     1.67169    1.67277     26    25.9990
213  100000     1.67169    1.67277     26    25.9989
214  100000     1.67169    1.67277     26    25.9990
215  100000     1.67170    1.67277     26    25.9992
216  100000     1.67170    1.67277     26    25.9995
217  100000     1.67170    1.67277     26    25.9994
218  100000     1.67170    1.67277     26    25.9996
219  100000     1.67170    1.67277     26    25.9995
220  100000     1.67170    1.67277     26    25.9995
221  100000     1.67171    1.67277     26    25.9997
222  100000     1.67170    1.67277     26    25.9997
223  100000     1.67170    1.67277     26    25.9996
224  100000     1.67170    1.67277     26    25.9997
225  100000     1.67170    1.67277     26    25.9996
226  100000     1.67170    1.67277     26    25.9997
227  100000     1.67170    1.67277     26    25.9996
228  100000     1.67171    1.67277     26    25.9997
229  100000     1.67171    1.67277     26    25.9998
230  100000     1.67171    1.67277     26    25.9998
231  100000     1.67171    1.67277     26    25.9998
232  100000     1.67172    1.67277     26    25.9998
233  100000     1.67172    1.67277     26    25.9999
234  100000     1.67172    1.67277     26    25.9998
235  100000     1.67172    1.67277     26    25.9999
236  100000     1.67172    1.67277     26    25.9999
237  100000     1.67171    1.67277     26    25.9999
238  100000     1.67171    1.67277     26    25.9998
239  100000     1.67171    1.67277     26    25.9999
240  100000     1.67172    1.67277     26    25.9999
241  100000     1.67171    1.67277     26    25.9999
242  100000     1.67171    1.67277     26    25.9999
243  100000     1.67171    1.67277     26    25.9999
244  100000     1.67171    1.67277     26    25.9998
245  100000     1.67171    1.67277     26    25.9998
246  100000     1.67171    1.67277     26    25.9997
247  100000     1.67171    1.67277     26    25.9998
248  100000     1.67171    1.67277     26    25.9998
249  100000     1.67171    1.67277     26    25.9998
250  100000     1.67171    1.67277     26    25.9998
251  100000     1.67171    1.67277     26    25.9998
252  100000     1.67171    1.67277     26    25.9997
253  100000     1.67171    1.67277     26    25.9998
254  100000     1.67171    1.67277     26    25.9998
255  100000     1.67172    1.67277     26    25.9998
256  100000     1.67171    1.67277     26    25.9997
257  100000     1.67171    1.67277     26    25.9998
258  100000     1.67172    1.67277     26    25.9998
259  100000     1.67172    1.67277     26    25.9998
260  100000     1.67171    1.67277     26    25.9997
261  100000     1.67171    1.67277     26    25.9996
262  100000     1.67171    1.67277     26    25.9996
263  100000     1.67171    1.67277     26    25.9996
264  100000     1.67172    1.67277     26    25.9997
265  100000     1.67172    1.67277     26    25.9997
266  100000     1.67172    1.67277     26    25.9997
267  100000     1.67172    1.67277     26    25.9998
268  100000     1.67172    1.67277     26    25.9999
269  100000     1.67172    1.67277     26    25.9999
270  100000     1.67172    1.67277     26    25.9999
271  100000     1.67172    1.67277     26    25.9999
272  100000     1.67173    1.67277     26    26.0000
273  100000     1.67173    1.67277     26    26.0000
274  100000     1.67173    1.67277     26    26.0000
275  100000     1.67173    1.67277     26    26.0000
276  100000     1.67173    1.67277     26    26.0000
277  100000     1.67173    1.67277     26    26.0000
278  100000     1.67173    1.67277     26    26.0000
279  100000     1.67173    1.67277     26    26.0000
280  100000     1.67173    1.67277     26    26.0000
281  100000     1.67173    1.67277     26    26.0000
282  100000     1.67173    1.67277     26    26.0000
283  100000     1.67173    1.67277     26    26.0000
284  100000     1.67173    1.67277     26    26.0000
285  100000     1.67173    1.67277     26    26.0000
286  100000     1.67173    1.67277     26    26.0000
287  100000     1.67173    1.67277     26    26.0000
288  100000     1.67173    1.67277     26    26.0000
289  100000     1.67173    1.67277     26    26.0000
290  100000     1.67173    1.67277     26    26.0000
291  100000     1.67173    1.67277     26    26.0000
292  100000     1.67173    1.67277     26    26.0000
293  100000     1.67173    1.67277     26    26.0000
294  100000     1.67173    1.67277     26    26.0000
295  100000     1.67173    1.67277     26    26.0000
296  100000     1.67173    1.67277     26    26.0000
297  100000     1.67173    1.67277     26    26.0000
298  100000     1.67173    1.67277     26    26.0000
299  100000     1.67173    1.67277     26    26.0000
300  100000     1.67173    1.67277     26    26.0000
301  100000     1.67173    1.67277     26    26.0000
302  100000     1.67173    1.67277     26    26.0000
303  100000     1.67173    1.67277     26    26.0000
304  100000     1.67173    1.67277     26    26.0000
305  100000     1.67173    1.67277     26    26.0000
306  100000     1.67173    1.67277     26    26.0000
307  100000     1.67174    1.67277     26    26.0000
308  100000     1.67174    1.67277     26    26.0000
309  100000     1.67174    1.67277     26    26.0000
310  100000     1.67174    1.67277     26    26.0000
311  100000     1.67173    1.67277     26    26.0000
312  100000     1.67173    1.67277     26    26.0000
313  100000     1.67173    1.67277     26    26.0000
314  100000     1.67173    1.67277     26    26.0000
315  100000     1.67173    1.67277     26    26.0000
316  100000     1.67173    1.67277     26    26.0000
317  100000     1.67173    1.67277     26    26.0000
318  100000     1.67173    1.67277     26    26.0000
319  100000     1.67173    1.67277     26    26.0000
320  100000     1.67173    1.67277     26    26.0000
321  100000     1.67173    1.67277     26    26.0000
322  100000     1.67173    1.67277     26    26.0000
323  100000     1.67173    1.67277     26    26.0000
324  100000     1.67173    1.67277     26    26.0000
325  100000     1.67173    1.67277     26    26.0000
326  100000     1.67173    1.67277     26    26.0000
327  100000     1.67173    1.67277     26    26.0000
328  100000     1.67173    1.67277     26    26.0000
329  100000     1.67173    1.67277     26    26.0000
330  100000     1.67173    1.67277     26    26.0000
331  100000     1.67173    1.67277     26    26.0000
332  100000     1.67173    1.67277     26    26.0000
333  100000     1.67173    1.67277     26    26.0000
334  100000     1.67173    1.67277     26    26.0000
335  100000     1.67173    1.67277     26    26.0000
336  100000     1.67173    1.67277     26    26.0000
337  100000     1.67173    1.67277     26    26.0000
338  100000     1.67173    1.67277     26    26.0000
339  100000     1.67173    1.67277     26    26.0000
340  100000     1.67173    1.67277     26    26.0000
341  100000     1.67173    1.67277     26    26.0000
342  100000     1.67173    1.67277     26    26.0000
343  100000     1.67173    1.67277     26    26.0000
344  100000     1.67173    1.67277     26    26.0000
345  100000     1.67173    1.67277     26    26.0000
346  100000     1.67173    1.67277     26    26.0000
347  100000     1.67173    1.67277     26    26.0000
348  100000     1.67173    1.67277     26    26.0000
349  100000     1.67173    1.67277     26    26.0000
350  100000     1.67173    1.67277     26    26.0000
351  100000     1.67173    1.67277     26    26.0000
352  100000     1.67173    1.67277     26    26.0000
353  100000     1.67173    1.67277     26    26.0000
354  100000     1.67173    1.67277     26    26.0000
355  100000     1.67173    1.67277     26    26.0000
356  100000     1.67173    1.67277     26    26.0000
357  100000     1.67173    1.67277     26    26.0000
358  100000     1.67173    1.67277     26    26.0000
359  100000     1.67173    1.67277     26    26.0000
360  100000     1.67173    1.67277     26    26.0000
361  100000     1.67173    1.67277     26    26.0000
362  100000     1.67173    1.67277     26    26.0000
363  100000     1.67173    1.67277     26    26.0000
364  100000     1.67173    1.67277     26    26.0000
365  100000     1.67173    1.67277     26    26.0000
366  100000     1.67173    1.67277     26    26.0000
367  100000     1.67173    1.67277     26    26.0000
368  100000     1.67173    1.67277     26    26.0000
369  100000     1.67173    1.67277     26    26.0000
370  100000     1.67173    1.67277     26    26.0000
371  100000     1.67173    1.67277     26    26.0000
372  100000     1.67173    1.67277     26    26.0000
373  100000     1.67173    1.67277     26    26.0000
374  100000     1.67173    1.67277     26    26.0000
375  100000     1.67173    1.67277     26    26.0000
376  100000     1.67173    1.67277     26    26.0000
377  100000     1.67173    1.67277     26    26.0000
378  100000     1.67173    1.67277     26    26.0000
379  100000     1.67173    1.67277     26    26.0000
380  100000     1.67173    1.67277     26    26.0000
381  100000     1.67172    1.67277     26    26.0000
382  100000     1.67173    1.67277     26    26.0000
383  100000     1.67173    1.67277     26    26.0000
384  100000     1.67173    1.67277     26    26.0000
385  100000     1.67173    1.67277     26    26.0000
386  100000     1.67173    1.67277     26    26.0000
387  100000     1.67173    1.67277     26    26.0000
388  100000     1.67173    1.67277     26    26.0000
389  100000     1.67173    1.67277     26    26.0000
390  100000     1.67173    1.67277     26    26.0000
391  100000     1.67173    1.67277     26    26.0000
392  100000     1.67173    1.67277     26    26.0000
393  100000     1.67173    1.67277     26    26.0000
394  100000     1.67173    1.67277     26    26.0000
395  100000     1.67173    1.67277     26    26.0000
396  100000     1.67173    1.67277     26    26.0000
397  100000     1.67173    1.67277     26    26.0000
398  100000     1.67173    1.67277     26    26.0000
399  100000     1.67173    1.67277     26    26.0000
400  100000     1.67173    1.67277     26    26.0000

Asexual Model with selection (random number based on
relative fitness to determine who survives).  Mutation rate is      1e-08,
Recomb rate is ONE per asexual individual per generation
Starting values at generation   0 are; population size 100000,
with avg fitness of    1.22146.

gen  popSize    avgFit     maxFit maxAlleles avgAlleles
  0  100000     1.22146    1.74035     28    10.0120
  1  100000     1.22146    1.74035     28    10.0120
  2  100000     1.22608    1.74035     28    10.2008
  3  100000     1.23095    1.74035     28    10.3992
  4  100000     1.23604    1.74035     28    10.6055
  5  100000     1.24076    1.74035     28    10.7964
  6  100000     1.24620    1.74035     28    11.0154
  7  100000     1.25108    1.74035     28    11.2110
  8  100000     1.25643    1.74035     28    11.4243
  9  100000     1.26200    1.74035     28    11.6456
 10  100000     1.26746    1.74035     28    11.8622
 11  100000     1.27318    1.74035     28    12.0882
 12  100000     1.27912    1.74035     28    12.3211
 13  100000     1.28538    1.74035     28    12.5655
 14  100000     1.29136    1.74035     28    12.7970
 15  100000     1.29755    1.74035     28    13.0366
 16  100000     1.30418    1.74035     28    13.2912
 17  100000     1.31123    1.74035     28    13.5596
 18  100000     1.31836    1.74035     28    13.8296
 19  100000     1.32597    1.74035     28    14.1163
 20  100000     1.33351    1.74035     28    14.3989
 21  100000     1.34126    1.74035     28    14.6873
 22  100000     1.34936    1.74035     28    14.9866
 23  100000     1.35792    1.74035     28    15.3010
 24  100000     1.36667    1.74035     28    15.6203
 25  100000     1.37626    1.74035     28    15.9659
 26  100000     1.38609    1.74035     28    16.3172
 27  100000     1.39702    1.74035     28    16.7047
 28  100000     1.40784    1.74035     28    17.0857
 29  100000     1.41949    1.74035     28    17.4938
 30  100000     1.43111    1.74035     28    17.8958
 31  100000     1.44394    1.74035     28    18.3370
 32  100000     1.45727    1.74035     28    18.7941
 33  100000     1.47066    1.74035     28    19.2497
 34  100000     1.48484    1.74035     28    19.7285
 35  100000     1.49986    1.74035     28    20.2331
 36  100000     1.51495    1.74035     28    20.7365
 37  100000     1.53057    1.74035     28    21.2547
 38  100000     1.54617    1.74035     28    21.7707
 39  100000     1.56131    1.74035     28    22.2701
 40  100000     1.57631    1.74035     28    22.7620
 41  100000     1.59122    1.74035     28    23.2488
 42  100000     1.60434    1.74035     28    23.6761
 43  100000     1.61739    1.74035     28    24.1001
 44  100000     1.63000    1.74035     28    24.5074
 45  100000     1.64120    1.74035     28    24.8684
 46  100000     1.65148    1.74035     28    25.1988
 47  100000     1.66099    1.74035     28    25.5038
 48  100000     1.67035    1.74035     28    25.8026
 49  100000     1.67831    1.74035     28    26.0561
 50  100000     1.68515    1.74035     28    26.2736
 51  100000     1.69119    1.74035     28    26.4655
 52  100000     1.69658    1.74035     28    26.6362
 53  100000     1.70134    1.74035     28    26.7863
 54  100000     1.70559    1.74035     28    26.9203
 55  100000     1.70976    1.74035     28    27.0507
 56  100000     1.71333    1.74035     28    27.1632
 57  100000     1.71663    1.74035     28    27.2665
 58  100000     1.71908    1.74035     28    27.3428
 59  100000     1.72146    1.74035     28    27.4171
 60  100000     1.72348    1.74035     28    27.4802
 61  100000     1.72517    1.74035     28    27.5328
 62  100000     1.72670    1.74035     28    27.5801
 63  100000     1.72804    1.74035     28    27.6217
 64  100000     1.72934    1.74035     28    27.6622
 65  100000     1.73053    1.74035     28    27.6992
 66  100000     1.73143    1.74035     28    27.7271
 67  100000     1.73221    1.74035     28    27.7511
 68  100000     1.73297    1.74035     28    27.7747
 69  100000     1.73390    1.74035     28    27.8033
 70  100000     1.73451    1.74035     28    27.8222
 71  100000     1.73500    1.74035     28    27.8371
 72  100000     1.73546    1.74035     28    27.8513
 73  100000     1.73593    1.74035     28    27.8656
 74  100000     1.73644    1.74035     28    27.8814
 75  100000     1.73675    1.74035     28    27.8908
 76  100000     1.73705    1.74035     28    27.9001
 77  100000     1.73731    1.74035     28    27.9080
 78  100000     1.73751    1.74035     28    27.9139
 79  100000     1.73772    1.74035     28    27.9204
 80  100000     1.73803    1.74035     28    27.9298
 81  100000     1.73817    1.74035     28    27.9342
 82  100000     1.73832    1.74035     28    27.9385
 83  100000     1.73849    1.74035     28    27.9438
 84  100000     1.73860    1.74035     28    27.9471
 85  100000     1.73876    1.74035     28    27.9520
 86  100000     1.73891    1.74035     28    27.9566
 87  100000     1.73902    1.74035     28    27.9599
 88  100000     1.73914    1.74035     28    27.9634
 89  100000     1.73928    1.74035     28    27.9676
 90  100000     1.73934    1.74035     28    27.9694
 91  100000     1.73940    1.74035     28    27.9713
 92  100000     1.73944    1.74035     28    27.9725
 93  100000     1.73954    1.74035     28    27.9756
 94  100000     1.73964    1.74035     28    27.9785
 95  100000     1.73970    1.74035     28    27.9802
 96  100000     1.73976    1.74035     28    27.9822
 97  100000     1.73980    1.74035     28    27.9835
 98  100000     1.73985    1.74035     28    27.9849
 99  100000     1.73986    1.74035     28    27.9853
100  100000     1.73987    1.74035     28    27.9855
101  100000     1.73991    1.74035     28    27.9867
102  100000     1.73997    1.74035     28    27.9886
103  100000     1.74000    1.74035     28    27.9893
104  100000     1.74003    1.74035     28    27.9903
105  100000     1.74006    1.74035     28    27.9913
106  100000     1.74008    1.74035     28    27.9918
107  100000     1.74011    1.74035     28    27.9926
108  100000     1.74012    1.74035     28    27.9929
109  100000     1.74015    1.74035     28    27.9939
110  100000     1.74017    1.74035     28    27.9944
111  100000     1.74019    1.74035     28    27.9950
112  100000     1.74021    1.74035     28    27.9956
113  100000     1.74023    1.74035     28    27.9961
114  100000     1.74025    1.74035     28    27.9968
115  100000     1.74025    1.74035     28    27.9968
116  100000     1.74025    1.74035     28    27.9968
117  100000     1.74025    1.74035     28    27.9969
118  100000     1.74025    1.74035     28    27.9969
119  100000     1.74024    1.74035     28    27.9966
120  100000     1.74025    1.74035     28    27.9969
121  100000     1.74025    1.74035     28    27.9969
122  100000     1.74026    1.74035     28    27.9971
123  100000     1.74026    1.74035     28    27.9973
124  100000     1.74026    1.74035     28    27.9971
125  100000     1.74026    1.74035     28    27.9972
126  100000     1.74026    1.74035     28    27.9972
127  100000     1.74026    1.74035     28    27.9973
128  100000     1.74026    1.74035     28    27.9973
129  100000     1.74027    1.74035     28    27.9974
130  100000     1.74026    1.74035     28    27.9971
131  100000     1.74025    1.74035     28    27.9969
132  100000     1.74025    1.74035     28    27.9970
133  100000     1.74027    1.74035     28    27.9976
134  100000     1.74028    1.74035     28    27.9978
135  100000     1.74029    1.74035     28    27.9980
136  100000     1.74030    1.74035     28    27.9983
137  100000     1.74029    1.74035     28    27.9981
138  100000     1.74029    1.74035     28    27.9980
139  100000     1.74029    1.74035     28    27.9982
140  100000     1.74031    1.74035     28    27.9986
141  100000     1.74031    1.74035     28    27.9986
142  100000     1.74030    1.74035     28    27.9984
143  100000     1.74031    1.74035     28    27.9986
144  100000     1.74031    1.74035     28    27.9986
145  100000     1.74030    1.74035     28    27.9984
146  100000     1.74030    1.74035     28    27.9983
147  100000     1.74030    1.74035     28    27.9984
148  100000     1.74031    1.74035     28    27.9987
149  100000     1.74031    1.74035     28    27.9988
150  100000     1.74032    1.74035     28    27.9988
151  100000     1.74031    1.74035     28    27.9988
152  100000     1.74031    1.74035     28    27.9988
153  100000     1.74032    1.74035     28    27.9991
154  100000     1.74032    1.74035     28    27.9991
155  100000     1.74032    1.74035     28    27.9990
156  100000     1.74032    1.74035     28    27.9991
157  100000     1.74033    1.74035     28    27.9991
158  100000     1.74033    1.74035     28    27.9992
159  100000     1.74033    1.74035     28    27.9993
160  100000     1.74033    1.74035     28    27.9994
161  100000     1.74033    1.74035     28    27.9994
162  100000     1.74034    1.74035     28    27.9995
163  100000     1.74034    1.74035     28    27.9995
164  100000     1.74034    1.74035     28    27.9995
165  100000     1.74034    1.74035     28    27.9995
166  100000     1.74034    1.74035     28    27.9996
167  100000     1.74034    1.74035     28    27.9997
168  100000     1.74034    1.74035     28    27.9996
169  100000     1.74034    1.74035     28    27.9996
170  100000     1.74034    1.74035     28    27.9996
171  100000     1.74034    1.74035     28    27.9997
172  100000     1.74034    1.74035     28    27.9996
173  100000     1.74034    1.74035     28    27.9996
174  100000     1.74035    1.74035     28    27.9998
175  100000     1.74035    1.74035     28    27.9998
176  100000     1.74035    1.74035     28    27.9998
177  100000     1.74035    1.74035     28    27.9998
178  100000     1.74035    1.74035     28    27.9998
179  100000     1.74035    1.74035     28    27.9999
180  100000     1.74035    1.74035     28    27.9998
181  100000     1.74035    1.74035     28    27.9999
182  100000     1.74035    1.74035     28    27.9999
183  100000     1.74035    1.74035     28    27.9999
184  100000     1.74035    1.74035     28    28.0000
185  100000     1.74035    1.74035     28    27.9999
186  100000     1.74035    1.74035     28    27.9999
187  100000     1.74035    1.74035     28    27.9999
188  100000     1.74035    1.74035     28    27.9999
189  100000     1.74035    1.74035     28    27.9999
190  100000     1.74035    1.74035     28    27.9999
191  100000     1.74035    1.74035     28    28.0000
192  100000     1.74035    1.74035     28    28.0000
193  100000     1.74035    1.74035     28    28.0000
194  100000     1.74035    1.74035     28    28.0000
195  100000     1.74035    1.74035     28    28.0000
196  100000     1.74035    1.74035     28    28.0000
197  100000     1.74035    1.74035     28    28.0000
198  100000     1.74035    1.74035     28    28.0000
199  100000     1.74035    1.74035     28    28.0000
200  100000     1.74035    1.74035     28    28.0000
201  100000     1.74035    1.74035     28    28.0000
202  100000     1.74035    1.74035     28    28.0000
203  100000     1.74035    1.74035     28    28.0000
204  100000     1.74035    1.74035     28    28.0000
205  100000     1.74035    1.74035     28    28.0000
206  100000     1.74035    1.74035     28    28.0000
207  100000     1.74035    1.74035     28    28.0000
208  100000     1.74035    1.74035     28    28.0000
209  100000     1.74035    1.74035     28    28.0000
210  100000     1.74035    1.74035     28    28.0000
211  100000     1.74035    1.74035     28    28.0000
212  100000     1.74035    1.74035     28    28.0000
213  100000     1.74035    1.74035     28    28.0000
214  100000     1.74035    1.74035     28    28.0000
215  100000     1.74035    1.74035     28    28.0000
216  100000     1.74035    1.74035     28    28.0000
217  100000     1.74035    1.74035     28    28.0000
218  100000     1.74035    1.74035     28    28.0000
219  100000     1.74035    1.74035     28    28.0000
220  100000     1.74035    1.74035     28    28.0000
221  100000     1.74035    1.74035     28    28.0000
222  100000     1.74035    1.74035     28    28.0000
223  100000     1.74035    1.74035     28    28.0000
224  100000     1.74035    1.74035     28    28.0000
225  100000     1.74035    1.74035     28    28.0000
226  100000     1.74035    1.74035     28    28.0000
227  100000     1.74035    1.74035     28    28.0000
228  100000     1.74035    1.74035     28    28.0000
229  100000     1.74035    1.74035     28    28.0000
230  100000     1.74035    1.74035     28    28.0000
231  100000     1.74035    1.74035     28    28.0000
232  100000     1.74035    1.74035     28    28.0000
233  100000     1.74035    1.74035     28    28.0000
234  100000     1.74035    1.74035     28    28.0000
235  100000     1.74035    1.74035     28    28.0000
236  100000     1.74035    1.74035     28    28.0000
237  100000     1.74035    1.74035     28    28.0000
238  100000     1.74035    1.74035     28    28.0000
239  100000     1.74035    1.74035     28    28.0000
240  100000     1.74035    1.74035     28    28.0000
241  100000     1.74035    1.74035     28    28.0000
242  100000     1.74035    1.74035     28    28.0000
243  100000     1.74035    1.74035     28    28.0000
244  100000     1.74035    1.74035     28    28.0000
245  100000     1.74035    1.74035     28    28.0000
246  100000     1.74035    1.74035     28    28.0000
247  100000     1.74035    1.74035     28    28.0000
248  100000     1.74035    1.74035     28    28.0000
249  100000     1.74035    1.74035     28    28.0000
250  100000     1.74035    1.74035     28    28.0000
251  100000     1.74035    1.74035     28    28.0000
252  100000     1.74035    1.74035     28    28.0000
253  100000     1.74035    1.74035     28    28.0000
254  100000     1.74035    1.74035     28    28.0000
255  100000     1.74035    1.74035     28    28.0000
256  100000     1.74035    1.74035     28    28.0000
257  100000     1.74035    1.74035     28    28.0000
258  100000     1.74035    1.74035     28    28.0000
259  100000     1.74035    1.74035     28    28.0000
260  100000     1.74035    1.74035     28    28.0000
261  100000     1.74035    1.74035     28    28.0000
262  100000     1.74035    1.74035     28    28.0000
263  100000     1.74035    1.74035     28    28.0000
264  100000     1.74035    1.74035     28    28.0000
265  100000     1.74035    1.74035     28    28.0000
266  100000     1.74035    1.74035     28    28.0000
267  100000     1.74035    1.74035     28    28.0000
268  100000     1.74035    1.74035     28    28.0000
269  100000     1.74035    1.74035     28    28.0000
270  100000     1.74035    1.74035     28    28.0000
271  100000     1.74035    1.74035     28    28.0000
272  100000     1.74035    1.74035     28    28.0000
273  100000     1.74035    1.74035     28    28.0000
274  100000     1.74035    1.74035     28    28.0000
275  100000     1.74035    1.74035     28    28.0000
276  100000     1.74035    1.74035     28    28.0000
277  100000     1.74035    1.74035     28    28.0000
278  100000     1.74035    1.74035     28    28.0000
279  100000     1.74035    1.74035     28    28.0000
280  100000     1.74035    1.74035     28    28.0000
281  100000     1.74035    1.74035     28    28.0000
282  100000     1.74035    1.74035     28    28.0000
283  100000     1.74035    1.74035     28    28.0000
284  100000     1.74035    1.74035     28    28.0000
285  100000     1.74035    1.74035     28    28.0000
286  100000     1.74035    1.74035     28    28.0000
287  100000     1.74035    1.74035     28    28.0000
288  100000     1.74035    1.74035     28    28.0000
289  100000     1.74035    1.74035     28    28.0000
290  100000     1.74035    1.74035     28    28.0000
291  100000     1.74035    1.74035     28    28.0000
292  100000     1.74035    1.74035     28    28.0000
293  100000     1.74035    1.74035     28    28.0000
294  100000     1.74035    1.74035     28    28.0000
295  100000     1.74035    1.74035     28    28.0000
296  100000     1.74035    1.74035     28    28.0000
297  100000     1.74035    1.74035     28    28.0000
298  100000     1.74035    1.74035     28    28.0000
299  100000     1.74035    1.74035     28    28.0000
300  100000     1.74035    1.74035     28    28.0000
301  100000     1.74035    1.74035     28    28.0000
302  100000     1.74035    1.74035     28    28.0000
303  100000     1.74035    1.74035     28    28.0000
304  100000     1.74035    1.74035     28    28.0000
305  100000     1.74035    1.74035     28    28.0000
306  100000     1.74035    1.74035     28    28.0000
307  100000     1.74035    1.74035     28    28.0000
308  100000     1.74035    1.74035     28    28.0000
309  100000     1.74035    1.74035     28    28.0000
310  100000     1.74035    1.74035     28    28.0000
311  100000     1.74035    1.74035     28    28.0000
312  100000     1.74035    1.74035     28    28.0000
313  100000     1.74035    1.74035     28    28.0000
314  100000     1.74035    1.74035     28    28.0000
315  100000     1.74035    1.74035     28    28.0000
316  100000     1.74035    1.74035     28    28.0000
317  100000     1.74035    1.74035     28    28.0000
318  100000     1.74035    1.74035     28    28.0000
319  100000     1.74035    1.74035     28    28.0000
320  100000     1.74035    1.74035     28    28.0000
321  100000     1.74035    1.74035     28    28.0000
322  100000     1.74035    1.74035     28    28.0000
323  100000     1.74035    1.74035     28    28.0000
324  100000     1.74035    1.74035     28    28.0000
325  100000     1.74035    1.74035     28    28.0000
326  100000     1.74035    1.74035     28    28.0000
327  100000     1.74035    1.74035     28    28.0000
328  100000     1.74035    1.74035     28    28.0000
329  100000     1.74035    1.74035     28    28.0000
330  100000     1.74035    1.74035     28    28.0000
331  100000     1.74035    1.74035     28    28.0000
332  100000     1.74035    1.74035     28    28.0000
333  100000     1.74035    1.74035     28    28.0000
334  100000     1.74035    1.74035     28    28.0000
335  100000     1.74035    1.74035     28    28.0000
336  100000     1.74035    1.74035     28    28.0000
337  100000     1.74035    1.74035     28    28.0000
338  100000     1.74035    1.74035     28    28.0000
339  100000     1.74035    1.74035     28    28.0000
340  100000     1.74035    1.74035     28    28.0000
341  100000     1.74035    1.74035     28    28.0000
342  100000     1.74035    1.74035     28    28.0000
343  100000     1.74035    1.74035     28    28.0000
344  100000     1.74035    1.74035     28    28.0000
345  100000     1.74035    1.74035     28    28.0000
346  100000     1.74035    1.74035     28    28.0000
347  100000     1.74035    1.74035     28    28.0000
348  100000     1.74035    1.74035     28    28.0000
349  100000     1.74035    1.74035     28    28.0000
350  100000     1.74035    1.74035     28    28.0000
351  100000     1.74035    1.74035     28    28.0000
352  100000     1.74035    1.74035     28    28.0000
353  100000     1.74035    1.74035     28    28.0000
354  100000     1.74035    1.74035     28    28.0000
355  100000     1.74035    1.74035     28    28.0000
356  100000     1.74035    1.74035     28    28.0000
357  100000     1.74035    1.74035     28    28.0000
358  100000     1.74035    1.74035     28    28.0000
359  100000     1.74035    1.74035     28    28.0000
360  100000     1.74035    1.74035     28    28.0000
361  100000     1.74035    1.74035     28    28.0000
362  100000     1.74035    1.74035     28    28.0000
363  100000     1.74035    1.74035     28    28.0000
364  100000     1.74035    1.74035     28    28.0000
365  100000     1.74035    1.74035     28    28.0000
366  100000     1.74035    1.74035     28    28.0000
367  100000     1.74035    1.74035     28    28.0000
368  100000     1.74035    1.74035     28    28.0000
369  100000     1.74035    1.74035     28    28.0000
370  100000     1.74035    1.74035     28    28.0000
371  100000     1.74035    1.74035     28    28.0000
372  100000     1.74035    1.74035     28    28.0000
373  100000     1.74035    1.74035     28    28.0000
374  100000     1.74035    1.74035     28    28.0000
375  100000     1.74035    1.74035     28    28.0000
376  100000     1.74035    1.74035     28    28.0000
377  100000     1.74035    1.74035     28    28.0000
378  100000     1.74035    1.74035     28    28.0000
379  100000     1.74035    1.74035     28    28.0000
380  100000     1.74035    1.74035     28    28.0000
381  100000     1.74035    1.74035     28    28.0000
382  100000     1.74035    1.74035     28    28.0000
383  100000     1.74035    1.74035     28    28.0000
384  100000     1.74035    1.74035     28    28.0000
385  100000     1.74035    1.74035     28    28.0000
386  100000     1.74035    1.74035     28    28.0000
387  100000     1.74035    1.74035     28    28.0000
388  100000     1.74035    1.74035     28    28.0000
389  100000     1.74035    1.74035     28    28.0000
390  100000     1.74035    1.74035     28    28.0000
391  100000     1.74035    1.74035     28    28.0000
392  100000     1.74035    1.74035     28    28.0000
393  100000     1.74035    1.74035     28    28.0000
394  100000     1.74035    1.74035     28    28.0000
395  100000     1.74035    1.74035     28    28.0000
396  100000     1.74035    1.74035     28    28.0000
397  100000     1.74035    1.74035     28    28.0000
398  100000     1.74035    1.74035     28    28.0000
399  100000     1.74035    1.74035     28    28.0000
400  100000     1.74035    1.74035     28    28.0000

Asexual Model with selection (random number based on
relative fitness to determine who survives).  Mutation rate is      1e-08,
Recomb rate is ONE per asexual individual per generation
Starting values at generation   0 are; population size 100000,
with avg fitness of    1.22141.

gen  popSize    avgFit     maxFit maxAlleles avgAlleles
  0  100000     1.22141    1.63934     25    10.0107
  1  100000     1.22141    1.63934     25    10.0107
  2  100000     1.22606    1.63934     25    10.2007
  3  100000     1.23098    1.63934     25    10.4019
  4  100000     1.23562    1.63934     25    10.5903
  5  100000     1.24052    1.63934     25    10.7877
  6  100000     1.24583    1.63934     25    11.0014
  7  100000     1.25091    1.63934     25    11.2052
  8  100000     1.25618    1.63934     25    11.4161
  9  100000     1.26171    1.63934     25    11.6365
 10  100000     1.26726    1.63934     25    11.8575
 11  100000     1.27262    1.63934     25    12.0693
 12  100000     1.27838    1.63934     25    12.2958
 13  100000     1.28423    1.63934     25    12.5247
 14  100000     1.29016    1.63934     25    12.7563
 15  100000     1.29630    1.63934     25    12.9952
 16  100000     1.30246    1.63934     25    13.2335
 17  100000     1.30861    1.63934     25    13.4695
 18  100000     1.31459    1.63934     25    13.6990
 19  100000     1.32105    1.63934     25    13.9458
 20  100000     1.32742    1.63934     25    14.1870
 21  100000     1.33400    1.63934     25    14.4359
 22  100000     1.34074    1.63934     25    14.6900
 23  100000     1.34743    1.63934     25    14.9401
 24  100000     1.35439    1.63934     25    15.1999
 25  100000     1.36150    1.63934     25    15.4629
 26  100000     1.36829    1.63934     25    15.7141
 27  100000     1.37571    1.63934     25    15.9867
 28  100000     1.38309    1.63934     25    16.2575
 29  100000     1.39043    1.63934     25    16.5242
 30  100000     1.39797    1.63934     25    16.7984
 31  100000     1.40564    1.63934     25    17.0751
 32  100000     1.41298    1.63934     25    17.3397
 33  100000     1.42036    1.63934     25    17.6045
 34  100000     1.42794    1.63934     25    17.8753
 35  100000     1.43511    1.63934     25    18.1301
 36  100000     1.44209    1.63934     25    18.3771
 37  100000     1.44927    1.63934     25    18.6309
 38  100000     1.45600    1.63934     25    18.8679
 39  100000     1.46316    1.63934     25    19.1188
 40  100000     1.47012    1.63934     25    19.3615
 41  100000     1.47678    1.63934     25    19.5927
 42  100000     1.48331    1.63934     25    19.8193
 43  100000     1.48961    1.63934     25    20.0371
 44  100000     1.49579    1.63934     25    20.2497
 45  100000     1.50197    1.63934     25    20.4608
 46  100000     1.50800    1.63934     25    20.6661
 47  100000     1.51389    1.63934     25    20.8669
 48  100000     1.51975    1.63934     25    21.0659
 49  100000     1.52523    1.63934     25    21.2513
 50  100000     1.53069    1.63934     25    21.4360
 51  100000     1.53581    1.63934     25    21.6086
 52  100000     1.54050    1.63934     25    21.7659
 53  100000     1.54502    1.63934     25    21.9176
 54  100000     1.54952    1.63934     25    22.0677
 55  100000     1.55396    1.63934     25    22.2159
 56  100000     1.55836    1.63934     25    22.3626
 57  100000     1.56237    1.63934     25    22.4959
 58  100000     1.56611    1.63934     25    22.6199
 59  100000     1.56980    1.63934     25    22.7417
 60  100000     1.57289    1.63934     25    22.8439
 61  100000     1.57624    1.63934     25    22.9546
 62  100000     1.57942    1.63934     25    23.0594
 63  100000     1.58257    1.63934     25    23.1632
 64  100000     1.58564    1.63934     25    23.2640
 65  100000     1.58839    1.63934     25    23.3542
 66  100000     1.59108    1.63934     25    23.4426
 67  100000     1.59378    1.63934     25    23.5307
 68  100000     1.59630    1.63934     25    23.6134
 69  100000     1.59849    1.63934     25    23.6847
 70  100000     1.60076    1.63934     25    23.7584
 71  100000     1.60277    1.63934     25    23.8241
 72  100000     1.60469    1.63934     25    23.8863
 73  100000     1.60655    1.63934     25    23.9471
 74  100000     1.60836    1.63934     25    24.0060
 75  100000     1.61015    1.63934     25    24.0639
 76  100000     1.61158    1.63934     25    24.1104
 77  100000     1.61277    1.63934     25    24.1489
 78  100000     1.61411    1.63934     25    24.1922
 79  100000     1.61528    1.63934     25    24.2302
 80  100000     1.61643    1.63934     25    24.2673
 81  100000     1.61769    1.63934     25    24.3082
 82  100000     1.61862    1.63934     25    24.3380
 83  100000     1.61965    1.63934     25    24.3713
 84  100000     1.62072    1.63934     25    24.4059
 85  100000     1.62163    1.63934     25    24.4353
 86  100000     1.62256    1.63934     25    24.4652
 87  100000     1.62328    1.63934     25    24.4883
 88  100000     1.62416    1.63934     25    24.5166
 89  100000     1.62488    1.63934     25    24.5400
 90  100000     1.62562    1.63934     25    24.5638
 91  100000     1.62617    1.63934     25    24.5815
 92  100000     1.62688    1.63934     25    24.6043
 93  100000     1.62750    1.63934     25    24.6244
 94  100000     1.62797    1.63934     25    24.6395
 95  100000     1.62854    1.63934     25    24.6577
 96  100000     1.62909    1.63934     25    24.6756
 97  100000     1.62947    1.63934     25    24.6878
 98  100000     1.62991    1.63934     25    24.7019
 99  100000     1.63031    1.63934     25    24.7147
100  100000     1.63066    1.63934     25    24.7259
101  100000     1.63096    1.63934     25    24.7357
102  100000     1.63131    1.63934     25    24.7468
103  100000     1.63153    1.63934     25    24.7539
104  100000     1.63183    1.63934     25    24.7635
105  100000     1.63214    1.63934     25    24.7733
106  100000     1.63237    1.63934     25    24.7809
107  100000     1.63267    1.63934     25    24.7902
108  100000     1.63294    1.63934     25    24.7989
109  100000     1.63311    1.63934     25    24.8045
110  100000     1.63345    1.63934     25    24.8151
111  100000     1.63365    1.63934     25    24.8217
112  100000     1.63385    1.63934     25    24.8282
113  100000     1.63404    1.63934     25    24.8342
114  100000     1.63424    1.63934     25    24.8405
115  100000     1.63446    1.63934     25    24.8474
116  100000     1.63456    1.63934     25    24.8509
117  100000     1.63478    1.63934     25    24.8578
118  100000     1.63494    1.63934     25    24.8630
119  100000     1.63504    1.63934     25    24.8663
120  100000     1.63522    1.63934     25    24.8720
121  100000     1.63535    1.63934     25    24.8761
122  100000     1.63549    1.63934     25    24.8804
123  100000     1.63563    1.63934     25    24.8851
124  100000     1.63573    1.63934     25    24.8883
125  100000     1.63589    1.63934     25    24.8932
126  100000     1.63599    1.63934     25    24.8965
127  100000     1.63608    1.63934     25    24.8993
128  100000     1.63620    1.63934     25    24.9030
129  100000     1.63631    1.63934     25    24.9066
130  100000     1.63639    1.63934     25    24.9092
131  100000     1.63646    1.63934     25    24.9115
132  100000     1.63655    1.63934     25    24.9144
133  100000     1.63659    1.63934     25    24.9157
134  100000     1.63669    1.63934     25    24.9189
135  100000     1.63674    1.63934     25    24.9205
136  100000     1.63679    1.63934     25    24.9219
137  100000     1.63687    1.63934     25    24.9245
138  100000     1.63691    1.63934     25    24.9257
139  100000     1.63698    1.63934     25    24.9280
140  100000     1.63700    1.63934     25    24.9286
141  100000     1.63707    1.63934     25    24.9307
142  100000     1.63714    1.63934     25    24.9330
143  100000     1.63717    1.63934     25    24.9339
144  100000     1.63721    1.63934     25    24.9353
145  100000     1.63728    1.63934     25    24.9376
146  100000     1.63736    1.63934     25    24.9400
147  100000     1.63747    1.63934     25    24.9434
148  100000     1.63749    1.63934     25    24.9443
149  100000     1.63754    1.63934     25    24.9458
150  100000     1.63758    1.63934     25    24.9469
151  100000     1.63767    1.63934     25    24.9499
152  100000     1.63772    1.63934     25    24.9516
153  100000     1.63777    1.63934     25    24.9529
154  100000     1.63777    1.63934     25    24.9532
155  100000     1.63779    1.63934     25    24.9536
156  100000     1.63783    1.63934     25    24.9550
157  100000     1.63786    1.63934     25    24.9558
158  100000     1.63788    1.63934     25    24.9566
159  100000     1.63793    1.63934     25    24.9581
160  100000     1.63793    1.63934     25    24.9583
161  100000     1.63797    1.63934     25    24.9594
162  100000     1.63802    1.63934     25    24.9610
163  100000     1.63804    1.63934     25    24.9618
164  100000     1.63806    1.63934     25    24.9622
165  100000     1.63811    1.63934     25    24.9638
166  100000     1.63815    1.63934     25    24.9650
167  100000     1.63818    1.63934     25    24.9661
168  100000     1.63820    1.63934     25    24.9667
169  100000     1.63821    1.63934     25    24.9670
170  100000     1.63824    1.63934     25    24.9680
171  100000     1.63827    1.63934     25    24.9690
172  100000     1.63829    1.63934     25    24.9695
173  100000     1.63830    1.63934     25    24.9700
174  100000     1.63832    1.63934     25    24.9704
175  100000     1.63834    1.63934     25    24.9711
176  100000     1.63838    1.63934     25    24.9723
177  100000     1.63840    1.63934     25    24.9729
178  100000     1.63841    1.63934     25    24.9734
179  100000     1.63845    1.63934     25    24.9745
180  100000     1.63846    1.63934     25    24.9749
181  100000     1.63847    1.63934     25    24.9753
182  100000     1.63848    1.63934     25    24.9755
183  100000     1.63850    1.63934     25    24.9762
184  100000     1.63853    1.63934     25    24.9771
185  100000     1.63855    1.63934     25    24.9778
186  100000     1.63856    1.63934     25    24.9781
187  100000     1.63856    1.63934     25    24.9780
188  100000     1.63855    1.63934     25    24.9779
189  100000     1.63855    1.63934     25    24.9780
190  100000     1.63857    1.63934     25    24.9786
191  100000     1.63858    1.63934     25    24.9789
192  100000     1.63860    1.63934     25    24.9795
193  100000     1.63862    1.63934     25    24.9800
194  100000     1.63864    1.63934     25    24.9806
195  100000     1.63865    1.63934     25    24.9811
196  100000     1.63868    1.63934     25    24.9820
197  100000     1.63870    1.63934     25    24.9825
198  100000     1.63872    1.63934     25    24.9831
199  100000     1.63873    1.63934     25    24.9834
200  100000     1.63876    1.63934     25    24.9843
201  100000     1.63877    1.63934     25    24.9846
202  100000     1.63877    1.63934     25    24.9847
203  100000     1.63877    1.63934     25    24.9848
204  100000     1.63877    1.63934     25    24.9847
205  100000     1.63878    1.63934     25    24.9850
206  100000     1.63877    1.63934     25    24.9848
207  100000     1.63876    1.63934     25    24.9846
208  100000     1.63876    1.63934     25    24.9845
209  100000     1.63877    1.63934     25    24.9847
210  100000     1.63877    1.63934     25    24.9848
211  100000     1.63879    1.63934     25    24.9852
212  100000     1.63879    1.63934     25    24.9853
213  100000     1.63880    1.63934     25    24.9856
214  100000     1.63880    1.63934     25    24.9856
215  100000     1.63881    1.63934     25    24.9861
216  100000     1.63884    1.63934     25    24.9870
217  100000     1.63886    1.63934     25    24.9874
218  100000     1.63887    1.63934     25    24.9878
219  100000     1.63887    1.63934     25    24.9880
220  100000     1.63889    1.63934     25    24.9885
221  100000     1.63891    1.63934     25    24.9890
222  100000     1.63891    1.63934     25    24.9892
223  100000     1.63891    1.63934     25    24.9892
224  100000     1.63892    1.63934     25    24.9895
225  100000     1.63892    1.63934     25    24.9894
226  100000     1.63893    1.63934     25    24.9897
227  100000     1.63893    1.63934     25    24.9899
228  100000     1.63894    1.63934     25    24.9900
229  100000     1.63893    1.63934     25    24.9899
230  100000     1.63894    1.63934     25    24.9902
231  100000     1.63894    1.63934     25    24.9900
232  100000     1.63894    1.63934     25    24.9902
233  100000     1.63895    1.63934     25    24.9905
234  100000     1.63896    1.63934     25    24.9907
235  100000     1.63896    1.63934     25    24.9909
236  100000     1.63898    1.63934     25    24.9914
237  100000     1.63899    1.63934     25    24.9915
238  100000     1.63899    1.63934     25    24.9918
239  100000     1.63900    1.63934     25    24.9920
240  100000     1.63901    1.63934     25    24.9922
241  100000     1.63901    1.63934     25    24.9923
242  100000     1.63902    1.63934     25    24.9925
243  100000     1.63901    1.63934     25    24.9924
244  100000     1.63902    1.63934     25    24.9926
245  100000     1.63903    1.63934     25    24.9929
246  100000     1.63904    1.63934     25    24.9933
247  100000     1.63905    1.63934     25    24.9935
248  100000     1.63906    1.63934     25    24.9938
249  100000     1.63907    1.63934     25    24.9942
250  100000     1.63907    1.63934     25    24.9943
251  100000     1.63908    1.63934     25    24.9946
252  100000     1.63909    1.63934     25    24.9949
253  100000     1.63908    1.63934     25    24.9947
254  100000     1.63909    1.63934     25    24.9948
255  100000     1.63909    1.63934     25    24.9950
256  100000     1.63910    1.63934     25    24.9951
257  100000     1.63910    1.63934     25    24.9952
258  100000     1.63911    1.63934     25    24.9954
259  100000     1.63911    1.63934     25    24.9956
260  100000     1.63913    1.63934     25    24.9960
261  100000     1.63913    1.63934     25    24.9961
262  100000     1.63914    1.63934     25    24.9963
263  100000     1.63914    1.63934     25    24.9965
264  100000     1.63915    1.63934     25    24.9966
265  100000     1.63915    1.63934     25    24.9967
266  100000     1.63915    1.63934     25    24.9967
267  100000     1.63915    1.63934     25    24.9968
268  100000     1.63915    1.63934     25    24.9967
269  100000     1.63915    1.63934     25    24.9967
270  100000     1.63916    1.63934     25    24.9969
271  100000     1.63916    1.63934     25    24.9970
272  100000     1.63916    1.63934     25    24.9970
273  100000     1.63917    1.63934     25    24.9972
274  100000     1.63916    1.63934     25    24.9971
275  100000     1.63917    1.63934     25    24.9972
276  100000     1.63917    1.63934     25    24.9972
277  100000     1.63917    1.63934     25    24.9974
278  100000     1.63917    1.63934     25    24.9974
279  100000     1.63917    1.63934     25    24.9973
280  100000     1.63917    1.63934     25    24.9973
281  100000     1.63917    1.63934     25    24.9972
282  100000     1.63917    1.63934     25    24.9973
283  100000     1.63917    1.63934     25    24.9973
284  100000     1.63917    1.63934     25    24.9973
285  100000     1.63917    1.63934     25    24.9973
286  100000     1.63917    1.63934     25    24.9973
287  100000     1.63917    1.63934     25    24.9974
288  100000     1.63918    1.63934     25    24.9976
289  100000     1.63918    1.63934     25    24.9976
290  100000     1.63918    1.63934     25    24.9977
291  100000     1.63918    1.63934     25    24.9977
292  100000     1.63919    1.63934     25    24.9978
293  100000     1.63919    1.63934     25    24.9979
294  100000     1.63919    1.63934     25    24.9980
295  100000     1.63920    1.63934     25    24.9981
296  100000     1.63920    1.63934     25    24.9982
297  100000     1.63920    1.63934     25    24.9984
298  100000     1.63921    1.63934     25    24.9985
299  100000     1.63921    1.63934     25    24.9985
300  100000     1.63921    1.63934     25    24.9987
301  100000     1.63921    1.63934     25    24.9986
302  100000     1.63921    1.63934     25    24.9987
303  100000     1.63921    1.63934     25    24.9986
304  100000     1.63922    1.63934     25    24.9988
305  100000     1.63922    1.63934     25    24.9989
306  100000     1.63922    1.63934     25    24.9989
307  100000     1.63922    1.63934     25    24.9989
308  100000     1.63922    1.63934     25    24.9988
309  100000     1.63922    1.63934     25    24.9989
310  100000     1.63922    1.63934     25    24.9990
311  100000     1.63922    1.63934     25    24.9990
312  100000     1.63922    1.63934     25    24.9990
313  100000     1.63922    1.63934     25    24.9990
314  100000     1.63923    1.63934     25    24.9991
315  100000     1.63923    1.63934     25    24.9992
316  100000     1.63923    1.63934     25    24.9993
317  100000     1.63923    1.63934     25    24.9993
318  100000     1.63923    1.63934     25    24.9993
319  100000     1.63924    1.63934     25    24.9994
320  100000     1.63924    1.63934     25    24.9995
321  100000     1.63924    1.63934     25    24.9995
322  100000     1.63924    1.63934     25    24.9995
323  100000     1.63924    1.63934     25    24.9995
324  100000     1.63924    1.63934     25    24.9995
325  100000     1.63924    1.63934     25    24.9995
326  100000     1.63924    1.63934     25    24.9995
327  100000     1.63924    1.63934     25    24.9996
328  100000     1.63924    1.63934     25    24.9996
329  100000     1.63924    1.63934     25    24.9996
330  100000     1.63924    1.63934     25    24.9996
331  100000     1.63925    1.63934     25    24.9996
332  100000     1.63924    1.63934     25    24.9996
333  100000     1.63924    1.63934     25    24.9996
334  100000     1.63925    1.63934     25    24.9996
335  100000     1.63925    1.63934     25    24.9997
336  100000     1.63925    1.63934     25    24.9997
337  100000     1.63925    1.63934     25    24.9997
338  100000     1.63925    1.63934     25    24.9997
339  100000     1.63925    1.63934     25    24.9997
340  100000     1.63925    1.63934     25    24.9997
341  100000     1.63925    1.63934     25    24.9997
342  100000     1.63925    1.63934     25    24.9997
343  100000     1.63925    1.63934     25    24.9996
344  100000     1.63925    1.63934     25    24.9997
345  100000     1.63925    1.63934     25    24.9997
346  100000     1.63924    1.63934     25    24.9997
347  100000     1.63924    1.63934     25    24.9996
348  100000     1.63924    1.63934     25    24.9996
349  100000     1.63924    1.63934     25    24.9996
350  100000     1.63925    1.63934     25    24.9996
351  100000     1.63925    1.63934     25    24.9997
352  100000     1.63925    1.63934     25    24.9997
353  100000     1.63925    1.63934     25    24.9997
354  100000     1.63925    1.63934     25    24.9997
355  100000     1.63925    1.63934     25    24.9997
356  100000     1.63925    1.63934     25    24.9997
357  100000     1.63925    1.63934     25    24.9997
358  100000     1.63925    1.63934     25    24.9997
359  100000     1.63925    1.63934     25    24.9997
360  100000     1.63925    1.63934     25    24.9996
361  100000     1.63925    1.63934     25    24.9997
362  100000     1.63925    1.63934     25    24.9997
363  100000     1.63924    1.63934     25    24.9996
364  100000     1.63925    1.63934     25    24.9996
365  100000     1.63925    1.63934     25    24.9997
366  100000     1.63924    1.63934     25    24.9996
367  100000     1.63924    1.63934     25    24.9996
368  100000     1.63924    1.63934     25    24.9995
369  100000     1.63924    1.63934     25    24.9996
370  100000     1.63924    1.63934     25    24.9996
371  100000     1.63924    1.63934     25    24.9996
372  100000     1.63924    1.63934     25    24.9996
373  100000     1.63925    1.63934     25    24.9997
374  100000     1.63925    1.63934     25    24.9997
375  100000     1.63925    1.63934     25    24.9998
376  100000     1.63925    1.63934     25    24.9998
377  100000     1.63925    1.63934     25    24.9998
378  100000     1.63925    1.63934     25    24.9998
379  100000     1.63925    1.63934     25    24.9998
380  100000     1.63925    1.63934     25    24.9998
381  100000     1.63925    1.63934     25    24.9998
382  100000     1.63925    1.63934     25    24.9998
383  100000     1.63925    1.63934     25    24.9998
384  100000     1.63925    1.63934     25    24.9998
385  100000     1.63925    1.63934     25    24.9998
386  100000     1.63925    1.63934     25    24.9998
387  100000     1.63924    1.63934     25    24.9998
388  100000     1.63924    1.63934     25    24.9997
389  100000     1.63924    1.63934     25    24.9997
390  100000     1.63924    1.63934     25    24.9996
391  100000     1.63924    1.63934     25    24.9997
392  100000     1.63924    1.63934     25    24.9997
393  100000     1.63925    1.63934     25    24.9998
394  100000     1.63925    1.63934     25    24.9998
395  100000     1.63925    1.63934     25    24.9998
396  100000     1.63925    1.63934     25    24.9998
397  100000     1.63925    1.63934     25    24.9998
398  100000     1.63925    1.63934     25    24.9998
399  100000     1.63925    1.63934     25    24.9998
400  100000     1.63925    1.63934     25    24.9998

Asexual Model with selection (random number based on
relative fitness to determine who survives).  Mutation rate is      1e-08,
Recomb rate is ONE per asexual individual per generation
Starting values at generation   0 are; population size 100000,
with avg fitness of    1.22091.

gen  popSize    avgFit     maxFit maxAlleles avgAlleles
  0  100000     1.22091    1.70623     27     9.9900
  1  100000     1.22091    1.70623     27     9.9900
  2  100000     1.22545    1.70623     27    10.1761
  3  100000     1.23008    1.70623     27    10.3651
  4  100000     1.23487    1.70623     27    10.5598
  5  100000     1.24001    1.70623     27    10.7678
  6  100000     1.24488    1.70623     27    10.9651
  7  100000     1.25012    1.70623     27    11.1756
  8  100000     1.25520    1.70623     27    11.3788
  9  100000     1.26066    1.70623     27    11.5968
 10  100000     1.26619    1.70623     27    11.8163
 11  100000     1.27165    1.70623     27    12.0318
 12  100000     1.27689    1.70623     27    12.2373
 13  100000     1.28273    1.70623     27    12.4656
 14  100000     1.28878    1.70623     27    12.7020
 15  100000     1.29480    1.70623     27    12.9349
 16  100000     1.30145    1.70623     27    13.1921
 17  100000     1.30819    1.70623     27    13.4513
 18  100000     1.31479    1.70623     27    13.7040
 19  100000     1.32139    1.70623     27    13.9551
 20  100000     1.32768    1.70623     27    14.1939
 21  100000     1.33472    1.70623     27    14.4603
 22  100000     1.34158    1.70623     27    14.7180
 23  100000     1.34865    1.70623     27    14.9834
 24  100000     1.35618    1.70623     27    15.2642
 25  100000     1.36336    1.70623     27    15.5303
 26  100000     1.37043    1.70623     27    15.7915
 27  100000     1.37764    1.70623     27    16.0556
 28  100000     1.38499    1.70623     27    16.3239
 29  100000     1.39274    1.70623     27    16.6064
 30  100000     1.40026    1.70623     27    16.8784
 31  100000     1.40835    1.70623     27    17.1703
 32  100000     1.41601    1.70623     27    17.4436
 33  100000     1.42365    1.70623     27    17.7164
 34  100000     1.43129    1.70623     27    17.9875
 35  100000     1.43861    1.70623     27    18.2468
 36  100000     1.44607    1.70623     27    18.5087
 37  100000     1.45376    1.70623     27    18.7770
 38  100000     1.46125    1.70623     27    19.0366
 39  100000     1.46899    1.70623     27    19.3035
 40  100000     1.47677    1.70623     27    19.5709
 41  100000     1.48454    1.70623     27    19.8366
 42  100000     1.49200    1.70623     27    20.0899
 43  100000     1.49963    1.70623     27    20.3478
 44  100000     1.50764    1.70623     27    20.6180
 45  100000     1.51534    1.70623     27    20.8761
 46  100000     1.52309    1.70623     27    21.1342
 47  100000     1.53097    1.70623     27    21.3959
 48  100000     1.53830    1.70623     27    21.6380
 49  100000     1.54588    1.70623     27    21.8880
 50  100000     1.55340    1.70623     27    22.1339
 51  100000     1.56072    1.70623     27    22.3727
 52  100000     1.56857    1.70623     27    22.6289
 53  100000     1.57604    1.70623     27    22.8709
 54  100000     1.58370    1.70623     27    23.1185
 55  100000     1.59113    1.70623     27    23.3583
 56  100000     1.59799    1.70623     27    23.5800
 57  100000     1.60493    1.70623     27    23.8025
 58  100000     1.61153    1.70623     27    24.0139
 59  100000     1.61792    1.70623     27    24.2179
 60  100000     1.62395    1.70623     27    24.4102
 61  100000     1.62958    1.70623     27    24.5893
 62  100000     1.63528    1.70623     27    24.7704
 63  100000     1.64029    1.70623     27    24.9293
 64  100000     1.64503    1.70623     27    25.0797
 65  100000     1.64973    1.70623     27    25.2285
 66  100000     1.65450    1.70623     27    25.3788
 67  100000     1.65853    1.70623     27    25.5061
 68  100000     1.66256    1.70623     27    25.6330
 69  100000     1.66587    1.70623     27    25.7371
 70  100000     1.66921    1.70623     27    25.8426
 71  100000     1.67201    1.70623     27    25.9309
 72  100000     1.67477    1.70623     27    26.0176
 73  100000     1.67747    1.70623     27    26.1024
 74  100000     1.67973    1.70623     27    26.1734
 75  100000     1.68181    1.70623     27    26.2386
 76  100000     1.68399    1.70623     27    26.3070
 77  100000     1.68610    1.70623     27    26.3727
 78  100000     1.68764    1.70623     27    26.4209
 79  100000     1.68922    1.70623     27    26.4703
 80  100000     1.69066    1.70623     27    26.5156
 81  100000     1.69200    1.70623     27    26.5576
 82  100000     1.69298    1.70623     27    26.5881
 83  100000     1.69419    1.70623     27    26.6257
 84  100000     1.69520    1.70623     27    26.6573
 85  100000     1.69603    1.70623     27    26.6830
 86  100000     1.69685    1.70623     27    26.7088
 87  100000     1.69763    1.70623     27    26.7330
 88  100000     1.69826    1.70623     27    26.7525
 89  100000     1.69902    1.70623     27    26.7764
 90  100000     1.69969    1.70623     27    26.7972
 91  100000     1.70015    1.70623     27    26.8114
 92  100000     1.70068    1.70623     27    26.8279
 93  100000     1.70124    1.70623     27    26.8452
 94  100000     1.70161    1.70623     27    26.8569
 95  100000     1.70196    1.70623     27    26.8676
 96  100000     1.70223    1.70623     27    26.8761
 97  100000     1.70251    1.70623     27    26.8848
 98  100000     1.70277    1.70623     27    26.8927
 99  100000     1.70306    1.70623     27    26.9018
100  100000     1.70331    1.70623     27    26.9097
101  100000     1.70351    1.70623     27    26.9159
102  100000     1.70373    1.70623     27    26.9226
103  100000     1.70387    1.70623     27    26.9271
104  100000     1.70397    1.70623     27    26.9301
105  100000     1.70410    1.70623     27    26.9341
106  100000     1.70426    1.70623     27    26.9391
107  100000     1.70446    1.70623     27    26.9454
108  100000     1.70458    1.70623     27    26.9491
109  100000     1.70474    1.70623     27    26.9539
110  100000     1.70490    1.70623     27    26.9589
111  100000     1.70495    1.70623     27    26.9604
112  100000     1.70499    1.70623     27    26.9616
113  100000     1.70503    1.70623     27    26.9630
114  100000     1.70520    1.70623     27    26.9682
115  100000     1.70526    1.70623     27    26.9701
116  100000     1.70531    1.70623     27    26.9716
117  100000     1.70536    1.70623     27    26.9731
118  100000     1.70540    1.70623     27    26.9745
119  100000     1.70547    1.70623     27    26.9765
120  100000     1.70553    1.70623     27    26.9783
121  100000     1.70560    1.70623     27    26.9806
122  100000     1.70562    1.70623     27    26.9813
123  100000     1.70568    1.70623     27    26.9829
124  100000     1.70571    1.70623     27    26.9840
125  100000     1.70576    1.70623     27    26.9854
126  100000     1.70579    1.70623     27    26.9864
127  100000     1.70581    1.70623     27    26.9870
128  100000     1.70581    1.70623     27    26.9871
129  100000     1.70584    1.70623     27    26.9879
130  100000     1.70586    1.70623     27    26.9887
131  100000     1.70589    1.70623     27    26.9896
132  100000     1.70593    1.70623     27    26.9909
133  100000     1.70594    1.70623     27    26.9911
134  100000     1.70596    1.70623     27    26.9917
135  100000     1.70599    1.70623     27    26.9926
136  100000     1.70602    1.70623     27    26.9936
137  100000     1.70603    1.70623     27    26.9937
138  100000     1.70605    1.70623     27    26.9944
139  100000     1.70607    1.70623     27    26.9952
140  100000     1.70607    1.70623     27    26.9951
141  100000     1.70608    1.70623     27    26.9953
142  100000     1.70609    1.70623     27    26.9956
143  100000     1.70609    1.70623     27    26.9957
144  100000     1.70611    1.70623     27    26.9963
145  100000     1.70611    1.70623     27    26.9963
146  100000     1.70613    1.70623     27    26.9969
147  100000     1.70613    1.70623     27    26.9969
148  100000     1.70613    1.70623     27    26.9971
149  100000     1.70615    1.70623     27    26.9976
150  100000     1.70615    1.70623     27    26.9975
151  100000     1.70616    1.70623     27    26.9978
152  100000     1.70616    1.70623     27    26.9977
153  100000     1.70617    1.70623     27    26.9981
154  100000     1.70618    1.70623     27    26.9984
155  100000     1.70618    1.70623     27    26.9985
156  100000     1.70618    1.70623     27    26.9986
157  100000     1.70619    1.70623     27    26.9987
158  100000     1.70619    1.70623     27    26.9989
159  100000     1.70620    1.70623     27    26.9991
160  100000     1.70620    1.70623     27    26.9991
161  100000     1.70620    1.70623     27    26.9992
162  100000     1.70621    1.70623     27    26.9993
163  100000     1.70621    1.70623     27    26.9992
164  100000     1.70621    1.70623     27    26.9993
165  100000     1.70621    1.70623     27    26.9992
166  100000     1.70620    1.70623     27    26.9992
167  100000     1.70621    1.70623     27    26.9993
168  100000     1.70621    1.70623     27    26.9993
169  100000     1.70621    1.70623     27    26.9994
170  100000     1.70620    1.70623     27    26.9992
171  100000     1.70620    1.70623     27    26.9992
172  100000     1.70620    1.70623     27    26.9992
173  100000     1.70621    1.70623     27    26.9994
174  100000     1.70621    1.70623     27    26.9995
175  100000     1.70621    1.70623     27    26.9995
176  100000     1.70621    1.70623     27    26.9994
177  100000     1.70621    1.70623     27    26.9994
178  100000     1.70622    1.70623     27    26.9995
179  100000     1.70622    1.70623     27    26.9997
180  100000     1.70623    1.70623     27    26.9999
181  100000     1.70623    1.70623     27    26.9999
182  100000     1.70623    1.70623     27    26.9999
183  100000     1.70623    1.70623     27    26.9999
184  100000     1.70623    1.70623     27    27.0000
185  100000     1.70623    1.70623     27    27.0000
186  100000     1.70623    1.70623     27    27.0000
187  100000     1.70623    1.70623     27    26.9999
188  100000     1.70623    1.70623     27    27.0000
189  100000     1.70623    1.70623     27    27.0000
190  100000     1.70623    1.70623     27    27.0000
191  100000     1.70623    1.70623     27    27.0000
192  100000     1.70623    1.70623     27    27.0000
193  100000     1.70623    1.70623     27    27.0000
194  100000     1.70623    1.70623     27    27.0000
195  100000     1.70623    1.70623     27    27.0000
196  100000     1.70623    1.70623     27    27.0000
197  100000     1.70623    1.70623     27    27.0000
198  100000     1.70623    1.70623     27    27.0000
199  100000     1.70623    1.70623     27    27.0000
200  100000     1.70623    1.70623     27    27.0000
201  100000     1.70623    1.70623     27    27.0000
202  100000     1.70623    1.70623     27    27.0000
203  100000     1.70623    1.70623     27    27.0000
204  100000     1.70623    1.70623     27    27.0000
205  100000     1.70623    1.70623     27    27.0000
206  100000     1.70623    1.70623     27    27.0000
207  100000     1.70623    1.70623     27    27.0000
208  100000     1.70623    1.70623     27    27.0000
209  100000     1.70623    1.70623     27    27.0000
210  100000     1.70623    1.70623     27    27.0000
211  100000     1.70623    1.70623     27    27.0000
212  100000     1.70623    1.70623     27    27.0000
213  100000     1.70623    1.70623     27    27.0000
214  100000     1.70623    1.70623     27    27.0000
215  100000     1.70623    1.70623     27    27.0000
216  100000     1.70623    1.70623     27    27.0000
217  100000     1.70623    1.70623     27    27.0000
218  100000     1.70623    1.70623     27    27.0000
219  100000     1.70623    1.70623     27    27.0000
220  100000     1.70623    1.70623     27    27.0000
221  100000     1.70623    1.70623     27    27.0000
222  100000     1.70623    1.70623     27    27.0000
223  100000     1.70623    1.70623     27    27.0000
224  100000     1.70623    1.70623     27    27.0000
225  100000     1.70623    1.70623     27    27.0000
226  100000     1.70623    1.70623     27    27.0000
227  100000     1.70623    1.70623     27    27.0000
228  100000     1.70623    1.70623     27    27.0000
229  100000     1.70623    1.70623     27    27.0000
230  100000     1.70623    1.70623     27    27.0000
231  100000     1.70623    1.70623     27    27.0000
232  100000     1.70623    1.70623     27    27.0000
233  100000     1.70623    1.70623     27    27.0000
234  100000     1.70623    1.70623     27    27.0000
235  100000     1.70623    1.70623     27    27.0000
236  100000     1.70623    1.70623     27    27.0000
237  100000     1.70623    1.70623     27    27.0000
238  100000     1.70623    1.70623     27    27.0000
239  100000     1.70623    1.70623     27    27.0000
240  100000     1.70623    1.70623     27    27.0000
241  100000     1.70623    1.70623     27    27.0000
242  100000     1.70623    1.70623     27    27.0000
243  100000     1.70623    1.70623     27    27.0000
244  100000     1.70623    1.70623     27    27.0000
245  100000     1.70623    1.70623     27    27.0000
246  100000     1.70623    1.70623     27    27.0000
247  100000     1.70623    1.70623     27    27.0000
248  100000     1.70623    1.70623     27    27.0000
249  100000     1.70623    1.70623     27    27.0000
250  100000     1.70623    1.70623     27    27.0000
251  100000     1.70623    1.70623     27    27.0000
252  100000     1.70623    1.70623     27    27.0000
253  100000     1.70623    1.70623     27    27.0000
254  100000     1.70623    1.70623     27    27.0000
255  100000     1.70623    1.70623     27    27.0000
256  100000     1.70623    1.70623     27    27.0000
257  100000     1.70623    1.70623     27    27.0000
258  100000     1.70623    1.70623     27    27.0000
259  100000     1.70623    1.70623     27    27.0000
260  100000     1.70623    1.70623     27    27.0000
261  100000     1.70623    1.70623     27    27.0000
262  100000     1.70623    1.70623     27    27.0000
263  100000     1.70623    1.70623     27    27.0000
264  100000     1.70623    1.70623     27    27.0000
265  100000     1.70623    1.70623     27    27.0000
266  100000     1.70623    1.70623     27    27.0000
267  100000     1.70623    1.70623     27    27.0000
268  100000     1.70623    1.70623     27    27.0000
269  100000     1.70623    1.70623     27    27.0000
270  100000     1.70623    1.70623     27    27.0000
271  100000     1.70623    1.70623     27    27.0000
272  100000     1.70623    1.70623     27    27.0000
273  100000     1.70623    1.70623     27    27.0000
274  100000     1.70623    1.70623     27    27.0000
275  100000     1.70623    1.70623     27    27.0000
276  100000     1.70623    1.70623     27    27.0000
277  100000     1.70623    1.70623     27    27.0000
278  100000     1.70623    1.70623     27    27.0000
279  100000     1.70623    1.70623     27    27.0000
280  100000     1.70623    1.70623     27    27.0000
281  100000     1.70623    1.70623     27    27.0000
282  100000     1.70623    1.70623     27    27.0000
283  100000     1.70623    1.70623     27    27.0000
284  100000     1.70623    1.70623     27    27.0000
285  100000     1.70623    1.70623     27    27.0000
286  100000     1.70623    1.70623     27    27.0000
287  100000     1.70623    1.70623     27    27.0000
288  100000     1.70623    1.70623     27    27.0000
289  100000     1.70623    1.70623     27    27.0000
290  100000     1.70623    1.70623     27    27.0000
291  100000     1.70623    1.70623     27    27.0000
292  100000     1.70623    1.70623     27    27.0000
293  100000     1.70623    1.70623     27    27.0000
294  100000     1.70623    1.70623     27    27.0000
295  100000     1.70623    1.70623     27    27.0000
296  100000     1.70623    1.70623     27    27.0000
297  100000     1.70623    1.70623     27    27.0000
298  100000     1.70623    1.70623     27    27.0000
299  100000     1.70623    1.70623     27    27.0000
300  100000     1.70623    1.70623     27    27.0000
301  100000     1.70623    1.70623     27    27.0000
302  100000     1.70623    1.70623     27    27.0000
303  100000     1.70623    1.70623     27    27.0000
304  100000     1.70623    1.70623     27    27.0000
305  100000     1.70623    1.70623     27    27.0000
306  100000     1.70623    1.70623     27    27.0000
307  100000     1.70623    1.70623     27    27.0000
308  100000     1.70623    1.70623     27    27.0000
309  100000     1.70623    1.70623     27    27.0000
310  100000     1.70623    1.70623     27    27.0000
311  100000     1.70623    1.70623     27    27.0000
312  100000     1.70623    1.70623     27    27.0000
313  100000     1.70623    1.70623     27    27.0000
314  100000     1.70623    1.70623     27    27.0000
315  100000     1.70623    1.70623     27    26.9999
316  100000     1.70623    1.70623     27    26.9999
317  100000     1.70623    1.70623     27    27.0000
318  100000     1.70623    1.70623     27    27.0000
319  100000     1.70623    1.70623     27    27.0000
320  100000     1.70623    1.70623     27    27.0000
321  100000     1.70623    1.70623     27    27.0000
322  100000     1.70623    1.70623     27    27.0000
323  100000     1.70623    1.70623     27    27.0000
324  100000     1.70623    1.70623     27    27.0000
325  100000     1.70623    1.70623     27    27.0000
326  100000     1.70623    1.70623     27    27.0000
327  100000     1.70623    1.70623     27    27.0000
328  100000     1.70623    1.70623     27    27.0000
329  100000     1.70623    1.70623     27    27.0000
330  100000     1.70623    1.70623     27    27.0000
331  100000     1.70623    1.70623     27    27.0000
332  100000     1.70623    1.70623     27    27.0000
333  100000     1.70623    1.70623     27    27.0000
334  100000     1.70623    1.70623     27    27.0000
335  100000     1.70623    1.70623     27    27.0000
336  100000     1.70623    1.70623     27    27.0000
337  100000     1.70623    1.70623     27    27.0000
338  100000     1.70623    1.70623     27    27.0000
339  100000     1.70623    1.70623     27    27.0000
340  100000     1.70623    1.70623     27    27.0000
341  100000     1.70623    1.70623     27    27.0000
342  100000     1.70623    1.70623     27    27.0000
343  100000     1.70623    1.70623     27    27.0000
344  100000     1.70623    1.70623     27    27.0000
345  100000     1.70623    1.70623     27    27.0000
346  100000     1.70623    1.70623     27    27.0000
347  100000     1.70623    1.70623     27    27.0000
348  100000     1.70623    1.70623     27    27.0000
349  100000     1.70623    1.70623     27    27.0000
350  100000     1.70623    1.70623     27    27.0000
351  100000     1.70623    1.70623     27    27.0000
352  100000     1.70623    1.70623     27    27.0000
353  100000     1.70623    1.70623     27    27.0000
354  100000     1.70623    1.70623     27    27.0000
355  100000     1.70623    1.70623     27    27.0000
356  100000     1.70623    1.70623     27    27.0000
357  100000     1.70623    1.70623     27    27.0000
358  100000     1.70623    1.70623     27    27.0000
359  100000     1.70623    1.70623     27    27.0000
360  100000     1.70623    1.70623     27    27.0000
361  100000     1.70623    1.70623     27    27.0000
362  100000     1.70623    1.70623     27    27.0000
363  100000     1.70623    1.70623     27    27.0000
364  100000     1.70623    1.70623     27    27.0000
365  100000     1.70623    1.70623     27    27.0000
366  100000     1.70623    1.70623     27    27.0000
367  100000     1.70623    1.70623     27    27.0000
368  100000     1.70623    1.70623     27    27.0000
369  100000     1.70623    1.70623     27    27.0000
370  100000     1.70623    1.70623     27    27.0000
371  100000     1.70623    1.70623     27    27.0000
372  100000     1.70623    1.70623     27    27.0000
373  100000     1.70623    1.70623     27    27.0000
374  100000     1.70623    1.70623     27    27.0000
375  100000     1.70623    1.70623     27    27.0000
376  100000     1.70623    1.70623     27    27.0000
377  100000     1.70623    1.70623     27    27.0000
378  100000     1.70623    1.70623     27    27.0000
379  100000     1.70623    1.70623     27    27.0000
380  100000     1.70623    1.70623     27    27.0000
381  100000     1.70623    1.70623     27    27.0000
382  100000     1.70623    1.70623     27    27.0000
383  100000     1.70623    1.70623     27    27.0000
384  100000     1.70623    1.70623     27    27.0000
385  100000     1.70623    1.70623     27    27.0000
386  100000     1.70623    1.70623     27    27.0000
387  100000     1.70623    1.70623     27    27.0000
388  100000     1.70623    1.70623     27    27.0000
389  100000     1.70623    1.70623     27    27.0000
390  100000     1.70623    1.70623     27    27.0000
391  100000     1.70623    1.70623     27    27.0000
392  100000     1.70623    1.70623     27    27.0000
393  100000     1.70623    1.70623     27    27.0000
394  100000     1.70623    1.70623     27    27.0000
395  100000     1.70623    1.70623     27    27.0000
396  100000     1.70623    1.70623     27    27.0000
397  100000     1.70623    1.70623     27    27.0000
398  100000     1.70623    1.70623     27    27.0000
399  100000     1.70623    1.70623     27    27.0000
400  100000     1.70623    1.70623     27    27.0000
```
