## Supplementary Table 2 for "Sex Solves Haldane’s Dilemma"

```
Supplementary Table S2. Simulation results for ten replicates of a sexual population.
	Recombination rate is ONE per sexual individual per generation
Starting values are: population size 100000, number of loci 100, allele frequencies 0.05, selective advantage 0.02 with multiplicative effects between loci.


gen  popSize    avgFit     maxFit maxAlleles avgAlleles
  0  100000     1.22116    1.70689     27    10.0006
  1  100000     1.22116    1.70689     27    10.0006
  2  100000     1.22588    1.67213     26    10.1937
  3  100000     1.23046    1.70689     27    10.3803
  4  100000     1.23507    1.80927     30    10.5682
  5  100000     1.23969    1.70557     27    10.7557
  6  100000     1.24482    1.67277     26    10.9622
  7  100000     1.24986    1.70623     27    11.1647
  8  100000     1.25532    1.70426     27    11.3836
  9  100000     1.26055    1.70689     27    11.5917
 10  100000     1.26642    1.70689     27    11.8246
 11  100000     1.27205    1.77312     29    12.0474
 12  100000     1.27776    1.77584     29    12.2713
 13  100000     1.28319    1.80927     30    12.4838
 14  100000     1.28937    1.77312     29    12.7253
 15  100000     1.29538    1.84759     31    12.9568
 16  100000     1.30144    1.88309     32    13.1908
 17  100000     1.30778    1.99682     35    13.4340
 18  100000     1.31485    1.88454     32    13.7043
 19  100000     1.32104    1.99758     35    13.9399
 20  100000     1.32773    1.88382     32    14.1924
 21  100000     1.33539    1.95842     34    14.4807
 22  100000     1.34213    1.91854     33    14.7330
 23  100000     1.34984    1.88382     32    15.0201
 24  100000     1.35695    1.95917     34    15.2844
 25  100000     1.36489    1.88382     32    15.5786
 26  100000     1.37275    2.03754     36    15.8664
 27  100000     1.38100    2.03675     36    16.1672
 28  100000     1.38884    1.99758     35    16.4513
 29  100000     1.39738    1.99912     35    16.7573
 30  100000     1.40628    1.99912     35    17.0768
 31  100000     1.41540    2.16059     39    17.4021
 32  100000     1.42459    2.03675     36    17.7269
 33  100000     1.43328    2.16059     39    18.0317
 34  100000     1.44199    2.07829     37    18.3339
 35  100000     1.45145    2.07989     37    18.6635
 36  100000     1.46146    2.16059     39    19.0086
 37  100000     1.47213    2.20549     40    19.3738
 38  100000     1.48177    2.16142     39    19.7004
 39  100000     1.49264    2.20465     40    20.0686
 40  100000     1.50370    2.43317     45    20.4388
 41  100000     1.51514    2.38455     44    20.8195
 42  100000     1.52638    2.33869     43    21.1917
 43  100000     1.53760    2.38455     44    21.5604
 44  100000     1.54922    2.43224     45    21.9377
 45  100000     1.56103    2.34049     43    22.3194
 46  100000     1.57238    2.38638     44    22.6822
 47  100000     1.58478    2.53147     47    23.0781
 48  100000     1.59732    2.52855     47    23.4738
 49  100000     1.60962    2.52953     47    23.8564
 50  100000     1.62345    2.43411     45    24.2878
 51  100000     1.63714    2.48374     46    24.7105
 52  100000     1.65086    2.48088     46    25.1304
 53  100000     1.66437    2.63172     49    25.5393
 54  100000     1.67948    2.63577     49    25.9942
 55  100000     1.69424    2.63273     49    26.4333
 56  100000     1.70984    2.58309     48    26.8939
 57  100000     1.72527    2.68745     50    27.3454
 58  100000     1.74132    2.63374     49    27.8125
 59  100000     1.75831    2.79495     52    28.3017
 60  100000     1.77444    2.84866     53    28.7601
 61  100000     1.79162    2.96374     55    29.2445
 62  100000     1.80917    3.02302     56    29.7323
 63  100000     1.82685    2.90787     54    30.2221
 64  100000     1.84598    2.96146     55    30.7465
 65  100000     1.86354    2.90898     54    31.2232
 66  100000     1.88219    2.96602     55    31.7229
 67  100000     1.90143    3.46983     63    32.2325
 68  100000     1.92340    2.96602     55    32.8133
 69  100000     1.94432    3.20558     59    33.3575
 70  100000     1.96541    3.26467     60    33.8988
 71  100000     1.98826    3.20928     59    34.4817
 72  100000     2.01099    3.40310     62    35.0571
 73  100000     2.03334    3.33766     61    35.6109
 74  100000     2.05631    3.26718     60    36.1773
 75  100000     2.07891    3.34151     61    36.7280
 76  100000     2.10358    3.46849     63    37.3209
 77  100000     2.12906    3.47116     63    37.9266
 78  100000     2.15356    3.82361     68    38.5041
 79  100000     2.18094    3.75585     67    39.1407
 80  100000     2.20722    3.68079     66    39.7478
 81  100000     2.23428    3.82950     68    40.3608
 82  100000     2.26181    4.05453     71    40.9776
 83  100000     2.29167    3.82361     68    41.6351
 84  100000     2.32377    3.75296     67    42.3386
 85  100000     2.35418    3.98727     70    42.9969
 86  100000     2.38473    3.75441     67    43.6482
 87  100000     2.41841    4.06545     71    44.3575
 88  100000     2.45008    4.13244     72    45.0138
 89  100000     2.48436    4.22645     73    45.7142
 90  100000     2.51744    4.39044     75    46.3781
 91  100000     2.55424    4.66096     78    47.1123
 92  100000     2.58895    4.65738     78    47.7932
 93  100000     2.62554    4.47652     76    48.5045
 94  100000     2.66280    4.65738     78    49.2134
 95  100000     2.70013    4.84181     80    49.9178
 96  100000     2.73917    4.84926     80    50.6403
 97  100000     2.78152    4.56430     77    51.4178
 98  100000     2.82275    4.75418     79    52.1636
 99  100000     2.86593    4.84926     80    52.9333
100  100000     2.90847    4.94435     81    53.6770
101  100000     2.95558    5.04711     82    54.4876
102  100000     3.00234    5.23489     84    55.2846
103  100000     3.04756    5.23489     84    56.0392
104  100000     3.09506    5.24698     84    56.8242
105  100000     3.14581    5.35192     85    57.6459
106  100000     3.19462    5.44847     86    58.4197
107  100000     3.24472    5.67077     88    59.2093
108  100000     3.29918    5.55744     87    60.0526
109  100000     3.35218    5.77752     89    60.8618
110  100000     3.40493    6.00631     91    61.6546
111  100000     3.45774    5.89080     90    62.4286
112  100000     3.51585    6.25137     93    63.2765
113  100000     3.57478    6.12643     92    64.1148
114  100000     3.63287    6.25858     93    64.9281
115  100000     3.69737    6.25617     93    65.8228
116  100000     3.75997    6.64166     96    66.6768
117  100000     3.82359    6.37639     94    67.5276
118  100000     3.88767    6.63400     96    68.3679
119  100000     3.95574    6.89671     98    69.2474
120  100000     4.02140    6.76668     97    70.0817
121  100000     4.09062    7.92519    105    70.9435
122  100000     4.16413    7.45088    102    71.8548
123  100000     4.23519    8.08059    106    72.7100
124  100000     4.31261    7.45948    102    73.6305
125  100000     4.38730    7.59989    103    74.5022
126  100000     4.46565    7.76084    104    75.4009
127  100000     4.54354    8.07127    106    76.2789
128  100000     4.62382    8.39735    108    77.1670
129  100000     4.70366    8.88397    111    78.0379
130  100000     4.78836    8.23587    107    78.9456
131  100000     4.87793    9.07909    112    79.8852
132  100000     4.96419    8.39412    108    80.7828
133  100000     5.05335    8.89080    111    81.6849
134  100000     5.14255    9.08607    112    82.5764
135  100000     5.23540    9.43136    114    83.4826
136  100000     5.33084   10.02404    117    84.4047
137  100000     5.42513    9.62369    115    85.2954
138  100000     5.52481    9.81239    116    86.2213
139  100000     5.62647   10.00479    117    87.1459
140  100000     5.73062   10.39698    119    88.0801
141  100000     5.83814   10.03176    117    89.0294
142  100000     5.95055   10.41699    119    89.9975
143  100000     6.05503   10.41699    119    90.8864
144  100000     6.16706   11.05034    122    91.8218
145  100000     6.28009   10.82534    121    92.7461
146  100000     6.38928   11.71319    125    93.6259
147  100000     6.50884   11.27135    123    94.5702
148  100000     6.62640   12.17704    127    95.4852
149  100000     6.74616   11.93827    126    96.3937
150  100000     6.86767   11.73122    125    97.3028
151  100000     6.99465   12.17236    127    98.2373
152  100000     7.12874   12.41103    128    99.2052
153  100000     7.26191   13.70278    133   100.1525
154  100000     7.39898   12.92734    130   101.1085
155  100000     7.53269   12.92734    130   102.0214
156  100000     7.67533   13.15550    131   102.9819
157  100000     7.82317   13.97146    134   103.9576
158  100000     7.95831   13.67646    133   104.8250
159  100000     8.10465   13.43926    132   105.7558
160  100000     8.25055   14.24541    135   106.6654
161  100000     8.40396   14.80953    137   107.6081
162  100000     8.55505   15.11734    138   108.5211
163  100000     8.71198   16.02415    141   109.4423
164  100000     8.86444   15.68581    140   110.3275
165  100000     9.02451   15.09992    138   111.2439
166  100000     9.18803   16.31951    142   112.1661
167  100000     9.35398   16.01183    141   113.0819
168  100000     9.51764   16.35092    142   113.9689
169  100000     9.69132   16.31951    142   114.8954
170  100000     9.86891   17.31174    145   115.8244
171  100000    10.03744   17.32506    145   116.6881
172  100000    10.21792   18.00421    147   117.6010
173  100000    10.39446   17.32506    145   118.4743
174  100000    10.58141   17.33839    145   119.3852
175  100000    10.76384   19.06951    150   120.2630
176  100000    10.95530   19.85518    152   121.1723
177  100000    11.13932   18.72438    149   122.0279
178  100000    11.32796   18.37136    148   122.8875
179  100000    11.51617   20.21338    153   123.7315
180  100000    11.70760   21.87120    157   124.5749
181  100000    11.90207   21.03809    155   125.4174
182  100000    12.09962   20.64939    154   126.2619
183  100000    12.29512   20.64939    154   127.0883
184  100000    12.49610   21.05428    155   127.9197
185  100000    12.70109   21.03809    155   128.7478
186  100000    12.91326   22.72857    159   129.5991
187  100000    13.12947   21.45060    156   130.4454
188  100000    13.34704   22.30863    158   131.2909
189  100000    13.55857   23.18314    160   132.0979
190  100000    13.77892   23.17422    160   132.9247
191  100000    13.99777   24.09193    162   133.7330
192  100000    14.22915   23.64680    161   134.5757
193  100000    14.45089   23.62862    161   135.3645
194  100000    14.68949   23.63771    161   136.2029
195  100000    14.93068   25.56655    165   137.0429
196  100000    15.17292   26.06785    166   137.8659
197  100000    15.41790   24.59267    163   138.6872
198  100000    15.65663   27.61027    169   139.4840
199  100000    15.90085   26.56877    167   140.2795
200  100000    16.13962   28.18415    170   141.0464
201  100000    16.38380   27.08972    168   141.8159
202  100000    16.63669   26.58921    167   142.6007
203  100000    16.88349   28.20583    170   143.3585
204  100000    17.13525   28.18415    170   144.1213
205  100000    17.38952   27.11056    168   144.8777
206  100000    17.64927   28.18415    170   145.6381
207  100000    17.90967   28.18415    170   146.3958
208  100000    18.15183   29.87476    173   147.0852
209  100000    18.39643   28.73678    171   147.7740
210  100000    18.63963   30.47225    174   148.4463
211  100000    18.89173   29.88625    173   149.1396
212  100000    19.15010   29.30024    172   149.8405
213  100000    19.42437   31.05780    175   150.5698
214  100000    19.68766   32.93345    178   151.2639
215  100000    19.95966   33.57920    179   151.9686
216  100000    20.22187   32.90813    178   152.6401
217  100000    20.50149   32.30012    177   153.3459
218  100000    20.78292   31.69114    176   154.0504
219  100000    21.03735   32.31254    177   154.6767
220  100000    21.28260   32.28770    177   155.2771
221  100000    21.55237   32.31254    177   155.9259
222  100000    21.83955   34.25079    180   156.6107
223  100000    22.11545   34.92237    181   157.2551
224  100000    22.38176   34.23762    180   157.8746
225  100000    22.65160   36.29135    183   158.4914
226  100000    22.93006   36.30530    183   159.1183
227  100000    23.19673   34.90895    181   159.7148
228  100000    23.46549   36.30530    183   160.3090
229  100000    23.74693   34.90895    181   160.9216
230  100000    24.00684   38.46826    186   161.4821
231  100000    24.30659   35.60713    182   162.1227
232  100000    24.57032   38.48306    186   162.6807
233  100000    24.83214   37.01717    184   163.2278
234  100000    25.09169   37.01717    184   163.7661
235  100000    25.34399   39.23763    187   164.2810
236  100000    25.60984   37.74300    185   164.8215
237  100000    25.87767   37.72849    185   165.3569
238  100000    26.13739   38.48306    186   165.8746
239  100000    26.40396   40.77576    189   166.3973
240  100000    26.66153   38.48306    186   166.8966
241  100000    26.94258   39.99161    188   167.4375
242  100000    27.22278   39.23763    187   167.9731
243  100000    27.48226   40.00699    188   168.4627
244  100000    27.74224   41.59128    190   168.9479
245  100000    27.99663   40.79144    189   169.4192
246  100000    28.25631   42.40679    191   169.8977
247  100000    28.52229   40.77576    189   170.3798
248  100000    28.80276   40.00699    188   170.8847
249  100000    29.07516   40.79144    189   171.3729
250  100000    29.33991   40.79144    189   171.8424
251  100000    29.60282   42.40679    191   172.3045
252  100000    29.86256   43.23830    192   172.7576
253  100000    30.10700   43.23830    192   173.1784
254  100000    30.35879   43.23830    192   173.6081
255  100000    30.62045   42.40679    191   174.0518
256  100000    30.86820   43.23830    192   174.4686
257  100000    31.12822   44.08611    193   174.9024
258  100000    31.38619   44.08611    193   175.3286
259  100000    31.63441   44.95054    194   175.7377
260  100000    31.88560   44.08611    193   176.1474
261  100000    32.12609   44.06916    193   176.5362
262  100000    32.37821   44.08611    193   176.9400
263  100000    32.61645   43.23830    192   177.3186
264  100000    32.87834   44.08611    193   177.7330
265  100000    33.12598   46.73059    196   178.1220
266  100000    33.36069   45.83192    195   178.4866
267  100000    33.60701   46.73059    196   178.8678
268  100000    33.84817   46.71262    196   179.2375
269  100000    34.07497   45.83192    195   179.5833
270  100000    34.29923   46.73059    196   179.9237
271  100000    34.50872   46.73059    196   180.2391
272  100000    34.72149   46.73059    196   180.5549
273  100000    34.95278   46.73059    196   180.8990
274  100000    35.17545   45.83192    195   181.2293
275  100000    35.39655   45.83192    195   181.5515
276  100000    35.62420   45.83192    195   181.8846
277  100000    35.85475   47.64688    197   182.2194
278  100000    36.07242   46.73059    196   182.5348
279  100000    36.28450   46.73059    196   182.8370
280  100000    36.49334   47.64688    197   183.1344
281  100000    36.69446   47.64688    197   183.4190
282  100000    36.88920   47.64688    197   183.6941
283  100000    37.08703   49.53370    199   183.9698
284  100000    37.29756   47.64688    197   184.2629
285  100000    37.49465   48.58113    198   184.5363
286  100000    37.67688   48.58113    198   184.7885
287  100000    37.86320   48.58113    198   185.0436
288  100000    38.04909   48.58113    198   185.2981
289  100000    38.24018   47.64688    197   185.5565
290  100000    38.41884   49.53370    199   185.7987
291  100000    38.61102   49.53370    199   186.0579
292  100000    38.79171   49.53370    199   186.2995
293  100000    38.97189   49.53370    199   186.5395
294  100000    39.15288   49.53370    199   186.7788
295  100000    39.33128   49.53370    199   187.0141
296  100000    39.51052   49.53370    199   187.2512
297  100000    39.70387   50.50495    200   187.5051
298  100000    39.88874   49.53370    199   187.7454
299  100000    40.05964   50.50495    200   187.9659
300  100000    40.21750   49.53370    199   188.1696
301  100000    40.38060   50.50495    200   188.3796
302  100000    40.55578   49.53370    199   188.6039
303  100000    40.71360   49.53370    199   188.8060
304  100000    40.86617   50.50495    200   188.9999
305  100000    41.01862   49.53370    199   189.1935
306  100000    41.15851   50.50495    200   189.3702
307  100000    41.30616   50.50495    200   189.5555
308  100000    41.46454   50.50495    200   189.7543
309  100000    41.62589   50.50495    200   189.9557
310  100000    41.75823   50.50495    200   190.1201
311  100000    41.89265   50.50495    200   190.2871
312  100000    42.03085   50.50495    200   190.4583
313  100000    42.17214   50.50495    200   190.6331
314  100000    42.31659   50.50495    200   190.8104
315  100000    42.45890   50.50495    200   190.9839
316  100000    42.58814   50.50495    200   191.1423
317  100000    42.72600   50.50495    200   191.3103
318  100000    42.87142   50.50495    200   191.4864
319  100000    42.99800   50.50495    200   191.6397
320  100000    43.12816   50.50495    200   191.7963
321  100000    43.24773   50.50495    200   191.9405
322  100000    43.36670   50.50495    200   192.0829
323  100000    43.49805   50.50495    200   192.2393
324  100000    43.61822   50.50495    200   192.3834
325  100000    43.73680   50.50495    200   192.5246
326  100000    43.84787   50.50495    200   192.6567
327  100000    43.95445   50.50495    200   192.7828
328  100000    44.06032   50.50495    200   192.9069
329  100000    44.17297   50.50495    200   193.0399
330  100000    44.28371   50.50495    200   193.1696
331  100000    44.39674   50.50495    200   193.3014
332  100000    44.49281   50.50495    200   193.4141
333  100000    44.59304   50.50495    200   193.5303
334  100000    44.70963   50.50495    200   193.6659
335  100000    44.81008   50.50495    200   193.7827
336  100000    44.92945   50.50495    200   193.9208
337  100000    45.02062   50.50495    200   194.0268
338  100000    45.11660   50.50495    200   194.1374
339  100000    45.20110   50.50495    200   194.2345
340  100000    45.29368   50.50495    200   194.3407
341  100000    45.38990   50.50495    200   194.4508
342  100000    45.47020   50.50495    200   194.5424
343  100000    45.55341   50.50495    200   194.6374
344  100000    45.64415   50.50495    200   194.7404
345  100000    45.72620   50.50495    200   194.8337
346  100000    45.81683   50.50495    200   194.9367
347  100000    45.90324   50.50495    200   195.0346
348  100000    45.98986   50.50495    200   195.1328
349  100000    46.06455   50.50495    200   195.2173
350  100000    46.13602   50.50495    200   195.2976
351  100000    46.20868   50.50495    200   195.3791
352  100000    46.27820   50.50495    200   195.4576
353  100000    46.35928   50.50495    200   195.5482
354  100000    46.42966   50.50495    200   195.6271
355  100000    46.49097   50.50495    200   195.6962
356  100000    46.56793   50.50495    200   195.7821
357  100000    46.62374   50.50495    200   195.8441
358  100000    46.69369   50.50495    200   195.9222
359  100000    46.76228   50.50495    200   195.9982
360  100000    46.83789   50.50495    200   196.0820
361  100000    46.90710   50.50495    200   196.1586
362  100000    46.98283   50.50495    200   196.2424
363  100000    47.04227   50.50495    200   196.3079
364  100000    47.10323   50.50495    200   196.3754
365  100000    47.16743   50.50495    200   196.4461
366  100000    47.22202   50.50495    200   196.5065
367  100000    47.27950   50.50495    200   196.5698
368  100000    47.33127   50.50495    200   196.6268
369  100000    47.39160   50.50495    200   196.6927
370  100000    47.45238   50.50495    200   196.7595
371  100000    47.50574   50.50495    200   196.8179
372  100000    47.56387   50.50495    200   196.8814
373  100000    47.61747   50.50495    200   196.9400
374  100000    47.66687   50.50495    200   196.9938
375  100000    47.71471   50.50495    200   197.0458
376  100000    47.76666   50.50495    200   197.1026
377  100000    47.81721   50.50495    200   197.1575
378  100000    47.86628   50.50495    200   197.2103
379  100000    47.92056   50.50495    200   197.2695
380  100000    47.97522   50.50495    200   197.3288
381  100000    48.02904   50.50495    200   197.3872
382  100000    48.07808   50.50495    200   197.4400
383  100000    48.11729   50.50495    200   197.4825
384  100000    48.16128   50.50495    200   197.5301
385  100000    48.20563   50.50495    200   197.5779
386  100000    48.24773   50.50495    200   197.6231
387  100000    48.29088   50.50495    200   197.6696
388  100000    48.33092   50.50495    200   197.7126
389  100000    48.36876   50.50495    200   197.7535
390  100000    48.40917   50.50495    200   197.7967
391  100000    48.44411   50.50495    200   197.8341
392  100000    48.48347   50.50495    200   197.8761
393  100000    48.52459   50.50495    200   197.9200
394  100000    48.56761   50.50495    200   197.9663
395  100000    48.60512   50.50495    200   198.0067
396  100000    48.63897   50.50495    200   198.0428
397  100000    48.67189   50.50495    200   198.0780
398  100000    48.69695   50.50495    200   198.1047
399  100000    48.73266   50.50495    200   198.1428
400  100000    48.76278   50.50495    200   198.1749

Diploid Model with selection (random number based on
relative fitness to determine who survives).  Mutation rate is      1e-08,
Recomb rate is ONE per sexual individual per generation
Starting values at generation   0 are; population size 100000,
with avg fitness of    1.22144.

gen  popSize    avgFit     maxFit maxAlleles avgAlleles
  0  100000     1.22144    1.74102     28    10.0116
  1  100000     1.22144    1.74102     28    10.0116
  2  100000     1.22598    1.70623     27    10.1963
  3  100000     1.23083    1.67213     26    10.3944
  4  100000     1.23573    1.70623     27    10.5935
  5  100000     1.24060    1.67149     26    10.7911
  6  100000     1.24523    1.74035     28    10.9777
  7  100000     1.25043    1.74035     28    11.1864
  8  100000     1.25586    1.70689     27    11.4031
  9  100000     1.26130    1.70689     27    11.6198
 10  100000     1.26674    1.70689     27    11.8360
 11  100000     1.27237    1.74035     28    12.0592
 12  100000     1.27824    1.84404     31    12.2892
 13  100000     1.28391    1.80927     30    12.5115
 14  100000     1.29001    1.84546     31    12.7485
 15  100000     1.29584    1.81067     30    12.9743
 16  100000     1.30285    1.91928     33    13.2444
 17  100000     1.30944    1.92002     33    13.4977
 18  100000     1.31593    1.84546     31    13.7456
 19  100000     1.32305    1.88309     32    14.0162
 20  100000     1.32955    1.84546     31    14.2624
 21  100000     1.33651    1.92149     33    14.5246
 22  100000     1.34339    1.99835     35    14.7811
 23  100000     1.35036    1.99682     35    15.0408
 24  100000     1.35804    2.03832     36    15.3250
 25  100000     1.36527    1.95992     34    15.5910
 26  100000     1.37323    1.95842     34    15.8839
 27  100000     1.38121    2.07829     37    16.1752
 28  100000     1.38944    1.99835     35    16.4720
 29  100000     1.39740    2.03675     36    16.7591
 30  100000     1.40596    2.03832     36    17.0655
 31  100000     1.41449    2.11904     38    17.3704
 32  100000     1.42366    2.16225     39    17.6942
 33  100000     1.43248    2.20465     40    18.0043
 34  100000     1.44148    2.07909     37    18.3190
 35  100000     1.44985    2.07749     37    18.6090
 36  100000     1.45945    2.16225     39    18.9412
 37  100000     1.46868    2.34049     43    19.2575
 38  100000     1.47846    2.20549     40    19.5912
 39  100000     1.48865    2.20549     40    19.9362
 40  100000     1.49970    2.24960     41    20.3061
 41  100000     1.51110    2.20380     40    20.6863
 42  100000     1.52141    2.24528     41    21.0262
 43  100000     1.53359    2.24874     41    21.4278
 44  100000     1.54435    2.25047     41    21.7788
 45  100000     1.55613    2.29460     42    22.1611
 46  100000     1.56871    2.33959     43    22.5676
 47  100000     1.58077    2.57912     48    22.9510
 48  100000     1.59259    2.48088     46    23.3261
 49  100000     1.60535    2.38638     44    23.7277
 50  100000     1.61868    2.43224     45    24.1429
 51  100000     1.63237    2.53050     47    24.5654
 52  100000     1.64573    2.43411     45    24.9747
 53  100000     1.65912    2.58111     48    25.3836
 54  100000     1.67386    2.48183     46    25.8266
 55  100000     1.68860    2.63476     49    26.2662
 56  100000     1.70387    2.63273     49    26.7160
 57  100000     1.72014    2.73804     51    27.1947
 58  100000     1.73652    2.73909     51    27.6699
 59  100000     1.75236    2.89893     54    28.1280
 60  100000     1.76899    2.90451     54    28.6017
 61  100000     1.78723    2.79280     52    29.1171
 62  100000     1.80532    2.85085     53    29.6263
 63  100000     1.82382    2.84428     53    30.1399
 64  100000     1.84231    2.96602     55    30.6470
 65  100000     1.86197    3.07992     57    31.1810
 66  100000     1.88156    3.02534     56    31.7081
 67  100000     1.90113    3.14515     58    32.2306
 68  100000     1.92212    2.96374     55    32.7825
 69  100000     1.94284    3.14636     58    33.3228
 70  100000     1.96408    3.33253     61    33.8708
 71  100000     1.98543    3.20928     59    34.4162
 72  100000     2.00525    3.33253     61    34.9178
 73  100000     2.02814    3.20189     59    35.4884
 74  100000     2.05028    3.46716     63    36.0329
 75  100000     2.07389    3.60862     65    36.6105
 76  100000     2.09760    3.67938     66    37.1845
 77  100000     2.12295    3.47383     63    37.7860
 78  100000     2.14763    3.46583     63    38.3683
 79  100000     2.17282    3.67655     66    38.9502
 80  100000     2.20026    3.60307     65    39.5835
 81  100000     2.22683    3.90008     69    40.1903
 82  100000     2.25277    3.60862     65    40.7756
 83  100000     2.28177    3.75585     67    41.4197
 84  100000     2.31084    3.67938     66    42.0594
 85  100000     2.34046    3.90759     69    42.7020
 86  100000     2.37159    3.90158     69    43.3680
 87  100000     2.40187    4.22320     73    44.0093
 88  100000     2.43330    3.90609     69    44.6660
 89  100000     2.46775    3.98574     70    45.3763
 90  100000     2.50089    4.14198     72    46.0465
 91  100000     2.53588    4.30601     74    46.7507
 92  100000     2.57174    4.22320     73    47.4557
 93  100000     2.60812    4.47997     76    48.1641
 94  100000     2.64526    4.84553     80    48.8769
 95  100000     2.68207    4.65201     78    49.5760
 96  100000     2.72127    4.64843     78    50.3135
 97  100000     2.75948    4.66455     78    51.0218
 98  100000     2.80148    5.04323     82    51.7834
 99  100000     2.84396    4.75235     79    52.5415
100  100000     2.88788    5.34369     85    53.3179
101  100000     2.93339    5.34369     85    54.1047
102  100000     2.97879    5.23489     84    54.8869
103  100000     3.02253    5.04129     82    55.6200
104  100000     3.07019    5.23891     84    56.4103
105  100000     3.11850    5.34164     85    57.1984
106  100000     3.16870    6.00631     91    58.0084
107  100000     3.21830    5.45057     86    58.7939
108  100000     3.26770    5.55530     87    59.5609
109  100000     3.31951    5.77086     89    60.3616
110  100000     3.37359    5.77752     89    61.1774
111  100000     3.42743    6.37885     94    61.9794
112  100000     3.48083    6.02018     91    62.7624
113  100000     3.54038    6.26822     93    63.6156
114  100000     3.59967    6.38375     94    64.4619
115  100000     3.65935    6.62380     96    65.2946
116  100000     3.72303    6.63910     96    66.1673
117  100000     3.78648    6.49892     95    67.0263
118  100000     3.85054    6.64166     96    67.8769
119  100000     3.91445    6.50392     95    68.7120
120  100000     3.98234    7.17809    100    69.5890
121  100000     4.05277    7.03194     99    70.4797
122  100000     4.12497    7.04005     99    71.3730
123  100000     4.19607    7.45661    102    72.2419
124  100000     4.27256    7.33011    101    73.1586
125  100000     4.34478    8.23587    107    74.0036
126  100000     4.42178    8.07748    106    74.8967
127  100000     4.49786    8.07127    106    75.7605
128  100000     4.58122    8.07127    106    76.6948
129  100000     4.66395    8.07438    106    77.6072
130  100000     4.74575    8.24220    107    78.4898
131  100000     4.82876    8.73325    110    79.3724
132  100000     4.92057    8.39412    108    80.3310
133  100000     5.01065    8.72989    110    81.2566
134  100000     5.09971    9.44588    114    82.1517
135  100000     5.18901    9.09306    112    83.0317
136  100000     5.28565    9.09656    112    83.9733
137  100000     5.38103   10.41299    119    84.8820
138  100000     5.47926    9.63109    115    85.8007
139  100000     5.57617    9.81994    116    86.6919
140  100000     5.67844   10.19704    118    87.6188
141  100000     5.78447   10.01249    117    88.5611
142  100000     5.89554   10.01249    117    89.5285
143  100000     6.00749   10.61308    120    90.4873
144  100000     6.11700   10.40898    119    91.4062
145  100000     6.23273   12.90747    130    92.3645
146   96404     6.35155   11.47911    124    93.3267
147  100000     6.47036   11.92909    126    94.2686
148  100000     6.58920   11.48794    124    95.1993
149  100000     6.70518   11.46587    124    96.0945
150  100000     6.82857   11.46587    124    97.0158
151  100000     6.95216   12.41103    128    97.9372
152  100000     7.08122   11.94746    126    98.8728
153  100000     7.20863   12.19109    127    99.7808
154  100000     7.34016   12.67386    129   100.7051
155  100000     7.46863   12.92734    130   101.5915
156  100000     7.59572   12.67386    129   102.4509
157  100000     7.73579   13.67646    133   103.3795
158  100000     7.87774   13.70805    133   104.3093
159  100000     8.02077   14.81523    137   105.2249
160  100000     8.17804   14.22352    135   106.2174
161  100000     8.32210   14.82663    137   107.1025
162  100000     8.46829   14.53591    136   107.9968
163  100000     8.62181   15.40191    139   108.9151
164  100000     8.78320   14.80384    137   109.8646
165  100000     8.94697   16.61393    143   110.8097
166  100000     9.10481   15.11734    138   111.7047
167  100000     9.26120   16.30697    142   112.5752
168  100000     9.42322   15.71599    140   113.4625
169  100000     9.58953   17.29843    145   114.3578
170  100000     9.75776   17.65118    146   115.2468
171  100000     9.92404   17.96962    147   116.1073
172  100000    10.10693   18.70279    149   117.0421
173  100000    10.28652   17.99729    147   117.9419
174  100000    10.46715   17.99037    147   118.8372
175  100000    10.63893   17.65118    146   119.6672
176  100000    10.82231   18.01113    147   120.5405
177  100000    11.00593   18.36429    148   121.4059
178  100000    11.18825   19.86282    152   122.2414
179  100000    11.37738   18.35723    148   123.1007
180  100000    11.57435   21.03000    155   123.9749
181  100000    11.76946   19.84755    152   124.8327
182  100000    11.98200   21.04618    155   125.7459
183  100000    12.18663   21.04618    155   126.6188
184  100000    12.40235   21.04618    155   127.5138
185  100000    12.61659   21.88803    157   128.3931
186  100000    12.83390   21.46711    156   129.2715
187  100000    13.04983   21.85439    157   130.1277
188  100000    13.27369   22.28291    158   131.0027
189  100000    13.48436   22.73731    159   131.8112
190  100000    13.69913   23.62862    161   132.6240
191  100000    13.92304   25.05560    164   133.4548
192  100000    14.14923   24.11046    162   134.2794
193  100000    14.37590   25.06524    164   135.0960
194  100000    14.59150   25.09417    164   135.8611
195  100000    14.81275   25.08453    164   136.6306
196  100000    15.05407   25.55672    165   137.4610
197  100000    15.27499   24.60213    163   138.2120
198  100000    15.50803   24.60213    163   138.9860
199  100000    15.75525   26.08791    166   139.7976
200  100000    15.98980   27.10014    168   140.5591
201  100000    16.23344   26.07788    166   141.3348
202  100000    16.48663   27.65278    169   142.1298
203  100000    16.73635   28.17331    170   142.9015
204  100000    16.98657   27.63152    169   143.6696
205  100000    17.24622   27.66341    169   144.4458
206  100000    17.49674   29.31151    172   145.1871
207  100000    17.75417   31.05780    175   145.9360
208  100000    18.00673   29.26646    172   146.6642
209  100000    18.26571   30.47225    174   147.3965
210  100000    18.51870   29.87476    173   148.1041
211  100000    18.78249   29.89774    173   148.8333
212  100000    19.05072   29.88625    173   149.5641
213  100000    19.30394   31.05780    175   150.2456
214  100000    19.57309   31.06975    175   150.9588
215  100000    19.84625   31.05780    175   151.6741
216  100000    20.13194   30.48397    174   152.4092
217  100000    20.39017   32.28770    177   153.0645
218  100000    20.66081   31.69114    176   153.7409
219  100000    20.93688   34.90895    181   154.4266
220  100000    21.20225   33.59212    179   155.0761
221  100000    21.46698   33.56629    179   155.7144
222  100000    21.74187   36.30530    183   156.3712
223  100000    22.01967   33.57920    179   157.0240
224  100000    22.29074   34.88211    181   157.6556
225  100000    22.57183   34.90895    181   158.3011
226  100000    22.84129   34.90895    181   158.9124
227  100000    23.09222   37.71398    185   159.4735
228  100000    23.37912   34.92237    181   160.1106
229  100000    23.65954   37.74300    185   160.7259
230  100000    23.93104   35.59344    182   161.3149
231  100000    24.22917   36.29135    183   161.9532
232  100000    24.51557   38.48306    186   162.5601
233  100000    24.77948   36.30530    183   163.1135
234  100000    25.04654   39.23763    187   163.6664
235  100000    25.30155   39.22254    187   164.1897
236  100000    25.57437   37.74300    185   164.7430
237  100000    25.83819   40.00699    188   165.2738
238  100000    26.12080   39.99161    188   165.8350
239  100000    26.39546   38.48306    186   166.3748
240  100000    26.67070   41.57529    190   166.9117
241  100000    26.94100   39.23763    187   167.4331
242  100000    27.19286   40.79144    189   167.9149
243  100000    27.46544   40.00699    188   168.4302
244  100000    27.72759   40.79144    189   168.9179
245  100000    28.00741   41.57529    190   169.4383
246  100000    28.26232   42.40679    191   169.9079
247  100000    28.51981   41.59128    190   170.3774
248  100000    28.79192   41.59128    190   170.8701
249  100000    29.03120   45.83192    195   171.2981
250  100000    29.28778   41.59128    190   171.7514
251  100000    29.55762   41.59128    190   172.2245
252  100000    29.83357   43.23830    192   172.7052
253  100000    30.10557   42.40679    191   173.1764
254  100000    30.37251   42.40679    191   173.6318
255  100000    30.61689   43.23830    192   174.0458
256  100000    30.87240   44.08611    193   174.4752
257  100000    31.12060   44.08611    193   174.8885
258  100000    31.38707   43.23830    192   175.3282
259  100000    31.65110   44.06916    193   175.7620
260  100000    31.89558   44.08611    193   176.1610
261  100000    32.14018   46.73059    196   176.5549
262  100000    32.39936   44.08611    193   176.9718
263  100000    32.63934   44.08611    193   177.3543
264  100000    32.90842   45.83192    195   177.7791
265  100000    33.14673   44.95054    194   178.1527
266  100000    33.39271   46.73059    196   178.5344
267  100000    33.62499   46.73059    196   178.8942
268  100000    33.87394   47.64688    197   179.2746
269  100000    34.11224   46.73059    196   179.6384
270  100000    34.35081   44.95054    194   179.9991
271  100000    34.57997   46.73059    196   180.3423
272  100000    34.80226   48.58113    198   180.6739
273  100000    35.04879   47.64688    197   181.0406
274  100000    35.26368   48.58113    198   181.3568
275  100000    35.48254   46.73059    196   181.6769
276  100000    35.71403   48.58113    198   182.0131
277  100000    35.91753   48.58113    198   182.3083
278  100000    36.10038   47.64688    197   182.5727
279  100000    36.30270   48.58113    198   182.8628
280  100000    36.50784   48.58113    198   183.1536
281  100000    36.72144   48.58113    198   183.4556
282  100000    36.91460   48.58113    198   183.7277
283  100000    37.12307   49.53370    199   184.0187
284  100000    37.32484   48.58113    198   184.2995
285  100000    37.52802   48.58113    198   184.5814
286  100000    37.72234   48.58113    198   184.8506
287  100000    37.91138   48.58113    198   185.1102
288  100000    38.10407   48.58113    198   185.3723
289  100000    38.28991   49.53370    199   185.6237
290  100000    38.46950   48.58113    198   185.8670
291  100000    38.65662   48.58113    198   186.1181
292  100000    38.84437   49.53370    199   186.3695
293  100000    39.02531   48.58113    198   186.6104
294  100000    39.21338   49.53370    199   186.8587
295  100000    39.38980   49.53370    199   187.0905
296  100000    39.56913   50.50495    200   187.3265
297  100000    39.74044   49.53370    199   187.5522
298  100000    39.92248   49.53370    199   187.7882
299  100000    40.08948   50.50495    200   188.0051
300  100000    40.25702   49.53370    199   188.2218
301  100000    40.41816   50.50495    200   188.4299
302  100000    40.59813   50.50495    200   188.6601
303  100000    40.75620   50.50495    200   188.8614
304  100000    40.91424   50.50495    200   189.0620
305  100000    41.07525   49.53370    199   189.2656
306  100000    41.23758   50.50495    200   189.4700
307  100000    41.38205   50.50495    200   189.6514
308  100000    41.53113   50.50495    200   189.8381
309  100000    41.66301   50.50495    200   190.0019
310  100000    41.80781   50.50495    200   190.1824
311  100000    41.95344   50.50495    200   190.3632
312  100000    42.10680   50.50495    200   190.5521
313  100000    42.25206   50.50495    200   190.7308
314  100000    42.40194   50.50495    200   190.9149
315  100000    42.54629   50.50495    200   191.0908
316  100000    42.68120   50.50495    200   191.2559
317  100000    42.82195   50.50495    200   191.4264
318  100000    42.96606   50.50495    200   191.6007
319  100000    43.09620   50.50495    200   191.7576
320  100000    43.23534   50.50495    200   191.9249
321  100000    43.36521   50.50495    200   192.0810
322  100000    43.48618   50.50495    200   192.2255
323  100000    43.59436   50.50495    200   192.3549
324  100000    43.72087   50.50495    200   192.5055
325  100000    43.83513   50.50495    200   192.6411
326  100000    43.94987   50.50495    200   192.7765
327  100000    44.06057   50.50495    200   192.9069
328  100000    44.16527   50.50495    200   193.0306
329  100000    44.28160   50.50495    200   193.1675
330  100000    44.38308   50.50495    200   193.2858
331  100000    44.49626   50.50495    200   193.4185
332  100000    44.60260   50.50495    200   193.5426
333  100000    44.70798   50.50495    200   193.6652
334  100000    44.81110   50.50495    200   193.7850
335  100000    44.90636   50.50495    200   193.8943
336  100000    45.01352   50.50495    200   194.0186
337  100000    45.11515   50.50495    200   194.1358
338  100000    45.20758   50.50495    200   194.2422
339  100000    45.30434   50.50495    200   194.3532
340  100000    45.39761   50.50495    200   194.4592
341  100000    45.48878   50.50495    200   194.5634
342  100000    45.57240   50.50495    200   194.6586
343  100000    45.65885   50.50495    200   194.7573
344  100000    45.74389   50.50495    200   194.8537
345  100000    45.83082   50.50495    200   194.9523
346  100000    45.90729   50.50495    200   195.0388
347  100000    45.99105   50.50495    200   195.1339
348  100000    46.07537   50.50495    200   195.2286
349  100000    46.14863   50.50495    200   195.3113
350  100000    46.21406   50.50495    200   195.3850
351  100000    46.30130   50.50495    200   195.4827
352  100000    46.37897   50.50495    200   195.5701
353  100000    46.46038   50.50495    200   195.6610
354  100000    46.52394   50.50495    200   195.7322
355  100000    46.60229   50.50495    200   195.8200
356  100000    46.67216   50.50495    200   195.8982
357  100000    46.73666   50.50495    200   195.9697
358  100000    46.80469   50.50495    200   196.0454
359  100000    46.86401   50.50495    200   196.1113
360  100000    46.93073   50.50495    200   196.1852
361  100000    46.99317   50.50495    200   196.2540
362  100000    47.05833   50.50495    200   196.3258
363  100000    47.12029   50.50495    200   196.3941
364  100000    47.19219   50.50495    200   196.4734
365  100000    47.25414   50.50495    200   196.5415
366  100000    47.30132   50.50495    200   196.5932
367  100000    47.35749   50.50495    200   196.6550
368  100000    47.41283   50.50495    200   196.7156
369  100000    47.46828   50.50495    200   196.7763
370  100000    47.51829   50.50495    200   196.8308
371  100000    47.57693   50.50495    200   196.8949
372  100000    47.62793   50.50495    200   196.9505
373  100000    47.68421   50.50495    200   197.0121
374  100000    47.73425   50.50495    200   197.0665
375  100000    47.77843   50.50495    200   197.1149
376  100000    47.83521   50.50495    200   197.1767
377  100000    47.87898   50.50495    200   197.2240
378  100000    47.92467   50.50495    200   197.2737
379  100000    47.97440   50.50495    200   197.3277
380  100000    48.01318   50.50495    200   197.3694
381  100000    48.06737   50.50495    200   197.4281
382  100000    48.11084   50.50495    200   197.4751
383  100000    48.15761   50.50495    200   197.5256
384  100000    48.20373   50.50495    200   197.5753
385  100000    48.24705   50.50495    200   197.6223
386  100000    48.28967   50.50495    200   197.6682
387  100000    48.33079   50.50495    200   197.7122
388  100000    48.36828   50.50495    200   197.7526
389  100000    48.40605   50.50495    200   197.7932
390  100000    48.44735   50.50495    200   197.8373
391  100000    48.49144   50.50495    200   197.8845
392  100000    48.52847   50.50495    200   197.9242
393  100000    48.56681   50.50495    200   197.9654
394  100000    48.60127   50.50495    200   198.0023
395  100000    48.63057   50.50495    200   198.0335
396  100000    48.66895   50.50495    200   198.0745
397  100000    48.70866   50.50495    200   198.1170
398  100000    48.75727   50.50495    200   198.1687
399  100000    48.79140   50.50495    200   198.2054
400  100000    48.83102   50.50495    200   198.2475

Diploid Model with selection (random number based on
relative fitness to determine who survives).  Mutation rate is      1e-08,
Recomb rate is ONE per sexual individual per generation
Starting values at generation   0 are; population size 100000,
with avg fitness of    1.22101.

gen  popSize    avgFit     maxFit maxAlleles avgAlleles
  0  100000     1.22101    1.63934     25     9.9943
  1  100000     1.22101    1.63934     25     9.9943
  2  100000     1.22588    1.67277     26    10.1941
  3  100000     1.23081    1.73969     28    10.3952
  4  100000     1.23568    1.63998     25    10.5926
  5  100000     1.24061    1.67213     26    10.7915
  6  100000     1.24516    1.74035     28    10.9763
  7  100000     1.25050    1.67277     26    11.1911
  8  100000     1.25532    1.73969     28    11.3842
  9  100000     1.26063    1.74035     28    11.5965
 10  100000     1.26579    1.81067     30    11.8009
 11  100000     1.27165    1.73969     28    12.0332
 12  100000     1.27702    1.95766     34    12.2433
 13  100000     1.28284    1.77448     29    12.4713
 14  100000     1.28831    1.80997     30    12.6840
 15  100000     1.29420    1.81067     30    12.9129
 16  100000     1.30090    1.88164     32    13.1713
 17  100000     1.30736    1.95842     34    13.4198
 18  100000     1.31422    1.92002     33    13.6820
 19  100000     1.32119    1.84546     31    13.9468
 20  100000     1.32731    1.84688     31    14.1777
 21  100000     1.33443    1.88309     32    14.4458
 22  100000     1.34198    1.91854     33    14.7289
 23  100000     1.34929    1.88164     32    15.0010
 24  100000     1.35672    1.92075     33    15.2765
 25  100000     1.36524    1.92002     33    15.5909
 26  100000     1.37317    1.95992     34    15.8812
 27  100000     1.38101    1.92149     33    16.1676
 28  100000     1.38941    1.99758     35    16.4718
 29  100000     1.39748    1.99451     35    16.7622
 30  100000     1.40661    1.99912     35    17.0888
 31  100000     1.41531    2.11822     38    17.3979
 32  100000     1.42404    2.03754     36    17.7059
 33  100000     1.43332    2.12067     38    18.0294
 34  100000     1.44325    2.16142     39    18.3773
 35  100000     1.45330    2.20549     40    18.7276
 36  100000     1.46295    2.16142     39    19.0615
 37  100000     1.47374    2.16059     39    19.4291
 38  100000     1.48329    2.20549     40    19.7534
 39  100000     1.49401    2.24874     41    20.1138
 40  100000     1.50412    2.20634     40    20.4526
 41  100000     1.51454    2.33959     43    20.7979
 42  100000     1.52601    2.20634     40    21.1774
 43  100000     1.53763    2.29195     42    21.5566
 44  100000     1.54971    2.29548     42    21.9526
 45  100000     1.56149    2.33779     43    22.3321
 46  100000     1.57322    2.43224     45    22.7090
 47  100000     1.58610    2.52855     47    23.1174
 48  100000     1.59801    2.48183     46    23.4926
 49  100000     1.61077    2.58309     48    23.8914
 50  100000     1.62465    2.43224     45    24.3244
 51  100000     1.63829    2.53439     47    24.7461
 52  100000     1.65327    2.68539     50    25.2029
 53  100000     1.66775    2.62868     49    25.6397
 54  100000     1.68403    2.68332     50    26.1276
 55  100000     1.69915    2.74120     51    26.5758
 56  100000     1.71617    2.68642     50    27.0777
 57  100000     1.73143    2.68642     50    27.5237
 58  100000     1.74823    2.74015     51    28.0112
 59  100000     1.76499    2.74120     51    28.4904
 60  100000     1.78139    3.08229     57    28.9561
 61  100000     1.79999    2.85085     53    29.4779
 62  100000     1.81805    3.08111     57    29.9796
 63  100000     1.83608    3.02302     56    30.4770
 64  100000     1.85438    2.79602     52    30.9764
 65  100000     1.87328    3.26970     60    31.4871
 66  100000     1.89184    3.07401     57    31.9827
 67  100000     1.91148    3.20682     59    32.5003
 68  100000     1.93225    3.08348     57    33.0442
 69  100000     1.95303    3.27221     60    33.5836
 70  100000     1.97541    3.08229     57    34.1536
 71  100000     1.99731    3.39656     62    34.7101
 72  100000     2.01955    3.20805     59    35.2682
 73  100000     2.04438    3.40179     62    35.8815
 74  100000     2.06833    3.47116     63    36.4702
 75  100000     2.09403    3.40441     62    37.0908
 76  100000     2.11886    3.67938     66    37.6856
 77  100000     2.14618    3.47250     63    38.3280
 78  100000     2.17226    3.75296     67    38.9403
 79  100000     2.19985    3.68079     66    39.5752
 80  100000     2.22782    4.05765     71    40.2095
 81  100000     2.25592    3.98268     70    40.8398
 82  100000     2.28566    3.82802     68    41.5016
 83  100000     2.31706    3.90308     69    42.1878
 84  100000     2.34900    3.82950     68    42.8787
 85  100000     2.38013    3.75730     67    43.5478
 86  100000     2.41299    3.90158     69    44.2388
 87  100000     2.44567    3.97961     70    44.9158
 88  100000     2.47916    4.30601     74    45.6036
 89  100000     2.51443    4.13880     72    46.3169
 90  100000     2.55104    4.14836     72    47.0490
 91  100000     2.58859    4.39213     75    47.7863
 92  100000     2.62432    4.39720     75    48.4796
 93  100000     2.66126    4.75784     79    49.1832
 94  100000     2.70208    4.93675     81    49.9519
 95  100000     2.74059    4.56606     77    50.6685
 96  100000     2.78074    4.57484     77    51.4038
 97  100000     2.82251    4.76333     79    52.1576
 98  100000     2.86506    5.13817     83    52.9121
 99  100000     2.90851    5.35398     85    53.6705
100  100000     2.95122    5.35398     85    54.4106
101  100000     2.99731    5.24093     84    55.1949
102  100000     3.04483    5.14014     83    55.9872
103  100000     3.09497    5.24295     84    56.8125
104  100000     3.14473    5.56600     87    57.6183
105  100000     3.19584    5.67732     88    58.4309
106  100000     3.25145    5.67513     88    59.3036
107  100000     3.30849    6.13822     92    60.1851
108  100000     3.36441    5.77530     89    61.0360
109  100000     3.42279    6.00631     91    61.9038
110  100000     3.48000    6.27545     93    62.7425
111  100000     3.54087    6.02018     91    63.6267
112  100000     3.59784    6.01555     91    64.4395
113  100000     3.65882    6.13822     92    65.2915
114  100000     3.72166    6.26099     93    66.1537
115  100000     3.78387    6.38130     94    66.9974
116  100000     3.84622    6.63655     96    67.8242
117  100000     3.91550    7.05360     99    68.7335
118  100000     3.98160    6.90467     98    69.5770
119  100000     4.05217    7.17533    100    70.4667
120  100000     4.12408    6.89936     98    71.3571
121  100000     4.19651    7.31321    101    72.2435
122  100000     4.27081    7.75786    104    73.1314
123  100000     4.34546    7.76084    104    74.0088
124  100000     4.42478    7.76980    104    74.9305
125  100000     4.50589    8.23587    107    75.8469
126  100000     4.58715    8.39412    108    76.7564
127  100000     4.66916    8.07127    106    77.6587
128  100000     4.75479    8.57519    109    78.5795
129  100000     4.83997    8.55214    109    79.4813
130  100000     4.92435    8.56530    109    80.3621
131  100000     5.01047    8.89422    111    81.2412
132  100000     5.10721    9.07909    112    82.2121
133  100000     5.19705    9.64221    115    83.0954
134  100000     5.29582    9.44588    114    84.0516
135  100000     5.40014   10.00479    117    85.0384
136  100000     5.50202    9.63109    115    85.9909
137  100000     5.60771   10.60492    120    86.9597
138  100000     5.71050   10.20881    118    87.8895
139  100000     5.82034   10.62125    120    88.8634
140  100000     5.92665   10.63351    120    89.7718
141  100000     6.03712   11.70419    125    90.7159
142  100000     6.15398   11.70869    125    91.6937
143  100000     6.26996   11.23241    123    92.6505
144  100000     6.39154   11.27135    123    93.6274
145  100000     6.51353   11.71770    125    94.5935
146  100000     6.63816   11.69519    125    95.5587
147  100000     6.76580   11.94286    126    96.5269
148  100000     6.89088   11.93827    126    97.4602
149  100000     7.02213   12.91244    130    98.4219
150  100000     7.15329   12.65925    129    99.3633
151  100000     7.28684   13.43410    132   100.3075
152  100000     7.42195   13.16056    131   101.2433
153  100000     7.56437   13.42893    132   102.2185
154  100000     7.70804   12.92237    130   103.1781
155  100000     7.85317   14.52473    136   104.1299
156  100000     8.00178   14.25089    135   105.0892
157  100000     8.15323   14.21258    135   106.0474
158  100000     8.30477   13.97146    134   106.9851
159  100000     8.46392   15.10572    138   107.9589
160  100000     8.62703   15.09992    138   108.9360
161  100000     8.78449   15.08831    138   109.8638
162  100000     8.94976   16.00568    141   110.8121
163  100000     9.12131   15.69787    140   111.7812
164  100000     9.28723   16.64590    143   112.7034
165  100000     9.44842   16.64590    143   113.5896
166  100000     9.61086   17.65797    146   114.4618
167  100000     9.78065   16.65230    143   115.3522
168  100000     9.95402   17.62405    146   116.2492
169  100000    10.13944   17.32506    145   117.1953
170  100000    10.32351   17.65118    146   118.1147
171  100000    10.50039   17.96962    147   118.9848
172  100000    10.67968   20.26007    153   119.8489
173  100000    10.87053   19.83992    152   120.7578
174  100000    11.06190   19.08418    150   121.6551
175  100000    11.26315   19.48084    151   122.5794
176  100000    11.46061   20.63351    154   123.4672
177  100000    11.65246   19.84755    152   124.3163
178  100000    11.84967   21.43411    156   125.1762
179  100000    12.06135   21.05428    155   126.0844
180  100000    12.27781   21.46711    156   126.9960
181  100000    12.48975   21.45885    156   127.8692
182  100000    12.71488   22.74605    159   128.7873
183  100000    12.93207   21.87120    157   129.6561
184  100000    13.14908   22.30863    158   130.5091
185  100000    13.36202   22.75480    159   131.3347
186  100000    13.58588   23.64680    161   132.1900
187  100000    13.81632   23.20990    160   133.0497
188  100000    14.03806   24.62106    163   133.8629
189  100000    14.27172   24.58322    163   134.7130
190  100000    14.51636   25.07488    164   135.5843
191  100000    14.75310   24.11974    162   136.4122
192  100000    15.00053   25.03634    164   137.2688
193  100000    15.23106   25.09417    164   138.0541
194  100000    15.47974   26.05783    166   138.8856
195  100000    15.70366   27.62089    169   139.6223
196  100000    15.95350   26.07788    166   140.4290
197  100000    16.20269   28.17331    170   141.2278
198  100000    16.44544   27.12099    168   141.9906
199  100000    16.68793   28.70365    171   142.7456
200  100000    16.94892   28.74783    171   143.5416
201  100000    17.20107   27.63152    169   144.3059
202  100000    17.46809   28.74783    171   145.0957
203  100000    17.72163   29.87476    173   145.8375
204  100000    17.98292   29.28898    172   146.5902
205  100000    18.23240   29.87476    173   147.2978
206  100000    18.50083   29.30024    172   148.0498
207  100000    18.77518   29.30024    172   148.8046
208  100000    19.04705   29.88625    173   149.5451
209  100000    19.31285   31.66678    176   150.2598
210  100000    19.58293   32.28770    177   150.9767
211  100000    19.84287   33.59212    179   151.6557
212  100000    20.12095   32.93345    178   152.3725
213  100000    20.38707   31.67896    176   153.0522
214  100000    20.65463   32.94612    178   153.7247
215  100000    20.93300   32.30012    177   154.4140
216  100000    21.21214   34.25079    180   155.0962
217  100000    21.49973   32.93345    178   155.7901
218  100000    21.77611   32.94612    178   156.4477
219  100000    22.05421   32.94612    178   157.1043
220  100000    22.32810   33.57920    179   157.7388
221  100000    22.62002   34.90895    181   158.4086
222  100000    22.89455   37.01717    184   159.0317
223  100000    23.17443   35.59344    182   159.6558
224  100000    23.45623   36.30530    183   160.2785
225  100000    23.73805   37.74300    185   160.8931
226  100000    24.02755   36.30530    183   161.5233
227  100000    24.29155   38.48306    186   162.0861
228  100000    24.55828   37.74300    185   162.6510
229  100000    24.84936   37.74300    185   163.2597
230  100000    25.12766   37.74300    185   163.8319
231  100000    25.41739   39.23763    187   164.4275
232  100000    25.69841   38.48306    186   164.9942
233  100000    25.98313   37.74300    185   165.5618
234  100000    26.25375   40.79144    189   166.0973
235  100000    26.52519   39.23763    187   166.6255
236  100000    26.80567   40.79144    189   167.1723
237  100000    27.08106   39.23763    187   167.6966
238  100000    27.35866   40.00699    188   168.2214
239  100000    27.62858   40.00699    188   168.7284
240  100000    27.90167   39.23763    187   169.2390
241  100000    28.17233   40.00699    188   169.7377
242  100000    28.44587   41.59128    190   170.2381
243  100000    28.72047   41.57529    190   170.7349
244  100000    28.98100   42.40679    191   171.2016
245  100000    29.26113   43.23830    192   171.6978
246  100000    29.53438   42.40679    191   172.1804
247  100000    29.79881   42.40679    191   172.6405
248  100000    30.06830   42.40679    191   173.1080
249  100000    30.31972   43.23830    192   173.5362
250  100000    30.58398   43.23830    192   173.9860
251  100000    30.82608   43.22167    192   174.3934
252  100000    31.09314   44.08611    193   174.8383
253  100000    31.35714   44.08611    193   175.2763
254  100000    31.61692   43.23830    192   175.7047
255  100000    31.86957   44.95054    194   176.1142
256  100000    32.10295   46.73059    196   176.4958
257  100000    32.36959   45.83192    195   176.9235
258  100000    32.61500   44.95054    194   177.3124
259  100000    32.85147   44.95054    194   177.6864
260  100000    33.07734   45.83192    195   178.0401
261  100000    33.29515   45.83192    195   178.3806
262  100000    33.52770   44.95054    194   178.7416
263  100000    33.77718   47.64688    197   179.1255
264  100000    34.02098   46.73059    196   179.4985
265  100000    34.24748   45.83192    195   179.8406
266  100000    34.48333   45.83192    195   180.1974
267  100000    34.71752   47.64688    197   180.5465
268  100000    34.95815   47.64688    197   180.9035
269  100000    35.19563   46.73059    196   181.2540
270  100000    35.41438   47.64688    197   181.5767
271  100000    35.64553   47.64688    197   181.9137
272  100000    35.85580   46.73059    196   182.2171
273  100000    36.08839   47.64688    197   182.5521
274  100000    36.30327   46.73059    196   182.8604
275  100000    36.51669   47.64688    197   183.1650
276  100000    36.72854   48.58113    198   183.4635
277  100000    36.90838   48.58113    198   183.7184
278  100000    37.10976   48.58113    198   184.0002
279  100000    37.30123   48.58113    198   184.2674
280  100000    37.50945   48.58113    198   184.5563
281  100000    37.71240   47.64688    197   184.8350
282  100000    37.90478   47.64688    197   185.0987
283  100000    38.10122   48.58113    198   185.3671
284  100000    38.29337   49.53370    199   185.6275
285  100000    38.48663   49.53370    199   185.8884
286  100000    38.67349   49.53370    199   186.1397
287  100000    38.86588   49.53370    199   186.3962
288  100000    39.04702   49.53370    199   186.6377
289  100000    39.22296   49.53370    199   186.8701
290  100000    39.40397   49.53370    199   187.1085
291  100000    39.57591   49.53370    199   187.3346
292  100000    39.74070   49.53370    199   187.5507
293  100000    39.93768   49.53370    199   187.8064
294  100000    40.11570   50.50495    200   188.0377
295  100000    40.29385   50.50495    200   188.2672
296  100000    40.44767   50.50495    200   188.4653
297  100000    40.61891   50.50495    200   188.6834
298  100000    40.77231   50.50495    200   188.8797
299  100000    40.92500   50.50495    200   189.0735
300  100000    41.08916   50.50495    200   189.2804
301  100000    41.24190   50.50495    200   189.4737
302  100000    41.40434   50.50495    200   189.6774
303  100000    41.56571   50.50495    200   189.8790
304  100000    41.72230   50.50495    200   190.0744
305  100000    41.87636   50.50495    200   190.2670
306  100000    42.00816   50.50495    200   190.4295
307  100000    42.15614   50.50495    200   190.6122
308  100000    42.30223   50.50495    200   190.7915
309  100000    42.44393   50.50495    200   190.9655
310  100000    42.59233   50.50495    200   191.1466
311  100000    42.73329   50.50495    200   191.3178
312  100000    42.86496   50.50495    200   191.4782
313  100000    42.99765   50.50495    200   191.6388
314  100000    43.12797   50.50495    200   191.7958
315  100000    43.26194   50.50495    200   191.9566
316  100000    43.39733   50.50495    200   192.1193
317  100000    43.51284   50.50495    200   192.2574
318  100000    43.64256   50.50495    200   192.4117
319  100000    43.75969   50.50495    200   192.5498
320  100000    43.89143   50.50495    200   192.7062
321  100000    44.00950   50.50495    200   192.8461
322  100000    44.11691   50.50495    200   192.9735
323  100000    44.22692   50.50495    200   193.1031
324  100000    44.33885   50.50495    200   193.2343
325  100000    44.44813   50.50495    200   193.3611
326  100000    44.55188   50.50495    200   193.4823
327  100000    44.65074   50.50495    200   193.5973
328  100000    44.76422   50.50495    200   193.7291
329  100000    44.87616   50.50495    200   193.8588
330  100000    44.97993   50.50495    200   193.9789
331  100000    45.06734   50.50495    200   194.0796
332  100000    45.16511   50.50495    200   194.1923
333  100000    45.26889   50.50495    200   194.3111
334  100000    45.36694   50.50495    200   194.4235
335  100000    45.46273   50.50495    200   194.5334
336  100000    45.55757   50.50495    200   194.6421
337  100000    45.65174   50.50495    200   194.7489
338  100000    45.73866   50.50495    200   194.8480
339  100000    45.82248   50.50495    200   194.9429
340  100000    45.90536   50.50495    200   195.0365
341  100000    45.99078   50.50495    200   195.1332
342  100000    46.07682   50.50495    200   195.2306
343  100000    46.16275   50.50495    200   195.3272
344  100000    46.24187   50.50495    200   195.4168
345  100000    46.31796   50.50495    200   195.5023
346  100000    46.38019   50.50495    200   195.5718
347  100000    46.45019   50.50495    200   195.6502
348  100000    46.51965   50.50495    200   195.7275
349  100000    46.59382   50.50495    200   195.8102
350  100000    46.65387   50.50495    200   195.8773
351  100000    46.71351   50.50495    200   195.9437
352  100000    46.78725   50.50495    200   196.0256
353  100000    46.85750   50.50495    200   196.1037
354  100000    46.92993   50.50495    200   196.1839
355  100000    46.99498   50.50495    200   196.2561
356  100000    47.06777   50.50495    200   196.3365
357  100000    47.12999   50.50495    200   196.4048
358  100000    47.19511   50.50495    200   196.4769
359  100000    47.25056   50.50495    200   196.5378
360  100000    47.30878   50.50495    200   196.6020
361  100000    47.36817   50.50495    200   196.6669
362  100000    47.42170   50.50495    200   196.7258
363  100000    47.47502   50.50495    200   196.7841
364  100000    47.53000   50.50495    200   196.8447
365  100000    47.58997   50.50495    200   196.9100
366  100000    47.64515   50.50495    200   196.9703
367  100000    47.69402   50.50495    200   197.0237
368  100000    47.74925   50.50495    200   197.0837
369  100000    47.80041   50.50495    200   197.1395
370  100000    47.85788   50.50495    200   197.2016
371  100000    47.91012   50.50495    200   197.2584
372  100000    47.95621   50.50495    200   197.3086
373  100000    48.00798   50.50495    200   197.3645
374  100000    48.06097   50.50495    200   197.4218
375  100000    48.10108   50.50495    200   197.4651
376  100000    48.14523   50.50495    200   197.5127
377  100000    48.18359   50.50495    200   197.5541
378  100000    48.22828   50.50495    200   197.6020
379  100000    48.27900   50.50495    200   197.6568
380  100000    48.31838   50.50495    200   197.6993
381  100000    48.35950   50.50495    200   197.7435
382  100000    48.39786   50.50495    200   197.7847
383  100000    48.43880   50.50495    200   197.8287
384  100000    48.47522   50.50495    200   197.8677
385  100000    48.51342   50.50495    200   197.9088
386  100000    48.55059   50.50495    200   197.9485
387  100000    48.58813   50.50495    200   197.9886
388  100000    48.62216   50.50495    200   198.0250
389  100000    48.65881   50.50495    200   198.0642
390  100000    48.68897   50.50495    200   198.0964
391  100000    48.72802   50.50495    200   198.1380
392  100000    48.76035   50.50495    200   198.1725
393  100000    48.79646   50.50495    200   198.2109
394  100000    48.82860   50.50495    200   198.2452
395  100000    48.85980   50.50495    200   198.2783
396  100000    48.89190   50.50495    200   198.3124
397  100000    48.92752   50.50495    200   198.3503
398  100000    48.95661   50.50495    200   198.3812
399  100000    48.98660   50.50495    200   198.4131
400  100000    49.01679   50.50495    200   198.4452

Diploid Model with selection (random number based on
relative fitness to determine who survives).  Mutation rate is      1e-08,
Recomb rate is ONE per sexual individual per generation
Starting values at generation   0 are; population size 100000,
with avg fitness of    1.22127.

gen  popSize    avgFit     maxFit maxAlleles avgAlleles
  0  100000     1.22127    1.74035     28    10.0042
  1  100000     1.22127    1.74035     28    10.0042
  2  100000     1.22619    1.63998     25    10.2054
  3  100000     1.23068    1.67085     26    10.3890
  4  100000     1.23579    1.67085     26    10.5960
  5  100000     1.24054    1.67213     26    10.7877
  6  100000     1.24548    1.77380     29    10.9860
  7  100000     1.25058    1.67277     26    11.1912
  8  100000     1.25607    1.67342     26    11.4111
  9  100000     1.26187    1.73969     28    11.6427
 10  100000     1.26740    1.70623     27    11.8621
 11  100000     1.27310    1.74035     28    12.0860
 12  100000     1.27850    1.73902     28    12.2991
 13  100000     1.28488    1.84475     31    12.5481
 14  100000     1.29077    1.88164     32    12.7773
 15  100000     1.29731    1.92002     33    13.0313
 16  100000     1.30354    1.84688     31    13.2709
 17  100000     1.30970    1.92075     33    13.5069
 18  100000     1.31643    1.88309     32    13.7648
 19  100000     1.32293    1.84617     31    14.0118
 20  100000     1.33017    1.92075     33    14.2859
 21  100000     1.33708    1.84617     31    14.5446
 22  100000     1.34487    1.88237     32    14.8362
 23  100000     1.35281    1.88454     32    15.1319
 24  100000     1.36029    1.92075     33    15.4091
 25  100000     1.36765    1.88382     32    15.6783
 26  100000     1.37530    2.11985     38    15.9587
 27  100000     1.38330    2.03910     36    16.2501
 28  100000     1.39104    1.99912     35    16.5298
 29  100000     1.39957    2.11985     38    16.8366
 30  100000     1.40858    2.07989     37    17.1595
 31  100000     1.41725    2.11904     38    17.4669
 32  100000     1.42654    2.07909     37    17.7955
 33  100000     1.43533    2.11985     38    18.1029
 34  100000     1.44461    2.08069     37    18.4277
 35  100000     1.45430    2.12067     38    18.7622
 36  100000     1.46440    2.12067     38    19.1090
 37  100000     1.47363    2.20210     40    19.4249
 38  100000     1.48368    2.24874     41    19.7645
 39  100000     1.49365    2.16225     39    20.1015
 40  100000     1.50496    2.34049     43    20.4793
 41  100000     1.51522    2.29283     42    20.8207
 42  100000     1.52591    2.47802     46    21.1741
 43  100000     1.53695    2.34139     43    21.5375
 44  100000     1.54912    2.47993     46    21.9345
 45  100000     1.56172    2.53439     47    22.3405
 46  100000     1.57413    2.43317     45    22.7358
 47  100000     1.58685    2.53147     47    23.1399
 48  100000     1.59947    2.53342     47    23.5385
 49  100000     1.61272    2.47993     46    23.9554
 50  100000     1.62541    2.58309     48    24.3478
 51  100000     1.63950    2.53147     47    24.7804
 52  100000     1.65429    2.63678     49    25.2314
 53  100000     1.66917    2.84975     53    25.6815
 54  100000     1.68427    2.68642     50    26.1306
 55  100000     1.69986    2.79173     52    26.5933
 56  100000     1.71719    2.68642     50    27.1032
 57  100000     1.73417    2.63273     49    27.5984
 58  100000     1.74995    2.63476     49    28.0560
 59  100000     1.76694    2.74331     51    28.5401
 60  100000     1.78489    2.90340     54    29.0484
 61  100000     1.80291    2.84647     53    29.5550
 62  100000     1.82230    3.02534     56    30.0925
 63  100000     1.84183    2.96146     55    30.6298
 64  100000     1.86032    3.08348     57    31.1345
 65  100000     1.88088    3.02418     56    31.6843
 66  100000     1.90074    3.08111     57    32.2154
 67  100000     1.92245    3.20682     59    32.7822
 68  100000     1.94372    3.08466     57    33.3362
 69  100000     1.96711    3.20928     59    33.9420
 70  100000     1.98912    3.33894     61    34.5011
 71  100000     2.01210    3.27599     60    35.0818
 72  100000     2.03577    3.40703     62    35.6682
 73  100000     2.05932    3.46050     63    36.2476
 74  100000     2.08416    3.54195     64    36.8532
 75  100000     2.10791    3.53786     64    37.4249
 76  100000     2.13430    3.60723     65    38.0500
 77  100000     2.16119    3.46449     63    38.6837
 78  100000     2.18722    3.67513     66    39.2864
 79  100000     2.21610    4.06545     71    39.9458
 80  100000     2.24343    4.06702     71    40.5635
 81  100000     2.27272    3.75441     67    41.2214
 82  100000     2.30237    3.75296     67    41.8716
 83  100000     2.33141    4.13562     72    42.5031
 84  100000     2.36139    4.05297     71    43.1490
 85  100000     2.39169    4.22320     73    43.7917
 86  100000     2.42440    3.90308     69    44.4775
 87  100000     2.45638    4.06233     71    45.1370
 88  100000     2.48975    4.13562     72    45.8181
 89  100000     2.52525    4.57308     77    46.5332
 90  100000     2.55838    4.30435     74    47.1914
 91  100000     2.59547    4.48514     76    47.9204
 92  100000     2.63148    4.48342     76    48.6135
 93  100000     2.66967    4.94435     81    49.3457
 94  100000     2.70775    4.56781     77    50.0579
 95  100000     2.74715    4.74505     79    50.7856
 96  100000     2.78799    4.57308     77    51.5331
 97  100000     2.83058    4.85113     80    52.2948
 98  100000     2.87509    5.03548     82    53.0861
 99  100000     2.92021    4.94435     81    53.8718
100  100000     2.96403    4.94815     81    54.6200
101  100000     3.01018    5.34164     85    55.4045
102  100000     3.05836    5.34164     85    56.2078
103  100000     3.10823    5.14212     83    57.0249
104  100000     3.16047    5.34164     85    57.8655
105  100000     3.21430    6.01324     91    58.7197
106  100000     3.26439    5.89533     90    59.5008
107  100000     3.31894    5.78419     89    60.3468
108  100000     3.37205    5.78196     89    61.1516
109  100000     3.42692    6.00862     91    61.9667
110  100000     3.48080    6.13586     92    62.7550
111  100000     3.53983    6.02018     91    63.6074
112  100000     3.59860    6.38866     94    64.4436
113  100000     3.66146    6.64166     96    65.3168
114  100000     3.72384    6.26581     93    66.1735
115  100000     3.79072    6.63655     96    67.0753
116  100000     3.85627    6.89936     98    67.9420
117  100000     3.92539    6.89936     98    68.8424
118  100000     3.99214    7.18638    100    69.6974
119  100000     4.05857    7.31321    101    70.5372
120  100000     4.12933    7.31603    101    71.4131
121  100000     4.20577    7.32447    101    72.3499
122  100000     4.28115    7.31321    101    73.2560
123  100000     4.35407    7.91910    105    74.1083
124  100000     4.43552    7.91910    105    75.0470
125  100000     4.52090    7.76084    104    76.0152
126  100000     4.60569    8.39090    108    76.9589
127  100000     4.68851    8.08370    106    77.8603
128  100000     4.77026    9.08957    112    78.7353
129  100000     4.86051    8.72989    110    79.6921
130  100000     4.95151    8.71982    110    80.6289
131  100000     5.04564    9.24288    113    81.5893
132  100000     5.14290    9.43136    114    82.5577
133  100000     5.24292    9.81994    116    83.5380
134  100000     5.34226    9.08957    112    84.4929
135  100000     5.44382    9.79354    116    85.4492
136  100000     5.54854   10.41699    119    86.4211
137  100000     5.65620   10.03176    117    87.3979
138  100000     5.76158   10.40098    119    88.3344
139  100000     5.87777   10.59269    120    89.3520
140  100000     5.98613   10.83367    121    90.2780
141  100000     6.10290   11.49236    124    91.2651
142  100000     6.22248   11.04185    122    92.2527
143  100000     6.34142   11.25836    123    93.2158
144  100000     6.45945   11.04610    122    94.1614
145  100000     6.58582   11.72221    125    95.1502
146  100000     6.70604   11.93368    126    96.0735
147  100000     6.83647   12.41103    128    97.0605
148  100000     6.96684   12.64465    129    98.0248
149  100000     7.10125   12.65438    129    98.9985
150  100000     7.23702   13.42893    132    99.9646
151  100000     7.37856   13.15550    131   100.9504
152  100000     7.51455   15.68581    140   101.8835
153   95448     7.66373   14.21805    135   102.8827
154  100000     7.80825   13.41345    132   103.8386
155  100000     7.95365   13.70278    133   104.7805
156  100000     8.10329   14.80384    137   105.7336
157  100000     8.24951   13.95535    134   106.6472
158  100000     8.40416   14.50799    136   107.5936
159  100000     8.55950   15.11153    138   108.5331
160  100000     8.72045   15.09411    138   109.4840
161  100000     8.88070   16.00568    141   110.4083
162  100000     9.05010   16.31951    142   111.3760
163  100000     9.22224   16.31324    142   112.3379
164  100000     9.39555   17.99729    147   113.2911
165  100000     9.57142   17.66476    146   114.2396
166  100000     9.73746   16.99188    144   115.1161
167  100000     9.92024   17.31174    145   116.0761
168  100000    10.09175   18.34312    148   116.9535
169  100000    10.27570   18.35017    148   117.8802
170  100000    10.45610   18.71718    149   118.7707
171  100000    10.64511   18.72438    149   119.6822
172  100000    10.83365   19.07684    150   120.5853
173  100000    11.02335   18.01113    147   121.4725
174  100000    11.21281   20.63351    154   122.3420
175  100000    11.41858   19.44342    151   123.2759
176  100000    11.62332   19.83992    152   124.1858
177  100000    11.83166   19.46586    151   125.0973
178  100000    12.04118   20.22116    153   125.9964
179  100000    12.24498   21.45060    156   126.8540
180  100000    12.45006   21.45060    156   127.7041
181  100000    12.66717   21.03809    155   128.5940
182  100000    12.87836   22.30005    158   129.4444
183  100000    13.09828   21.87120    157   130.3115
184  100000    13.30922   22.30863    158   131.1287
185  100000    13.53783   23.20097    160   132.0031
186  100000    13.76466   22.75480    159   132.8552
187  100000    13.99332   24.10119    162   133.7060
188  100000    14.21613   23.20097    160   134.5178
189  100000    14.44856   24.12901    162   135.3526
190  100000    14.68599   24.60213    163   136.1873
191  100000    14.92193   26.07788    166   137.0047
192  100000    15.15402   25.55672    165   137.7972
193  100000    15.39742   25.57638    165   138.6145
194  100000    15.63318   25.57638    165   139.3953
195  100000    15.88045   26.07788    166   140.2040
196  100000    16.12324   26.08791    166   140.9820
197  100000    16.37034   25.57638    165   141.7627
198  100000    16.61050   28.75889    171   142.5080
199  100000    16.85056   28.75889    171   143.2478
200  100000    17.09677   27.64214    169   143.9974
201  100000    17.35006   28.72573    171   144.7568
202  100000    17.59965   28.75889    171   145.4890
203  100000    17.84989   28.71468    171   146.2156
204  100000    18.11284   28.73678    171   146.9654
205  100000    18.37446   30.46054    174   147.6984
206  100000    18.63684   30.44883    174   148.4289
207  100000    18.90988   32.30012    177   149.1799
208  100000    19.16793   31.05780    175   149.8778
209  100000    19.43030   30.47225    174   150.5779
210  100000    19.69624   32.92079    178   151.2803
211  100000    19.96617   31.08170    175   151.9798
212  100000    20.23987   31.06975    175   152.6814
213  100000    20.51648   31.65460    176   153.3805
214  100000    20.79682   32.30012    177   154.0779
215  100000    21.05859   32.31254    177   154.7247
216  100000    21.33184   34.90895    181   155.3905
217  100000    21.58987   33.57920    179   156.0101
218  100000    21.85584   33.59212    179   156.6432
219  100000    22.14060   34.90895    181   157.3081
220  100000    22.42430   34.25079    180   157.9640
221  100000    22.70582   34.23762    180   158.6076
222  100000    22.98499   36.27739    183   159.2344
223  100000    23.24735   34.90895    181   159.8188
224  100000    23.53282   35.60713    182   160.4480
225  100000    23.80789   36.29135    183   161.0509
226  100000    24.08250   36.30530    183   161.6421
227  100000    24.37513   37.74300    185   162.2649
228  100000    24.66002   38.46826    186   162.8632
229  100000    24.92136   37.72849    185   163.4078
230  100000    25.19292   38.48306    186   163.9653
231  100000    25.47234   38.46826    186   164.5364
232  100000    25.75019   38.48306    186   165.0961
233  100000    26.04644   40.79144    189   165.6853
234  100000    26.31785   38.46826    186   166.2211
235  100000    26.57273   39.23763    187   166.7203
236  100000    26.83878   38.48306    186   167.2338
237  100000    27.11260   40.00699    188   167.7590
238  100000    27.38574   40.00699    188   168.2762
239  100000    27.66646   40.00699    188   168.8023
240  100000    27.96005   45.83192    195   169.3513
241  100000    28.22372   40.00699    188   169.8359
242  100000    28.50247   40.79144    189   170.3430
243  100000    28.77592   41.57529    190   170.8370
244  100000    29.03032   41.59128    190   171.2926
245  100000    29.30092   41.59128    190   171.7725
246  100000    29.56506   41.59128    190   172.2362
247  100000    29.81280   41.59128    190   172.6683
248  100000    30.06584   42.40679    191   173.1060
249  100000    30.32320   42.40679    191   173.5473
250  100000    30.59964   42.40679    191   174.0148
251  100000    30.84576   43.23830    192   174.4304
252  100000    31.07844   43.23830    192   174.8197
253  100000    31.33356   44.95054    194   175.2430
254  100000    31.60832   44.08611    193   175.6933
255  100000    31.85871   44.08611    193   176.1028
256  100000    32.10255   44.08611    193   176.4963
257  100000    32.34766   44.95054    194   176.8895
258  100000    32.59580   44.95054    194   177.2824
259  100000    32.85468   44.08611    193   177.6935
260  100000    33.10744   44.95054    194   178.0907
261  100000    33.33134   44.95054    194   178.4402
262  100000    33.56557   45.83192    195   178.8032
263  100000    33.79472   44.95054    194   179.1562
264  100000    34.03780   45.83192    195   179.5262
265  100000    34.26259   45.83192    195   179.8652
266  100000    34.49118   46.73059    196   180.2117
267  100000    34.70938   48.58113    198   180.5385
268  100000    34.94448   46.73059    196   180.8870
269  100000    35.16933   46.73059    196   181.2188
270  100000    35.37630   47.64688    197   181.5219
271  100000    35.59655   48.58113    198   181.8442
272  100000    35.80959   46.73059    196   182.1538
273  100000    36.03141   47.64688    197   182.4732
274  100000    36.25314   46.73059    196   182.7918
275  100000    36.43716   48.58113    198   183.0529
276  100000    36.65544   47.64688    197   183.3636
277  100000    36.86762   47.64688    197   183.6630
278  100000    37.08091   47.64688    197   183.9616
279  100000    37.29783   48.58113    198   184.2629
280  100000    37.49118   49.53370    199   184.5321
281  100000    37.69030   48.58113    198   184.8064
282  100000    37.87571   49.53370    199   185.0602
283  100000    38.07458   48.58113    198   185.3305
284  100000    38.26659   48.58113    198   185.5921
285  100000    38.46967   49.53370    199   185.8667
286  100000    38.66041   49.53370    199   186.1233
287  100000    38.84892   49.53370    199   186.3754
288  100000    39.02161   49.53370    199   186.6048
289  100000    39.20991   49.53370    199   186.8550
290  100000    39.37713   49.53370    199   187.0750
291  100000    39.55363   49.53370    199   187.3074
292  100000    39.73503   49.53370    199   187.5439
293  100000    39.91123   49.53370    199   187.7734
294  100000    40.08470   50.50495    200   187.9975
295  100000    40.25425   50.50495    200   188.2165
296  100000    40.42849   49.53370    199   188.4404
297  100000    40.60923   50.50495    200   188.6719
298  100000    40.75953   50.50495    200   188.8642
299  100000    40.92982   50.50495    200   189.0811
300  100000    41.08818   50.50495    200   189.2813
301  100000    41.24979   50.50495    200   189.4849
302  100000    41.40199   50.50495    200   189.6754
303  100000    41.55987   50.50495    200   189.8740
304  100000    41.70179   50.50495    200   190.0510
305  100000    41.85136   50.50495    200   190.2371
306  100000    41.99730   50.50495    200   190.4175
307  100000    42.14032   50.50495    200   190.5932
308  100000    42.29143   50.50495    200   190.7791
309  100000    42.43797   50.50495    200   190.9594
310  100000    42.57907   50.50495    200   191.1308
311  100000    42.71005   50.50495    200   191.2904
312  100000    42.84324   50.50495    200   191.4527
313  100000    42.95780   50.50495    200   191.5912
314  100000    43.08146   50.50495    200   191.7394
315  100000    43.20963   50.50495    200   191.8935
316  100000    43.33865   50.50495    200   192.0489
317  100000    43.46522   50.50495    200   192.2005
318  100000    43.58833   50.50495    200   192.3474
319  100000    43.71186   50.50495    200   192.4950
320  100000    43.82868   50.50495    200   192.6332
321  100000    43.93523   50.50495    200   192.7592
322  100000    44.05317   50.50495    200   192.8981
323  100000    44.17206   50.50495    200   193.0386
324  100000    44.28521   50.50495    200   193.1712
325  100000    44.39454   50.50495    200   193.2997
326  100000    44.50646   50.50495    200   193.4306
327  100000    44.60797   50.50495    200   193.5489
328  100000    44.71111   50.50495    200   193.6686
329  100000    44.80430   50.50495    200   193.7766
330  100000    44.90673   50.50495    200   193.8954
331  100000    45.00208   50.50495    200   194.0054
332  100000    45.08310   50.50495    200   194.0987
333  100000    45.18146   50.50495    200   194.2119
334  100000    45.26504   50.50495    200   194.3077
335  100000    45.36498   50.50495    200   194.4222
336  100000    45.46075   50.50495    200   194.5320
337  100000    45.54109   50.50495    200   194.6238
338  100000    45.63678   50.50495    200   194.7330
339  100000    45.72058   50.50495    200   194.8280
340  100000    45.80330   50.50495    200   194.9218
341  100000    45.88614   50.50495    200   195.0160
342  100000    45.95927   50.50495    200   195.0987
343  100000    46.04988   50.50495    200   195.2012
344  100000    46.12051   50.50495    200   195.2809
345  100000    46.19776   50.50495    200   195.3680
346  100000    46.27247   50.50495    200   195.4517
347  100000    46.35301   50.50495    200   195.5416
348  100000    46.43084   50.50495    200   195.6287
349  100000    46.50484   50.50495    200   195.7117
350  100000    46.58049   50.50495    200   195.7959
351  100000    46.64828   50.50495    200   195.8713
352  100000    46.71610   50.50495    200   195.9469
353  100000    46.78946   50.50495    200   196.0282
354  100000    46.85446   50.50495    200   196.1003
355  100000    46.92154   50.50495    200   196.1750
356  100000    46.99526   50.50495    200   196.2566
357  100000    47.06242   50.50495    200   196.3305
358  100000    47.12568   50.50495    200   196.4004
359  100000    47.19086   50.50495    200   196.4720
360  100000    47.25423   50.50495    200   196.5420
361  100000    47.30763   50.50495    200   196.6002
362  100000    47.36414   50.50495    200   196.6624
363  100000    47.41969   50.50495    200   196.7236
364  100000    47.48232   50.50495    200   196.7921
365  100000    47.53795   50.50495    200   196.8530
366  100000    47.58892   50.50495    200   196.9087
367  100000    47.64650   50.50495    200   196.9711
368  100000    47.69434   50.50495    200   197.0234
369  100000    47.75130   50.50495    200   197.0853
370  100000    47.80199   50.50495    200   197.1405
371  100000    47.84867   50.50495    200   197.1913
372  100000    47.89705   50.50495    200   197.2437
373  100000    47.94406   50.50495    200   197.2948
374  100000    47.98832   50.50495    200   197.3428
375  100000    48.03785   50.50495    200   197.3964
376  100000    48.08799   50.50495    200   197.4503
377  100000    48.13131   50.50495    200   197.4973
378  100000    48.17516   50.50495    200   197.5446
379  100000    48.21404   50.50495    200   197.5865
380  100000    48.25176   50.50495    200   197.6271
381  100000    48.28678   50.50495    200   197.6649
382  100000    48.32295   50.50495    200   197.7039
383  100000    48.35826   50.50495    200   197.7419
384  100000    48.39547   50.50495    200   197.7821
385  100000    48.44032   50.50495    200   197.8302
386  100000    48.48266   50.50495    200   197.8755
387  100000    48.52229   50.50495    200   197.9181
388  100000    48.56824   50.50495    200   197.9672
389  100000    48.60305   50.50495    200   198.0046
390  100000    48.63131   50.50495    200   198.0346
391  100000    48.65880   50.50495    200   198.0640
392  100000    48.69572   50.50495    200   198.1036
393  100000    48.73112   50.50495    200   198.1413
394  100000    48.76586   50.50495    200   198.1784
395  100000    48.80505   50.50495    200   198.2202
396  100000    48.83246   50.50495    200   198.2494
397  100000    48.86517   50.50495    200   198.2842
398  100000    48.89158   50.50495    200   198.3121
399  100000    48.91727   50.50495    200   198.3394
400  100000    48.94403   50.50495    200   198.3680

Diploid Model with selection (random number based on
relative fitness to determine who survives).  Mutation rate is      1e-08,
Recomb rate is ONE per sexual individual per generation
Starting values at generation   0 are; population size 100000,
with avg fitness of    1.22133.

gen  popSize    avgFit     maxFit maxAlleles avgAlleles
  0  100000     1.22133    1.67277     26    10.0074
  1  100000     1.22133    1.67277     26    10.0074
  2  100000     1.22631    1.74035     28    10.2113
  3  100000     1.23028    1.70623     27    10.3723
  4  100000     1.23571    1.70426     27    10.5927
  5  100000     1.24101    1.67277     26    10.8081
  6  100000     1.24643    1.70689     27    11.0266
  7  100000     1.25153    1.88309     32    11.2325
  8  100000     1.25685    1.70492     27    11.4442
  9  100000     1.26194    1.77448     29    11.6464
 10  100000     1.26733    1.81136     30    11.8607
 11  100000     1.27284    1.80927     30    12.0778
 12  100000     1.27833    1.92002     33    12.2935
 13  100000     1.28409    1.81136     30    12.5185
 14  100000     1.29043    1.81067     30    12.7645
 15  100000     1.29656    1.84546     31    13.0021
 16  100000     1.30273    1.84617     31    13.2393
 17  100000     1.30948    1.84617     31    13.4976
 18  100000     1.31588    1.88309     32    13.7437
 19  100000     1.32206    1.92002     33    13.9778
 20  100000     1.32869    1.92075     33    14.2291
 21  100000     1.33593    2.03597     36    14.5004
 22  100000     1.34270    1.88454     32    14.7544
 23  100000     1.35008    2.03754     36    15.0303
 24  100000     1.35699    1.92149     33    15.2865
 25  100000     1.36460    2.03910     36    15.5664
 26  100000     1.37227    1.95917     34    15.8465
 27  100000     1.38064    1.96068     34    16.1498
 28  100000     1.38906    2.07829     37    16.4554
 29  100000     1.39718    2.07829     37    16.7486
 30  100000     1.40528    1.99835     35    17.0398
 31  100000     1.41378    2.03754     36    17.3435
 32  100000     1.42268    2.25047     41    17.6580
 33  100000     1.43215    2.07829     37    17.9898
 34  100000     1.44202    2.11985     38    18.3339
 35  100000     1.45206    2.16142     39    18.6806
 36  100000     1.46299    2.16225     39    19.0572
 37  100000     1.47329    2.16059     39    19.4082
 38  100000     1.48319    2.20380     40    19.7460
 39  100000     1.49355    2.20549     40    20.0985
 40  100000     1.50384    2.38638     44    20.4414
 41  100000     1.51405    2.16225     39    20.7830
 42  100000     1.52537    2.25047     41    21.1566
 43  100000     1.53621    2.33959     43    21.5137
 44  100000     1.54847    2.25047     41    21.9122
 45  100000     1.55998    2.43317     45    22.2825
 46  100000     1.57279    2.33869     43    22.6959
 47  100000     1.58408    2.43504     45    23.0571
 48  100000     1.59601    2.38363     44    23.4335
 49  100000     1.60888    2.43504     45    23.8357
 50  100000     1.62221    2.53245     47    24.2514
 51  100000     1.63555    2.68642     50    24.6630
 52  100000     1.64914    2.58012     48    25.0775
 53  100000     1.66357    2.57912     48    25.5138
 54  100000     1.67912    2.58012     48    25.9816
 55  100000     1.69362    2.68435     50    26.4126
 56  100000     1.70869    2.73699     51    26.8591
 57  100000     1.72435    2.73383     51    27.3165
 58  100000     1.73961    2.74225     51    27.7609
 59  100000     1.75474    2.74120     51    28.1983
 60  100000     1.77212    2.90787     54    28.6924
 61  100000     1.78974    2.79173     52    29.1910
 62  100000     1.80853    2.90563     54    29.7150
 63  100000     1.82713    2.84647     53    30.2304
 64  100000     1.84505    2.85194     53    30.7194
 65  100000     1.86443    2.96374     55    31.2476
 66  100000     1.88347    2.90451     54    31.7582
 67  100000     1.90349    3.08111     57    32.2925
 68  100000     1.92449    3.08111     57    32.8435
 69  100000     1.94500    3.14394     58    33.3777
 70  100000     1.96674    3.20189     59    33.9354
 71  100000     1.98899    3.14394     58    34.5015
 72  100000     2.01162    3.27221     60    35.0719
 73  100000     2.03389    3.20682     59    35.6279
 74  100000     2.05891    3.20682     59    36.2397
 75  100000     2.08332    3.46583     63    36.8322
 76  100000     2.10808    3.46849     63    37.4305
 77  100000     2.13261    3.53514     64    38.0148
 78  100000     2.15948    3.40179     62    38.6421
 79  100000     2.18586    3.53922     64    39.2545
 80  100000     2.21426    3.61279     65    39.9054
 81  100000     2.24268    3.53514     64    40.5488
 82  100000     2.27268    3.68646     66    41.2191
 83  100000     2.30242    3.90308     69    41.8755
 84  100000     2.33187    3.90458     69    42.5132
 85  100000     2.36132    3.82802     68    43.1443
 86  100000     2.39144    4.22645     73    43.7865
 87  100000     2.42589    4.56781     77    44.5087
 88  100000     2.45893    4.31098     74    45.1902
 89  100000     2.49452    4.14995     72    45.9130
 90  100000     2.52862    4.30270     74    46.5985
 91  100000     2.56480    4.30766     74    47.3140
 92  100000     2.60187    4.56430     77    48.0394
 93  100000     2.63954    4.48169     76    48.7670
 94  100000     2.67812    4.56606     77    49.5011
 95  100000     2.71743    4.65738     78    50.2390
 96  100000     2.75877    4.57660     77    51.0020
 97  100000     2.79780    4.57133     77    51.7125
 98  100000     2.84053    5.13619     83    52.4786
 99  100000     2.88368    4.74505     79    53.2414
100  100000     2.92787    5.23489     84    54.0112
101  100000     2.97339    5.24698     84    54.7904
102  100000     3.01713    5.24496     84    55.5285
103  100000     3.06601    5.23891     84    56.3398
104  100000     3.11395    5.45057     86    57.1284
105  100000     3.16127    5.44847     86    57.8932
106  100000     3.21110    5.67295     88    58.6836
107  100000     3.26378    5.77530     89    59.5053
108  100000     3.31348    6.01324     91    60.2715
109  100000     3.36764    5.90895     90    61.0880
110  100000     3.42055    5.89987     90    61.8783
111  100000     3.47886    6.25137     93    62.7332
112  100000     3.53772    6.14058     92    63.5838
113  100000     3.59577    6.37639     94    64.4066
114  100000     3.65785    6.25617     93    65.2708
115  100000     3.71984    6.50892     95    66.1223
116  100000     3.78408    6.38375     94    66.9901
117  100000     3.84820    6.63655     96    67.8413
118  100000     3.91163    6.64421     96    68.6672
119  100000     3.98180    6.89671     98    69.5689
120  100000     4.05348    7.04547     99    70.4760
121  100000     4.12547    7.04005     99    71.3727
122  100000     4.19672    7.75487    104    72.2439
123  100000     4.27194    7.45374    102    73.1417
124  100000     4.34872    8.06197    106    74.0478
125  100000     4.42808    7.61159    103    74.9679
126  100000     4.51163    7.91606    105    75.9125
127  100000     4.59763    8.23903    107    76.8753
128  100000     4.68288    7.91910    105    77.8103
129  100000     4.76801    8.39412    108    78.7241
130  100000     4.85835    9.26423    113    79.6805
131  100000     4.94670    8.89764    111    80.5998
132  100000     5.03207    8.73325    110    81.4647
133  100000     5.12415    9.62369    115    82.3893
134  100000     5.21426   10.00479    117    83.2733
135  100000     5.31113   10.22059    118    84.2131
136  100000     5.40676    9.63850    115    85.1196
137  100000     5.50363    9.81616    116    86.0292
138  100000     5.60345   10.60084    120    86.9386
139  100000     5.70877   10.00094    117    87.8875
140  100000     5.80872   10.22059    118    88.7674
141  100000     5.91854   10.42100    119    89.7230
142  100000     6.03133   10.61308    120    90.6823
143  100000     6.14997   11.04610    122    91.6699
144  100000     6.26817   10.82534    121    92.6415
145  100000     6.38368   11.49678    124    93.5722
146  100000     6.50664   11.25403    123    94.5415
147  100000     6.62518   11.70869    125    95.4642
148  100000     6.74809   12.42058    128    96.4012
149  100000     6.87041   11.95205    126    97.3117
150  100000     7.00215   13.16562    131    98.2827
151  100000     7.13035   14.26185    135    99.2119
152  100000     7.26366   13.15044    131   100.1473
153  100000     7.39464   13.15550    131   101.0620
154  100000     7.53467   13.43926    132   102.0223
155  100000     7.67814   13.68698    133   102.9857
156  100000     7.82191   13.69224    133   103.9293
157  100000     7.96644   14.80953    137   104.8674
158  100000     8.11443   14.80384    137   105.8041
159  100000     8.26490   14.51915    136   106.7432
160  100000     8.42028   14.24541    135   107.6920
161  100000     8.57338   15.10572    138   108.6059
162  100000     8.73786   14.79815    137   109.5785
163  100000     8.90030   16.01799    141   110.5267
164  100000     9.06941   16.01183    141   111.4907
165  100000     9.23363   15.69184    140   112.4088
166  100000     9.39441   16.33207    142   113.2917
167  100000     9.56732   16.97882    144   114.2233
168  100000     9.73969   16.98535    144   115.1346
169  100000     9.90802   17.30508    145   116.0076
170  100000    10.09179   18.35723    148   116.9452
171  100000    10.27019   18.72438    149   117.8469
172  100000    10.45420   18.68841    149   118.7551
173  100000    10.63751   18.70998    149   119.6489
174  100000    10.83030   19.06951    150   120.5676
175  100000    11.02263   18.74599    149   121.4714
176  100000    11.21676   20.23672    153   122.3672
177  100000    11.40628   20.60180    154   123.2221
178  100000    11.61074   20.22116    153   124.1356
179  100000    11.81127   21.45885    156   125.0186
180  100000    12.01831   20.63351    154   125.9075
181  100000    12.22830   21.87962    157   126.7953
182  100000    12.44273   20.64145    154   127.6836
183  100000    12.65268   21.87120    157   128.5388
184  100000    12.86954   21.90487    157   129.4114
185  100000    13.09644   22.27434    158   130.3047
186  100000    13.31740   23.21882    160   131.1634
187  100000    13.53510   23.18314    160   131.9959
188  100000    13.76643   22.32579    158   132.8611
189  100000    13.98862   25.56655    165   133.6891
190  100000    14.21674   23.20097    160   134.5200
191  100000    14.43992   23.65589    161   135.3162
192  100000    14.67487   24.60213    163   136.1422
193  100000    14.91190   24.59267    163   136.9660
194  100000    15.15908   24.58322    163   137.8085
195  100000    15.40075   25.06524    164   138.6198
196  100000    15.62786   25.56655    165   139.3725
197  100000    15.88513   25.54689    165   140.2126
198  100000    16.14202   27.11056    168   141.0361
199  100000    16.39522   27.64214    169   141.8358
200  100000    16.64754   27.62089    169   142.6237
201  100000    16.88333   27.62089    169   143.3430
202  100000    17.15372   27.65278    169   144.1632
203  100000    17.41866   28.72573    171   144.9496
204  100000    17.68146   28.73678    171   145.7214
205  100000    17.95264   29.27772    172   146.5057
206  100000    18.21250   29.31151    172   147.2478
207  100000    18.44737   29.31151    172   147.9042
208  100000    18.70581   29.30024    172   148.6188
209  100000    18.97429   29.88625    173   149.3540
210  100000    19.24812   30.46054    174   150.0889
211  100000    19.51376   31.08170    175   150.7965
212  100000    19.77755   31.08170    175   151.4883
213  100000    20.04628   32.28770    177   152.1857
214  100000    20.31898   31.08170    175   152.8781
215  100000    20.59168   32.30012    177   153.5649
216  100000    20.86973   32.26287    177   154.2563
217  100000    21.14774   35.59344    182   154.9369
218  100000    21.42545   32.94612    178   155.6109
219  100000    21.69399   32.92079    178   156.2564
220  100000    21.96026   34.90895    181   156.8842
221  100000    22.24199   33.59212    179   157.5417
222  100000    22.52782   35.59344    182   158.2016
223  100000    22.77224   33.57920    179   158.7574
224  100000    23.03542   34.90895    181   159.3529
225  100000    23.30671   36.30530    183   159.9572
226  100000    23.55220   36.30530    183   160.4960
227  100000    23.82154   37.01717    184   161.0823
228  100000    24.10698   34.92237    181   161.6958
229  100000    24.38678   36.98871    184   162.2900
230  100000    24.66567   37.74300    185   162.8793
231  100000    24.95794   37.01717    184   163.4872
232  100000    25.22281   37.72849    185   164.0289
233  100000    25.49925   37.74300    185   164.5918
234  100000    25.79070   38.48306    186   165.1778
235  100000    26.05601   39.22254    187   165.7099
236  100000    26.31682   38.46826    186   166.2223
237  100000    26.58794   38.46826    186   166.7549
238  100000    26.85415   38.48306    186   167.2678
239  100000    27.12604   40.77576    189   167.7870
240  100000    27.39687   40.79144    189   168.3002
241  100000    27.66648   40.00699    188   168.8044
242  100000    27.92633   40.00699    188   169.2879
243  100000    28.19829   40.00699    188   169.7899
244  100000    28.45798   40.79144    189   170.2625
245  100000    28.72988   41.59128    190   170.7525
246  100000    29.01115   43.22167    192   171.2572
247  100000    29.29035   41.59128    190   171.7501
248  100000    29.56645   42.40679    191   172.2374
249  100000    29.82013   41.59128    190   172.6795
250  100000    30.07671   44.08611    193   173.1222
251  100000    30.33595   42.40679    191   173.5694
252  100000    30.59333   43.23830    192   174.0040
253  100000    30.84860   43.23830    192   174.4349
254  100000    31.09187   43.23830    192   174.8415
255  100000    31.35185   43.23830    192   175.2734
256  100000    31.60759   43.23830    192   175.6935
257  100000    31.83816   44.95054    194   176.0685
258  100000    32.08041   44.06916    193   176.4594
259  100000    32.31405   44.95054    194   176.8339
260  100000    32.55237   44.95054    194   177.2158
261  100000    32.79117   45.83192    195   177.5942
262  100000    33.04660   44.08611    193   177.9956
263  100000    33.29005   45.83192    195   178.3762
264  100000    33.52894   44.95054    194   178.7462
265  100000    33.75374   45.83192    195   179.0915
266  100000    33.97655   44.95054    194   179.4326
267  100000    34.20497   46.73059    196   179.7796
268  100000    34.43433   47.64688    197   180.1250
269  100000    34.67058   46.73059    196   180.4801
270  100000    34.89985   46.73059    196   180.8199
271  100000    35.13223   46.73059    196   181.1624
272  100000    35.35519   46.73059    196   181.4908
273  100000    35.55645   47.64688    197   181.7841
274  100000    35.78903   47.64688    197   182.1232
275  100000    36.00353   47.64688    197   182.4330
276  100000    36.21759   48.58113    198   182.7422
277  100000    36.42028   47.64688    197   183.0301
278  100000    36.65364   47.64688    197   183.3610
279  100000    36.85253   47.64688    197   183.6410
280  100000    37.05168   47.64688    197   183.9204
281  100000    37.24849   48.58113    198   184.1941
282  100000    37.43274   48.58113    198   184.4503
283  100000    37.62153   48.58113    198   184.7113
284  100000    37.83172   48.58113    198   185.0002
285  100000    38.03407   49.53370    199   185.2760
286  100000    38.22743   48.58113    198   185.5379
287  100000    38.41694   50.50495    200   185.7957
288  100000    38.59893   49.53370    199   186.0405
289  100000    38.78783   49.53370    199   186.2932
290  100000    38.96696   49.53370    199   186.5331
291  100000    39.14446   48.58113    198   186.7700
292  100000    39.32819   50.50495    200   187.0115
293  100000    39.51406   49.53370    199   187.2553
294  100000    39.70385   49.53370    199   187.5047
295  100000    39.86771   50.50495    200   187.7183
296  100000    40.04066   49.53370    199   187.9434
297  100000    40.20305   50.50495    200   188.1525
298  100000    40.37626   49.53370    199   188.3758
299  100000    40.55346   50.50495    200   188.6031
300  100000    40.71971   50.50495    200   188.8154
301  100000    40.86882   50.50495    200   189.0047
302  100000    41.02483   50.50495    200   189.2010
303  100000    41.18816   49.53370    199   189.4084
304  100000    41.34468   50.50495    200   189.6055
305  100000    41.48579   50.50495    200   189.7822
306  100000    41.63505   50.50495    200   189.9680
307  100000    41.78895   50.50495    200   190.1594
308  100000    41.93423   50.50495    200   190.3386
309  100000    42.06791   50.50495    200   190.5040
310  100000    42.19914   50.50495    200   190.6660
311  100000    42.33994   50.50495    200   190.8386
312  100000    42.46632   50.50495    200   190.9933
313  100000    42.60134   50.50495    200   191.1587
314  100000    42.73046   50.50495    200   191.3153
315  100000    42.86627   50.50495    200   191.4805
316  100000    42.99796   50.50495    200   191.6399
317  100000    43.12342   50.50495    200   191.7909
318  100000    43.26067   50.50495    200   191.9551
319  100000    43.38912   50.50495    200   192.1094
320  100000    43.51619   50.50495    200   192.2610
321  100000    43.63555   50.50495    200   192.4039
322  100000    43.75801   50.50495    200   192.5494
323  100000    43.88396   50.50495    200   192.6984
324  100000    44.00682   50.50495    200   192.8431
325  100000    44.13369   50.50495    200   192.9929
326  100000    44.25165   50.50495    200   193.1318
327  100000    44.35660   50.50495    200   193.2549
328  100000    44.47452   50.50495    200   193.3923
329  100000    44.58760   50.50495    200   193.5244
330  100000    44.68894   50.50495    200   193.6424
331  100000    44.79212   50.50495    200   193.7621
332  100000    44.89340   50.50495    200   193.8795
333  100000    44.98915   50.50495    200   193.9901
334  100000    45.08368   50.50495    200   194.0989
335  100000    45.18331   50.50495    200   194.2138
336  100000    45.27512   50.50495    200   194.3194
337  100000    45.36208   50.50495    200   194.4184
338  100000    45.46286   50.50495    200   194.5338
339  100000    45.56272   50.50495    200   194.6479
340  100000    45.64245   50.50495    200   194.7389
341  100000    45.73601   50.50495    200   194.8450
342  100000    45.82021   50.50495    200   194.9406
343  100000    45.91422   50.50495    200   195.0470
344  100000    46.00696   50.50495    200   195.1520
345  100000    46.08597   50.50495    200   195.2408
346  100000    46.16679   50.50495    200   195.3319
347  100000    46.24998   50.50495    200   195.4257
348  100000    46.31887   50.50495    200   195.5030
349  100000    46.38309   50.50495    200   195.5751
350  100000    46.44923   50.50495    200   195.6493
351  100000    46.52501   50.50495    200   195.7339
352  100000    46.59610   50.50495    200   195.8132
353  100000    46.66567   50.50495    200   195.8902
354  100000    46.73722   50.50495    200   195.9700
355  100000    46.79724   50.50495    200   196.0367
356  100000    46.86701   50.50495    200   196.1143
357  100000    46.92821   50.50495    200   196.1824
358  100000    46.99129   50.50495    200   196.2517
359  100000    47.06212   50.50495    200   196.3299
360  100000    47.13091   50.50495    200   196.4061
361  100000    47.18765   50.50495    200   196.4688
362  100000    47.25527   50.50495    200   196.5431
363  100000    47.31872   50.50495    200   196.6127
364  100000    47.37161   50.50495    200   196.6709
365  100000    47.42650   50.50495    200   196.7310
366  100000    47.48550   50.50495    200   196.7957
367  100000    47.54751   50.50495    200   196.8633
368  100000    47.61041   50.50495    200   196.9318
369  100000    47.67097   50.50495    200   196.9981
370  100000    47.72772   50.50495    200   197.0600
371  100000    47.77575   50.50495    200   197.1122
372  100000    47.83025   50.50495    200   197.1715
373  100000    47.87660   50.50495    200   197.2222
374  100000    47.92595   50.50495    200   197.2757
375  100000    47.97288   50.50495    200   197.3260
376  100000    48.02182   50.50495    200   197.3792
377  100000    48.06748   50.50495    200   197.4288
378  100000    48.11735   50.50495    200   197.4825
379  100000    48.15925   50.50495    200   197.5279
380  100000    48.19506   50.50495    200   197.5662
381  100000    48.23473   50.50495    200   197.6091
382  100000    48.27566   50.50495    200   197.6532
383  100000    48.31797   50.50495    200   197.6988
384  100000    48.35802   50.50495    200   197.7418
385  100000    48.40035   50.50495    200   197.7871
386  100000    48.43659   50.50495    200   197.8258
387  100000    48.47613   50.50495    200   197.8685
388  100000    48.51271   50.50495    200   197.9076
389  100000    48.55398   50.50495    200   197.9517
390  100000    48.59292   50.50495    200   197.9935
391  100000    48.62888   50.50495    200   198.0320
392  100000    48.65889   50.50495    200   198.0640
393  100000    48.69585   50.50495    200   198.1035
394  100000    48.73204   50.50495    200   198.1421
395  100000    48.77037   50.50495    200   198.1830
396  100000    48.80205   50.50495    200   198.2168
397  100000    48.83264   50.50495    200   198.2494
398  100000    48.87460   50.50495    200   198.2941
399  100000    48.90707   50.50495    200   198.3286
400  100000    48.93438   50.50495    200   198.3576

Diploid Model with selection (random number based on
relative fitness to determine who survives).  Mutation rate is      1e-08,
Recomb rate is ONE per sexual individual per generation
Starting values at generation   0 are; population size 100000,
with avg fitness of    1.22136.

gen  popSize    avgFit     maxFit maxAlleles avgAlleles
  0  100000     1.22136    1.67020     26    10.0082
  1  100000     1.22136    1.67020     26    10.0082
  2  100000     1.22593    1.63998     25    10.1958
  3  100000     1.23023    1.84617     31    10.3712
  4  100000     1.23510    1.70689     27    10.5691
  5  100000     1.24037    1.67277     26    10.7817
  6  100000     1.24554    1.77448     29    10.9902
  7  100000     1.25076    1.74035     28    11.1999
  8  100000     1.25616    1.73969     28    11.4157
  9  100000     1.26104    1.73902     28    11.6104
 10  100000     1.26698    1.77516     29    11.8460
 11  100000     1.27228    1.80997     30    12.0553
 12  100000     1.27806    1.84546     31    12.2823
 13  100000     1.28343    1.81067     30    12.4922
 14  100000     1.28976    1.84617     31    12.7376
 15  100000     1.29566    1.84688     31    12.9671
 16  100000     1.30239    1.88237     32    13.2282
 17  100000     1.30896    1.88237     32    13.4808
 18  100000     1.31578    1.84688     31    13.7401
 19  100000     1.32269    1.81067     30    14.0025
 20  100000     1.32923    1.92002     33    14.2507
 21  100000     1.33615    1.84759     31    14.5102
 22  100000     1.34252    1.88092     32    14.7479
 23  100000     1.35012    1.92075     33    15.0322
 24  100000     1.35772    1.92075     33    15.3130
 25  100000     1.36584    2.07909     37    15.6127
 26  100000     1.37256    1.95465     34    15.8589
 27  100000     1.38124    2.03910     36    16.1737
 28  100000     1.38916    2.11904     38    16.4623
 29  100000     1.39790    2.11741     38    16.7747
 30  100000     1.40617    2.03832     36    17.0722
 31  100000     1.41448    2.03754     36    17.3695
 32  100000     1.42345    2.07829     37    17.6848
 33  100000     1.43268    2.15976     39    18.0102
 34  100000     1.44165    2.11904     38    18.3235
 35  100000     1.45136    2.16225     39    18.6597
 36  100000     1.46091    2.11985     38    18.9897
 37  100000     1.47111    2.43317     45    19.3375
 38  100000     1.48189    2.24615     41    19.7040
 39  100000     1.49170    2.24960     41    20.0336
 40  100000     1.50371    2.29371     42    20.4365
 41  100000     1.51474    2.24960     41    20.8041
 42  100000     1.52522    2.38546     44    21.1503
 43  100000     1.53711    2.29460     42    21.5403
 44  100000     1.54838    2.38455     44    21.9091
 45  100000     1.56008    2.38638     44    22.2872
 46  100000     1.57219    2.38546     44    22.6761
 47  100000     1.58446    2.38546     44    23.0665
 48  100000     1.59723    2.58210     48    23.4685
 49  100000     1.61020    2.53245     47    23.8730
 50  100000     1.62349    2.58210     48    24.2844
 51  100000     1.63850    2.58012     48    24.7462
 52  100000     1.65201    2.68539     50    25.1607
 53  100000     1.66631    2.53050     47    25.5937
 54  100000     1.68082    2.63071     49    26.0301
 55  100000     1.69565    2.68229     50    26.4714
 56  100000     1.71115    2.79280     52    26.9298
 57  100000     1.72700    2.73593     51    27.3914
 58  100000     1.74263    2.84537     53    27.8440
 59  100000     1.75978    2.74015     51    28.3347
 60  100000     1.77784    2.74225     51    28.8502
 61  100000     1.79559    2.90787     54    29.3499
 62  100000     1.81397    2.90675     54    29.8626
 63  100000     1.83304    2.85085     53    30.3893
 64  100000     1.85232    3.02185     56    30.9137
 65  100000     1.87172    2.90563     54    31.4385
 66  100000     1.89109    2.90340     54    31.9590
 67  100000     1.91214    3.20805     59    32.5134
 68  100000     1.93322    3.14273     58    33.0674
 69  100000     1.95348    3.20805     59    33.5921
 70  100000     1.97451    3.40048     62    34.1285
 71  100000     1.99686    3.33894     61    34.6979
 72  100000     2.01987    3.14515     58    35.2755
 73  100000     2.04323    3.27221     60    35.8539
 74  100000     2.06640    3.33509     61    36.4222
 75  100000     2.09046    3.53514     64    37.0047
 76  100000     2.11531    3.39787     62    37.5998
 77  100000     2.14115    3.40179     62    38.2137
 78  100000     2.16657    3.53786     64    38.8129
 79  100000     2.19171    3.74864     67    39.3928
 80  100000     2.21965    3.60585     65    40.0273
 81  100000     2.24660    3.75441     67    40.6398
 82  100000     2.27498    3.98421     70    41.2698
 83  100000     2.30435    3.75152     67    41.9163
 84  100000     2.33347    3.97808     70    42.5526
 85  100000     2.36270    4.06233     71    43.1810
 86  100000     2.39417    4.06545     71    43.8507
 87  100000     2.42706    4.14198     72    44.5365
 88  100000     2.45975    4.14517     72    45.2101
 89  100000     2.49301    4.84181     80    45.8892
 90  100000     2.52678    4.22320     73    46.5684
 91  100000     2.56224    4.47825     76    47.2697
 92  100000     2.59831    4.47308     76    47.9756
 93  100000     2.63369    4.48514     76    48.6595
 94  100000     2.67242    4.75418     79    49.3993
 95  100000     2.71165    4.56430     77    50.1350
 96  100000     2.75149    4.65917     78    50.8736
 97  100000     2.79206    4.65917     78    51.6101
 98  100000     2.83581    5.03548     82    52.3953
 99  100000     2.87960    4.74870     79    53.1678
100  100000     2.92434    4.93295     81    53.9465
101  100000     2.96914    5.55958     87    54.7149
102  100000     3.01562    5.24093     84    55.4990
103  100000     3.06384    5.24295     84    56.3006
104  100000     3.11351    5.34164     85    57.1174
105  100000     3.16282    5.55530     87    57.9120
106  100000     3.21387    5.89307     90    58.7245
107  100000     3.26752    5.67732     88    59.5599
108  100000     3.32157    5.67513     88    60.3945
109  100000     3.37463    6.02250     91    61.1958
110  100000     3.43051    5.89533     90    62.0300
111  100000     3.49032    5.89080     90    62.9004
112  100000     3.54724    6.48893     95    63.7186
113  100000     3.60615    6.25377     93    64.5561
114  100000     3.66628    6.38130     94    65.3920
115  100000     3.72883    6.50892     95    66.2495
116  100000     3.79103    6.51143     95    67.0891
117  100000     3.85612    6.63145     96    67.9519
118  100000     3.92315    7.31040    101    68.8272
119  100000     3.99061    6.90732     98    69.6917
120  100000     4.06190    7.31040    101    70.5957
121  100000     4.12827    7.17809    100    71.4169
122  100000     4.20200    7.74593    104    72.3128
123  100000     4.27317    7.76383    104    73.1685
124  100000     4.35001    7.61159    103    74.0732
125  100000     4.42579    7.76084    104    74.9488
126  100000     4.49980    7.75786    104    75.7926
127  100000     4.58163    8.39735    108    76.7055
128  100000     4.66292    8.22953    107    77.6023
129  100000     4.74569    8.22637    107    78.4988
130  100000     4.83195    8.71982    110    79.4093
131  100000     4.91772    8.90449    111    80.3089
132  100000     5.00742    9.06513    112    81.2219
133  100000     5.10087    9.26423    113    82.1539
134  100000     5.19084    8.72989    110    83.0480
135  100000     5.28769    9.64592    115    83.9912
136  100000     5.38395    9.26067    113    84.9096
137  100000     5.48437    9.62739    115    85.8472
138  100000     5.58280    9.81616    116    86.7545
139  100000     5.68666    9.81994    116    87.6913
140  100000     5.79103   10.20881    118    88.6191
141  100000     5.89721   10.40498    119    89.5421
142  100000     6.00599   10.82118    121    90.4743
143  100000     6.11687   11.24970    123    91.4058
144  100000     6.23037   11.02912    122    92.3384
145  100000     6.34095   11.05034    122    93.2373
146  100000     6.45986   11.04185    122    94.1839
147  100000     6.58514   11.05885    122    95.1603
148  100000     6.70498   11.48794    124    96.0760
149  100000     6.83265   12.17236    127    97.0407
150  100000     6.95791   12.43013    128    97.9636
151  100000     7.08339   11.93827    126    98.8788
152  100000     7.21997   12.66412    129    99.8524
153  100000     7.34680   12.65925    129   100.7433
154  100000     7.48522   13.97684    134   101.6939
155  100000     7.62198   14.24541    135   102.6210
156  100000     7.76047   13.17575    131   103.5360
157  100000     7.90445   14.23446    135   104.4734
158  100000     8.04887   14.80384    137   105.3986
159  100000     8.19938   15.10572    138   106.3393
160  100000     8.35784   14.52473    136   107.3156
161  100000     8.50749   14.51357    136   108.2200
162  100000     8.67058   14.82093    137   109.1917
163  100000     8.84178   16.03648    141   110.1955
164  100000     9.00348   15.69787    140   111.1225
165  100000     9.17631   16.31324    142   112.0955
166  100000     9.34486   17.30508    145   113.0256
167  100000     9.50969   16.62032    143   113.9236
168  100000     9.67726   17.28513    145   114.8199
169  100000     9.84382   16.33207    142   115.6962
170  100000    10.01264   17.32506    145   116.5627
171  100000    10.18669   18.33607    148   117.4437
172  100000    10.37579   18.01806    147   118.3813
173  100000    10.56649   17.96272    147   119.3154
174  100000    10.75780   18.32902    148   120.2360
175  100000    10.94513   18.70998    149   121.1182
176  100000    11.13279   19.81704    152   121.9903
177  100000    11.32418   19.08418    150   122.8572
178  100000    11.52515   19.45838    151   123.7637
179  100000    11.72589   19.82466    152   124.6491
180  100000    11.92587   21.01384    155   125.5124
181  100000    12.12666   20.24450    153   126.3635
182  100000    12.32462   20.22894    153   127.1933
183  100000    12.53640   21.89645    157   128.0678
184  100000    12.74681   22.27434    158   128.9167
185  100000    12.96756   22.30863    158   129.8015
186  100000    13.19219   21.87120    157   130.6827
187  100000    13.42236   25.04597    164   131.5679
188  100000    13.65315   23.20097    160   132.4425
189  100000    13.88086   23.19205    160   133.2864
190  100000    14.11082   23.19205    160   134.1348
191  100000    14.33572   23.19205    160   134.9495
192  100000    14.56377   25.06524    164   135.7585
193  100000    14.80199   24.11046    162   136.5893
194  100000    15.03552   24.55487    163   137.3899
195  100000    15.28136   25.56655    165   138.2290
196  100000    15.51547   26.55855    167   139.0081
197  100000    15.75483   27.63152    169   139.7939
198  100000    15.99950   27.11056    168   140.5872
199  100000    16.25048   27.66341    169   141.3818
200  100000    16.51118   27.64214    169   142.2000
201  100000    16.75429   28.20583    170   142.9494
202  100000    17.00259   28.17331    170   143.7068
203  100000    17.26677   28.19499    170   144.5060
204  100000    17.52239   29.31151    172   145.2591
205  100000    17.77748   28.71468    171   146.0003
206  100000    18.04325   30.48397    174   146.7645
207  100000    18.31937   28.74783    171   147.5477
208  100000    18.57257   30.48397    174   148.2532
209  100000    18.82000   29.30024    172   148.9323
210  100000    19.07939   29.85179    173   149.6386
211  100000    19.33667   31.06975    175   150.3323
212  100000    19.60766   31.67896    176   151.0433
213  100000    19.86866   31.08170    175   151.7262
214  100000    20.13689   32.30012    177   152.4159
215  100000    20.42610   32.30012    177   153.1505
216  100000    20.70862   31.69114    176   153.8581
217  100000    20.97978   32.92079    178   154.5293
218  100000    21.24587   34.23762    180   155.1794
219  100000    21.53011   34.90895    181   155.8632
220  100000    21.80465   33.56629    179   156.5162
221  100000    22.07568   34.25079    180   157.1537
222  100000    22.35428   35.60713    182   157.8016
223  100000    22.61379   34.92237    181   158.3957
224  100000    22.89940   36.26345    183   159.0398
225  100000    23.16934   37.01717    184   159.6492
226  100000    23.44829   36.30530    183   160.2654
227  100000    23.73173   35.59344    182   160.8836
228  100000    24.00270   36.29135    183   161.4725
229  100000    24.26842   39.22254    187   162.0383
230  100000    24.56582   37.74300    185   162.6675
231  100000    24.85261   37.01717    184   163.2650
232  100000    25.12064   36.30530    183   163.8191
233  100000    25.39677   37.01717    184   164.3835
234  100000    25.68814   37.72849    185   164.9718
235  100000    25.97680   38.46826    186   165.5474
236  100000    26.24165   40.00699    188   166.0727
237  100000    26.51220   39.23763    187   166.6026
238  100000    26.77854   39.99161    188   167.1187
239  100000    27.06006   40.00699    188   167.6608
240  100000    27.31469   40.77576    189   168.1437
241  100000    27.58023   40.00699    188   168.6441
242  100000    27.86589   41.59128    190   169.1761
243  100000    28.13691   42.40679    191   169.6735
244  100000    28.39237   42.40679    191   170.1410
245  100000    28.66590   41.59128    190   170.6371
246  100000    28.93100   41.59128    190   171.1147
247  100000    29.18663   41.59128    190   171.5672
248  100000    29.45251   41.59128    190   172.0364
249  100000    29.72249   42.40679    191   172.5074
250  100000    29.96111   43.23830    192   172.9208
251  100000    30.23107   43.23830    192   173.3849
252  100000    30.48432   44.08611    193   173.8187
253  100000    30.73922   42.40679    191   174.2491
254  100000    30.99604   44.95054    194   174.6803
255  100000    31.25454   43.23830    192   175.1081
256  100000    31.48964   44.08611    193   175.4942
257  100000    31.74153   44.93326    194   175.9060
258  100000    31.98800   44.08611    193   176.3063
259  100000    32.24375   44.08611    193   176.7185
260  100000    32.48949   44.95054    194   177.1119
261  100000    32.73682   45.83192    195   177.5051
262  100000    32.98100   44.95054    194   177.8898
263  100000    33.23406   44.95054    194   178.2869
264  100000    33.47173   45.83192    195   178.6568
265  100000    33.69806   45.83192    195   179.0049
266  100000    33.91863   45.83192    195   179.3424
267  100000    34.14031   45.83192    195   179.6789
268  100000    34.39408   45.83192    195   180.0618
269  100000    34.63347   47.64688    197   180.4219
270  100000    34.87140   46.73059    196   180.7777
271  100000    35.09085   47.64688    197   181.1020
272  100000    35.31744   47.64688    197   181.4341
273  100000    35.54842   46.73059    196   181.7730
274  100000    35.75356   46.73059    196   182.0717
275  100000    35.95458   47.64688    197   182.3618
276  100000    36.15731   47.64688    197   182.6526
277  100000    36.36690   49.53370    199   182.9518
278  100000    36.57029   48.58113    198   183.2407
279  100000    36.77757   48.58113    198   183.5347
280  100000    36.98230   48.58113    198   183.8229
281  100000    37.20390   47.64688    197   184.1316
282  100000    37.40572   49.53370    199   184.4117
283  100000    37.62233   48.58113    198   184.7097
284  100000    37.81973   48.58113    198   184.9813
285  100000    38.02546   48.58113    198   185.2628
286  100000    38.22906   48.58113    198   185.5393
287  100000    38.42152   48.58113    198   185.8002
288  100000    38.60801   49.53370    199   186.0529
289  100000    38.78482   48.58113    198   186.2887
290  100000    38.96338   48.58113    198   186.5262
291  100000    39.14149   49.53370    199   186.7631
292  100000    39.31429   49.53370    199   186.9928
293  100000    39.48123   50.50495    200   187.2120
294  100000    39.66747   49.53370    199   187.4564
295  100000    39.83364   49.53370    199   187.6731
296  100000    40.00785   49.53370    199   187.8990
297  100000    40.18272   50.50495    200   188.1262
298  100000    40.33040   49.53370    199   188.3162
299  100000    40.50176   49.53370    199   188.5366
300  100000    40.67195   50.50495    200   188.7525
301  100000    40.83402   50.50495    200   188.9591
302  100000    40.98671   50.50495    200   189.1534
303  100000    41.12949   50.50495    200   189.3342
304  100000    41.27706   49.53370    199   189.5195
305  100000    41.42491   50.50495    200   189.7045
306  100000    41.57006   50.50495    200   189.8867
307  100000    41.70082   50.50495    200   190.0496
308  100000    41.84929   50.50495    200   190.2339
309  100000    41.99915   50.50495    200   190.4195
310  100000    42.15477   50.50495    200   190.6113
311  100000    42.27046   50.50495    200   190.7540
312  100000    42.41402   50.50495    200   190.9307
313  100000    42.54214   50.50495    200   191.0871
314  100000    42.67334   50.50495    200   191.2467
315  100000    42.79783   50.50495    200   191.3975
316  100000    42.92800   50.50495    200   191.5554
317  100000    43.05772   50.50495    200   191.7121
318  100000    43.17916   50.50495    200   191.8584
319  100000    43.30719   50.50495    200   192.0122
320  100000    43.44103   50.50495    200   192.1721
321  100000    43.55620   50.50495    200   192.3093
322  100000    43.66699   50.50495    200   192.4408
323  100000    43.78569   50.50495    200   192.5815
324  100000    43.90233   50.50495    200   192.7200
325  100000    44.01707   50.50495    200   192.8552
326  100000    44.11609   50.50495    200   192.9719
327  100000    44.23091   50.50495    200   193.1071
328  100000    44.34674   50.50495    200   193.2426
329  100000    44.45699   50.50495    200   193.3718
330  100000    44.56433   50.50495    200   193.4965
331  100000    44.67757   50.50495    200   193.6282
332  100000    44.77866   50.50495    200   193.7459
333  100000    44.86706   50.50495    200   193.8481
334  100000    44.97063   50.50495    200   193.9680
335  100000    45.07017   50.50495    200   194.0826
336  100000    45.16527   50.50495    200   194.1921
337  100000    45.26783   50.50495    200   194.3107
338  100000    45.35704   50.50495    200   194.4128
339  100000    45.44698   50.50495    200   194.5158
340  100000    45.53707   50.50495    200   194.6188
341  100000    45.62432   50.50495    200   194.7179
342  100000    45.69914   50.50495    200   194.8031
343  100000    45.78242   50.50495    200   194.8973
344  100000    45.86759   50.50495    200   194.9940
345  100000    45.95643   50.50495    200   195.0944
346  100000    46.02997   50.50495    200   195.1776
347  100000    46.11877   50.50495    200   195.2778
348  100000    46.18927   50.50495    200   195.3571
349  100000    46.26711   50.50495    200   195.4447
350  100000    46.34374   50.50495    200   195.5303
351  100000    46.41842   50.50495    200   195.6142
352  100000    46.49111   50.50495    200   195.6953
353  100000    46.56931   50.50495    200   195.7828
354  100000    46.63208   50.50495    200   195.8527
355  100000    46.70920   50.50495    200   195.9389
356  100000    46.77346   50.50495    200   196.0104
357  100000    46.84770   50.50495    200   196.0927
358  100000    46.91546   50.50495    200   196.1680
359  100000    46.97265   50.50495    200   196.2309
360  100000    47.04211   50.50495    200   196.3077
361  100000    47.11137   50.50495    200   196.3838
362  100000    47.17600   50.50495    200   196.4554
363  100000    47.23405   50.50495    200   196.5193
364  100000    47.29867   50.50495    200   196.5907
365  100000    47.35159   50.50495    200   196.6485
366  100000    47.41375   50.50495    200   196.7168
367  100000    47.47175   50.50495    200   196.7804
368  100000    47.52760   50.50495    200   196.8413
369  100000    47.58441   50.50495    200   196.9033
370  100000    47.63283   50.50495    200   196.9563
371  100000    47.68601   50.50495    200   197.0145
372  100000    47.73116   50.50495    200   197.0638
373  100000    47.78752   50.50495    200   197.1250
374  100000    47.84109   50.50495    200   197.1831
375  100000    47.89096   50.50495    200   197.2373
376  100000    47.93665   50.50495    200   197.2871
377  100000    47.98730   50.50495    200   197.3417
378  100000    48.03362   50.50495    200   197.3919
379  100000    48.07791   50.50495    200   197.4398
380  100000    48.12217   50.50495    200   197.4875
381  100000    48.17047   50.50495    200   197.5397
382  100000    48.20739   50.50495    200   197.5798
383  100000    48.24758   50.50495    200   197.6230
384  100000    48.29062   50.50495    200   197.6693
385  100000    48.33234   50.50495    200   197.7141
386  100000    48.37489   50.50495    200   197.7598
387  100000    48.41902   50.50495    200   197.8071
388  100000    48.44683   50.50495    200   197.8371
389  100000    48.48292   50.50495    200   197.8759
390  100000    48.52003   50.50495    200   197.9153
391  100000    48.54929   50.50495    200   197.9469
392  100000    48.58959   50.50495    200   197.9899
393  100000    48.63049   50.50495    200   198.0337
394  100000    48.66231   50.50495    200   198.0678
395  100000    48.68757   50.50495    200   198.0948
396  100000    48.72190   50.50495    200   198.1315
397  100000    48.76024   50.50495    200   198.1723
398  100000    48.79139   50.50495    200   198.2054
399  100000    48.82114   50.50495    200   198.2371
400  100000    48.85349   50.50495    200   198.2716

Diploid Model with selection (random number based on
relative fitness to determine who survives).  Mutation rate is      1e-08,
Recomb rate is ONE per sexual individual per generation
Starting values at generation   0 are; population size 100000,
with avg fitness of    1.22090.

gen  popSize    avgFit     maxFit maxAlleles avgAlleles
  0  100000     1.22090    1.67149     26     9.9886
  1  100000     1.22090    1.67149     26     9.9886
  2  100000     1.22543    1.74035     28    10.1749
  3  100000     1.23029    1.67277     26    10.3728
  4  100000     1.23506    1.70689     27    10.5670
  5  100000     1.23976    1.74035     28    10.7575
  6  100000     1.24476    1.74035     28    10.9592
  7  100000     1.24978    1.70623     27    11.1612
  8  100000     1.25507    1.70557     27    11.3722
  9  100000     1.26043    1.70557     27    11.5865
 10  100000     1.26577    1.77243     29    11.7989
 11  100000     1.27141    1.80997     30    12.0219
 12  100000     1.27714    1.77312     29    12.2470
 13  100000     1.28305    1.77516     29    12.4793
 14  100000     1.28931    1.74102     28    12.7230
 15  100000     1.29545    1.77516     29    12.9602
 16  100000     1.30168    1.91928     33    13.2004
 17  100000     1.30812    1.84475     31    13.4482
 18  100000     1.31465    1.88092     32    13.6962
 19  100000     1.32183    1.91854     33    13.9695
 20  100000     1.32911    1.95766     34    14.2451
 21  100000     1.33566    1.95842     34    14.4928
 22  100000     1.34289    1.88309     32    14.7639
 23  100000     1.35003    1.88237     32    15.0291
 24  100000     1.35722    1.95992     34    15.2959
 25  100000     1.36476    1.95842     34    15.5750
 26  100000     1.37317    1.95992     34    15.8830
 27  100000     1.38177    2.03754     36    16.1954
 28  100000     1.38969    1.99835     35    16.4818
 29  100000     1.39812    2.07989     37    16.7842
 30  100000     1.40686    2.11985     38    17.0985
 31  100000     1.41551    2.03754     36    17.4054
 32  100000     1.42424    2.07829     37    17.7136
 33  100000     1.43354    2.03832     36    18.0402
 34  100000     1.44169    2.03989     36    18.3245
 35  100000     1.45128    2.20295     40    18.6573
 36  100000     1.46104    2.16059     39    18.9941
 37  100000     1.47110    2.16225     39    19.3371
 38  100000     1.48079    2.20634     40    19.6669
 39  100000     1.49184    2.24701     41    20.0385
 40  100000     1.50382    2.29371     42    20.4416
 41  100000     1.51493    2.29371     42    20.8120
 42  100000     1.52589    2.33869     43    21.1724
 43  100000     1.53747    2.29371     42    21.5521
 44  100000     1.54865    2.38546     44    21.9169
 45  100000     1.56095    2.33959     43    22.3128
 46  100000     1.57363    2.58409     48    22.7200
 47  100000     1.58629    2.43224     45    23.1251
 48  100000     1.59933    2.48088     46    23.5351
 49  100000     1.61235    2.48279     46    23.9413
 50  100000     1.62681    2.43317     45    24.3885
 51  100000     1.64060    2.52758     47    24.8135
 52  100000     1.65505    2.48374     46    25.2556
 53  100000     1.66936    2.84647     53    25.6849
 54  100000     1.68426    2.58409     48    26.1308
 55  100000     1.70053    2.63273     49    26.6154
 56  100000     1.71588    2.74015     51    27.0683
 57  100000     1.73059    2.73383     51    27.4963
 58  100000     1.74649    2.85085     53    27.9558
 59  100000     1.76292    2.74015     51    28.4260
 60  100000     1.78001    2.73909     51    28.9149
 61  100000     1.79733    2.79387     52    29.4026
 62  100000     1.81600    2.95464     55    29.9235
 63  100000     1.83369    2.90563     54    30.4138
 64  100000     1.85100    2.96260     55    30.8838
 65  100000     1.86755    2.90898     54    31.3322
 66  100000     1.88737    2.96716     55    31.8615
 67  100000     1.90743    3.07874     57    32.3929
 68  100000     1.92745    3.14515     58    32.9187
 69  100000     1.94927    3.14515     58    33.4828
 70  100000     1.97179    3.26970     60    34.0632
 71  100000     1.99266    3.39918     62    34.5904
 72  100000     2.01461    3.20312     59    35.1415
 73  100000     2.03850    3.46983     63    35.7356
 74  100000     2.06155    3.47383     63    36.3032
 75  100000     2.08632    3.60862     65    36.9050
 76  100000     2.11237    3.61140     65    37.5300
 77  100000     2.13779    3.47383     63    38.1349
 78  100000     2.16425    3.46716     63    38.7561
 79  100000     2.19250    3.53786     64    39.4076
 80  100000     2.21992    3.98114     70    40.0357
 81  100000     2.24754    3.75296     67    40.6563
 82  100000     2.27487    3.75152     67    41.2651
 83  100000     2.30344    3.91059     69    41.8954
 84  100000     2.33208    3.75585     67    42.5199
 85  100000     2.36356    3.98421     70    43.1975
 86  100000     2.39639    4.21833     73    43.8909
 87  100000     2.42624    4.13562     72    44.5187
 88  100000     2.45780    4.22482     73    45.1697
 89  100000     2.49151    4.21995     73    45.8563
 90  100000     2.52653    4.14198     72    46.5631
 91  100000     2.56367    4.47825     76    47.2982
 92  100000     2.59836    4.39044     75    47.9765
 93  100000     2.63393    4.30601     74    48.6624
 94  100000     2.67044    4.47825     76    49.3571
 95  100000     2.70908    4.56957     77    50.0841
 96  100000     2.74958    4.66634     78    50.8350
 97  100000     2.78990    4.75235     79    51.5701
 98  100000     2.83081    4.75601     79    52.3035
 99  100000     2.87187    4.74687     79    53.0317
100  100000     2.91507    5.66859     88    53.7874
101  100000     2.95830    5.33753     85    54.5314
102  100000     3.00353    5.14014     83    55.2989
103  100000     3.04983    5.34575     85    56.0733
104  100000     3.09688    5.57242     87    56.8456
105  100000     3.14737    5.55530     87    57.6659
106  100000     3.19726    5.45057     86    58.4627
107  100000     3.24683    5.78641     89    59.2385
108  100000     3.29677    5.78196     89    60.0127
109  100000     3.34832    6.25377     93    60.7999
110  100000     3.40295    5.89080     90    61.6173
111  100000     3.45960    6.13822     92    62.4496
112  100000     3.52042    6.50642     95    63.3331
113  100000     3.57831    6.37149     94    64.1586
114  100000     3.63643    6.76148     97    64.9730
115  100000     3.69731    6.25137     93    65.8180
116  100000     3.75476    6.89140     98    66.5968
117  100000     3.81979    6.76408     97    67.4639
118  100000     3.88643    6.90201     98    68.3401
119  100000     3.95197    7.31603    101    69.1860
120  100000     4.02470    6.89936     98    70.1120
121  100000     4.09415    7.32447    101    70.9776
122  100000     4.16694    7.61452    103    71.8718
123  100000     4.24178    7.90997    105    72.7723
124  100000     4.31958    7.47096    102    73.6967
125  100000     4.39449    7.46809    102    74.5721
126  100000     4.47687    8.22953    107    75.5112
127  100000     4.55341    8.23270    107    76.3723
128  100000     4.63648    8.37478    108    77.2916
129  100000     4.72212    9.07559    112    78.2214
130  100000     4.80967    8.71982    110    79.1544
131  100000     4.89571    8.91477    111    80.0558
132  100000     4.98852    8.74333    110    81.0138
133  100000     5.08415    9.08957    112    81.9737
134  100000     5.18345    9.61629    115    82.9557
135  100000     5.28049    9.44225    114    83.8959
136  100000     5.38113    9.80485    116    84.8500
137  100000     5.48716   10.21666    118    85.8499
138  100000     5.59391   10.00479    117    86.8275
139  100000     5.69608   10.62125    120    87.7465
140  100000     5.81098   10.20096    118    88.7598
141  100000     5.92649   10.82951    121    89.7673
142  100000     6.03612   11.47028    124    90.6994
143  100000     6.14892   11.06310    122    91.6443
144  100000     6.26577   11.49236    124    92.6003
145  100000     6.38578   11.47469    124    93.5723
146  100000     6.50444   11.69069    125    94.5100
147  100000     6.62910   11.93368    126    95.4787
148  100000     6.75132   12.91740    130    96.4023
149  100000     6.88570   11.70869    125    97.4075
150  100000     7.01550   12.41103    128    98.3644
151  100000     7.14861   12.92237    130    99.3280
152  100000     7.28209   12.40626    128   100.2650
153  100000     7.42127   13.18589    131   101.2336
154  100000     7.56473   15.37824    139   102.2112
155   98352     7.71217   13.96609    134   103.1920
156  100000     7.86900   14.21805    135   104.2212
157  100000     8.02225   13.70278    133   105.2039
158  100000     8.17615   14.24541    135   106.1770
159  100000     8.33763   15.98722    141   107.1721
160  100000     8.49279   16.01799    141   108.1169
161  100000     8.65408   15.39599    139   109.0748
162  100000     8.81854   16.00568    141   110.0395
163  100000     8.98947   15.40191    139   111.0216
164  100000     9.14752   15.70391    140   111.9133
165  100000     9.32133   16.32579    142   112.8738
166  100000     9.50197   16.94621    144   113.8581
167  100000     9.67785   17.30508    145   114.7917
168  100000     9.85871   17.56992    146   115.7449
169  100000    10.04389   17.65797    146   116.6943
170  100000    10.23008   18.32197    148   117.6341
171  100000    10.41724   18.69560    149   118.5593
172  100000    10.60611   18.36429    148   119.4848
173  100000    10.80365   19.09886    150   120.4242
174  100000    10.99724   19.85518    152   121.3339
175  100000    11.20364   21.04618    155   122.2960
176  100000    11.39609   20.19008    153   123.1710
177  100000    11.60730   21.45060    156   124.1163
178  100000    11.80495   19.84755    152   124.9763
179  100000    12.02077   19.85518    152   125.9087
180  100000    12.22482   20.63351    154   126.7684
181  100000    12.43741   20.62558    154   127.6493
182  100000    12.64727   21.46711    156   128.5094
183  100000    12.86205   21.45885    156   129.3743
184  100000    13.07278   22.29148    158   130.2097
185  100000    13.29977   21.87120    157   131.0920
186  100000    13.52450   23.20097    160   131.9475
187  100000    13.74201   24.58322    163   132.7664
188  100000    13.97841   23.20097    160   133.6427
189  100000    14.21247   23.66499    161   134.4939
190  100000    14.44772   25.57638    165   135.3309
191  100000    14.69102   23.68320    161   136.1927
192  100000    14.93582   25.06524    164   137.0438
193  100000    15.17682   25.07488    164   137.8640
194  100000    15.43039   25.07488    164   138.7160
195  100000    15.67760   25.09417    164   139.5336
196  100000    15.92468   26.06785    166   140.3394
197  100000    16.18063   26.58921    167   141.1537
198  100000    16.43096   26.60966    167   141.9469
199  100000    16.68571   27.63152    169   142.7354
200  100000    16.93890   27.11056    168   143.5108
201  100000    17.20526   28.75889    171   144.3165
202  100000    17.47286   28.75889    171   145.1064
203  100000    17.72960   30.46054    174   145.8572
204  100000    18.00499   31.05780    175   146.6518
205  100000    18.27143   29.32279    172   147.4065
206  100000    18.53249   29.30024    172   148.1401
207  100000    18.80471   30.47225    174   148.8858
208  100000    19.05937   29.88625    173   149.5809
209  100000    19.30418   30.46054    174   150.2376
210  100000    19.56771   31.67896    176   150.9340
211  100000    19.84294   32.31254    177   151.6521
212  100000    20.12750   31.03393    175   152.3868
213  100000    20.40730   34.92237    181   153.0962
214  100000    20.68711   32.90813    178   153.7963
215  100000    20.95442   32.30012    177   154.4625
216  100000    21.21547   32.90813    178   155.1006
217  100000    21.49566   32.92079    178   155.7752
218  100000    21.77592   34.90895    181   156.4439
219  100000    22.05795   33.59212    179   157.1082
220  100000    22.33014   36.30530    183   157.7393
221  100000    22.61922   34.90895    181   158.4033
222  100000    22.89461   34.90895    181   159.0302
223  100000    23.18547   37.00294    184   159.6847
224  100000    23.46126   36.29135    183   160.2940
225  100000    23.73335   34.92237    181   160.8883
226  100000    24.01575   36.29135    183   161.5001
227  100000    24.28487   37.01717    184   162.0730
228  100000    24.56994   36.30530    183   162.6753
229  100000    24.85887   38.48306    186   163.2778
230  100000    25.14370   37.72849    185   163.8673
231  100000    25.43330   37.74300    185   164.4585
232  100000    25.71400   38.48306    186   165.0274
233  100000    25.98492   38.48306    186   165.5675
234  100000    26.26723   39.22254    187   166.1281
235  100000    26.54739   38.48306    186   166.6742
236  100000    26.81755   39.22254    187   167.1953
237  100000    27.09274   40.79144    189   167.7236
238  100000    27.36057   40.79144    189   168.2318
239  100000    27.63200   40.00699    188   168.7416
240  100000    27.89803   41.57529    190   169.2356
241  100000    28.16907   40.00699    188   169.7347
242  100000    28.43924   41.59128    190   170.2296
243  100000    28.71399   43.23830    192   170.7273
244  100000    28.97714   40.79144    189   171.1998
245  100000    29.23741   42.40679    191   171.6614
246  100000    29.49120   42.40679    191   172.1086
247  100000    29.73997   44.08611    193   172.5420
248  100000    29.99113   41.59128    190   172.9766
249  100000    30.25856   43.22167    192   173.4363
250  100000    30.50845   44.06916    193   173.8632
251  100000    30.77709   43.23830    192   174.3145
252  100000    31.02953   44.08611    193   174.7363
253  100000    31.28545   44.08611    193   175.1614
254  100000    31.51293   43.23830    192   175.5362
255  100000    31.77419   44.95054    194   175.9624
256  100000    32.02752   44.95054    194   176.3726
257  100000    32.27190   44.08611    193   176.7653
258  100000    32.53058   44.08611    193   177.1802
259  100000    32.77468   44.95054    194   177.5665
260  100000    33.03264   44.95054    194   177.9748
261  100000    33.27982   46.73059    196   178.3605
262  100000    33.52121   45.83192    195   178.7324
263  100000    33.76093   46.73059    196   179.1029
264  100000    33.98800   45.81430    195   179.4494
265  100000    34.21711   46.73059    196   179.7970
266  100000    34.44549   46.73059    196   180.1412
267  100000    34.66665   45.83192    195   180.4714
268  100000    34.88389   46.73059    196   180.7959
269  100000    35.10566   46.73059    196   181.1233
270  100000    35.32766   46.73059    196   181.4498
271  100000    35.55545   46.73059    196   181.7822
272  100000    35.79237   46.73059    196   182.1273
273  100000    35.99639   47.64688    197   182.4214
274  100000    36.22354   47.64688    197   182.7472
275  100000    36.45074   47.64688    197   183.0718
276  100000    36.67109   47.64688    197   183.3845
277  100000    36.88849   47.64688    197   183.6912
278  100000    37.08310   49.53370    199   183.9635
279  100000    37.28250   49.53370    199   184.2416
280  100000    37.49044   49.53370    199   184.5299
281  100000    37.68187   48.58113    198   184.7938
282  100000    37.88909   50.50495    200   185.0775
283  100000    38.07554   48.58113    198   185.3334
284  100000    38.27221   49.53370    199   185.6003
285  100000    38.45697   49.53370    199   185.8487
286  100000    38.65101   48.58113    198   186.1108
287  100000    38.83045   49.53370    199   186.3501
288  100000    39.01706   49.53370    199   186.5984
289  100000    39.20187   48.58113    198   186.8439
290  100000    39.37306   49.53370    199   187.0705
291  100000    39.55166   49.53370    199   187.3057
292  100000    39.72649   49.53370    199   187.5337
293  100000    39.88679   49.53370    199   187.7431
294  100000    40.06218   50.50495    200   187.9701
295  100000    40.23708   49.53370    199   188.1970
296  100000    40.39968   50.50495    200   188.4063
297  100000    40.55097   50.50495    200   188.5997
298  100000    40.70908   49.53370    199   188.8013
299  100000    40.86893   50.50495    200   189.0047
300  100000    41.01209   49.53370    199   189.1863
301  100000    41.16696   50.50495    200   189.3811
302  100000    41.32728   50.50495    200   189.5836
303  100000    41.46902   50.50495    200   189.7610
304  100000    41.61209   50.50495    200   189.9400
305  100000    41.75514   50.50495    200   190.1180
306  100000    41.88235   50.50495    200   190.2754
307  100000    42.02557   50.50495    200   190.4521
308  100000    42.17750   50.50495    200   190.6397
309  100000    42.32213   50.50495    200   190.8176
310  100000    42.46619   50.50495    200   190.9943
311  100000    42.59724   50.50495    200   191.1549
312  100000    42.72592   50.50495    200   191.3114
313  100000    42.84774   50.50495    200   191.4580
314  100000    42.96427   50.50495    200   191.6000
315  100000    43.08951   50.50495    200   191.7501
316  100000    43.20897   50.50495    200   191.8936
317  100000    43.32773   50.50495    200   192.0364
318  100000    43.44139   50.50495    200   192.1723
319  100000    43.55185   50.50495    200   192.3043
320  100000    43.67153   50.50495    200   192.4463
321  100000    43.78895   50.50495    200   192.5857
322  100000    43.91260   50.50495    200   192.7325
323  100000    44.03173   50.50495    200   192.8732
324  100000    44.14288   50.50495    200   193.0039
325  100000    44.26072   50.50495    200   193.1429
326  100000    44.36368   50.50495    200   193.2638
327  100000    44.46712   50.50495    200   193.3843
328  100000    44.57362   50.50495    200   193.5080
329  100000    44.66961   50.50495    200   193.6196
330  100000    44.77529   50.50495    200   193.7425
331  100000    44.87164   50.50495    200   193.8540
332  100000    44.96350   50.50495    200   193.9605
333  100000    45.06705   50.50495    200   194.0798
334  100000    45.16415   50.50495    200   194.1917
335  100000    45.25089   50.50495    200   194.2914
336  100000    45.34209   50.50495    200   194.3959
337  100000    45.43957   50.50495    200   194.5078
338  100000    45.53382   50.50495    200   194.6155
339  100000    45.62383   50.50495    200   194.7179
340  100000    45.71622   50.50495    200   194.8227
341  100000    45.80423   50.50495    200   194.9226
342  100000    45.88798   50.50495    200   195.0178
343  100000    45.97056   50.50495    200   195.1113
344  100000    46.04516   50.50495    200   195.1955
345  100000    46.12001   50.50495    200   195.2797
346  100000    46.19687   50.50495    200   195.3664
347  100000    46.27979   50.50495    200   195.4594
348  100000    46.35703   50.50495    200   195.5462
349  100000    46.43654   50.50495    200   195.6350
350  100000    46.50147   50.50495    200   195.7075
351  100000    46.57358   50.50495    200   195.7883
352  100000    46.64745   50.50495    200   195.8705
353  100000    46.71048   50.50495    200   195.9406
354  100000    46.78077   50.50495    200   196.0189
355  100000    46.84324   50.50495    200   196.0881
356  100000    46.90921   50.50495    200   196.1612
357  100000    46.97330   50.50495    200   196.2323
358  100000    47.03095   50.50495    200   196.2958
359  100000    47.10169   50.50495    200   196.3738
360  100000    47.16606   50.50495    200   196.4447
361  100000    47.22233   50.50495    200   196.5067
362  100000    47.28324   50.50495    200   196.5737
363  100000    47.33998   50.50495    200   196.6359
364  100000    47.40107   50.50495    200   196.7030
365  100000    47.45345   50.50495    200   196.7603
366  100000    47.51196   50.50495    200   196.8244
367  100000    47.57820   50.50495    200   196.8968
368  100000    47.62810   50.50495    200   196.9511
369  100000    47.68302   50.50495    200   197.0111
370  100000    47.73441   50.50495    200   197.0672
371  100000    47.78293   50.50495    200   197.1200
372  100000    47.83486   50.50495    200   197.1764
373  100000    47.88965   50.50495    200   197.2360
374  100000    47.94184   50.50495    200   197.2924
375  100000    47.99550   50.50495    200   197.3507
376  100000    48.03233   50.50495    200   197.3904
377  100000    48.08444   50.50495    200   197.4466
378  100000    48.13910   50.50495    200   197.5057
379  100000    48.18171   50.50495    200   197.5519
380  100000    48.21950   50.50495    200   197.5925
381  100000    48.25962   50.50495    200   197.6355
382  100000    48.30662   50.50495    200   197.6863
383  100000    48.34887   50.50495    200   197.7316
384  100000    48.39111   50.50495    200   197.7770
385  100000    48.42985   50.50495    200   197.8187
386  100000    48.46771   50.50495    200   197.8593
387  100000    48.50468   50.50495    200   197.8989
388  100000    48.54012   50.50495    200   197.9369
389  100000    48.57120   50.50495    200   197.9701
390  100000    48.61130   50.50495    200   198.0132
391  100000    48.64538   50.50495    200   198.0495
392  100000    48.68022   50.50495    200   198.0868
393  100000    48.71413   50.50495    200   198.1228
394  100000    48.74825   50.50495    200   198.1591
395  100000    48.77862   50.50495    200   198.1916
396  100000    48.81191   50.50495    200   198.2271
397  100000    48.84125   50.50495    200   198.2583
398  100000    48.86924   50.50495    200   198.2881
399  100000    48.90110   50.50495    200   198.3219
400  100000    48.93176   50.50495    200   198.3546

Diploid Model with selection (random number based on
relative fitness to determine who survives).  Mutation rate is      1e-08,
Recomb rate is ONE per sexual individual per generation
Starting values at generation   0 are; population size 100000,
with avg fitness of    1.22133.

gen  popSize    avgFit     maxFit maxAlleles avgAlleles
  0  100000     1.22133    1.63998     25    10.0070
  1  100000     1.22133    1.63998     25    10.0070
  2  100000     1.22601    1.73768     28    10.1981
  3  100000     1.23048    1.67213     26    10.3809
  4  100000     1.23548    1.70557     27    10.5843
  5  100000     1.24043    1.67342     26    10.7846
  6  100000     1.24515    1.70689     27    10.9754
  7  100000     1.25046    1.70623     27    11.1888
  8  100000     1.25567    1.77448     29    11.3979
  9  100000     1.26160    1.74035     28    11.6333
 10  100000     1.26699    1.77516     29    11.8476
 11  100000     1.27224    1.73969     28    12.0547
 12  100000     1.27816    1.77312     29    12.2870
 13  100000     1.28446    1.80927     30    12.5332
 14  100000     1.29050    1.77516     29    12.7671
 15  100000     1.29689    1.88164     32    13.0150
 16  100000     1.30286    1.77312     29    13.2457
 17  100000     1.30923    1.80997     30    13.4905
 18  100000     1.31593    1.84617     31    13.7468
 19  100000     1.32260    1.95917     34    14.0005
 20  100000     1.32951    1.95917     34    14.2603
 21  100000     1.33673    1.92149     33    14.5313
 22  100000     1.34359    1.92002     33    14.7880
 23  100000     1.35106    1.96068     34    15.0671
 24  100000     1.35847    2.11985     38    15.3416
 25  100000     1.36606    2.03832     36    15.6190
 26  100000     1.37328    2.03832     36    15.8855
 27  100000     1.38122    2.16059     39    16.1757
 28  100000     1.38908    1.99989     35    16.4607
 29  100000     1.39753    1.99835     35    16.7643
 30  100000     1.40590    2.12067     38    17.0634
 31  100000     1.41470    2.11985     38    17.3769
 32  100000     1.42402    1.99835     35    17.7055
 33  100000     1.43380    2.20380     40    18.0501
 34  100000     1.44334    2.11904     38    18.3826
 35  100000     1.45225    2.12148     38    18.6932
 36  100000     1.46170    2.12230     38    19.0171
 37  100000     1.47137    2.12067     38    19.3471
 38  100000     1.48146    2.29107     42    19.6895
 39  100000     1.49282    2.16225     39    20.0745
 40  100000     1.50342    2.20465     40    20.4295
 41  100000     1.51467    2.29283     42    20.8043
 42  100000     1.52611    2.33959     43    21.1833
 43  100000     1.53750    2.34049     43    21.5544
 44  100000     1.54986    2.43411     45    21.9584
 45  100000     1.56136    2.42850     45    22.3316
 46  100000     1.57400    2.38455     44    22.7349
 47  100000     1.58579    2.33959     43    23.1093
 48  100000     1.59874    2.38546     44    23.5204
 49  100000     1.61150    2.58210     48    23.9183
 50  100000     1.62514    2.53245     47    24.3409
 51  100000     1.63975    2.58309     48    24.7918
 52  100000     1.65291    2.52953     47    25.1925
 53  100000     1.66692    2.73909     51    25.6173
 54  100000     1.68333    2.79280     52    26.1079
 55  100000     1.69843    2.53342     47    26.5572
 56  100000     1.71399    2.53342     47    27.0152
 57  100000     1.73043    2.68539     50    27.4957
 58  100000     1.74684    2.68745     50    27.9715
 59  100000     1.76422    2.90228     54    28.4691
 60  100000     1.78026    2.79280     52    28.9228
 61  100000     1.79854    2.84866     53    29.4363
 62  100000     1.81679    2.96146     55    29.9445
 63  100000     1.83515    3.02185     56    30.4492
 64  100000     1.85503    3.14394     58    30.9921
 65  100000     1.87479    3.14031     58    31.5276
 66  100000     1.89331    3.07874     57    32.0225
 67  100000     1.91321    3.08229     57    32.5476
 68  100000     1.93431    3.08229     57    33.0971
 69  100000     1.95549    3.02302     56    33.6435
 70  100000     1.97680    3.14394     58    34.1904
 71  100000     1.99844    3.53514     64    34.7365
 72  100000     2.02117    3.40310     62    35.3042
 73  100000     2.04481    3.40441     62    35.8925
 74  100000     2.06892    3.53786     64    36.4835
 75  100000     2.09386    3.54058     64    37.0844
 76  100000     2.11918    3.40310     62    37.6911
 77  100000     2.14499    3.47116     63    38.3002
 78  100000     2.17184    3.46983     63    38.9287
 79  100000     2.19981    3.61417     65    39.5739
 80  100000     2.22732    3.61001     65    40.1998
 81  100000     2.25557    3.82214     68    40.8333
 82  100000     2.28532    3.83097     68    41.4922
 83  100000     2.31618    3.90759     69    42.1697
 84  100000     2.34775    4.05141     71    42.8526
 85  100000     2.38111    3.82950     68    43.5664
 86  100000     2.41334    4.06233     71    44.2465
 87  100000     2.44510    4.21833     73    44.9073
 88  100000     2.48148    4.47480     76    45.6561
 89  100000     2.51603    4.22157     73    46.3515
 90  100000     2.55254    4.22320     73    47.0823
 91  100000     2.58962    4.56781     77    47.8047
 92  100000     2.62732    4.47997     76    48.5373
 93  100000     2.66528    4.75601     79    49.2623
 94  100000     2.70543    4.31263     74    50.0142
 95  100000     2.74244    4.65917     78    50.7013
 96  100000     2.78560    4.74870     79    51.4839
 97  100000     2.82780    4.66096     78    52.2455
 98  100000     2.87134    5.03936     82    53.0216
 99  100000     2.91678    4.84926     80    53.8159
100  100000     2.96095    5.14608     83    54.5750
101  100000     3.00702    5.65770     88    55.3547
102  100000     3.05540    5.66423     88    56.1604
103  100000     3.10432    5.56172     87    56.9657
104  100000     3.15603    5.66423     88    57.8004
105  100000     3.20698    5.34986     85    58.6101
106  100000     3.25910    5.78196     89    59.4304
107  100000     3.31273    5.67513     88    60.2546
108  100000     3.36716    5.66641     88    61.0771
109  100000     3.42176    6.00631     91    61.8907
110  100000     3.48104    5.88854     90    62.7599
111  100000     3.53927    6.76408     97    63.5960
112  100000     3.59803    6.38375     94    64.4307
113  100000     3.65988    6.51393     95    65.2958
114  100000     3.72225    7.17257    100    66.1501
115  100000     3.78913    6.51393     95    67.0533
116  100000     3.85514    6.63910     96    67.9296
117  100000     3.92461    6.88875     98    68.8344
118  100000     3.99527    7.01843     99    69.7377
119  100000     4.06667    7.62331    103    70.6349
120  100000     4.13938    7.75487    104    71.5349
121  100000     4.21177    7.45948    102    72.4137
122  100000     4.29051    7.61452    103    73.3522
123  100000     4.36951    7.60282    103    74.2752
124  100000     4.44773    7.75487    104    75.1795
125  100000     4.53244    8.07438    106    76.1393
126  100000     4.61700    8.07748    106    77.0757
127  100000     4.70123    8.40381    108    77.9961
128  100000     4.78683    8.55214    109    78.9124
129  100000     4.87550    8.38122    108    79.8500
130  100000     4.96954    9.62739    115    80.8257
131  100000     5.06334    9.44225    114    81.7726
132  100000     5.16464    9.43136    114    82.7778
133  100000     5.26422   10.22846    118    83.7527
134  100000     5.35997    9.82372    116    84.6691
135  100000     5.46007   10.60900    120    85.6030
136  100000     5.56577    9.63480    115    86.5853
137  100000     5.66938    9.82749    116    87.5198
138  100000     5.77653   11.04610    122    88.4721
139  100000     5.88500   10.42100    119    89.4193
140  100000     5.99774   11.02488    122    90.3889
141  100000     6.11558   11.04185    122    91.3806
142  100000     6.22737   11.48352    124    92.3016
143  100000     6.34711   11.25836    123    93.2715
144  100000     6.46446   11.71770    125    94.2038
145  100000     6.58612   11.69969    125    95.1552
146  100000     6.71350   11.69969    125    96.1351
147  100000     6.83855   11.71319    125    97.0735
148  100000     6.96407   12.40149    128    98.0018
149  100000     7.09134   12.91740    130    98.9259
150  100000     7.21788   13.17068    131    99.8286
151  100000     7.34722   12.64465    129   100.7336
152  100000     7.48666   13.15044    131   101.6952
153  100000     7.62733   12.93231    130   102.6458
154  100000     7.76490   13.17068    131   103.5574
155  100000     7.90797   13.95535    134   104.4823
156  100000     8.06159   13.96072    134   105.4720
157  100000     8.21675   15.10572    138   106.4524
158  100000     8.36944   15.70391    140   107.3909
159  100000     8.51936   14.52473    136   108.2996
160  100000     8.67282   15.38416    139   109.2082
161  100000     8.83898   15.68581    140   110.1767
162  100000     9.00631   15.38416    139   111.1400
163  100000     9.16840   15.70995    140   112.0537
164  100000     9.33692   16.01799    141   112.9852
165  100000     9.50827   16.65230    143   113.9144
166  100000     9.69598   17.99037    147   114.9102
167  100000     9.87462   16.98535    144   115.8412
168  100000    10.05610   17.65797    146   116.7734
169  100000    10.23990   16.99188    144   117.7011
170  100000    10.42075   17.97654    147   118.5995
171  100000    10.60756   18.36429    148   119.5078
172  100000    10.79325   18.36429    148   120.3999
173  100000    10.97607   19.47335    151   121.2644
174  100000    11.16415   19.42847    151   122.1310
175  100000    11.35482   19.86282    152   122.9960
176  100000    11.54930   20.23672    153   123.8647
177  100000    11.74181   19.47335    151   124.7131
178  100000    11.92658   20.64145    154   125.5126
179  100000    12.13137   20.62558    154   126.3900
180  100000    12.33658   21.03809    155   127.2464
181  100000    12.54498   21.45885    156   128.1021
182  100000    12.74313   21.02192    155   128.9066
183  100000    12.96309   21.43411    156   129.7868
184  100000    13.18639   23.19205    160   130.6654
185  100000    13.40021   23.65589    161   131.4893
186  100000    13.63015   23.20097    160   132.3617
187  100000    13.85978   23.64680    161   133.2217
188  100000    14.08550   23.65589    161   134.0476
189  100000    14.31035   23.65589    161   134.8631
190  100000    14.53723   25.07488    164   135.6725
191  100000    14.77317   24.11046    162   136.4975
192  100000    15.00589   25.08453    164   137.3005
193  100000    15.23988   25.09417    164   138.0908
194  100000    15.47417   25.08453    164   138.8754
195  100000    15.72186   27.11056    168   139.6953
196  100000    15.95123   26.06785    166   140.4417
197  100000    16.19838   27.66341    169   141.2328
198  100000    16.43859   26.56877    167   141.9895
199  100000    16.67510   28.73678    171   142.7218
200  100000    16.91242   28.70365    171   143.4487
201  100000    17.15109   29.25521    172   144.1689
202  100000    17.40540   28.19499    170   144.9229
203  100000    17.65343   29.28898    172   145.6481
204  100000    17.90875   28.74783    171   146.3857
205  100000    18.17094   30.46054    174   147.1350
206  100000    18.43421   30.48397    174   147.8759
207  100000    18.70019   29.27772    172   148.6147
208  100000    18.95451   29.87476    173   149.3098
209  100000    19.21801   30.47225    174   150.0188
210  100000    19.48453   32.28770    177   150.7295
211  100000    19.74187   31.69114    176   151.4055
212  100000    20.00334   31.66678    176   152.0826
213  100000    20.28454   32.31254    177   152.8017
214  100000    20.55459   31.67896    176   153.4814
215  100000    20.82589   32.92079    178   154.1552
216  100000    21.09406   32.93345    178   154.8160
217  100000    21.36097   33.56629    179   155.4634
218  100000    21.63173   32.31254    177   156.1127
219  100000    21.91095   36.29135    183   156.7757
220  100000    22.16791   33.57920    179   157.3756
221  100000    22.42441   34.90895    181   157.9706
222  100000    22.69247   35.60713    182   158.5817
223  100000    22.96760   34.92237    181   159.2031
224  100000    23.23995   34.90895    181   159.8126
225  100000    23.51065   35.59344    182   160.4084
226  100000    23.78586   37.74300    185   161.0088
227  100000    24.05509   35.60713    182   161.5890
228  100000    24.34300   38.46826    186   162.2026
229  100000    24.59584   37.01717    184   162.7322
230  100000    24.86629   38.45347    186   163.2977
231  100000    25.14399   37.72849    185   163.8726
232  100000    25.40868   38.48306    186   164.4138
233  100000    25.67239   40.00699    188   164.9459
234  100000    25.95007   40.00699    188   165.5011
235  100000    26.22673   37.74300    185   166.0492
236  100000    26.48341   39.23763    187   166.5518
237  100000    26.75578   40.00699    188   167.0789
238  100000    27.01495   39.23763    187   167.5792
239  100000    27.29278   40.00699    188   168.1078
240  100000    27.55723   39.23763    187   168.6027
241  100000    27.83509   40.79144    189   169.1230
242  100000    28.10270   40.79144    189   169.6172
243  100000    28.38192   41.59128    190   170.1272
244  100000    28.62854   42.40679    191   170.5723
245  100000    28.88308   40.77576    189   171.0286
246  100000    29.13814   42.40679    191   171.4835
247  100000    29.40577   43.23830    192   171.9572
248  100000    29.67204   42.40679    191   172.4225
249  100000    29.92250   42.40679    191   172.8569
250  100000    30.18355   42.40679    191   173.3038
251  100000    30.44644   44.08611    193   173.7545
252  100000    30.70645   42.40679    191   174.1942
253  100000    30.95399   43.23830    192   174.6091
254  100000    31.21380   44.08611    193   175.0434
255  100000    31.45748   44.08611    193   175.4449
256  100000    31.70875   43.23830    192   175.8578
257  100000    31.95167   44.93326    194   176.2529
258  100000    32.19000   45.83192    195   176.6384
259  100000    32.44304   44.95054    194   177.0427
260  100000    32.69312   44.08611    193   177.4404
261  100000    32.94480   45.83192    195   177.8371
262  100000    33.17310   46.73059    196   178.1954
263  100000    33.40413   44.95054    194   178.5542
264  100000    33.64577   45.83192    195   178.9258
265  100000    33.90071   45.83192    195   179.3177
266  100000    34.12415   45.83192    195   179.6574
267  100000    34.33866   45.83192    195   179.9810
268  100000    34.55253   46.73059    196   180.3022
269  100000    34.77704   46.73059    196   180.6375
270  100000    35.00732   46.73059    196   180.9795
271  100000    35.23558   46.73059    196   181.3168
272  100000    35.45397   47.64688    197   181.6382
273  100000    35.65134   47.64688    197   181.9249
274  100000    35.86852   47.64688    197   182.2399
275  100000    36.08415   47.64688    197   182.5502
276  100000    36.29591   47.64688    197   182.8522
277  100000    36.52402   48.58113    198   183.1775
278  100000    36.74645   47.64688    197   183.4925
279  100000    36.94105   47.64688    197   183.7661
280  100000    37.15643   48.58113    198   184.0674
281  100000    37.37373   47.64688    197   184.3698
282  100000    37.56290   47.64688    197   184.6323
283  100000    37.75386   49.53370    199   184.8939
284  100000    37.93981   48.58113    198   185.1487
285  100000    38.12073   49.53370    199   185.3947
286  100000    38.30889   49.53370    199   185.6516
287  100000    38.48084   48.58113    198   185.8836
288  100000    38.66579   47.64688    197   186.1326
289  100000    38.85962   48.58113    198   186.3903
290  100000    39.04077   48.58113    198   186.6313
291  100000    39.24389   49.53370    199   186.9005
292  100000    39.41734   49.53370    199   187.1293
293  100000    39.59075   49.53370    199   187.3563
294  100000    39.75938   49.53370    199   187.5775
295  100000    39.93836   49.53370    199   187.8106
296  100000    40.10698   49.53370    199   188.0290
297  100000    40.25427   49.53370    199   188.2191
298  100000    40.40634   50.50495    200   188.4141
299  100000    40.55830   50.50495    200   188.6082
300  100000    40.71764   49.53370    199   188.8119
301  100000    40.87427   50.50495    200   189.0103
302  100000    41.04084   50.50495    200   189.2222
303  100000    41.20213   50.50495    200   189.4255
304  100000    41.35472   50.50495    200   189.6179
305  100000    41.49946   50.50495    200   189.7990
306  100000    41.63949   49.53370    199   189.9736
307  100000    41.78678   50.50495    200   190.1576
308  100000    41.93064   50.50495    200   190.3358
309  100000    42.05801   50.50495    200   190.4927
310  100000    42.19256   50.50495    200   190.6588
311  100000    42.31950   50.50495    200   190.8145
312  100000    42.45412   50.50495    200   190.9791
313  100000    42.58785   50.50495    200   191.1426
314  100000    42.72258   50.50495    200   191.3073
315  100000    42.84996   50.50495    200   191.4621
316  100000    42.97932   50.50495    200   191.6185
317  100000    43.11434   50.50495    200   191.7810
318  100000    43.23450   50.50495    200   191.9256
319  100000    43.36004   50.50495    200   192.0758
320  100000    43.48777   50.50495    200   192.2287
321  100000    43.61755   50.50495    200   192.3828
322  100000    43.73637   50.50495    200   192.5245
323  100000    43.85899   50.50495    200   192.6692
324  100000    43.97389   50.50495    200   192.8046
325  100000    44.08972   50.50495    200   192.9413
326  100000    44.20148   50.50495    200   193.0729
327  100000    44.31431   50.50495    200   193.2054
328  100000    44.41645   50.50495    200   193.3247
329  100000    44.51453   50.50495    200   193.4390
330  100000    44.63000   50.50495    200   193.5734
331  100000    44.73182   50.50495    200   193.6920
332  100000    44.83566   50.50495    200   193.8128
333  100000    44.93313   50.50495    200   193.9250
334  100000    45.01143   50.50495    200   194.0156
335  100000    45.10490   50.50495    200   194.1232
336  100000    45.19587   50.50495    200   194.2281
337  100000    45.28774   50.50495    200   194.3334
338  100000    45.37186   50.50495    200   194.4300
339  100000    45.45487   50.50495    200   194.5248
340  100000    45.55140   50.50495    200   194.6349
341  100000    45.64296   50.50495    200   194.7392
342  100000    45.72879   50.50495    200   194.8368
343  100000    45.81498   50.50495    200   194.9348
344  100000    45.90090   50.50495    200   195.0319
345  100000    45.97590   50.50495    200   195.1167
346  100000    46.05215   50.50495    200   195.2029
347  100000    46.14217   50.50495    200   195.3043
348  100000    46.21801   50.50495    200   195.3895
349  100000    46.29791   50.50495    200   195.4797
350  100000    46.37278   50.50495    200   195.5635
351  100000    46.44571   50.50495    200   195.6451
352  100000    46.52476   50.50495    200   195.7335
353  100000    46.60369   50.50495    200   195.8215
354  100000    46.67071   50.50495    200   195.8962
355  100000    46.74848   50.50495    200   195.9824
356  100000    46.81842   50.50495    200   196.0606
357  100000    46.87078   50.50495    200   196.1188
358  100000    46.93963   50.50495    200   196.1951
359  100000    47.00498   50.50495    200   196.2669
360  100000    47.06598   50.50495    200   196.3344
361  100000    47.11734   50.50495    200   196.3909
362  100000    47.18257   50.50495    200   196.4627
363  100000    47.23902   50.50495    200   196.5249
364  100000    47.29149   50.50495    200   196.5828
365  100000    47.34860   50.50495    200   196.6451
366  100000    47.41556   50.50495    200   196.7188
367  100000    47.47364   50.50495    200   196.7823
368  100000    47.52678   50.50495    200   196.8405
369  100000    47.57548   50.50495    200   196.8937
370  100000    47.63147   50.50495    200   196.9547
371  100000    47.69016   50.50495    200   197.0189
372  100000    47.74609   50.50495    200   197.0798
373  100000    47.80245   50.50495    200   197.1412
374  100000    47.84299   50.50495    200   197.1851
375  100000    47.89767   50.50495    200   197.2444
376  100000    47.95213   50.50495    200   197.3034
377  100000    47.99792   50.50495    200   197.3530
378  100000    48.04567   50.50495    200   197.4050
379  100000    48.07999   50.50495    200   197.4419
380  100000    48.12567   50.50495    200   197.4914
381  100000    48.17038   50.50495    200   197.5397
382  100000    48.22453   50.50495    200   197.5981
383  100000    48.26585   50.50495    200   197.6426
384  100000    48.30528   50.50495    200   197.6849
385  100000    48.34949   50.50495    200   197.7326
386  100000    48.39155   50.50495    200   197.7776
387  100000    48.43241   50.50495    200   197.8217
388  100000    48.47340   50.50495    200   197.8658
389  100000    48.51194   50.50495    200   197.9070
390  100000    48.54846   50.50495    200   197.9459
391  100000    48.58455   50.50495    200   197.9845
392  100000    48.62203   50.50495    200   198.0246
393  100000    48.65446   50.50495    200   198.0594
394  100000    48.69250   50.50495    200   198.1001
395  100000    48.72502   50.50495    200   198.1348
396  100000    48.75607   50.50495    200   198.1677
397  100000    48.79486   50.50495    200   198.2092
398  100000    48.82739   50.50495    200   198.2437
399  100000    48.85365   50.50495    200   198.2717
400  100000    48.87820   50.50495    200   198.2978

Diploid Model with selection (random number based on
relative fitness to determine who survives).  Mutation rate is      1e-08,
Recomb rate is ONE per sexual individual per generation
Starting values at generation   0 are; population size 100000,
with avg fitness of    1.22124.

gen  popSize    avgFit     maxFit maxAlleles avgAlleles
  0  100000     1.22124    1.60782     24    10.0026
  1  100000     1.22124    1.60782     24    10.0026
  2  100000     1.22581    1.64061     25    10.1894
  3  100000     1.23068    1.73835     28    10.3882
  4  100000     1.23536    1.70689     27    10.5796
  5  100000     1.23972    1.80997     30    10.7567
  6  100000     1.24506    1.73835     28    10.9709
  7  100000     1.25039    1.73902     28    11.1852
  8  100000     1.25533    1.77448     29    11.3821
  9  100000     1.26059    1.74035     28    11.5921
 10  100000     1.26617    1.80997     30    11.8134
 11  100000     1.27157    1.74102     28    12.0264
 12  100000     1.27769    1.77448     29    12.2676
 13  100000     1.28352    1.77516     29    12.4961
 14  100000     1.28965    1.80858     30    12.7351
 15  100000     1.29630    1.91854     33    12.9926
 16  100000     1.30259    1.80927     30    13.2357
 17  100000     1.30898    1.81067     30    13.4806
 18  100000     1.31522    1.99758     35    13.7183
 19  100000     1.32184    1.88237     32    13.9701
 20  100000     1.32893    1.99758     35    14.2398
 21  100000     1.33625    1.92002     33    14.5153
 22  100000     1.34330    1.84759     31    14.7799
 23  100000     1.34979    1.95917     34    15.0216
 24  100000     1.35732    1.92075     33    15.3006
 25  100000     1.36437    1.92002     33    15.5603
 26  100000     1.37241    2.03754     36    15.8549
 27  100000     1.38083    2.03754     36    16.1624
 28  100000     1.38888    1.99451     35    16.4534
 29  100000     1.39753    2.03675     36    16.7650
 30  100000     1.40571    2.11659     38    17.0577
 31  100000     1.41448    2.07749     37    17.3698
 32  100000     1.42286    2.07829     37    17.6654
 33  100000     1.43195    2.11904     38    17.9860
 34  100000     1.44125    2.07669     37    18.3117
 35  100000     1.45028    2.11904     38    18.6246
 36  100000     1.45995    2.20210     40    18.9577
 37  100000     1.47008    2.20210     40    19.3056
 38  100000     1.47899    2.25047     41    19.6085
 39  100000     1.48848    2.24960     41    19.9285
 40  100000     1.49919    2.29283     42    20.2889
 41  100000     1.50980    2.20295     40    20.6426
 42  100000     1.52136    2.24960     41    21.0260
 43  100000     1.53201    2.29460     42    21.3743
 44  100000     1.54343    2.33959     43    21.7459
 45  100000     1.55563    2.29460     42    22.1429
 46  100000     1.56773    2.34049     43    22.5339
 47  100000     1.57980    2.38455     44    22.9182
 48  100000     1.59251    2.48183     46    23.3205
 49  100000     1.60524    2.38730     44    23.7204
 50  100000     1.61915    2.43224     45    24.1533
 51  100000     1.63272    2.48088     46    24.5721
 52  100000     1.64786    2.48470     46    25.0378
 53  100000     1.66194    2.73699     51    25.4669
 54  100000     1.67643    2.63273     49    25.9027
 55  100000     1.69159    2.63172     49    26.3552
 56  100000     1.70639    2.85085     53    26.7930
 57  100000     1.72290    2.79495     52    27.2759
 58  100000     1.73984    2.79387     52    27.7652
 59  100000     1.75681    2.68848     50    28.2537
 60  100000     1.77441    2.84756     53    28.7579
 61  100000     1.79075    2.84647     53    29.2210
 62  100000     1.80920    2.90675     54    29.7361
 63  100000     1.82769    2.79495     52    30.2479
 64  100000     1.84647    2.91010     54    30.7619
 65  100000     1.86514    3.02185     56    31.2696
 66  100000     1.88382    2.96830     55    31.7708
 67  100000     1.90333    3.08111     57    32.2893
 68  100000     1.92313    3.02534     56    32.8102
 69  100000     1.94518    3.14152     58    33.3863
 70  100000     1.96626    3.14515     58    33.9271
 71  100000     1.98750    3.20805     59    34.4671
 72  100000     2.01080    3.14636     58    35.0554
 73  100000     2.03228    3.27347     60    35.5904
 74  100000     2.05510    3.39787     62    36.1516
 75  100000     2.07884    3.60862     65    36.7311
 76  100000     2.10289    3.40179     62    37.3089
 77  100000     2.12838    3.47116     63    37.9172
 78  100000     2.15502    3.47383     63    38.5444
 79  100000     2.18099    3.61140     65    39.1502
 80  100000     2.20833    3.54058     64    39.7747
 81  100000     2.23561    3.66949     66    40.3925
 82  100000     2.26506    3.68362     66    41.0532
 83  100000     2.29460    3.82655     68    41.7096
 84  100000     2.32444    3.98421     70    42.3636
 85  100000     2.35436    3.98421     70    43.0087
 86  100000     2.38486    3.82950     68    43.6606
 87  100000     2.41597    3.90609     69    44.3134
 88  100000     2.44937    3.98268     70    45.0083
 89  100000     2.48184    4.22320     73    45.6730
 90  100000     2.51529    4.05609     71    46.3500
 91  100000     2.54999    4.30601     74    47.0429
 92  100000     2.58503    4.22482     73    47.7304
 93  100000     2.62045    4.47997     76    48.4160
 94  100000     2.65574    4.65738     78    49.0911
 95  100000     2.69315    4.57484     77    49.7987
 96  100000     2.73053    4.56606     77    50.4997
 97  100000     2.76876    4.56781     77    51.1969
 98  100000     2.80808    4.75418     79    51.9090
 99  100000     2.84893    5.14410     83    52.6397
100  100000     2.88962    4.66096     78    53.3572
101  100000     2.93280    5.03936     82    54.1079
102  100000     2.97822    4.85299     80    54.8823
103  100000     3.02081    4.94815     81    55.6000
104  100000     3.06556    5.55744     87    56.3402
105  100000     3.11237    5.23891     84    57.1062
106  100000     3.16160    5.66423     88    57.8960
107  100000     3.21038    5.57028     87    58.6784
108  100000     3.25905    5.56600     87    59.4434
109  100000     3.30854    5.90668     90    60.2024
110  100000     3.35966    6.25858     93    60.9788
111  100000     3.41174    5.90214     90    61.7623
112  100000     3.46636    6.37149     94    62.5659
113  100000     3.52279    6.13115     92    63.3828
114  100000     3.58125    6.26340     93    64.2173
115  100000     3.63975    6.37885     94    65.0356
116  100000     3.69761    6.62890     96    65.8311
117  100000     3.75739    6.77970     97    66.6453
118  100000     3.82210    6.90201     98    67.5108
119  100000     3.88884    6.76408     97    68.3852
120  100000     3.95503    7.31321    101    69.2436
121  100000     4.02366    6.89936     98    70.1166
122  100000     4.08965    7.30478    101    70.9397
123  100000     4.16343    7.32729    101    71.8415
124  100000     4.24067    7.61159    103    72.7777
125  100000     4.31440    7.32447    101    73.6533
126  100000     4.39199    7.62331    103    74.5597
127  100000     4.47094    7.76084    104    75.4599
128  100000     4.55458    7.61452    103    76.4048
129  100000     4.63741    8.57189    109    77.3199
130  100000     4.72142    8.07748    106    78.2299
131  100000     4.80592    8.72653    110    79.1347
132  100000     4.89568    9.26423    113    80.0767
133  100000     4.98682    8.57519    109    81.0131
134  100000     5.07375    8.91477    111    81.8909
135  100000     5.16352    9.25711    113    82.7765
136  100000     5.25822    9.07559    112    83.7010
137  100000     5.35756    9.61999    115    84.6513
138  100000     5.45602   10.02019    117    85.5802
139  100000     5.55861   10.42501    119    86.5299
140  100000     5.65882    9.81994    116    87.4407
141  100000     5.76657   10.20881    118    88.3979
142  100000     5.87186   10.22059    118    89.3207
143  100000     5.98078   10.83367    121    90.2518
144  100000     6.09268   10.63351    120    91.1981
145  100000     6.20479   11.23673    123    92.1283
146  100000     6.31511   11.27135    123    93.0235
147  100000     6.43135   11.49236    124    93.9550
148  100000     6.54146   11.50562    124    94.8209
149  100000     6.66514   11.69969    125    95.7776
150  100000     6.78488   11.92451    126    96.6836
151  100000     6.91208   11.69069    125    97.6302
152  100000     7.04166   12.65438    129    98.5781
153  100000     7.17011   12.92734    130    99.4982
154  100000     7.30217   12.67386    129   100.4315
155  100000     7.43597   12.90747    130   101.3611
156  100000     7.56828   13.16056    131   102.2558
157  100000     7.70098   13.42893    132   103.1459
158  100000     7.84395   13.68698    133   104.0819
159  100000     7.98822   13.93390    134   105.0122
160  100000     8.13771   14.22352    135   105.9563
161  100000     8.29112   14.23993    135   106.9071
162  100000     8.44139   14.22352    135   107.8282
163  100000     8.60000   15.99337    141   108.7817
164  100000     8.75727   15.39007    139   109.7073
165  100000     8.91089   15.41969    139   110.5898
166  100000     9.07587   15.39007    139   111.5326
167  100000     9.24571   16.03031    141   112.4799
168  100000     9.41329   16.00568    141   113.3978
169  100000     9.58501   16.63950    143   114.3248
170  100000     9.75850   17.63761    146   115.2358
171  100000     9.92993   17.65118    146   116.1273
172  100000    10.10817   17.31174    145   117.0383
173  100000    10.29159   17.32506    145   117.9591
174  100000    10.48210   18.72438    149   118.8939
175  100000    10.67062   17.97654    147   119.8145
176  100000    10.85178   19.44342    151   120.6766
177  100000    11.04119   19.84755    152   121.5569
178  100000    11.23693   19.43595    151   122.4566
179  100000    11.44274   19.47335    151   123.3878
180  100000    11.64500   19.45838    151   124.2857
181  100000    11.84230   20.60972    154   125.1437
182  100000    12.04105   21.06238    155   125.9972
183  100000    12.24306   21.43411    156   126.8520
184  100000    12.45383   20.63351    154   127.7281
185  100000    12.66316   21.85439    157   128.5812
186  100000    12.87787   21.02192    155   129.4455
187  100000    13.09687   21.87120    157   130.3110
188  100000    13.32235   22.75480    159   131.1878
189  100000    13.54345   23.63771    161   132.0334
190  100000    13.75964   23.67410    161   132.8418
191  100000    13.99347   24.11974    162   133.7107
192  100000    14.21684   24.11046    162   134.5231
193  100000    14.44786   25.56655    165   135.3490
194  100000    14.68197   27.07931    168   136.1743
195  100000    14.92254   24.59267    163   137.0126
196  100000    15.16617   25.57638    165   137.8440
197  100000    15.40061   24.57377    163   138.6312
198  100000    15.64011   26.08791    166   139.4250
199  100000    15.87761   25.56655    165   140.1962
200  100000    16.11442   26.57898    167   140.9607
201  100000    16.36676   26.56877    167   141.7590
202  100000    16.60420   27.63152    169   142.4955
203  100000    16.86225   27.64214    169   143.2905
204  100000    17.10113   28.18415    170   144.0134
205  100000    17.33819   28.75889    171   144.7207
206  100000    17.59472   27.65278    169   145.4745
207  100000    17.84461   29.87476    173   146.2009
208  100000    18.11312   29.30024    172   146.9653
209  100000    18.36789   29.30024    172   147.6840
210  100000    18.62177   31.66678    176   148.3905
211  100000    18.88865   31.04586    175   149.1259
212  100000    19.16191   29.87476    173   149.8659
213  100000    19.44162   31.66678    176   150.6127
214  100000    19.70367   31.67896    176   151.3009
215  100000    19.97470   30.47225    174   152.0035
216  100000    20.23480   32.30012    177   152.6704
217  100000    20.51596   32.90813    178   153.3786
218  100000    20.79096   32.31254    177   154.0624
219  100000    21.06412   32.92079    178   154.7348
220  100000    21.32141   33.56629    179   155.3594
221  100000    21.60896   34.23762    180   156.0535
222  100000    21.86963   34.21130    180   156.6728
223  100000    22.14660   34.25079    180   157.3177
224  100000    22.41837   34.25079    180   157.9473
225  100000    22.69631   34.90895    181   158.5841
226  100000    22.96748   34.90895    181   159.1957
227  100000    23.22622   34.92237    181   159.7702
228  100000    23.50301   37.00294    184   160.3835
229  100000    23.78956   36.29135    183   161.0097
230  100000    24.05633   38.48306    186   161.5852
231  100000    24.32618   37.01717    184   162.1602
232  100000    24.57766   37.01717    184   162.6908
233  100000    24.85512   37.01717    184   163.2729
234  100000    25.12595   37.72849    185   163.8335
235  100000    25.39424   37.74300    185   164.3828
236  100000    25.67191   40.00699    188   164.9447
237  100000    25.94411   39.23763    187   165.4862
238  100000    26.22266   38.48306    186   166.0373
239  100000    26.48831   39.97624    188   166.5572
240  100000    26.76926   40.00699    188   167.1026
241  100000    27.04368   39.23763    187   167.6298
242  100000    27.31720   39.23763    187   168.1505
243  100000    27.59119   40.79144    189   168.6651
244  100000    27.86291   40.77576    189   169.1730
245  100000    28.13484   40.77576    189   169.6757
246  100000    28.40571   40.00699    188   170.1699
247  100000    28.65760   41.59128    190   170.6255
248  100000    28.91022   42.39049    191   171.0790
249  100000    29.16533   40.79144    189   171.5313
250  100000    29.42922   43.23830    192   171.9976
251  100000    29.67847   43.23830    192   172.4340
252  100000    29.94337   42.40679    191   172.8933
253  100000    30.19390   43.23830    192   173.3237
254  100000    30.45703   44.08611    193   173.7743
255  100000    30.71568   43.23830    192   174.2123
256  100000    30.97617   44.08611    193   174.6480
257  100000    31.23520   44.08611    193   175.0788
258  100000    31.48277   44.06916    193   175.4874
259  100000    31.72028   44.08611    193   175.8745
260  100000    31.95445   44.95054    194   176.2557
261  100000    32.19122   44.08611    193   176.6387
262  100000    32.44474   44.08611    193   177.0436
263  100000    32.66513   44.95054    194   177.3941
264  100000    32.92849   44.95054    194   177.8089
265  100000    33.16500   44.95054    194   178.1796
266  100000    33.41101   44.95054    194   178.5640
267  100000    33.64431   46.73059    196   178.9231
268  100000    33.87687   46.73059    196   179.2810
269  100000    34.09934   46.73059    196   179.6195
270  100000    34.32644   47.64688    197   179.9621
271  100000    34.55700   46.71262    196   180.3070
272  100000    34.78951   46.73059    196   180.6573
273  100000    34.99311   46.73059    196   180.9581
274  100000    35.20884   47.64688    197   181.2767
275  100000    35.42203   46.73059    196   181.5903
276  100000    35.63182   46.73059    196   181.8962
277  100000    35.83840   47.64688    197   182.1967
278  100000    36.04899   47.64688    197   182.4977
279  100000    36.25755   47.64688    197   182.7967
280  100000    36.47227   48.58113    198   183.1032
281  100000    36.67950   47.64688    197   183.3958
282  100000    36.91066   48.58113    198   183.7212
283  100000    37.09184   48.58113    198   183.9756
284  100000    37.30169   48.58113    198   184.2673
285  100000    37.51438   48.58113    198   184.5626
286  100000    37.70065   48.58113    198   184.8191
287  100000    37.91729   48.58113    198   185.1170
288  100000    38.12084   48.58113    198   185.3932
289  100000    38.32881   49.53370    199   185.6761
290  100000    38.52612   48.58113    198   185.9423
291  100000    38.70641   48.58113    198   186.1855
292  100000    38.89244   48.58113    198   186.4338
293  100000    39.05505   48.58113    198   186.6502
294  100000    39.22809   49.53370    199   186.8793
295  100000    39.40453   50.50495    200   187.1125
296  100000    39.57451   49.53370    199   187.3359
297  100000    39.75346   50.50495    200   187.5688
298  100000    39.92514   50.50495    200   187.7926
299  100000    40.10330   49.53370    199   188.0233
300  100000    40.27504   49.53370    199   188.2447
301  100000    40.43408   50.50495    200   188.4497
302  100000    40.60130   50.50495    200   188.6624
303  100000    40.75942   50.50495    200   188.8658
304  100000    40.90842   49.53370    199   189.0546
305  100000    41.05251   50.50495    200   189.2370
306  100000    41.20417   50.50495    200   189.4281
307  100000    41.35894   50.50495    200   189.6230
308  100000    41.52245   50.50495    200   189.8279
309  100000    41.66645   50.50495    200   190.0070
310  100000    41.81998   50.50495    200   190.1974
311  100000    41.96466   50.50495    200   190.3766
312  100000    42.10542   50.50495    200   190.5508
313  100000    42.24533   50.50495    200   190.7229
314  100000    42.39024   50.50495    200   190.9012
315  100000    42.51312   50.50495    200   191.0518
316  100000    42.64348   50.50495    200   191.2106
317  100000    42.76620   50.50495    200   191.3602
318  100000    42.89272   50.50495    200   191.5128
319  100000    43.02832   50.50495    200   191.6763
320  100000    43.15766   50.50495    200   191.8320
321  100000    43.29709   50.50495    200   192.0000
322  100000    43.42473   50.50495    200   192.1528
323  100000    43.55178   50.50495    200   192.3043
324  100000    43.67545   50.50495    200   192.4517
325  100000    43.79288   50.50495    200   192.5915
326  100000    43.91936   50.50495    200   192.7416
327  100000    44.03255   50.50495    200   192.8749
328  100000    44.15872   50.50495    200   193.0228
329  100000    44.26978   50.50495    200   193.1535
330  100000    44.37679   50.50495    200   193.2786
331  100000    44.49116   50.50495    200   193.4121
332  100000    44.59750   50.50495    200   193.5358
333  100000    44.69374   50.50495    200   193.6479
334  100000    44.78816   50.50495    200   193.7579
335  100000    44.87982   50.50495    200   193.8638
336  100000    44.98306   50.50495    200   193.9829
337  100000    45.08574   50.50495    200   194.1010
338  100000    45.19003   50.50495    200   194.2211
339  100000    45.27181   50.50495    200   194.3151
340  100000    45.35848   50.50495    200   194.4148
341  100000    45.44625   50.50495    200   194.5151
342  100000    45.52920   50.50495    200   194.6099
343  100000    45.61179   50.50495    200   194.7039
344  100000    45.69713   50.50495    200   194.8009
345  100000    45.78385   50.50495    200   194.8995
346  100000    45.87089   50.50495    200   194.9980
347  100000    45.95729   50.50495    200   195.0960
348  100000    46.04397   50.50495    200   195.1935
349  100000    46.13035   50.50495    200   195.2912
350  100000    46.21428   50.50495    200   195.3856
351  100000    46.28920   50.50495    200   195.4694
352  100000    46.36925   50.50495    200   195.5593
353  100000    46.44839   50.50495    200   195.6480
354  100000    46.52142   50.50495    200   195.7293
355  100000    46.59677   50.50495    200   195.8136
356  100000    46.66183   50.50495    200   195.8861
357  100000    46.73970   50.50495    200   195.9728
358  100000    46.80522   50.50495    200   196.0457
359  100000    46.87380   50.50495    200   196.1217
360  100000    46.93079   50.50495    200   196.1851
361  100000    46.99577   50.50495    200   196.2569
362  100000    47.06106   50.50495    200   196.3290
363  100000    47.12749   50.50495    200   196.4022
364  100000    47.18848   50.50495    200   196.4692
365  100000    47.25384   50.50495    200   196.5411
366  100000    47.30929   50.50495    200   196.6018
367  100000    47.36016   50.50495    200   196.6578
368  100000    47.42483   50.50495    200   196.7287
369  100000    47.47914   50.50495    200   196.7885
370  100000    47.53172   50.50495    200   196.8459
371  100000    47.58315   50.50495    200   196.9019
372  100000    47.64122   50.50495    200   196.9655
373  100000    47.68602   50.50495    200   197.0143
374  100000    47.74129   50.50495    200   197.0744
375  100000    47.79470   50.50495    200   197.1329
376  100000    47.84800   50.50495    200   197.1907
377  100000    47.89934   50.50495    200   197.2466
378  100000    47.94293   50.50495    200   197.2938
379  100000    47.98863   50.50495    200   197.3432
380  100000    48.03492   50.50495    200   197.3933
381  100000    48.08258   50.50495    200   197.4449
382  100000    48.12765   50.50495    200   197.4937
383  100000    48.16421   50.50495    200   197.5331
384  100000    48.21208   50.50495    200   197.5848
385  100000    48.24821   50.50495    200   197.6237
386  100000    48.29195   50.50495    200   197.6706
387  100000    48.33405   50.50495    200   197.7158
388  100000    48.37344   50.50495    200   197.7584
389  100000    48.40743   50.50495    200   197.7951
390  100000    48.45051   50.50495    200   197.8410
391  100000    48.48874   50.50495    200   197.8822
392  100000    48.52821   50.50495    200   197.9245
393  100000    48.56356   50.50495    200   197.9623
394  100000    48.60043   50.50495    200   198.0018
395  100000    48.63962   50.50495    200   198.0436
396  100000    48.67291   50.50495    200   198.0791
397  100000    48.70353   50.50495    200   198.1119
398  100000    48.73686   50.50495    200   198.1475
399  100000    48.76806   50.50495    200   198.1809
400  100000    48.79897   50.50495    200   198.2138

Diploid Model with selection (random number based on
relative fitness to determine who survives).  Mutation rate is      1e-08,
Recomb rate is ONE per sexual individual per generation
Starting values at generation   0 are; population size 100000,
with avg fitness of    1.22125.

gen  popSize    avgFit     maxFit maxAlleles avgAlleles
  0  100000     1.22125    1.64061     25    10.0033
  1  100000     1.22125    1.64061     25    10.0033
  2  100000     1.22600    1.67085     26    10.1980
  3  100000     1.23123    1.67149     26    10.4115
  4  100000     1.23636    1.74035     28    10.6206
  5  100000     1.24118    1.70689     27    10.8153
  6  100000     1.24633    1.70426     27    11.0235
  7  100000     1.25143    1.77516     29    11.2282
  8  100000     1.25638    1.80927     30    11.4252
  9  100000     1.26142    1.77448     29    11.6255
 10  100000     1.26663    1.74102     28    11.8328
 11  100000     1.27237    1.77380     29    12.0588
 12  100000     1.27813    1.77584     29    12.2847
 13  100000     1.28347    1.77584     29    12.4938
 14  100000     1.28926    1.80997     30    12.7208
 15  100000     1.29538    1.81067     30    12.9579
 16  100000     1.30138    1.84475     31    13.1894
 17  100000     1.30743    1.84688     31    13.4228
 18  100000     1.31367    1.84617     31    13.6595
 19  100000     1.32038    1.88164     32    13.9146
 20  100000     1.32753    1.84546     31    14.1853
 21  100000     1.33459    1.95766     34    14.4530
 22  100000     1.34170    1.88382     32    14.7180
 23  100000     1.34912    1.95992     34    14.9955
 24  100000     1.35634    1.92075     33    15.2631
 25  100000     1.36375    1.88309     32    15.5364
 26  100000     1.37152    1.92223     33    15.8218
 27  100000     1.37966    1.95917     34    16.1181
 28  100000     1.38815    2.12148     38    16.4246
 29  100000     1.39672    2.12067     38    16.7332
 30  100000     1.40470    2.16142     39    17.0195
 31  100000     1.41415    1.99989     35    17.3582
 32  100000     1.42340    2.07989     37    17.6851
 33  100000     1.43307    2.03832     36    18.0250
 34  100000     1.44320    2.07749     37    18.3797
 35  100000     1.45251    2.16142     39    18.7025
 36  100000     1.46263    2.11985     38    19.0504
 37  100000     1.47297    2.16308     39    19.4040
 38  100000     1.48287    2.11985     38    19.7404
 39  100000     1.49173    2.16142     39    20.0392
 40  100000     1.50319    2.24787     41    20.4237
 41  100000     1.51388    2.34049     43    20.7799
 42  100000     1.52423    2.29195     42    21.1226
 43  100000     1.53503    2.24960     41    21.4785
 44  100000     1.54653    2.25047     41    21.8535
 45  100000     1.55938    2.29371     42    22.2698
 46  100000     1.57066    2.38730     44    22.6315
 47  100000     1.58312    2.38730     44    23.0290
 48  100000     1.59522    2.38271     44    23.4129
 49  100000     1.60805    2.47993     46    23.8146
 50  100000     1.62161    2.43130     45    24.2351
 51  100000     1.63522    2.43317     45    24.6549
 52  100000     1.64838    2.43598     45    25.0590
 53  100000     1.66232    2.53147     47    25.4833
 54  100000     1.67750    2.53245     47    25.9381
 55  100000     1.69200    2.58111     48    26.3726
 56  100000     1.70702    2.68642     50    26.8149
 57  100000     1.72257    2.63476     49    27.2687
 58  100000     1.73933    2.63476     49    27.7550
 59  100000     1.75554    2.85194     53    28.2226
 60  100000     1.77296    2.73699     51    28.7192
 61  100000     1.79024    2.96374     55    29.2061
 62  100000     1.80744    2.79280     52    29.6870
 63  100000     1.82481    2.84975     53    30.1669
 64  100000     1.84340    2.79387     52    30.6781
 65  100000     1.86255    2.96716     55    31.1989
 66  100000     1.88206    3.14394     58    31.7215
 67  100000     1.90115    2.96488     55    32.2312
 68  100000     1.92197    3.08585     57    32.7765
 69  100000     1.94310    3.08229     57    33.3274
 70  100000     1.96499    3.08585     57    33.8899
 71  100000     1.98646    3.14636     58    34.4362
 72  100000     2.00773    3.75441     67    34.9738
 73  100000     2.03107    3.20558     59    35.5559
 74  100000     2.05344    3.39918     62    36.1093
 75  100000     2.07743    3.33509     61    36.6938
 76  100000     2.10113    3.53786     64    37.2647
 77  100000     2.12605    3.47250     63    37.8596
 78  100000     2.15064    3.46983     63    38.4391
 79  100000     2.17624    3.60307     65    39.0368
 80  100000     2.20226    3.67513     66    39.6341
 81  100000     2.22953    3.60862     65    40.2545
 82  100000     2.25714    3.83539     68    40.8734
 83  100000     2.28665    4.05921     71    41.5280
 84  100000     2.31508    4.30435     74    42.1524
 85  100000     2.34305    4.14039     72    42.7602
 86  100000     2.37520    3.98574     70    43.4451
 87  100000     2.40580    4.22320     73    44.0880
 88  100000     2.43814    4.13721     72    44.7649
 89  100000     2.47078    4.06545     71    45.4337
 90  100000     2.50545    4.39551     75    46.1356
 91  100000     2.54072    4.31098     74    46.8419
 92  100000     2.57805    4.38875     75    47.5783
 93  100000     2.61713    4.30932     74    48.3374
 94  100000     2.65430    4.39044     75    49.0493
 95  100000     2.69136    4.75235     79    49.7492
 96  100000     2.73207    4.56430     77    50.5097
 97  100000     2.77168    4.75418     79    51.2332
 98  100000     2.81480    5.13817     83    52.0125
 99  100000     2.85958    5.14608     83    52.8091
100  100000     2.90443    4.94244     81    53.5971
101  100000     2.94934    4.94244     81    54.3716
102  100000     2.99653    5.24496     84    55.1768
103  100000     3.04362    5.44847     86    55.9624
104  100000     3.09279    5.24900     84    56.7751
105  100000     3.14296    5.76864     89    57.5902
106  100000     3.19536    5.55744     87    58.4285
107  100000     3.24870    5.55958     87    59.2638
108  100000     3.30027    6.02018     91    60.0657
109  100000     3.35262    5.55958     87    60.8624
110  100000     3.40931    6.12879     92    61.7051
111  100000     3.46802    6.01555     91    62.5709
112  100000     3.52499    5.90441     90    63.3956
113  100000     3.58405    6.51143     95    64.2364
114  100000     3.64550    6.50142     95    65.0998
115  100000     3.70835    6.37639     94    65.9678
116  100000     3.77205    6.51644     95    66.8295
117  100000     3.84059    6.76148     97    67.7427
118  100000     3.90902    7.04005     99    68.6338
119  100000     3.97725    7.03735     99    69.5126
120  100000     4.04753    6.90998     98    70.4044
121  100000     4.12218    7.46809    102    71.3284
122  100000     4.20069    7.48246    102    72.2908
123  100000     4.27882    7.75786    104    73.2242
124  100000     4.35317    7.60574    103    74.0946
125  100000     4.43036    8.06817    106    74.9869
126  100000     4.51028    8.39412    108    75.8946
127  100000     4.59640    8.40381    108    76.8501
128  100000     4.68282    8.40058    108    77.8038
129  100000     4.76969    8.72318    110    78.7356
130  100000     4.85920    8.38445    108    79.6814
131  100000     4.94729    9.07211    112    80.5929
132  100000     5.04348    9.07559    112    81.5680
133  100000     5.13873    9.62739    115    82.5196
134  100000     5.23276    9.82749    116    83.4484
135  100000     5.32718    9.61999    115    84.3562
136  100000     5.42362    9.81994    116    85.2705
137  100000     5.52186   10.02790    117    86.1817
138  100000     5.62528    9.62369    115    87.1234
139  100000     5.73261    9.80485    116    88.0910
140  100000     5.83035   10.41699    119    88.9514
141  100000     5.94080   10.60900    120    89.9110
142  100000     6.05513   11.46587    124    90.8851
143  100000     6.16825   10.60900    120    91.8256
144  100000     6.28462   11.46587    124    92.7796
145  100000     6.40142   11.69519    125    93.7134
146  100000     6.52062   11.04610    122    94.6588
147  100000     6.64462   12.17704    127    95.6251
148  100000     6.76499   12.16300    127    96.5350
149  100000     6.89095   11.96125    126    97.4764
150  100000     7.01386   12.18172    127    98.3802
151  100000     7.15013   12.15832    127    99.3615
152  100000     7.28209   12.66412    129   100.2952
153  100000     7.42020   13.42377    132   101.2553
154  100000     7.55813   12.89755    130   102.1970
155  100000     7.70271   12.91244    130   103.1626
156  100000     7.84075   13.97684    134   104.0667
157  100000     7.98649   15.11153    138   105.0043
158  100000     8.14176   14.22899    135   105.9881
159  100000     8.28686   14.80953    137   106.8912
160  100000     8.43813   14.81523    137   107.8161
161  100000     8.58774   15.09411    138   108.7151
162  100000     8.74414   15.68581    140   109.6383
163  100000     8.90266   16.33207    142   110.5570
164  100000     9.06278   15.37233    139   111.4655
165  100000     9.23295   16.62671    143   112.4146
166  100000     9.40388   16.67153    143   113.3524
167  100000     9.56790   16.01799    141   114.2360
168  100000     9.73507   17.99037    147   115.1208
169  100000     9.90915   17.31840    145   116.0272
170  100000    10.08411   18.32197    148   116.9267
171  100000    10.25545   17.30508    145   117.7889
172  100000    10.44079   19.09152    150   118.7070
173  100000    10.62134   18.74599    149   119.5843
174  100000    10.80924   18.36429    148   120.4832
175  100000    10.99674   19.08418    150   121.3640
176  100000    11.18997   19.47335    151   122.2585
177  100000    11.38300   19.83229    152   123.1332
178  100000    11.58549   19.06951    150   124.0351
179  100000    11.77824   21.04618    155   124.8802
180  100000    11.96681   20.60972    154   125.6911
181  100000    12.16521   21.78726    157   126.5375
182  100000    12.36454   21.05428    155   127.3685
183  100000    12.57409   22.72857    159   128.2298
184  100000    12.78705   22.31721    158   129.0882
185  100000    12.99919   21.87962    157   129.9346
186  100000    13.20818   21.47536    156   130.7539
187  100000    13.43192   21.88803    157   131.6137
188  100000    13.64318   22.75480    159   132.4138
189  100000    13.86418   23.66499    161   133.2313
190  100000    14.09444   24.09193    162   134.0792
191  100000    14.33061   24.11974    162   134.9292
192  100000    14.57266   25.08453    164   135.7901
193  100000    14.79218   24.58322    163   136.5571
194  100000    15.01892   24.60213    163   137.3397
195  100000    15.25529   25.59606    165   138.1411
196  100000    15.49514   26.08791    166   138.9419
197  100000    15.73848   28.19499    170   139.7406
198  100000    15.97978   28.74783    171   140.5224
199  100000    16.21090   26.57898    167   141.2645
200  100000    16.44976   27.12099    168   142.0162
201  100000    16.70349   27.64214    169   142.8041
202  100000    16.95579   28.19499    170   143.5736
203  100000    17.20622   28.17331    170   144.3295
204  100000    17.46017   28.18415    170   145.0825
205  100000    17.72264   27.65278    169   145.8488
206  100000    17.97248   28.17331    170   146.5672
207  100000    18.21658   30.44883    174   147.2625
208  100000    18.47983   29.32279    172   147.9980
209  100000    18.75339   32.92079    178   148.7552
210  100000    19.01690   33.57920    179   149.4752
211  100000    19.27860   31.66678    176   150.1751
212  100000    19.53827   31.69114    176   150.8626
213  100000    19.79308   31.06975    175   151.5321
214  100000    20.05034   31.66678    176   152.1968
215  100000    20.32730   32.94612    178   152.9052
216  100000    20.60468   32.31254    177   153.6020
217  100000    20.88142   32.93345    178   154.2882
218  100000    21.14348   34.22446    180   154.9333
219  100000    21.39614   34.25079    180   155.5439
220  100000    21.68261   32.92079    178   156.2301
221  100000    21.97036   33.57920    179   156.9071
222  100000    22.23777   37.00294    184   157.5301
223  100000    22.50878   36.26345    183   158.1542
224  100000    22.77138   35.57975    182   158.7503
225  100000    23.04941   34.92237    181   159.3791
226  100000    23.34112   38.46826    186   160.0299
227  100000    23.60835   35.60713    182   160.6185
228  100000    23.88325   35.60713    182   161.2132
229  100000    24.16438   37.00294    184   161.8178
230  100000    24.44150   36.29135    183   162.4069
231  100000    24.72809   40.79144    189   163.0076
232  100000    24.99645   37.74300    185   163.5660
233  100000    25.28134   36.30530    183   164.1470
234  100000    25.56341   38.48306    186   164.7202
235  100000    25.83514   38.48306    186   165.2674
236  100000    26.11645   39.22254    187   165.8273
237  100000    26.38544   39.22254    187   166.3571
238  100000    26.64969   41.57529    190   166.8684
239  100000    26.91462   39.99161    188   167.3818
240  100000    27.19764   40.00699    188   167.9218
241  100000    27.46762   38.48306    186   168.4310
242  100000    27.73681   39.23763    187   168.9342
243  100000    28.00335   40.77576    189   169.4279
244  100000    28.27045   40.79144    189   169.9182
245  100000    28.53322   40.00699    188   170.3977
246  100000    28.78766   42.39049    191   170.8578
247  100000    29.03530   40.79144    189   171.3041
248  100000    29.29594   43.23830    192   171.7654
249  100000    29.54209   42.40679    191   172.1967
250  100000    29.79926   43.23830    192   172.6442
251  100000    30.06558   43.23830    192   173.1029
252  100000    30.33229   43.23830    192   173.5599
253  100000    30.56019   43.23830    192   173.9476
254  100000    30.82756   42.40679    191   174.3977
255  100000    31.08140   42.40679    191   174.8214
256  100000    31.34374   43.22167    192   175.2572
257  100000    31.59680   44.95054    194   175.6737
258  100000    31.85299   45.83192    195   176.0933
259  100000    32.08503   44.95054    194   176.4677
260  100000    32.35316   44.08611    193   176.8976
261  100000    32.58760   44.95054    194   177.2707
262  100000    32.83905   44.95054    194   177.6699
263  100000    33.08487   44.95054    194   178.0560
264  100000    33.33539   46.73059    196   178.4470
265  100000    33.59346   46.73059    196   178.8478
266  100000    33.83818   46.73059    196   179.2235
267  100000    34.06584   46.73059    196   179.5700
268  100000    34.30304   46.73059    196   179.9287
269  100000    34.52085   46.71262    196   180.2567
270  100000    34.77190   45.83192    195   180.6307
271  100000    35.00863   46.73059    196   180.9822
272  100000    35.23078   47.64688    197   181.3103
273  100000    35.46491   47.64688    197   181.6536
274  100000    35.67022   47.64688    197   181.9533
275  100000    35.88968   48.58113    198   182.2711
276  100000    36.11046   47.64688    197   182.5891
277  100000    36.29918   46.73059    196   182.8593
278  100000    36.50986   48.58113    198   183.1576
279  100000    36.71630   48.58113    198   183.4498
280  100000    36.92845   48.58113    198   183.7483
281  100000    37.14335   47.64688    197   184.0485
282  100000    37.34642   48.58113    198   184.3323
283  100000    37.53838   49.53370    199   184.5967
284  100000    37.74236   48.58113    198   184.8778
285  100000    37.93692   48.58113    198   185.1440
286  100000    38.14397   49.53370    199   185.4259
287  100000    38.35483   49.53370    199   185.7124
288  100000    38.54740   48.58113    198   185.9715
289  100000    38.72207   50.50495    200   186.2064
290  100000    38.90748   49.53370    199   186.4535
291  100000    39.07862   50.50495    200   186.6801
292  100000    39.25898   49.53370    199   186.9190
293  100000    39.43116   49.53370    199   187.1451
294  100000    39.59424   49.53370    199   187.3604
295  100000    39.77005   50.50495    200   187.5897
296  100000    39.93488   50.50495    200   187.8038
297  100000    40.10349   50.50495    200   188.0235
298  100000    40.27283   50.50495    200   188.2409
299  100000    40.43720   49.53370    199   188.4519
300  100000    40.59668   50.50495    200   188.6568
301  100000    40.76119   50.50495    200   188.8671
302  100000    40.91626   50.50495    200   189.0644
303  100000    41.06473   50.50495    200   189.2526
304  100000    41.21201   50.50495    200   189.4379
305  100000    41.35677   50.50495    200   189.6198
306  100000    41.51658   50.50495    200   189.8197
307  100000    41.67751   50.50495    200   190.0204
308  100000    41.81564   50.50495    200   190.1921
309  100000    41.94687   50.50495    200   190.3545
310  100000    42.08066   50.50495    200   190.5197
311  100000    42.21416   50.50495    200   190.6853
312  100000    42.35605   50.50495    200   190.8585
313  100000    42.49196   50.50495    200   191.0243
314  100000    42.61648   50.50495    200   191.1762
315  100000    42.75424   50.50495    200   191.3440
316  100000    42.89086   50.50495    200   191.5096
317  100000    43.02039   50.50495    200   191.6666
318  100000    43.14558   50.50495    200   191.8176
319  100000    43.27909   50.50495    200   191.9779
320  100000    43.40510   50.50495    200   192.1290
321  100000    43.52068   50.50495    200   192.2673
322  100000    43.63836   50.50495    200   192.4071
323  100000    43.75628   50.50495    200   192.5470
324  100000    43.86790   50.50495    200   192.6792
325  100000    43.98010   50.50495    200   192.8118
326  100000    44.08201   50.50495    200   192.9322
327  100000    44.19513   50.50495    200   193.0650
328  100000    44.31482   50.50495    200   193.2048
329  100000    44.42538   50.50495    200   193.3348
330  100000    44.53484   50.50495    200   193.4625
331  100000    44.63092   50.50495    200   193.5747
332  100000    44.72430   50.50495    200   193.6836
333  100000    44.82027   50.50495    200   193.7950
334  100000    44.92324   50.50495    200   193.9138
335  100000    45.00984   50.50495    200   194.0138
336  100000    45.10500   50.50495    200   194.1232
337  100000    45.20520   50.50495    200   194.2390
338  100000    45.30144   50.50495    200   194.3495
339  100000    45.40048   50.50495    200   194.4632
340  100000    45.48739   50.50495    200   194.5627
341  100000    45.57137   50.50495    200   194.6584
342  100000    45.67343   50.50495    200   194.7744
343  100000    45.76278   50.50495    200   194.8758
344  100000    45.84334   50.50495    200   194.9672
345  100000    45.92185   50.50495    200   195.0561
346  100000    45.99799   50.50495    200   195.1421
347  100000    46.08236   50.50495    200   195.2376
348  100000    46.15469   50.50495    200   195.3189
349  100000    46.24516   50.50495    200   195.4202
350  100000    46.32646   50.50495    200   195.5111
351  100000    46.39721   50.50495    200   195.5907
352  100000    46.46720   50.50495    200   195.6694
353  100000    46.54914   50.50495    200   195.7605
354  100000    46.62813   50.50495    200   195.8486
355  100000    46.69471   50.50495    200   195.9230
356  100000    46.76374   50.50495    200   195.9999
357  100000    46.81820   50.50495    200   196.0604
358  100000    46.88529   50.50495    200   196.1348
359  100000    46.94019   50.50495    200   196.1953
360  100000    47.00622   50.50495    200   196.2682
361  100000    47.06942   50.50495    200   196.3380
362  100000    47.12403   50.50495    200   196.3986
363  100000    47.18719   50.50495    200   196.4681
364  100000    47.24923   50.50495    200   196.5364
365  100000    47.30182   50.50495    200   196.5943
366  100000    47.35333   50.50495    200   196.6507
367  100000    47.41490   50.50495    200   196.7183
368  100000    47.46475   50.50495    200   196.7729
369  100000    47.52624   50.50495    200   196.8403
370  100000    47.58248   50.50495    200   196.9016
371  100000    47.63379   50.50495    200   196.9576
372  100000    47.69130   50.50495    200   197.0202
373  100000    47.74508   50.50495    200   197.0787
374  100000    47.79988   50.50495    200   197.1385
375  100000    47.84735   50.50495    200   197.1901
376  100000    47.88775   50.50495    200   197.2338
377  100000    47.93676   50.50495    200   197.2869
378  100000    47.98808   50.50495    200   197.3425
379  100000    48.02841   50.50495    200   197.3864
380  100000    48.07850   50.50495    200   197.4407
381  100000    48.11946   50.50495    200   197.4848
382  100000    48.16069   50.50495    200   197.5292
383  100000    48.19759   50.50495    200   197.5689
384  100000    48.24420   50.50495    200   197.6191
385  100000    48.27791   50.50495    200   197.6556
386  100000    48.31467   50.50495    200   197.6952
387  100000    48.35524   50.50495    200   197.7389
388  100000    48.38969   50.50495    200   197.7759
389  100000    48.43213   50.50495    200   197.8214
390  100000    48.46609   50.50495    200   197.8578
391  100000    48.50128   50.50495    200   197.8955
392  100000    48.53709   50.50495    200   197.9337
393  100000    48.57215   50.50495    200   197.9713
394  100000    48.61414   50.50495    200   198.0164
395  100000    48.65069   50.50495    200   198.0555
396  100000    48.68113   50.50495    200   198.0879
397  100000    48.71910   50.50495    200   198.1283
398  100000    48.75327   50.50495    200   198.1647
399  100000    48.78807   50.50495    200   198.2018
400  100000    48.82191   50.50495    200   198.2379
```
